# Comparative analysis of retroviral Gag-host cell interactions: focus on the nuclear interactome

**DOI:** 10.1101/2024.01.18.575255

**Authors:** Gregory S. Lambert, Breanna L. Rice, Rebecca J. Kaddis Maldonado, Jordan Chang, Leslie J. Parent

## Abstract

Retroviruses exploit a variety of host proteins to assemble and release virions from infected cells. To date, most studies that examined possible interacting partners of retroviral Gag proteins focused on host proteins that localize primarily to the cytoplasm or plasma membrane. Given the recent findings that several full-length Gag proteins localize to the nucleus, identifying the Gag-nuclear interactome has high potential for novel findings that reveal previously unknown host processes. In this study, we systematically compared nuclear factors identified in published HIV-1 proteomic studies which had used a variety of experimental approaches. In addition, to contribute to this body of knowledge, we report results from a mass spectrometry approach using affinity-tagged (His_6_) HIV-1 and RSV Gag proteins mixed with nuclear extracts. Taken together, the previous studies—as well as our own—identified potential binding partners of HIV-1 and RSV Gag involved in several nuclear processes, including transcription, splicing, RNA modification, and chromatin remodeling. Although a subset of host proteins interacted with both Gag proteins, there were also unique host proteins belonging to each interactome dataset. To validate one of the novel findings, we demonstrated the interaction of RSV Gag with a member of the Mediator complex, Med26, which is required for RNA polymerase II-mediated transcription. These results provide a strong premise for future functional studies to investigate roles for these nuclear host factors that may have shared functions in the biology of both retroviruses, as well as functions specific to RSV and HIV-1, given their distinctive hosts and molecular pathology.

## 1. Introduction

Retroviral replication depends upon the selection and packaging of unspliced viral RNA (USvRNA) as the genome by the Gag polyprotein, the major structural protein shared by retroviruses. Historically, it was thought that the Gag protein performed this function in the cytoplasm before trafficking to the plasma membrane where budding of virions occurs. However, there is now compelling evidence that a population of retroviral Gag proteins enter the nucleus where they may initiate selection of genomic RNA (gRNA). The strongest evidence for this paradigm shift is based on studies of the Rous sarcoma virus (RSV) Gag protein, which depends on transient nucleocytoplasmic trafficking facilitated by host transport proteins to ensure efficient gRNA packaging (1–6). This requirement for nuclear trafficking was demonstrated by a mutant of Gag that bypassed the nucleus, resulting in decreased gRNA packaging which was increased with restoration of nuclear localization (7). RSV Gag co-opts the host karyopherins importin α/β, importin-11, and transportin-3 (TNPO3) to enter the nucleus (1–3, 6), and nuclear egress is mediated by binding of a hydrophobic nuclear export signal (NES) in the p10 domain to the CRM1-RanGTP nuclear export complex (3–5). A more recent study demonstrated that RSV Gag binds to newly synthesized USvRNA in discrete ribonucleoprotein complexes (RNPs) in the nucleus, and these Gag-vRNA RNPs have been observed trafficking across the nuclear envelope into the cytoplasm (8).

A population of human immunodeficiency virus type-1 (HIV-1) Gag, like that of RSV Gag, also undergoes nuclear localization (9–13). HIV-1 Gag forms focal RNP complexes with nascent USvRNA in the nucleus, and traffics to the major viral RNA transcription site in T cells reactivated from latency (9). Nuclear localization of HIV-1 Gag occurs in a concentra-tion-independent manner shortly after Gag synthesis begins, and Gag colocalizes with transcriptionally-active euchromatin near the nuclear periphery (10). The function(s) of these nuclear RNPs has yet to be thoroughly investigated, although these data demonstrate that both RSV and HIV-1 Gag proteins traffic to transcription sites and associate with their cognate USvRNAs. Although one possibility is that Gag-USvRNA binding in the nucleus could initiate the genomic RNA packaging process, nuclear localization of Gag could also influence other important cellular processes, such as regulation of viral or host transcription, RNA modification or processing, splicing, chromatin remodeling, or RNA export.

In addition to RSV and HIV-1, the Gag proteins of other retroviruses, including murine leukemia virus (MLV), prototype foamy virus (PFV), feline immunodeficiency virus (FIV), mouse mammary tumor virus (MMTV), and Mason-Pfizer monkey virus (MPMV) also have been shown to localize to the nucleus (11, 14–27). For example, the PFV Gag protein is involved in proviral integration through its interaction with chromatin (27). The finding that nuclear trafficking of Gag is a feature conserved among many retroviruses raises the likelihood that nuclear-localized Gag proteins participate in functions important for virus replication. There are also Gag cleavage products that undergo nuclear localization, including the nucleocapsid (NC) proteins of HIV-1, RSV, MLV, and MMTV, which localize to the nucleolus (11, 19, 22, 28), and the p12 protein of MLV, which binds to chromatin and influences proviral integration (23, 29).

It is well known that retroviruses exploit a variety of host pathways during replication, but previous investigation of host factors that bind to Gag have focused on factors localized to the cytoplasm and plasma membrane. However, the nuclear localization of retroviral Gag proteins raises important questions concerning their functions, which can be informed by identifying nuclear host partners. To gain further insight into what nuclear processes Gag could be influencing, we comprehensively analyzed and systematically compared six previously published HIV-1 proteomic studies performed by other laboratories, which used various experimental approaches to identify novel host proteins that interact with HIV-1 Gag. A variety of techniques are represented in this analysis, including affinity purifications of GFP-tagged Gag, tandem affinity purification of Gag, and BirA* Gag complexes (30–35). To complement those datasets, we performed affinity-tagged purification of both RSV and HIV-1 Gag, and identified nuclear interacting partners using mass spectrometry. To further explore one of the novel hits, we utilized immunoprecipitation and quantitative imaging approaches to validate the interaction of RSV Gag with Mediator complex subunit 26 (Med26), a critical component of the transcriptional Mediator complex, which is exploited by other viruses and endogenous retroelements (36–43). Together, published studies combined with our results suggest that Gag proteins may interface with host nuclear factors to facilitate genomic RNA selection and/or influence cellular processes, including gene expression, RNA processing, splicing, nucleic acid metabolism, and/or chromatin modification.

## 2. Materials and Methods

### Cells, Plasmids, and Purified Proteins

DF1 chicken embryo fibroblast cells, HeLa human cervical cancer cells, and QT6 quail fibroblast cells were maintained as described (11, 44, 45). The RSV Gag expression constructs pGag.ΔPR (referred to herein as RSV Gag), pGag.L219A.ΔPR (referred to herein as RSV Gag.L219A), pGag.ΔNC, and pGag.ΔPR-GFP (referred to herein as RSV Gag-GFP) (4, 11, 46) and plasmids encoding for *Escherichia coli* (*E. coli*) expression of His-tagged RSV Gag (pET28.TEV-Gag.3h) and HIV Gag (pET28a.WT.HIV.Gag.Δp6) were previously described (47, 48).

### Subcellular Fractionation

QT6 cells were transfected with untagged RSV Gag constructs using the calcium phosphate method (49). Sixteen hours later, the medium was changed to fresh primary growth medium (PGM) and the cells were allowed to recover for 24 hours. All subsequent steps were performed on ice or at 4°C with cold buffers unless otherwise stated. Cells were fractionated using the method described in (50) with some minor modifications, as below. Cells were removed from the plates using trypsin and then washed in cold PBS. The cell pellet was resuspended in sucrose buffer (10 mM HEPES pH 7.9, 10 mM KCl, 2 mM magnesium acetate, 3 mM CaCl_2,_ 340 mM sucrose, 1 mM DTT, 100 μg/ml phenylmethanesulfonyl fluoride (PMSF), 1 μg/ml pepstatin, and Roche Complete Protease Inhibitor Cocktail) and incubated on ice for 10 minutes. IGEPAL Nonidet P-40 was added to the final concentration of 0.5% and cells were vortexed on high for 15 seconds, and then spun for 10 minutes at 3,500 x g at 4°C. The supernatant (cytoplasm fraction) was collected, and the pelleted nuclei were resuspended in nucleoplasm extraction buffer (50 mM HEPES pH 7.9, 150 mM potassium acetate, 1.5 mM MgCl_2,_ 0.1% IGEPAL Nonidet P-40, 1 mM DTT, 100 μg/ml PMSF, 1 μg/ml pepstatin, and Roche Complete Protease Inhibitor Cocktail) and transferred to a Dounce homogenizer and homogenized with 20 slow strokes. The homogenates were checked under a light microscope for completion of nuclear lysis, then transferred to a new tube and rotated at 4°C for 20 minutes. The lysates were spun at 16,000 x *g* for 10 minutes at 4°C. The supernatant (nucleoplasmic fraction) was collected and the remaining chromatin-containing pellet was resuspended in nuclease incubation buffer (50 mM HEPES pH 7.9, 10 mM NaCl, 1.5 mM MgCl_2_, 1 mM DTT, 100 μg/ml PMSF, 1 μg/ml pepstatin, and Roche Complete Protease Inhibitor Cocktail) with 100 U/ml of OmniCleave nuclease (Epicentre) for 10 minutes at 37°C. NaCl was added to a final concentration of 150 mM and the lysates were incubated on ice for 20 minutes and spun for 10 minutes at 16,000 x *g* at 4°C. The supernatant (low-salt chromatin fraction) was collected and the pellet was resuspended in chromatin extraction buffer (50 mM HEPES pH 7.9, 500 mM NaCl, 1.5 mM MgCl_2_, 0.1% Triton X-100, 1 mM DTT, 100 μg/ml PMSF, 1 μg/ml pepstatin, and Roche Complete Protease Inhibitor Cocktail), incubated for 20 minutes on ice, spun for 10 minutes at 16,000 x g at 4°C, and the supernatant (high-salt chromatin fraction) was collected.

### Western Blot Analysis of Subcellular Fractions

Proteins from the subcellular fractions were analyzed via SDS-PAGE. Aliquots of the fractions were heated to 90°C in 4X SDS-PAGE sample buffer (250 mM Tris-HCl, pH 6.8, 40% glycerol, 0.4% bromophenol blue, 8% SDS, and 8% β-mercaptoethanol) for 10 minutes prior to loading on a 10% SDS-PAGE gel and analyzed by western blot. Proteins were detected using antibodies against RSV Gag (51), Calnexin (Enzo Life Sciences ADI-SPA-865), Med4 (Abcam ab129170), RCC1 (Abcam ab54600), Histone H2B (Abcam ab52484), GAPDH (UBP Bio Y1040), and the appropriate HRP-conjugated secondary antibodies (Invitrogen).

Signal densities of the protein bands on the antibody-stained membranes were analyzed using Bio-Rad Image Lab Software on a ChemiDoc MP system. Rectangles were drawn around each band, as well as a blank background region, using the volume tools feature to quantify the signal intensity of each band. The background subtraction method was set to local, and the blank region that was highlighted by a rectangle was labelled as the background volume. The volumes report table was exported to Microsoft Excel. For each band corresponding to the Gag signal, the adjusted volumes for each fraction were added together to calculate the total adjusted volume. Then the percentages of each fraction were calculated by subtracting the fraction’s adjusted volume from the total adjusted volume. Averages and standard deviations were calculated for each fraction for each Gag protein from three separate experiments.

### Purified RSV Gag and HIV Gag Pulldowns

Lysate Preparation: DF1 and HeLa cells were fractionated using the NE-PER Nuclear and Cytoplasmic Extraction kit (ThermoFisher Scientific). All steps and buffers used were performed on ice or at 4°C unless otherwise stated. Cells were lysed in CERI buffer containing the Complete Protease Inhibitor Cocktail (Roche). Cells were vortexed on the highest setting for 15 seconds and incubated on ice for 10 minutes. Ice-cold CERII buffer was added and cells were vortexed on high for 5 seconds then centrifuged for 5 minutes at 16,000 x *g* in a microcentrifuge. The supernatant was collected (cytoplasmic fraction), and the pelleted nuclei were resuspended in ice-cold NER buffer with protease inhibitor cocktail added. The nuclei were vortexed on high for 15 seconds and incubated on ice for 10 minutes, then vortexed for 15 seconds every 10 minutes for a total of 40 minutes. The lysed nuclei were centrifuged at 16,000 x *g* for 10 minutes. The supernatant (nuclear fractionation) was diluted to 14 ml with Buffer A (25 mM Tris-HCl pH 8.0, 200 mM NaCl, 2 mM 2-Mercaptoethanol (BME), and protease inhibitor cocktail). The nuclear fraction was concentrated to ∼1 ml in a 3 kD MWCO Amicon column and then was diluted to 14 ml and concentrated once more to ∼1.2 ml.

Nickel Affinity Purifications: Three reactions were performed using 6 μg of RSV H6.Gag.3h or HIV-1 WT.Gag.Δp6.H6, and a no protein control for DF1 and HeLa nuclear lysates, respectively. The proteins and no protein control were incubated with pre-washed nickel beads for 1 hour at 4°C with rotation. The beads were then washed three times in Wash Buffer (300 mM NaCl, 50 mM NaH_2_PO_4_, pH 8.0), followed by incubation with 500 μg of nuclear extract for 2 hours at 4°C with rotation. The beads were washed again three times in Wash Buffer, and bound proteins were eluted from the beads using Wash Buffer + 300 mM imidazole for 15 minutes while rotating at 4°C. The eluates were buffer exchanged into water using Zeba Spin Desalting Columns (ThermoFisher Scientific) and 20 ug of each sample was used for mass spectrometry analysis.

Sample Preparation for Mass Spectrometry: The samples were prepared and processed at the Mass Spectrometry and Proteomics Core Research Facility at Penn State College of Medicine using an ABSciex 5600 TripleTOF. In a final volume of 100 μl, the samples were incubated in 50 mM NH_4_HCO_3_, pH 8.0, 10% v/v acetronitrile, and 0.1 μg trypsin for at least 3 hours at 48°C. To evaporate off the NH_4_HCO_3_ and acetronitrile, samples were dried down using a SpeedVac, and then resuspended in 200 μl H_2_O with vortexing. The drying was repeated 3X total, but the final resuspension volume was 10 μl. To each sample, a 1/9^th^ volume of 1% formic acid was added.

### Mass Spectrometry

The following mass spectrometry workflows were performed two separate times and data from both instances were combined to create the set of interactors presented herein. 2D-LC Separations: SCX (strong cation-exchange) separations were performed on a passivated Waters 600E HPLC system, using a 4.6 X 250 mm PolySULFOETHYL Aspartamide column (PolyLC, Columbia, MD) at a flow rate of 1 ml/min. The gradient was 100% Buffer A (10 mM ammonium formate, pH 2.7, in 20% acetonitrile/80% water) (0-22 minutes following sample injection), 0%→40% Buffer B (666 mM ammonium formate, pH 2.7, in 20% acetonitrile/80% water) (16-48 min), 40%→100% Buffer B (48-49 min), isocratic 100% Buffer B (49-56 min), then at 56 min switched back to 100% Buffer A to re-equilibrate for the next injection. One milliliter fractions were collected and were dried down then resuspended in 9 µl of 2% (v/v) acetonitrile, 0.1% (v/v) formic acid, and were filtered prior to reverse phase C18 nanoflow-LC separation.

Mass Spectrometry Analysis: Each SCX fraction was analyzed following a calibration run using trypsin-digested β-Gal as a calibrant, then a blank run using the ABSciex 5600 TripleTOF. MS Spectra were then acquired from each sample using the newly updated default calibration, using a 60-minute gradient from an Eksigent NanoLC-Ultra-2D Plus and Eksigent cHiPLC Nanoflex through a 200 µm x 0.5 mm Chrom XP C18-CL 3 µm 120 Å Trap Column and elution through a 75 µm x 15 cm Chrom XP C18-CL 3 µm 120 Å Nano cHiPLC Column.

Protein Identification and Analysis: Protein identification and quantitation were performed using the Paragon algorithm as implemented in ProteinPilot 5.0 software (ProteinPilot 5.0, which contains the Paragon Algorithm 5.0.0.0, build 4632 from ABI/MDS-Sciex) (52). Spectra were searched against *Homo sapien* or *Gallus gallus* RefSeq subsets (plus 389 common contaminants) of the NCBInr database concatenated with a reversed “decoy” version of itself. For the ProteinPilot analyses, the preset Thorough Identification Search settings were used, and identifications needed to have a ProteinPilot Unused Score > 1.3 (>95% confidence interval) to be accepted. In addition, the only protein identifications (IDs) accepted were required to have a “Local False Discovery Rate” estimation of no higher than 5%, as calculated from the slope of the accumulated Decoy database hits by the PSPEP (<u>P</u>roteomics <u>S</u>ystem <u>P</u>erformance <u>E</u>valuation <u>P</u>ipeline) (53). Proteins that were labelled as contaminants or reversed were removed from the analysis. The mass spectrometry proteomics data have been deposited to the ProteomeXchange Consortium via the PRIDE (54) partner repository with the dataset identifier PXD048774.

### Analysis of Proteomics

The Database for Annotation, Visualization, and Integrated Discovery (DAVID, version 6.8) (55, 56) was used to assign each protein to its cellular compartment(s) and biological process categories. Proteins were organized by their gene name for entries into DAVID and the *Homo sapiens* species database was used. Data presented in the tables were generated using the Gene Ontology GOTERM_BP_ALL to categorize proteins by their biological function, and GOTERM_CC_ALL to first identify the proteins present in the nucleus. Categories with a p-value of ≤ 0.05, as determined by modified Fisher’s Exact Test, were considered statistically overrepresented, and any redundant categories (same p-value and proteins) were removed.

The Bioinformatics and Evolutionary Genomics online comparison tool was used to generate the Venn diagram (http://bioinformatics.psb.ugent.be/webtools/Venn/). Ingenuity Pathway Analysis (IPA) (QIAGEN Inc., https://www.qiagenbioinformatics.com/products/ingenuitypathway-analysis) was performed to categorize the functions of the identified proteins. Core Analysis was performed on the gene IDs that could be mapped by IPA, as some gene IDs were not recognized by IPA, on each separate proteomic list. Under the Core Analysis, Expression analysis was selected; direct and indirect relationships were examined. No endogenous chemicals were included in the analysis. The filters that were used included: all molecule types and data sources; confidence = experimentally observed; species = human only; no tissues or cell lines or mutations were included. Only examined categories associated with molecular and cellular functions, as outlined by (57). Additionally, protein interaction maps were generated using STRING consortium (https://string-db.org/) to visualize clusters of protein-protein interactions.

### Generation of RC.V8-Infected QT6 Nuclear Lysates

QT6 cells were infected for 4 hours with cell culture medium obtained from a separate culture of QT6 cells transfected with pRC.V8. Cells were fractionated using the method described in (50) with minor modifications, as described below. Cells were trypsinized, pelleted at low speed, and washed in cold PBS. The cell pellet was resuspended in lysis buffer (10 mM HEPES pH 7.9, 10 mM KCl, 0.1 mM EDTA, 0.3% Nonidet P-40, 1 mM DTT, 100 μg/ml phenylmethanesulfonyl fluoride (PMSF), 1 μg/ml pepstatin, 100U/ml Omnicleave (Epicentre), and Roche Complete Protease Inhibitor Cocktail) and incubated on ice for 5 minutes. Cells were then spun for 5 minutes at 3,000 rpm at 4°C to pellet nuclei, and the supernatant (cytoplasmic fraction) was collected. The pelleted nuclei were washed once with lysis buffer, then resuspended in nuclear extract buffer (20 mM HEPES pH 7.9, 10 mM NaCl, 1 mM DTT, 100 μg/ml PMSF, 1 μg/ml pepstatin, 100U/ml Omnicleave (Epicentre), and Roche Complete Protease Inhibitor Cocktail) and incubated at 37°C in a water bath for 10 minutes. Nuclear lysate was placed on ice, and solution was brought to 400 mM NaCl and 1 mM EDTA, followed by vortexing on high for 15 seconds and 20 minutes of rotation at 4°C. Debris was pelleted at 13,000 rpm for 10 minutes at 4°C, and supernatant was transferred to a fresh tube (nuclear fraction). Protein concentration in lysates was determined by Bradford assay.

### RSV Gag-Med26 Co-immunoprecipitation

RC.V8-infected nuclear lysates (500 µg) were pre-incubated for 2 hours with α-RSV-CA antibody (mouse α-RSV CA.A11, gift from Neil Christensen, Penn State College of Medicine) in low salt NET2 buffer (50 mM Tris pH 7.4, 150 mM NaCl, 0.05% Nonidet P-40; (58)) at 4°C with rotation. With 1 hour remaining, Pierce™ Protein G Magnetic Beads (60 µl of a 50% slurry per reaction) were washed 4 times with high salt NET2 buffer (50 mM Tris pH 7.4, 400 mM NaCl, 0.05% Nonidet P-40; (58)), then blocked with 5% w/v BSA in low salt NET2 buffer at 4°C with rotation. At the end of the 2 hours, blocking buffer was removed and beads were resuspended in low salt NET2 buffer. An equal amount (∼30 µl) was added to each reaction, and tubes were rotated at 4°C overnight.

After overnight incubation, buffer was removed and beads were washed 4 times with high salt NET2 buffer. Bound proteins were then eluted by boiling beads in 50 µl of 1X SDS-PAGE buffer for 10 minutes at 100°C. Beads were pelleted at 13,000 rpm for 5 minutes, and supernatant was taken for analysis.

Samples were run on 10% SDS-PAGE gels, transferred to PVDF, blocked for 30 minutes with 5% Milk/0.1% TBS-Tween, and then incubated with primary antibody (rabbit α-Med26, Proteintech, 21043-1-AP) in 0.5% Milk/0.1% TBS-Tween at 4°C with rocking overnight. Membranes were washed 3 times for 5 minutes with 0.1% TBS-Tween, then incubated with secondary antibody (goat α-rabbit-HRP, Sigma A0545) for 1 hour at room temperature. Washes were repeated, and membranes incubated with ECL 2 for 5 minutes. Western blots were imaged using a BioRad ChemiDoc MP imager. Blots were stripped and reprobed for RSV Gag using rabbit α-RSV-W [gift from John Wills, Penn State College of Medicine (51) and secondary antibody (goat α-rabbit-HRP, Sigma A0545)].

### Confocal Imaging

QT6 cells were plated at a density of 0.5 x 10^6^ cells/well in 6-well tissue culture dishes containing #1.5 coverslips and were allowed to settle overnight. The following afternoon, wells were transfected with 500 ng of RSV Gag-GFP (11) and 125 ng of FLAG-tagged Med26 (a gift from Joan Conaway and Ronald Conaway [Addgene plasmid #15367; http://n2t.net/addgene:15367; RRID:Addgene_15367)(59)] expression vectors. The following morning, cells were washed 2X quickly with warm PBS, and fixed in 3.7% formaldehyde in PBS for 10 minutes at RT. Slides were then washed 3X with PBS (5 minutes per wash), permeabilized for 15 minutes in 0.25% Triton X-100/PBS, washed 3X, and blocked for 30 minutes at RT in 10% BSA/PBS. Relevant slides were incubated for 2 hours with mouse α-FLAG-M2 antibody (Sigma, F1804) in 3% BSA/PBS in a humid chamber at 37°C followed by 3X washes. Secondary antibody staining with donkey α-mouse-AlexaFluor647 (AF647) was carried out for 2 hours in 3% BSA/PBS in a humid chamber at 37°C, followed by 3X washes. Slides were DAPI stained for 1 minute at RT, mounted with ProLong™ Diamond (Life Technologies), and cured for 24 hours. Slides were imaged on a Leica AOBS SP8 confocal microscope with a 63x/1.4 oil objective, with pinhole set to 1 airy unit and frame averaging set to four. FLAG-Med26-AF647 was excited using a white light laser (WLL) tuned to 647 nm at 2% laser power, and emission was detected via hybrid detector. RSV Gag-GFP was excited by WLL tuned to 488 nm at 3% laser power and detected via hybrid detector. DAPI was excited using a 405 nm diode laser at 8% power and detected using a photomultiplier tube.

Colocalization between RSV Gag and Med26 was assessed using Imaris image analysis software (Oxford Instruments). Briefly, images were Gaussian filtered and surfaces were generated around cell nuclei using the DAPI channel as reference. Masks were created for RSV Gag and Med26 signal within nuclear surfaces, and colocalization of this signal was assessed using the Imaris colocalization tool. Manders’ Overlap Coefficients were exported from Imaris and data was assessed for outliers by Grubbs’ test with an α = 0.05 using GraphPad Prism (GraphPad Software, Inc.). Statistical analysis and generation of **Fig 4C** was also done with this software. A total of 17 M1 (Med26∩Gag) and 18 M2 (Gag∩Med26) individual data points were plotted. Representative images (**Fig 4B**) and video (**Video S1**) were created using Imaris software.

## 3. Results

### 3.1. Subcellular localization

Prior experiments using microscopy revealed that both RSV and HIV-1 Gag proteins localize to the perichromatin compartment of the nucleus where they associate with their cognate USvRNAs at active transcription sites (3, 9, 11). In addition, HIV-1 Gag associates preferentially with marks associated with transcriptionally-active euchromatin compared to heterochromatin (10). To further define the nuclear subcompartments where RSV Gag is localized, cells expressing wild-type or mutant Gag proteins were separated into cytoplasmic, nucleoplasmic, and chromatin-associated protein fractions (**Fig 1**). The first chromatin fraction was extracted using a NaCl concentration of 150 mM, which removes proteins that are loosely associated with chromatin (i.e., the euchromatin fraction). In the second chromatin fraction, a higher salt concentration was used (500 mM NaCl) along with detergent, to remove proteins that are more tightly bound to chromatin (50) (i.e., the heterochromatin fraction).

**Figure 1.**
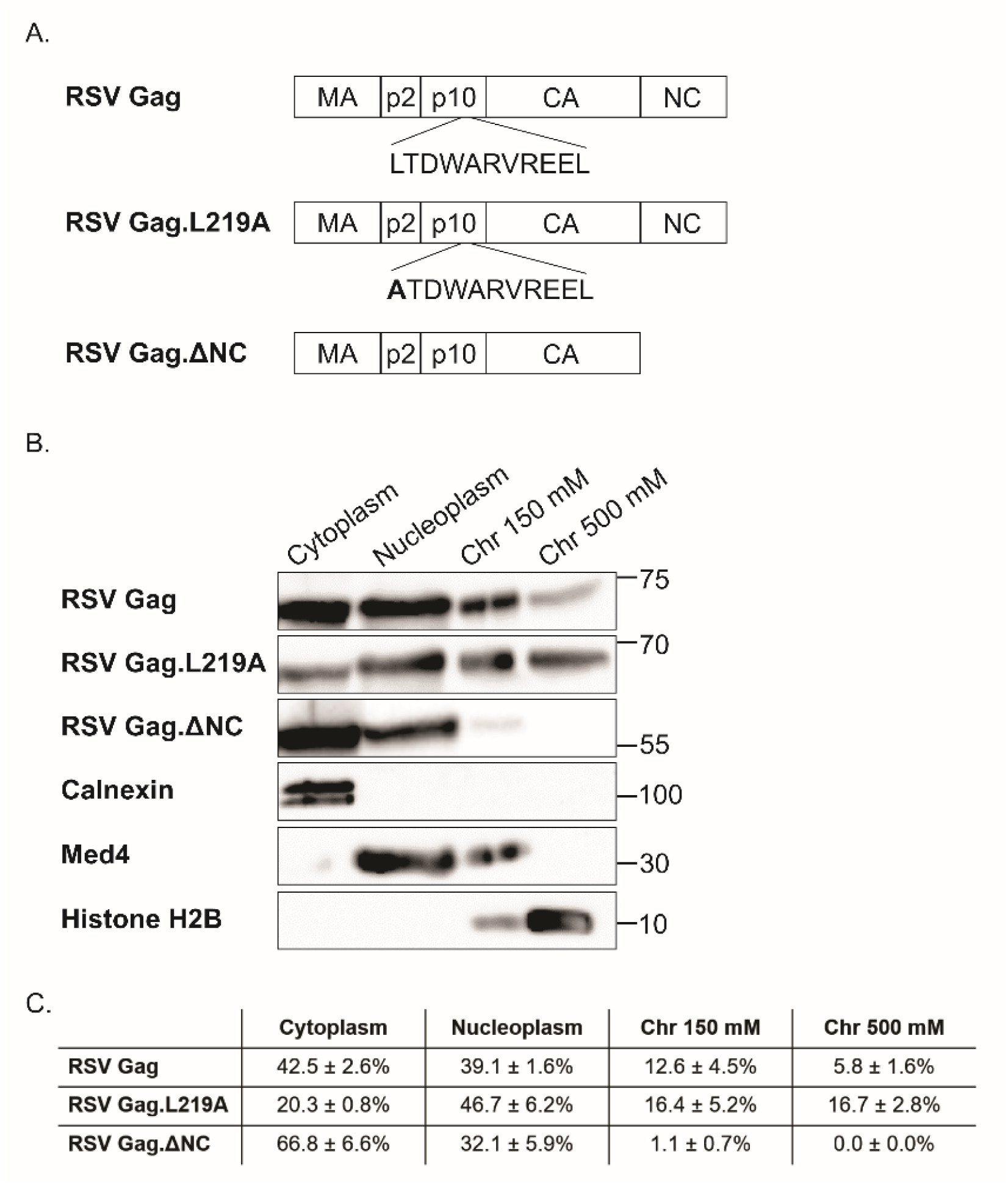
Schematic diagram and western blot with quantitation. (**A**) RSV Gag constructs. (**B**) Subcellular fractionations were performed to separate the cytoplasm and nucleoplasm from the chromatin fractions, which were further separated using differential NaCl concentrations. The 150 mM chromatin fraction (Chr 150 mM) contains proteins associated with open chromatin (euchromatin). The 500 mM chromatin fraction (Chr 500 mM) contains proteins that are associated with condensed chromatin (heterochromatin). Wild type RSV Gag and Gag.L219A were detected in all of the fractions at different ratios. Gag.ΔNC was primarily detected in the cytoplasm with very little in the chromatin fractions. (**C**) Band densities were determined for each Gag construct for each fraction and are displayed as mean ± standard error of the mean (SEM) for three biological replicates. To assess fraction purity, cellular proteins were detected using antibodies against calnexin (cytoplasm), Med4 (nucleoplasm and euchromatin), and Histone H2B (euchromatin and heterochromatin). The position of molecular weight markers, in kilodaltons, are indicated on the right.

The signals of the protein bands were quantified to yield the relative ratio of Gag protein in each fraction, demonstrating that RSV Gag was present in the cytoplasm (42.5 ± 2.6%) and the nucleoplasm (39.1 ± 1.6%) of cells (**Fig 1A**). Interestingly, RSV Gag was also present in both chromatin-associated protein fractions (euchromatin, 12.6 ± 4.5%; heterochromatin, 5.8 ± 1.6%). Examination of the nuclear-restricted mutant Gag.L219A, which contains a point mutation in the p10 NES, demonstrated decreased signal in the cytoplasm (20.3 ± 0.8%) and increased signal in both the nucleoplasm (46.7 ± 6.2%) and heterochromatin-associated fractions (16.7 ± 2.8%), compared to wild type Gag. Analysis of mutant Gag.ΔNC, which lacks the NC domain required for genomic RNA packaging, demonstrated an increase in cytoplasmic Gag compared to wild type (66.8 ± 6.6%), with little to no Gag.ΔNC in either of the chromatin fractions. Controls for the fractions included Calnexin (endoplasmic reticulum/cytoplasm), Mediator subunit 4 (Med4; nucleoplasm and euchromatin), and histone 2B (chromatin fractions). We have observed similar localization of HIV-1 Gag to all four subcellular compartments, as reported (9).

### 3.2. Affinity purification and proteomic analysis

We next set out to identify potential nuclear interacting partners of RSV and HIV-1 Gag to provide clues about their possible role(s) in the nucleus. Recombinant His-tagged RSV and HIV-1 Gag proteins purified from *E. coli* were incubated with nuclear lysates from DF1 chicken fibroblast cells or HeLa cells, respectively. A beads-only control was also performed using DF1 lysates and HeLa lysates incubated with nickel beads. The affinity purifications were performed twice for both Gag proteins, as well as the beads-only controls. Proteins identified using mass spectrometry were combined into a single list for each Gag protein and analyzed using DAVID and Ingenuity Pathway Analysis, as described in Materials and Methods. Proteins identified from the beads-only purifications were removed from the lists of RSV and HIV-1 affinity-tagged purifications. Proteins that had an unused score, as defined by ProteinPilot, of less than 1.3 were removed, along with common contaminants (see Supplemental files S1, S2, S3, and S4 for raw data).

DAVID was used to assign each protein from the two final Gag proteome lists to their cellular compartment(s) and biological process categories (16, 19). We used the functional annotation tool to determine the most relevant gene ontology (GO) terms associated with the proteomics lists. The order of the GO terms that were enriched in the Gag proteome lists are dependent upon the p-values for each GO term. The smaller the p-value, the more that particular GO term was enriched. **Table S1** shows the results of using the DAVID analysis software to analyze the list of proteins obtained from the RSV Gag affinity purification using DF1 nuclear lysates, and the top 10 biological processes’ GO terms are displayed. Only nuclear proteins, as determined by DAVID, were analyzed to determine the enriched biological functions, and **Table S2** shows the top 10 results. Included in the results were generic GO terms such as: GO:0034641∼cellular nitrogen compound metabolic process, GO:1901360∼organic cyclic compound metabolic process, and GO:0006807∼nitrogen compound metabolic process. There were also more specific GO terms that were enriched, including GO:0010467∼gene expression and GO:0006396∼RNA processing.

Next, the potential binding partners of HIV-1 Gag isolated from HeLa nuclear lysates were examined. The terms GO:0006396∼RNA processing and GO:0010467∼gene expression were among the top 10 GO biological process terms identified after DAVID analysis (**Table S3)**. When only nuclear proteins were examined, the top 10 GO terms again included GO:0010467∼gene expression and GO:0006396∼RNA processing (**Table S4)**.

The proteins identified for each RSV and HIV-1 Gag affinity purification were compared, and we found there were 57 proteins that overlapped from 317 total proteins from the RSV list and 436 total proteins from the HIV-1 list (**Fig 2**). **Table 1** contains the names and functions of the 57 proteins in common between the RSV and HIV-1 Gag. The functions of the 57 proteins are varied, encompassing cell cycle progression, RNA methylation, RNA processing and splicing, cytoskeletal regulation, mRNA nuclear export, transcription, DNA repair, and chromatin modification.

**Figure 2.**
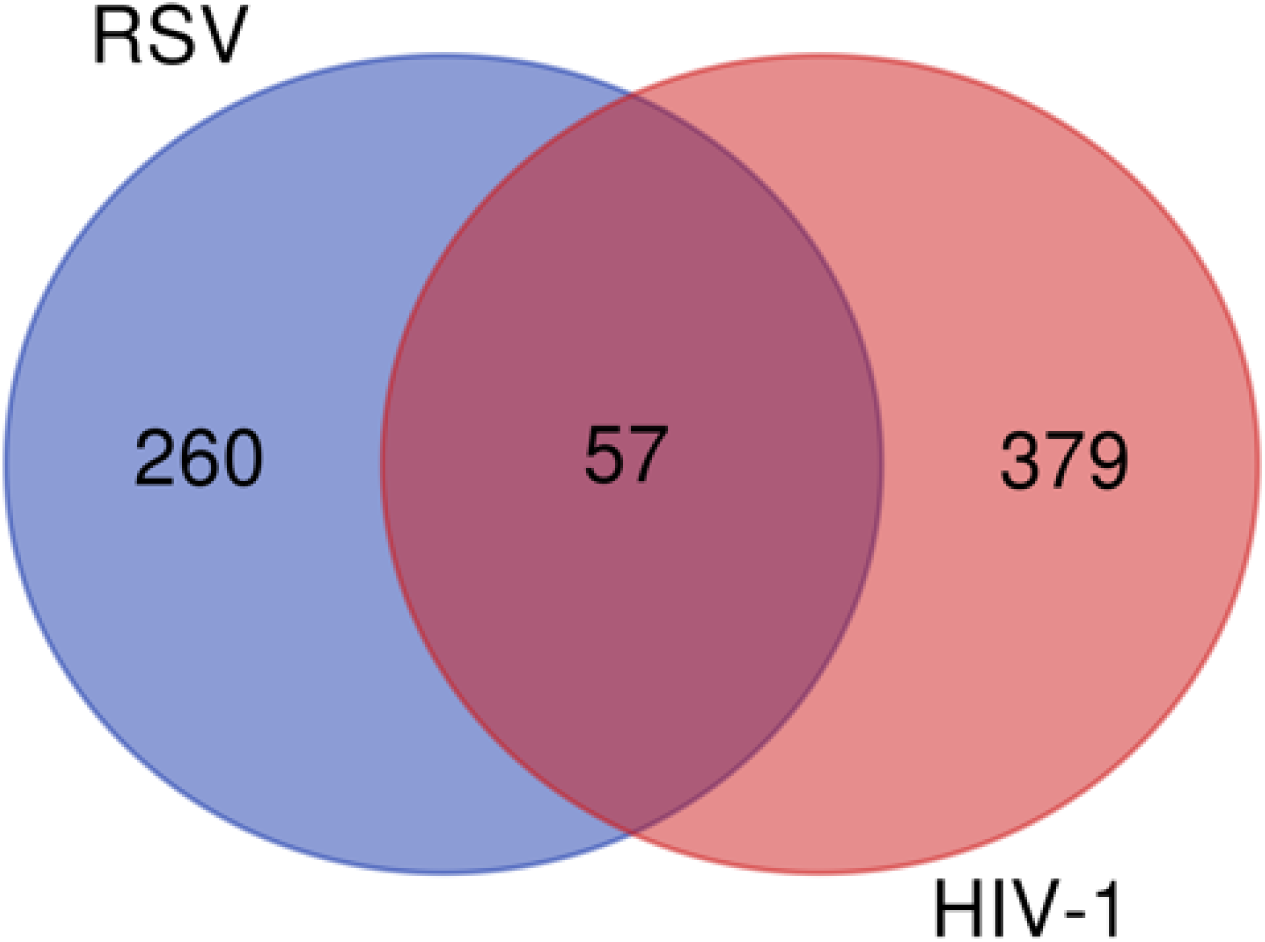
The number of proteins identified in both the RSV and HIV-1 Gag affinity tagged purifications. 57 proteins were found to be in common between RSV and HIV-1 Gag.

**Table 1.**
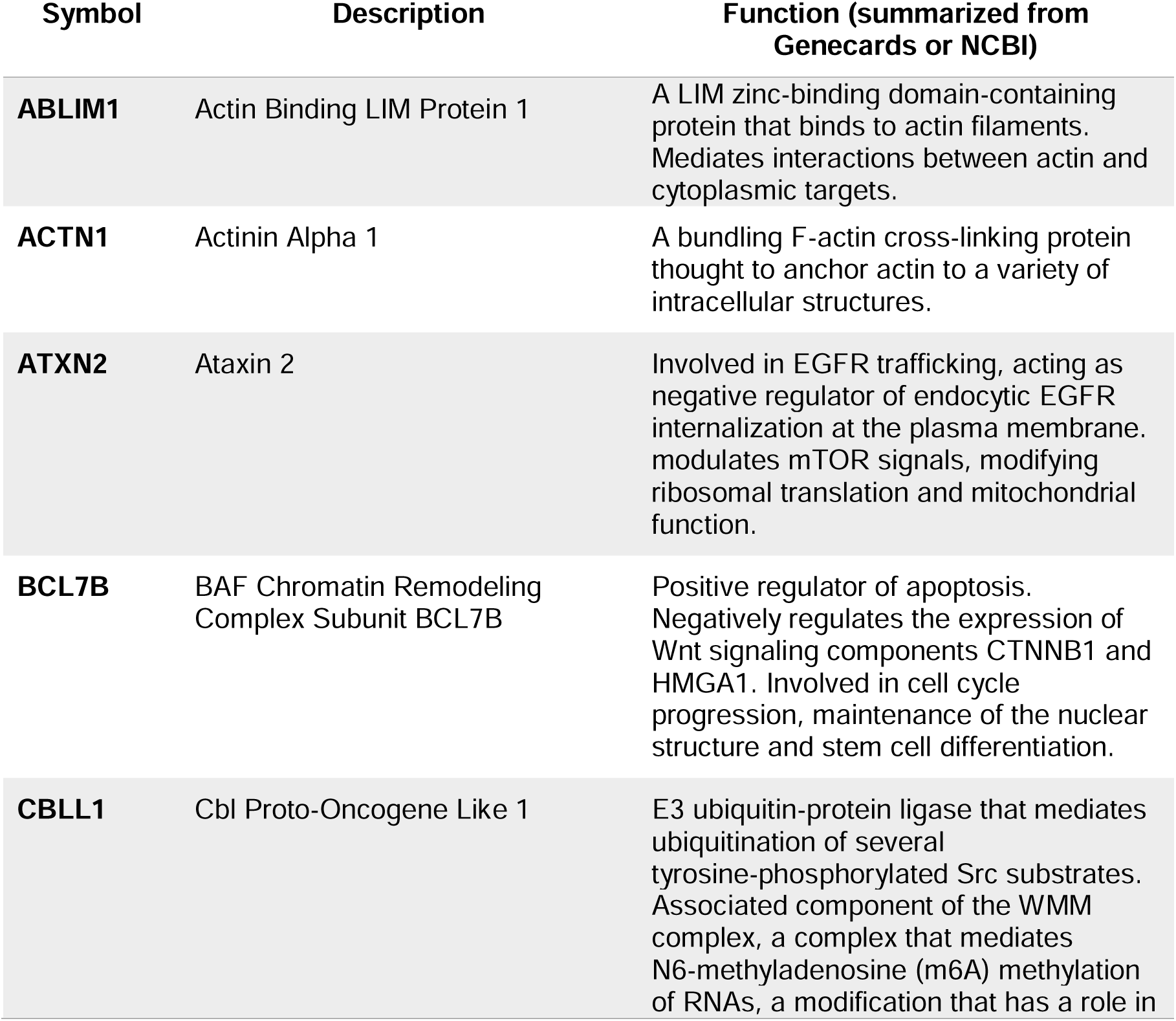
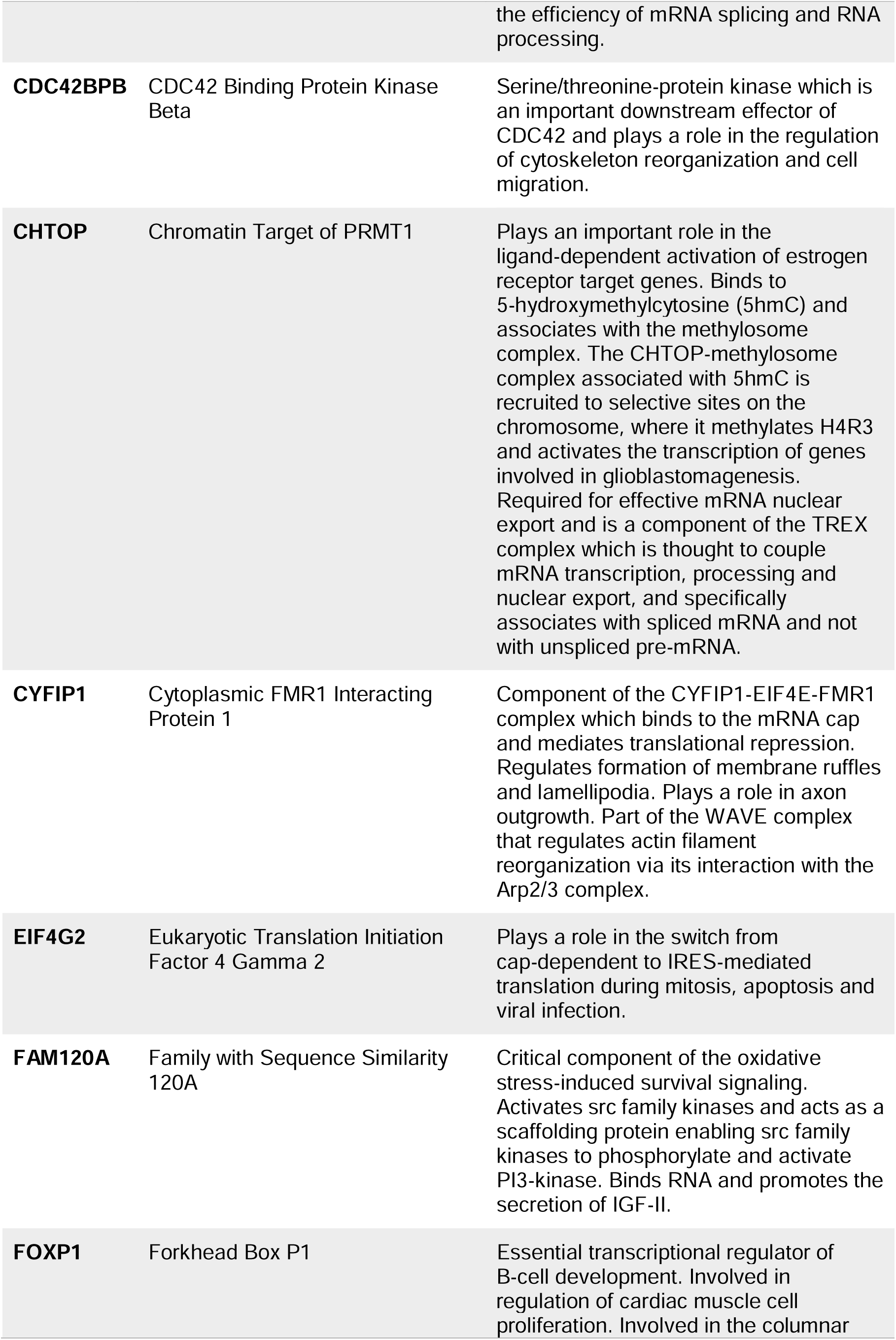
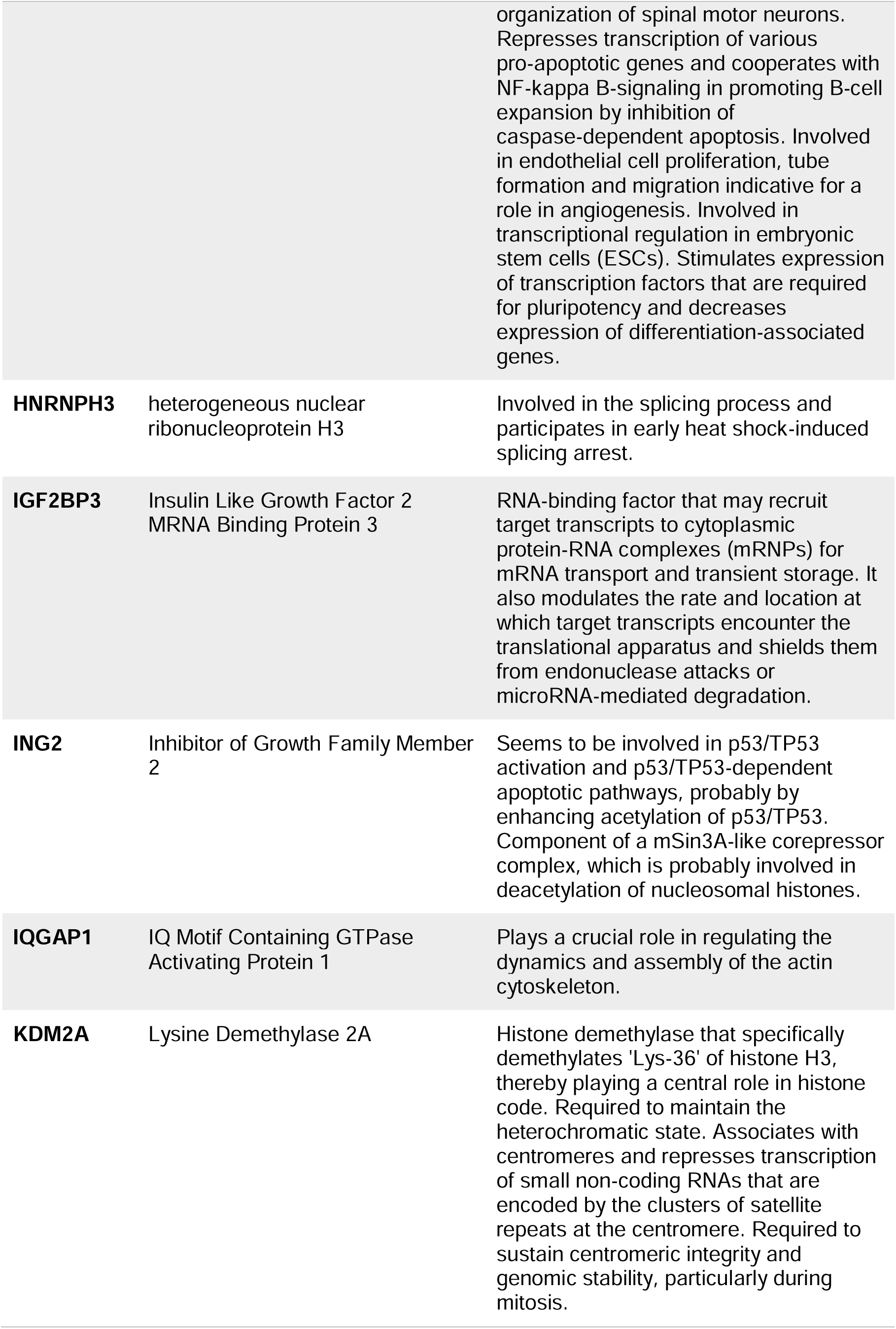
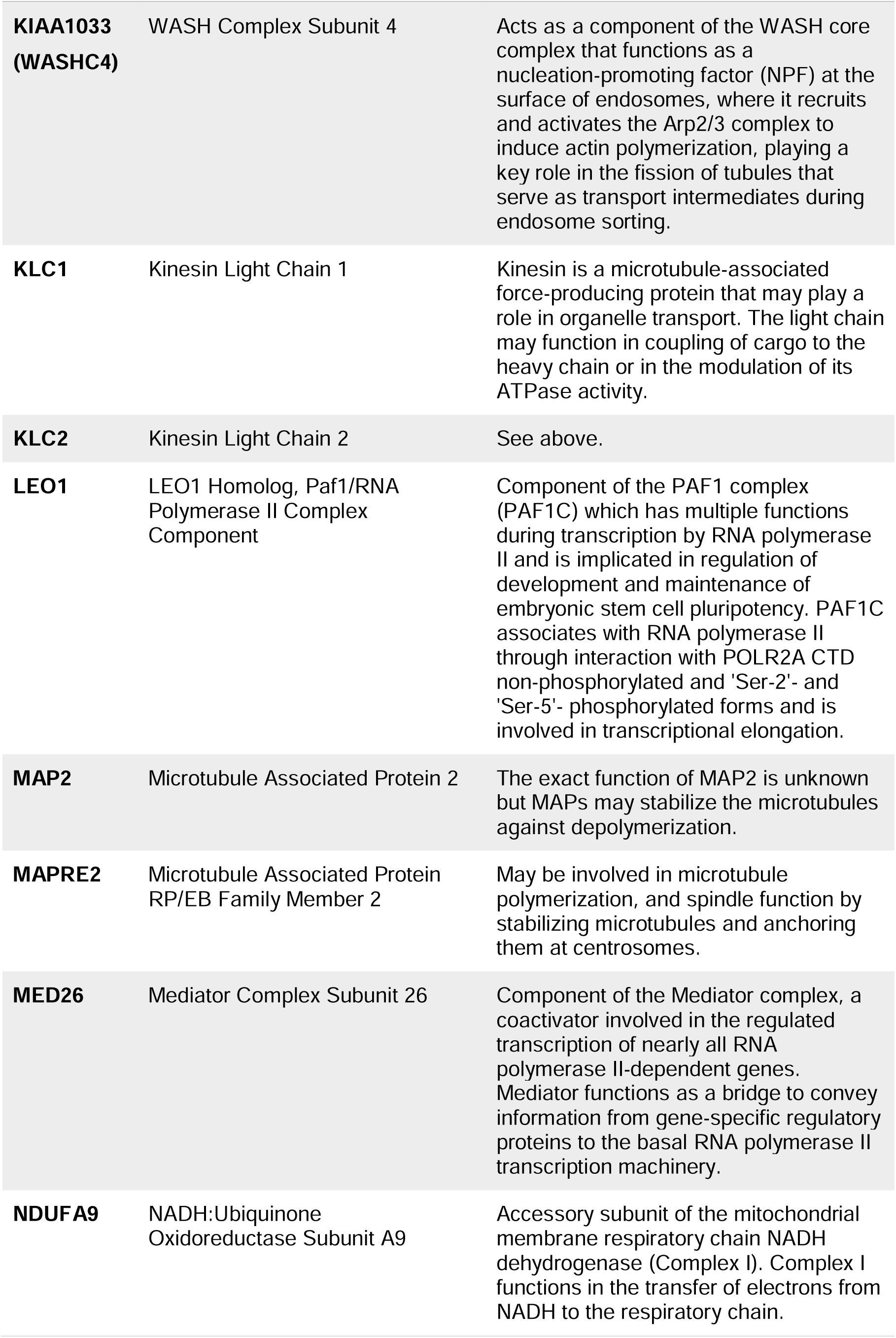
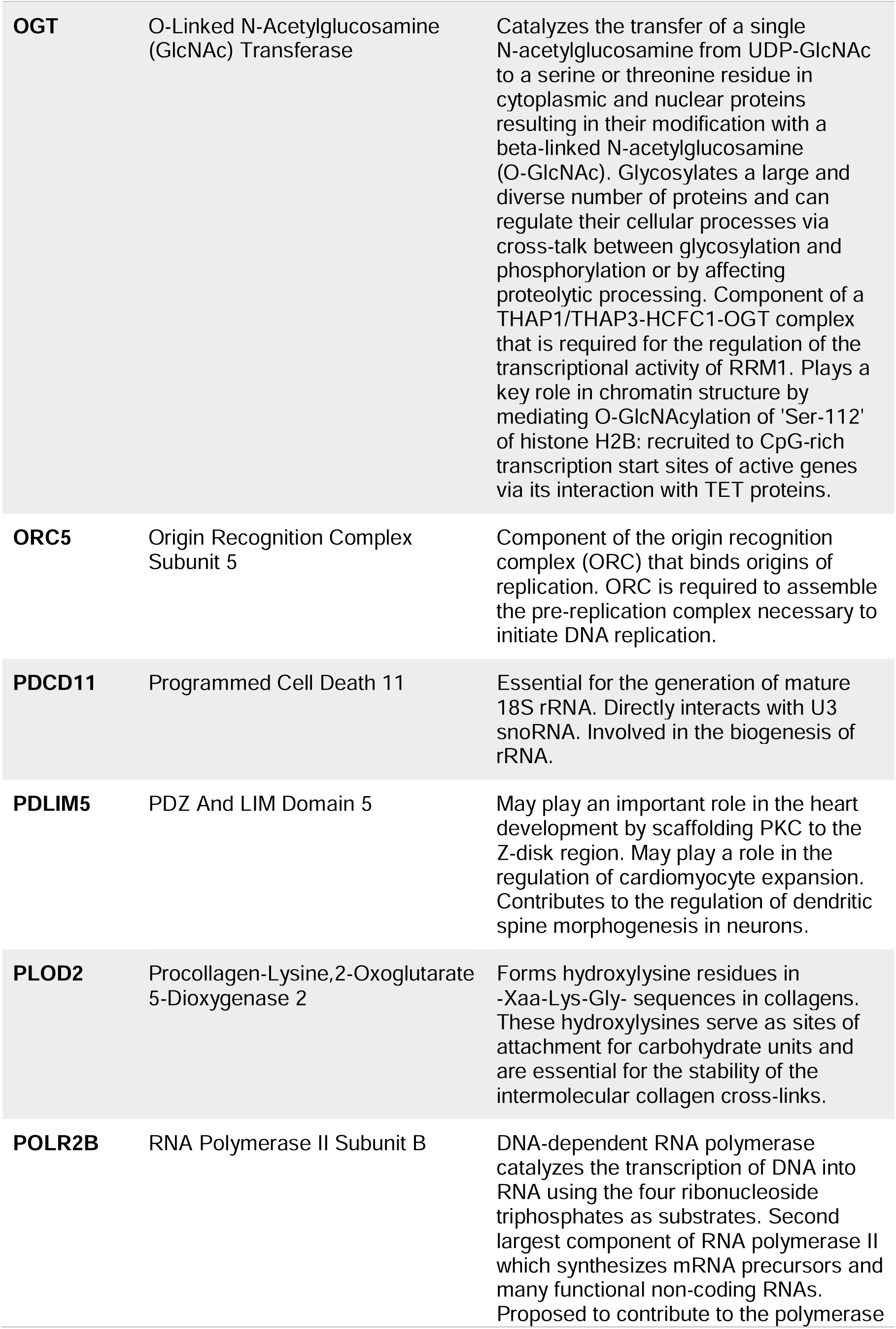
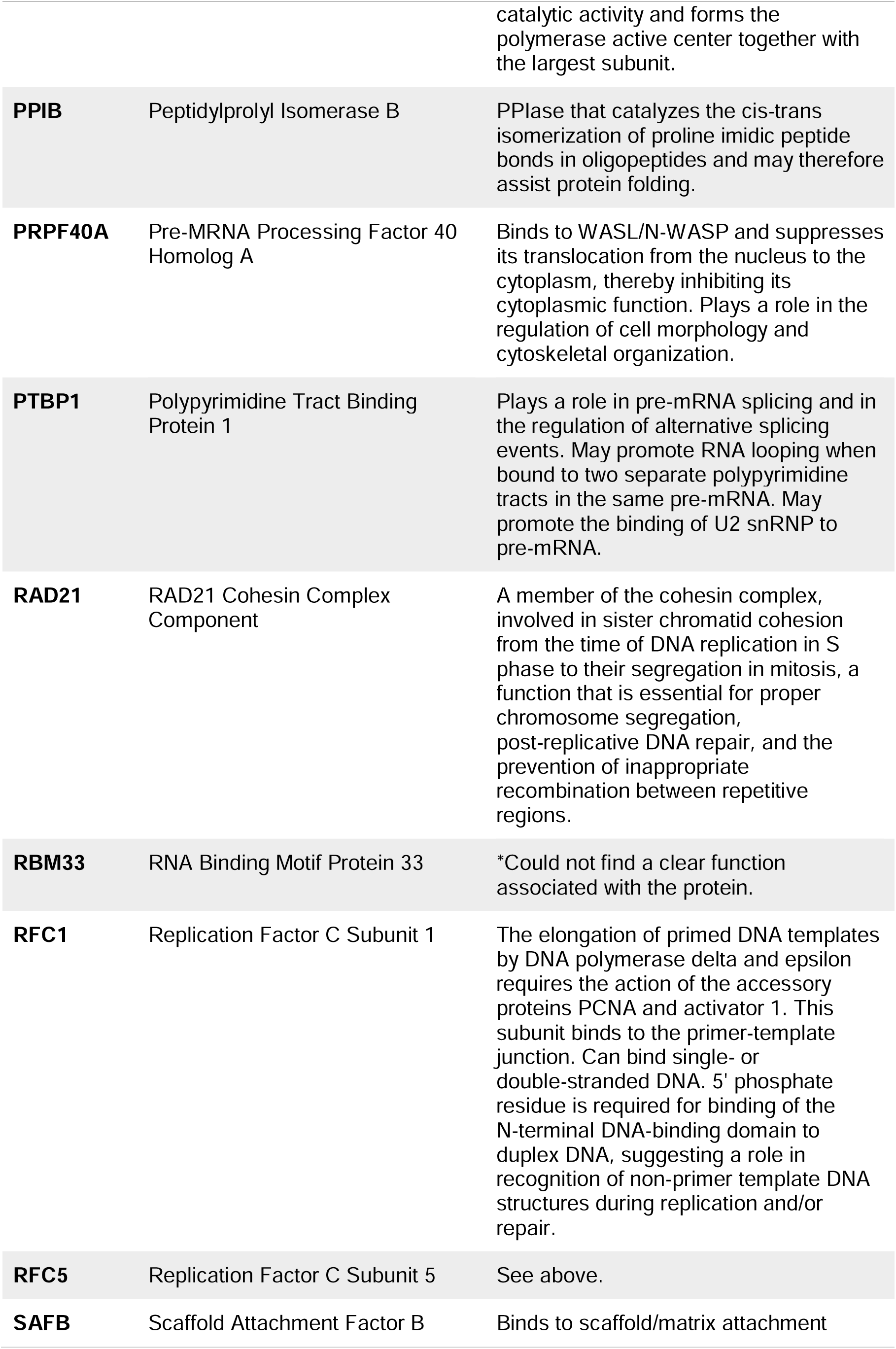
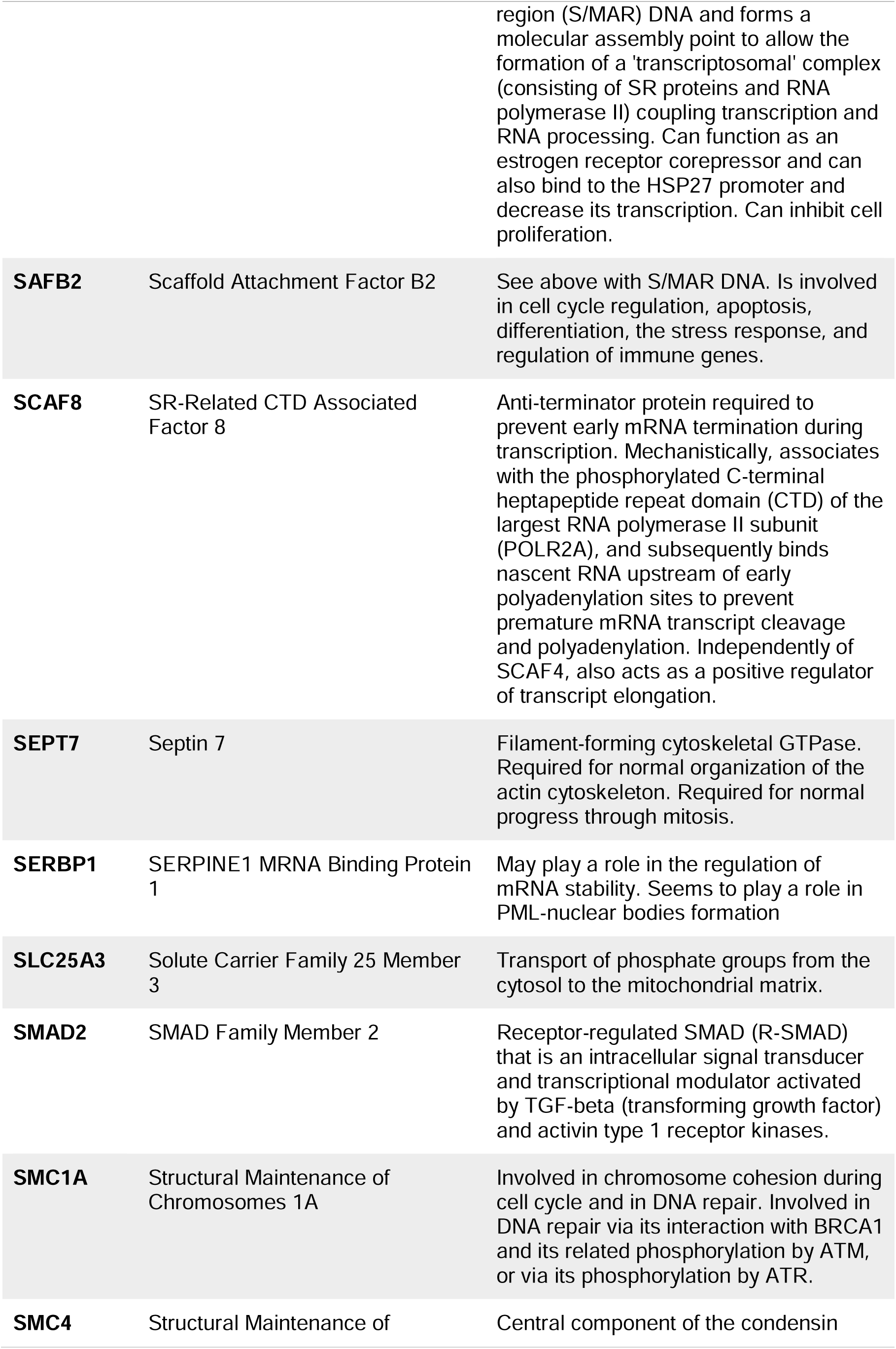
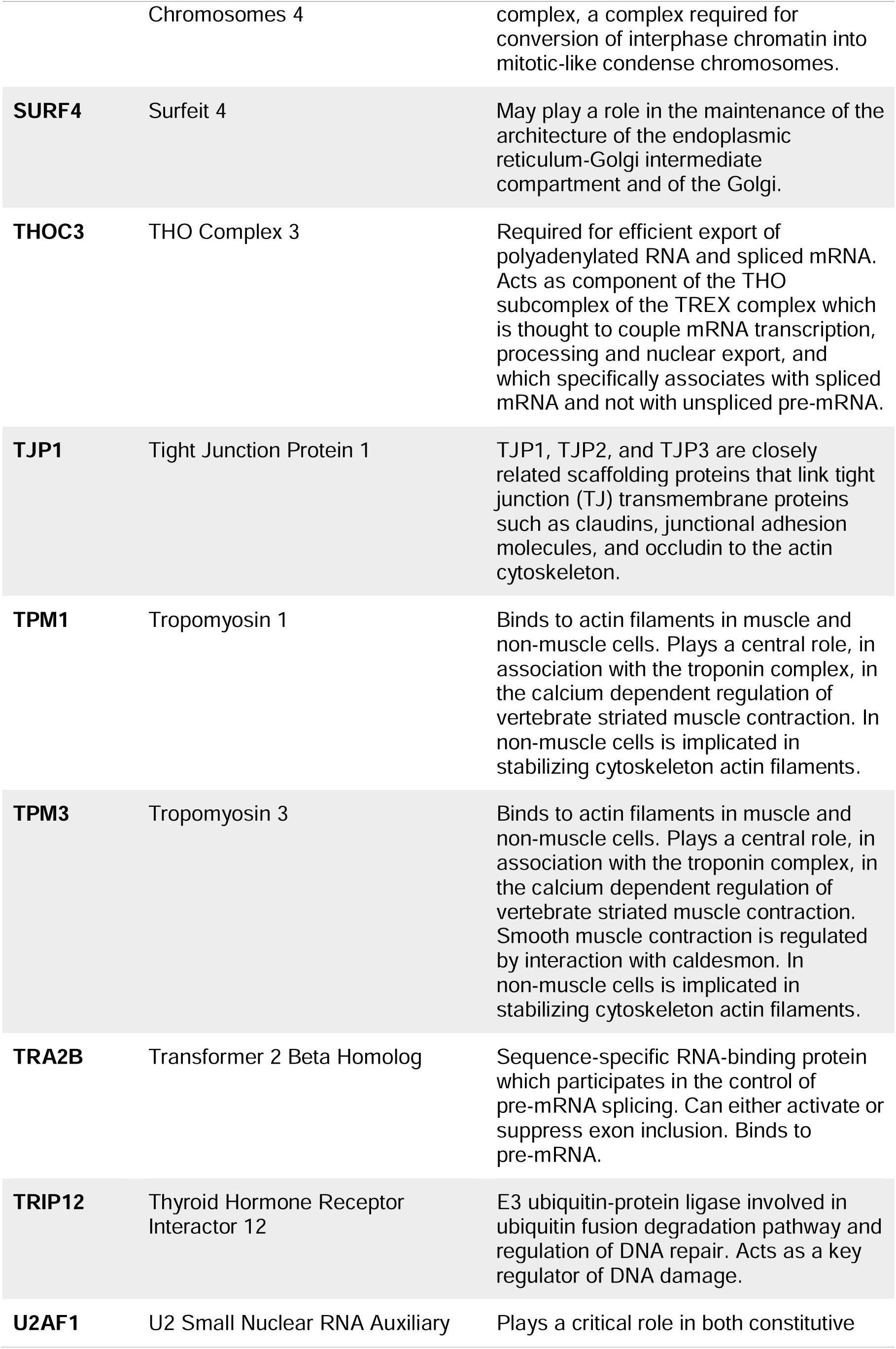
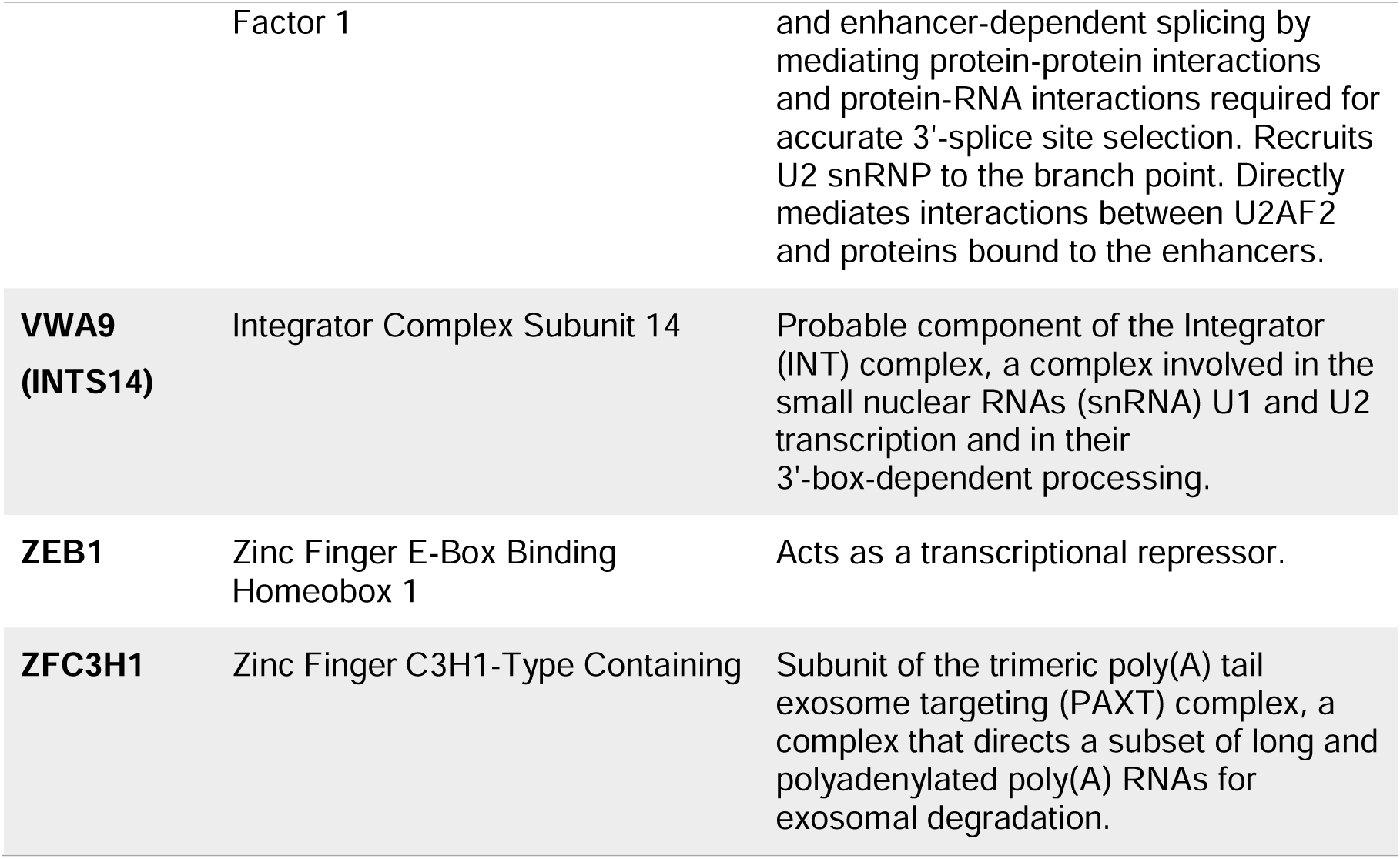
Names and functions of the 57 cellular proteins identified in both the RSV and HIV-1 Gag protein interactomes.

### 3.3. Comparative proteomic analysis of previously published HIV-1 Gag mass spectrometry experiments

Mass spectrometry experiments focusing on HIV-1 Gag cellular binding partners have been previously reported by other laboratories; however, these investigators did not focus on the nuclear proteins identified in these experiments. Engeland *et al.,* 2011 (31) performed five independent affinity-tagged purification experiments to identify cellular proteins that interact with HIV-1 Gag. The techniques used consisted of a tandem affinity purification (TAP) tag for a C-terminally tagged Gag, GFP-TRAP A beads, and GFP microbeads for Gags with GFP fused either internally to the MA domain or the C-terminus of Gag. Each of these Gag constructs were transfected into 293T cells and whole cell lysates were used for the affinity pulldowns. The authors found 31 proteins that were identified in at least 3 of the experiments, and of these 31 proteins, 24 localize to the nucleus. Using DAVID to analyze these 24 proteins, GO:0006396∼RNA processing and GO:0010467∼gene expression were the top two biological functions identified (**Table S5**). In 2014, Engeland and colleagues (30) examined the HIV-1 Gag interactome again and found 944 proteins that met their inclusion criteria. Using DAVID to analyze these 944 proteins, 186 were found to be nuclear. The authors performed a GO enrichment analysis using DAVID to determine the most enriched biological processes and found nuclear processes including GO:0006396∼RNA processing, GO:0008380∼RNA splicing, and GO:0006334∼nucleosome assembly. We performed a GO analysis using solely the 186 nuclear proteins identified and found similar results (**Table S6**).

In Jäger *et al.,* 2012 (32), the authors identified host proteins that interact with each of the HIV-1 polyproteins, processed proteins, and accessory proteins, in a systematic and quantitative manner. A purification strategy was used that consisted of appending two Strep tags and three FLAG tags at the C-terminal end of each HIV-1 protein, which were expressed separately in HEK293 or Jurkat cells. When examining their raw data, we found that 1,134 unique proteins were identified from affinity purifications using full-length Gag plus the proteolytic cleavage products of Gag (MA, CA, NC, and p6). Of these interacting proteins, 180 were identified as being nuclear by DAVID analysis. Limiting our GO analysis using DAVID to these nuclear proteins, GO:0006396∼RNA processing was the top biological function term identified, followed by protein targeting functions and RNA metabolism (**Table S7**).

Ritchie *et al.,* 2015 (35) provided further information on the possible protein interactors of HIV-1 Gag using the *E. coli* biotin ligase BirA* to permit identification of protein partners in close proximity to the bait protein (60–62). The BirA* tag was inserted within the MA domain of Gag, and the Gag-BirA* construct was expressed in cells. The authors found 53 proteins after their exclusion criteria eliminated nonspecific interactors, and from these 53 proteins, we identified 17 nuclear proteins by DAVID analysis. **Table S8** shows DAVID analysis of these 17 proteins, identifying the top biological categories of GO:0098609∼cell-cell adhesion, GO:0090304∼nucleic acid metabolic process, and GO:0010608∼posttranscriptional regulation of gene expression.

Le Sage *et al.,* 2015 (33) also identified potential host factors that interact with HIV-1 Gag. Similar to Ritchie *et al.*, they used the BirA* tagging system, except the tag was placed at the N-terminus of Gag. They found a total of 42 proteins, of which 19 were nuclear. When these 19 nuclear proteins alone were analyzed by DAVID, the top hits included protein targeting and RNA processing and metabolism (**Table S9**).

Li *et al.,* 2016 (34) examined binding partners of the MA domain of HIV-1 Gag by inserting a Strep tag at the C-terminus of MA and collecting MA-interacting complexes after HIV-1 infection. There were 97 proteins identified that met their inclusion criteria and were not present in the lysate-only control. Of these proteins, 63 were nuclear. When only the nuclear proteins were further analyzed by DAVID, the top categories were ER targeting and viral transcription/gene expression (**Table S10**).

There is one other publication that performed a tandem immunoprecipitation to identify HIV-1 Gag binding partners (63) that was not included in the comparative analysis presented here due to the small number of proteins identified. Gao *et al.,* identified 12 individual proteins and each of these were present in at least one of the previously discussed publications or found in the experiment we presented earlier in this report. They identified numerous ribosomal proteins, HNRNP proteins, and other nuclear proteins including histone proteins, EF1-α, nucleolin, B23, Nopp34, and SNRPD3.

To provide a comprehensive view of the set of proteins identified in multiple studies, proteins identified in the previously published laboratories’ reports were compared to the list of proteins found in our HIV-1 Gag affinity purifications. **Table 2** shows the list of proteins identified in our HIV-1 Gag pulldown as well as at least one of the publications discussed above. **Table 2** also indicates whether each protein was previously demonstrated to have a role in HIV-1 replication based on the HIV-1 human protein interaction dataset (64–66). A total of 129 proteins identified in our mass spectrometry analysis were also present in at least one of the prior publications.

**Table 2.**
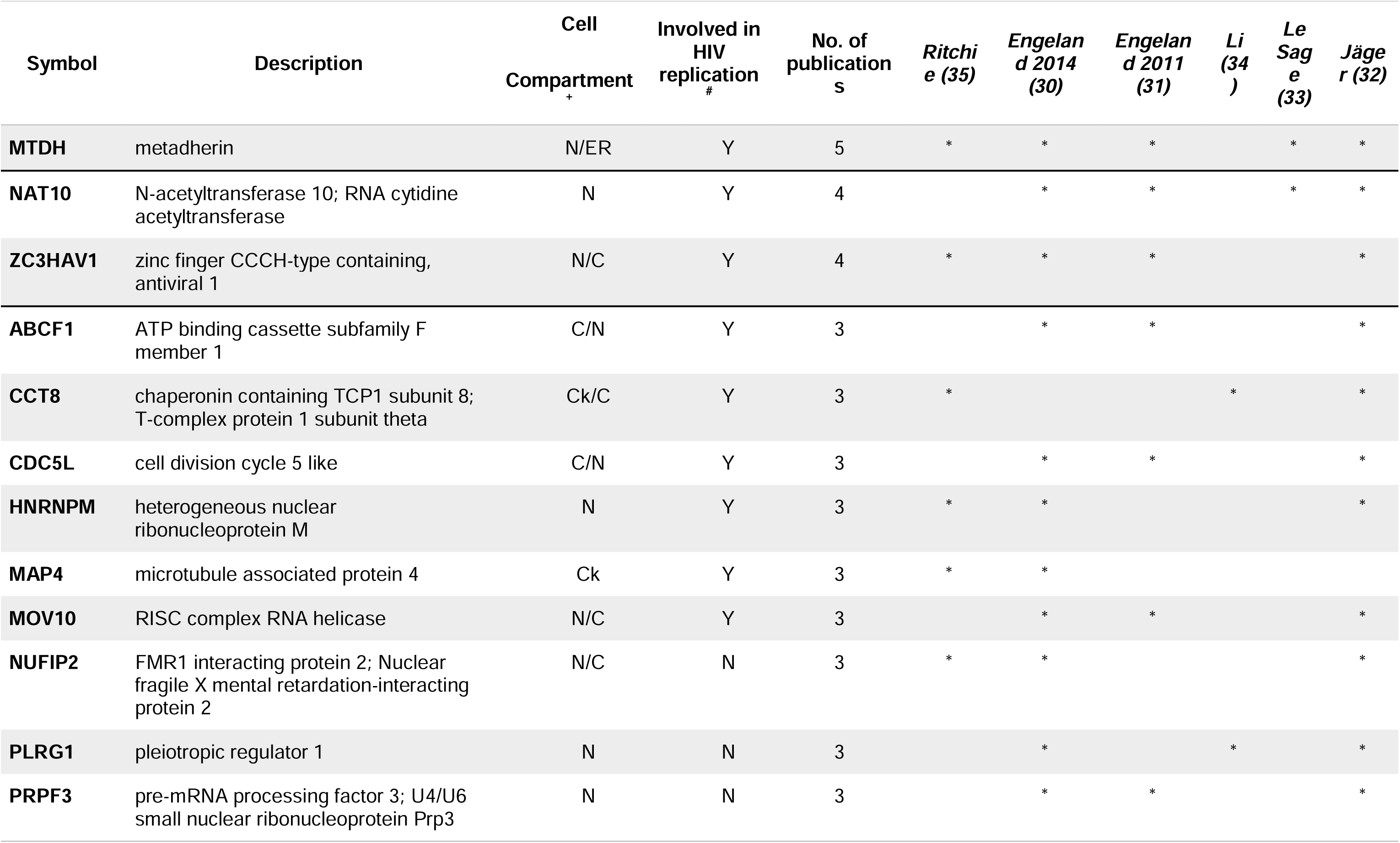
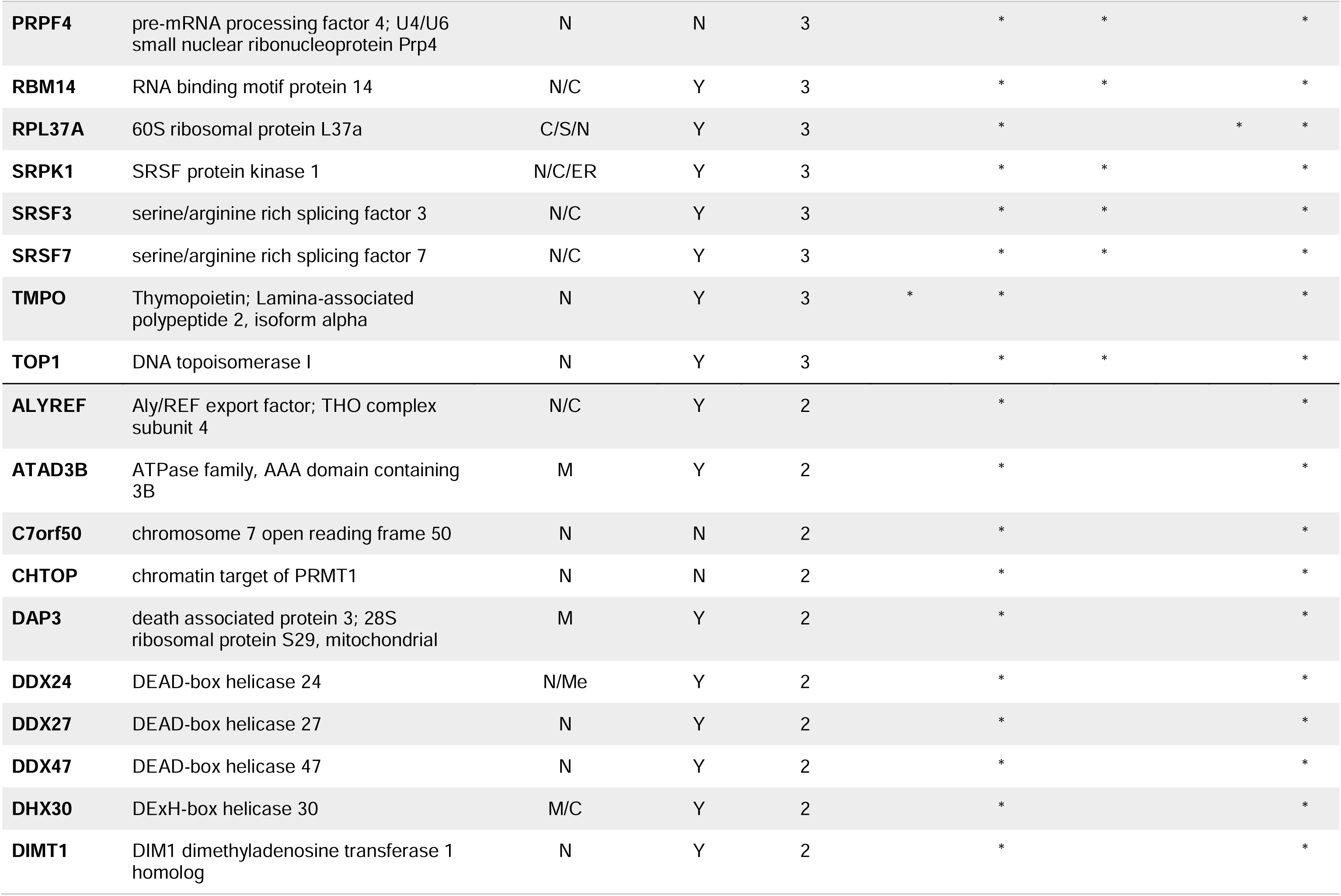
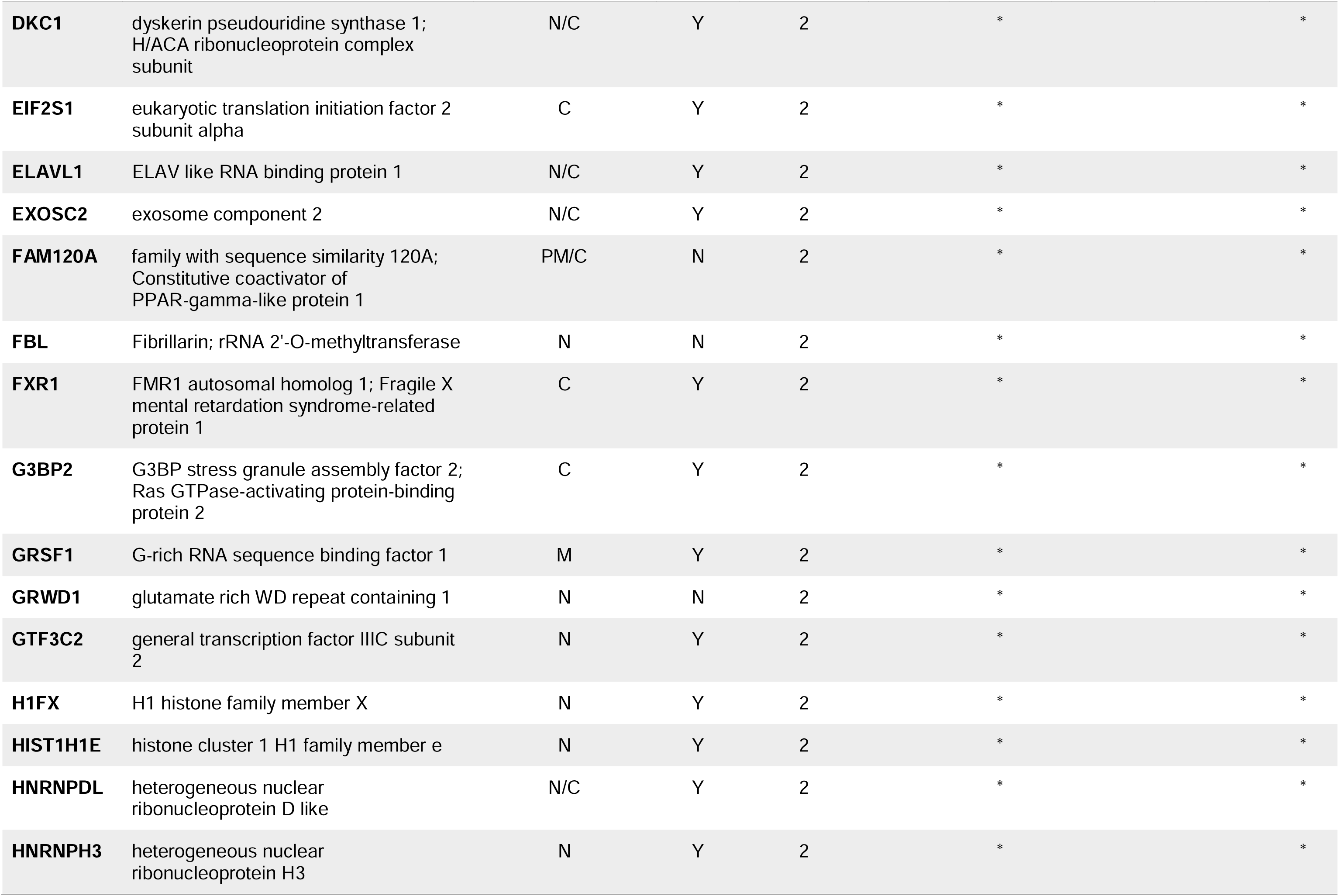
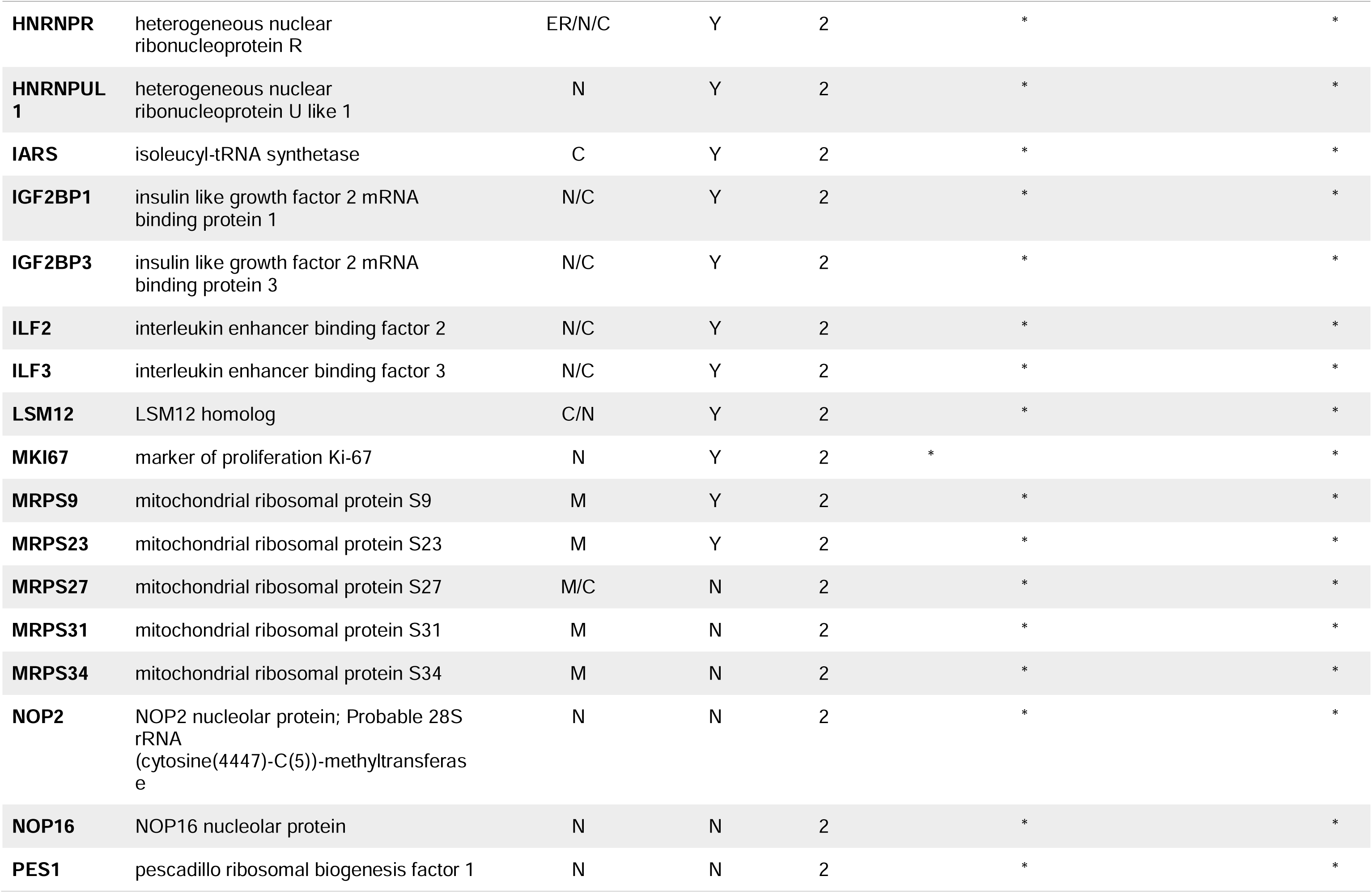
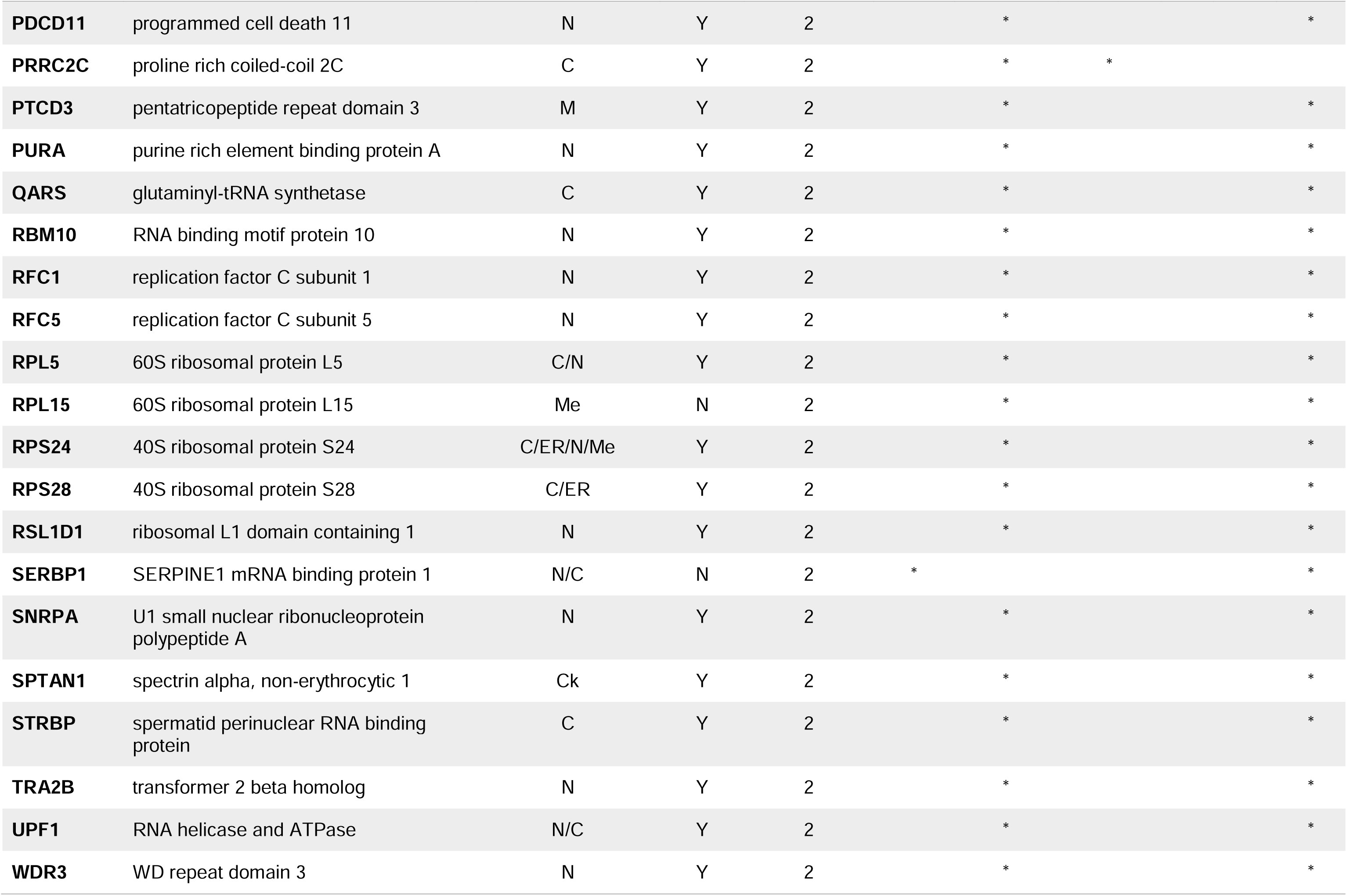
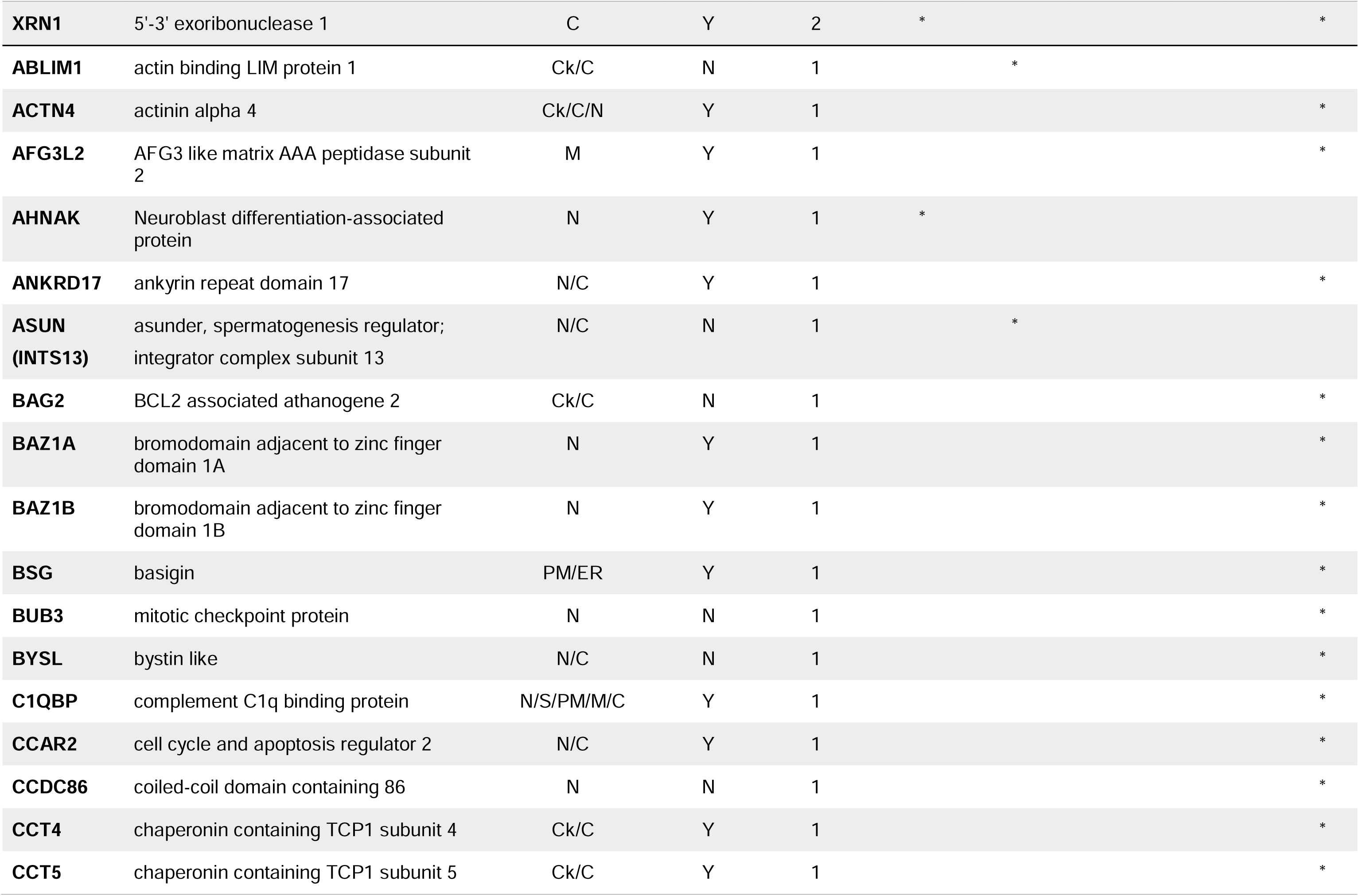
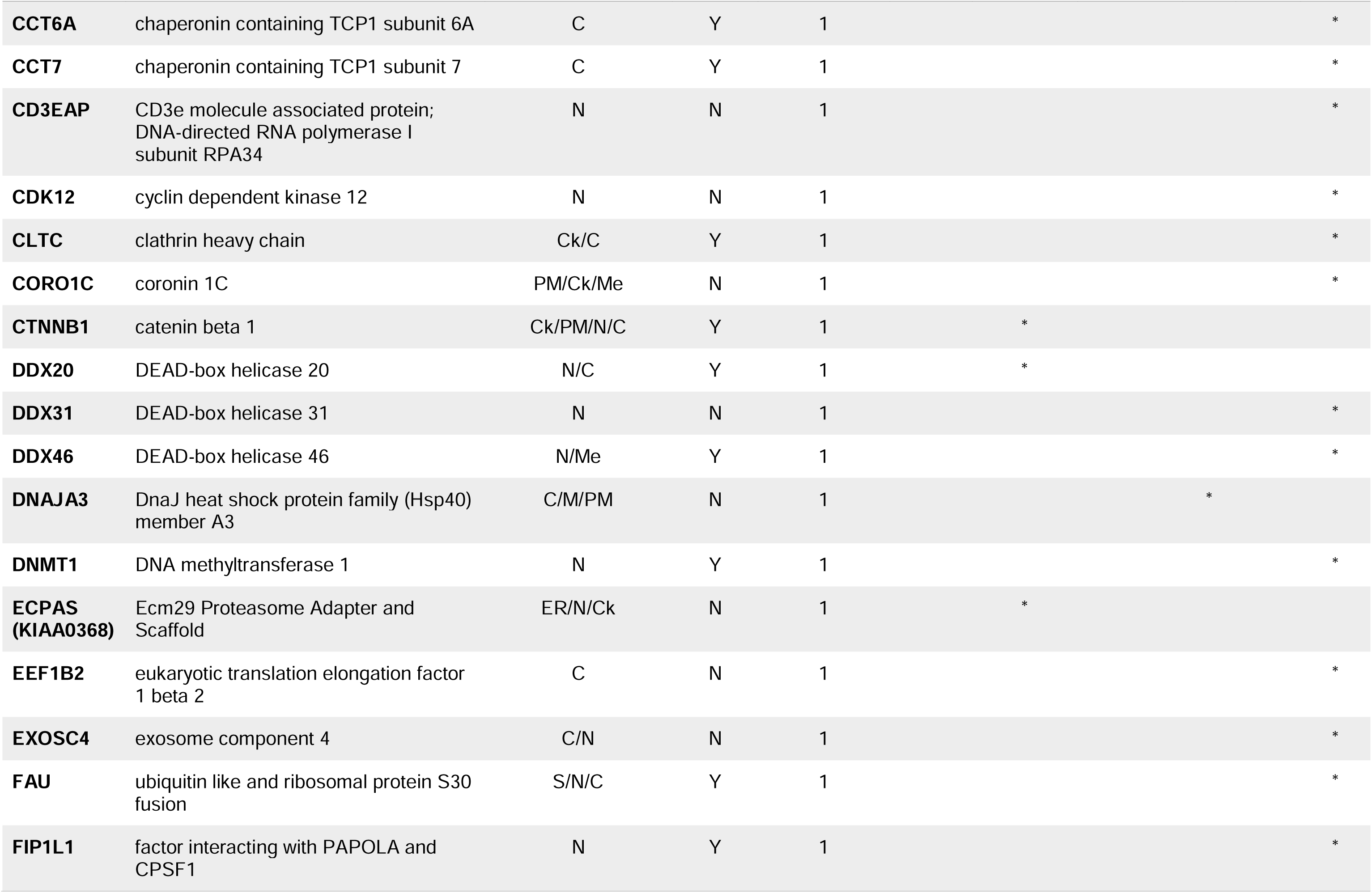
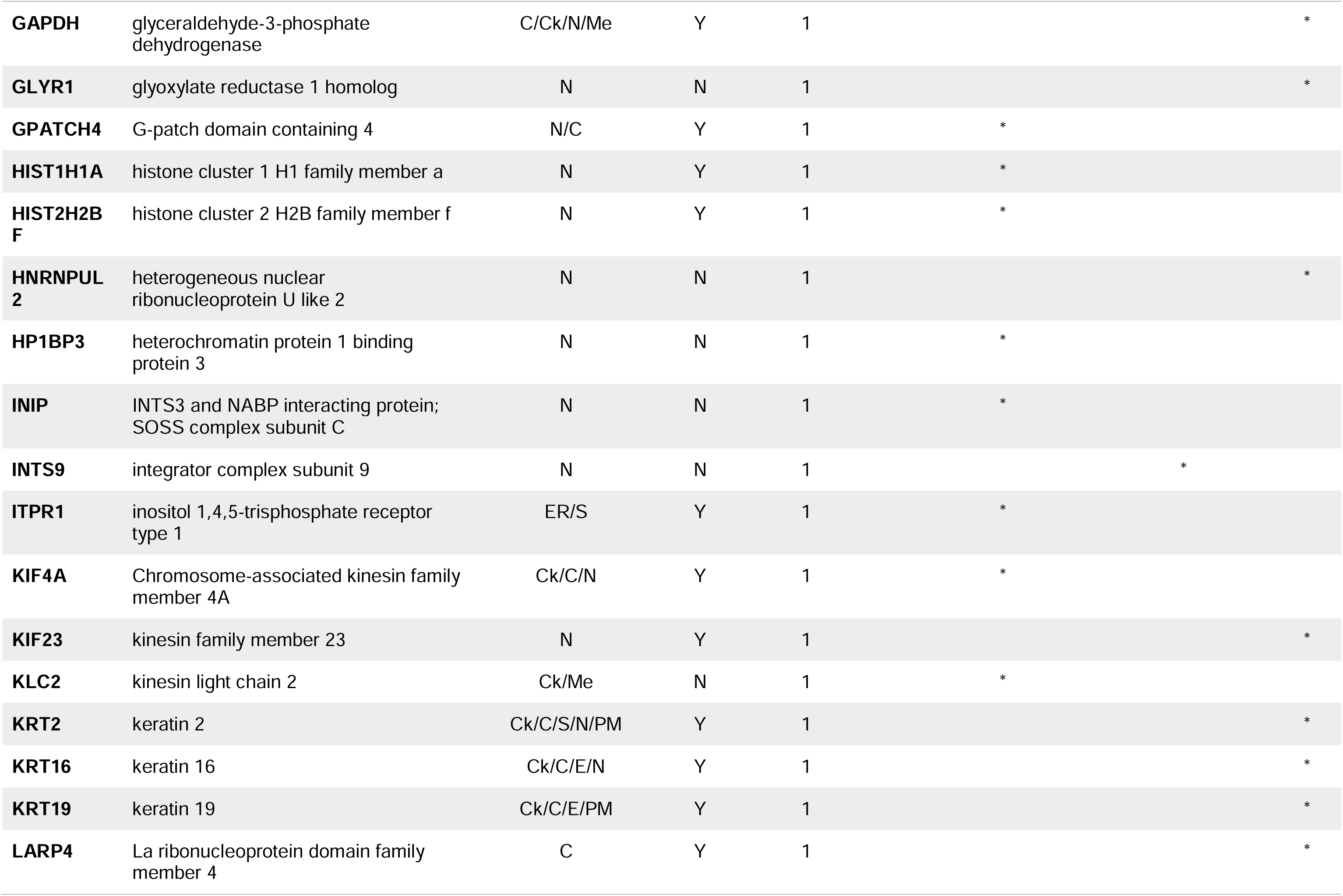
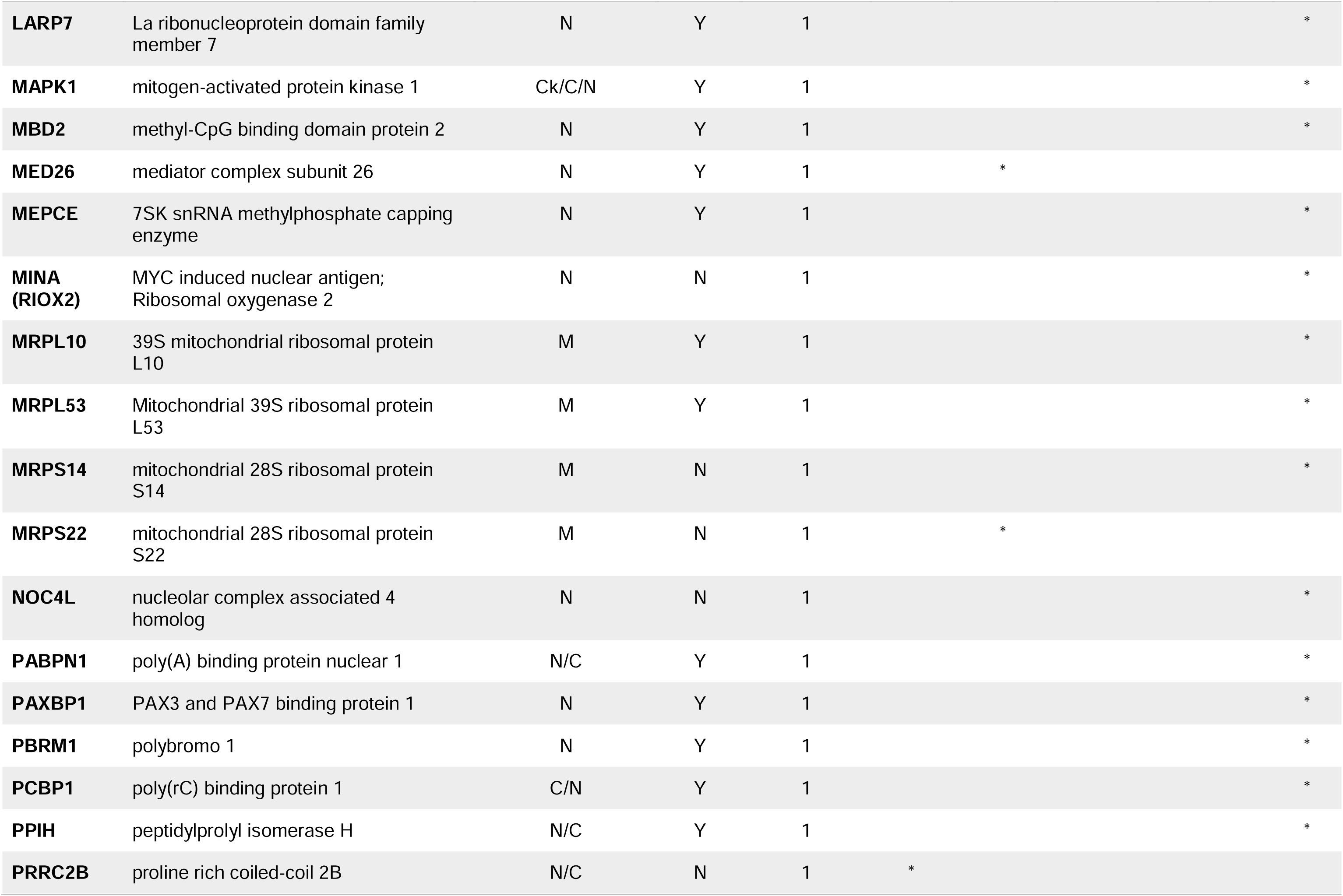
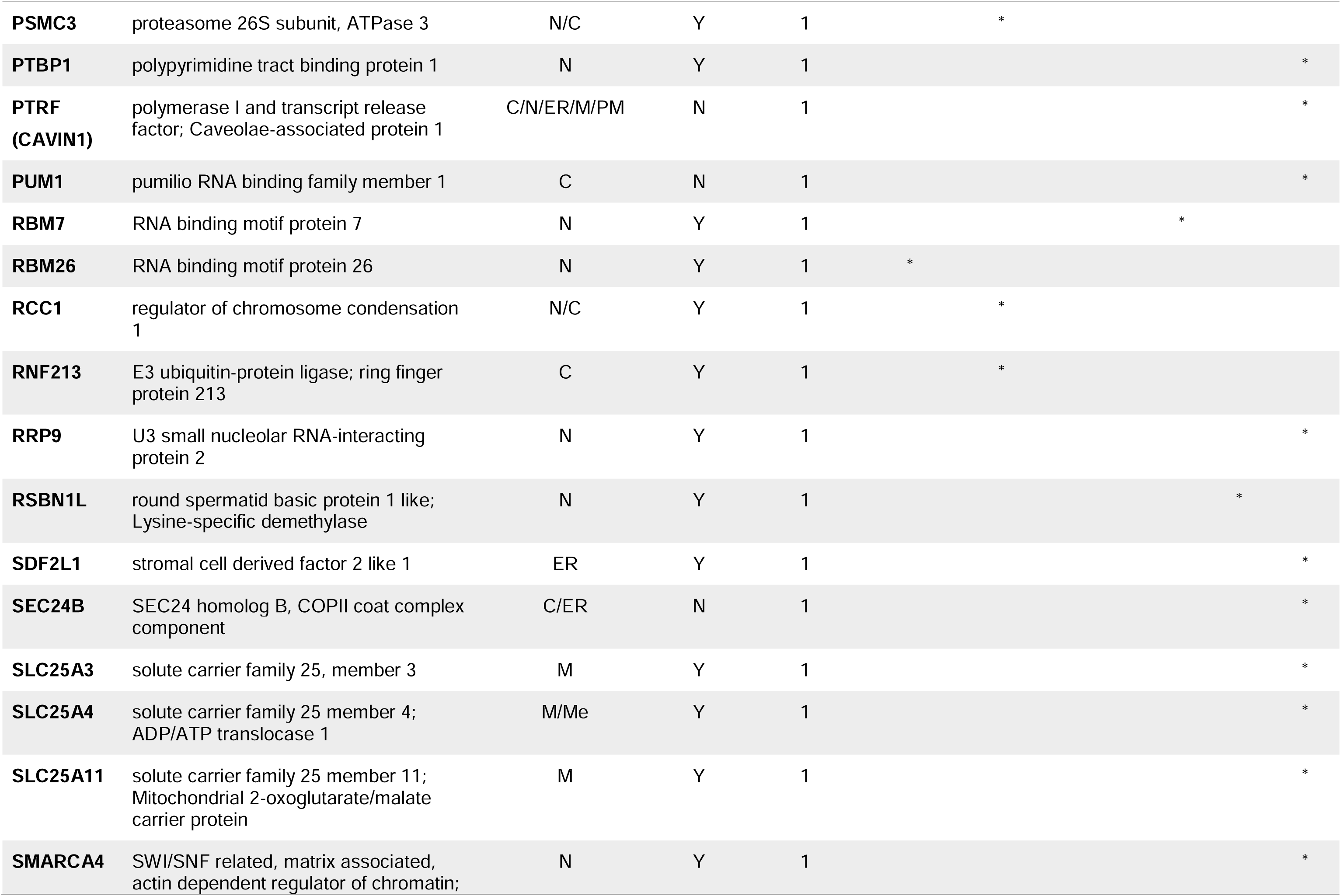
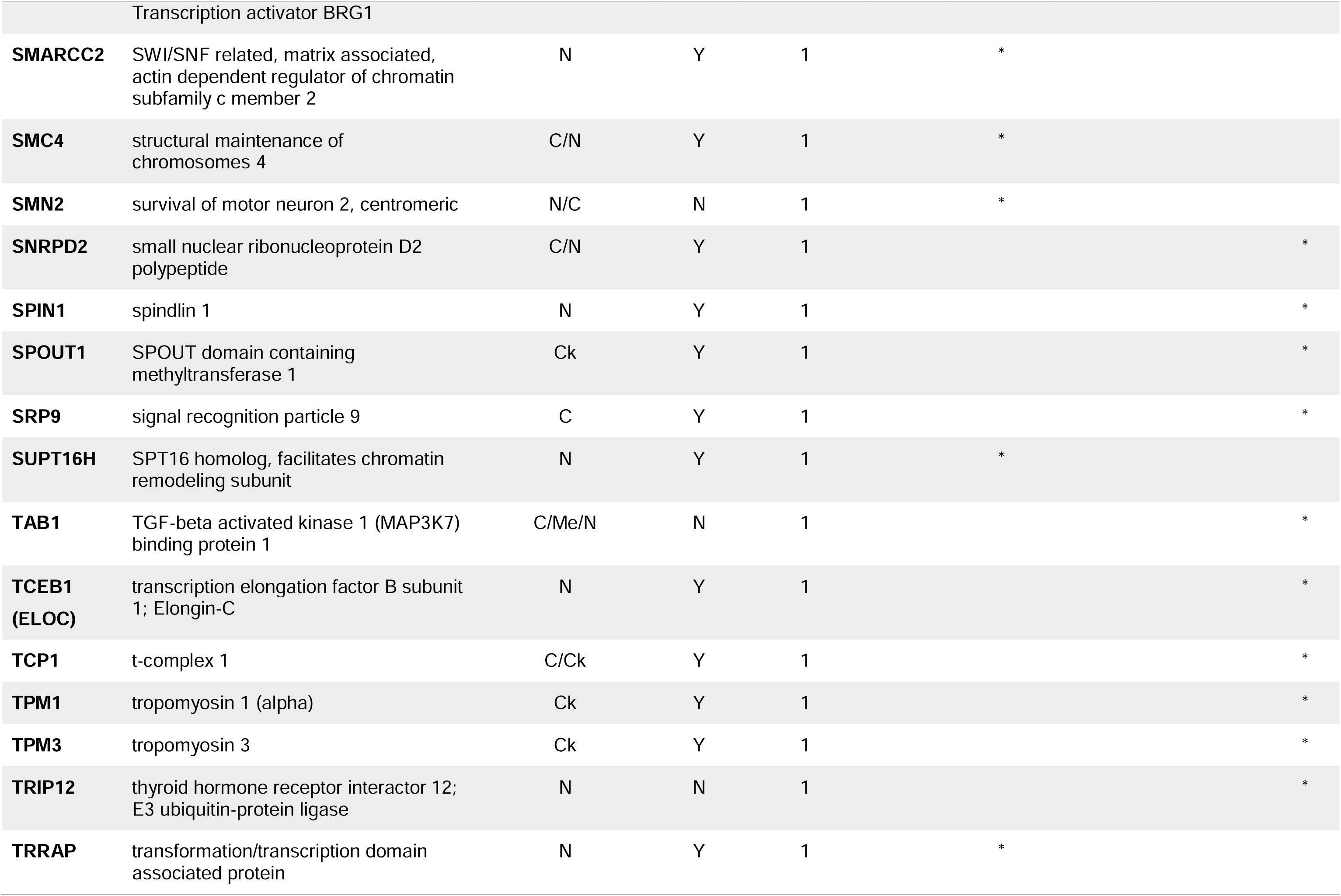
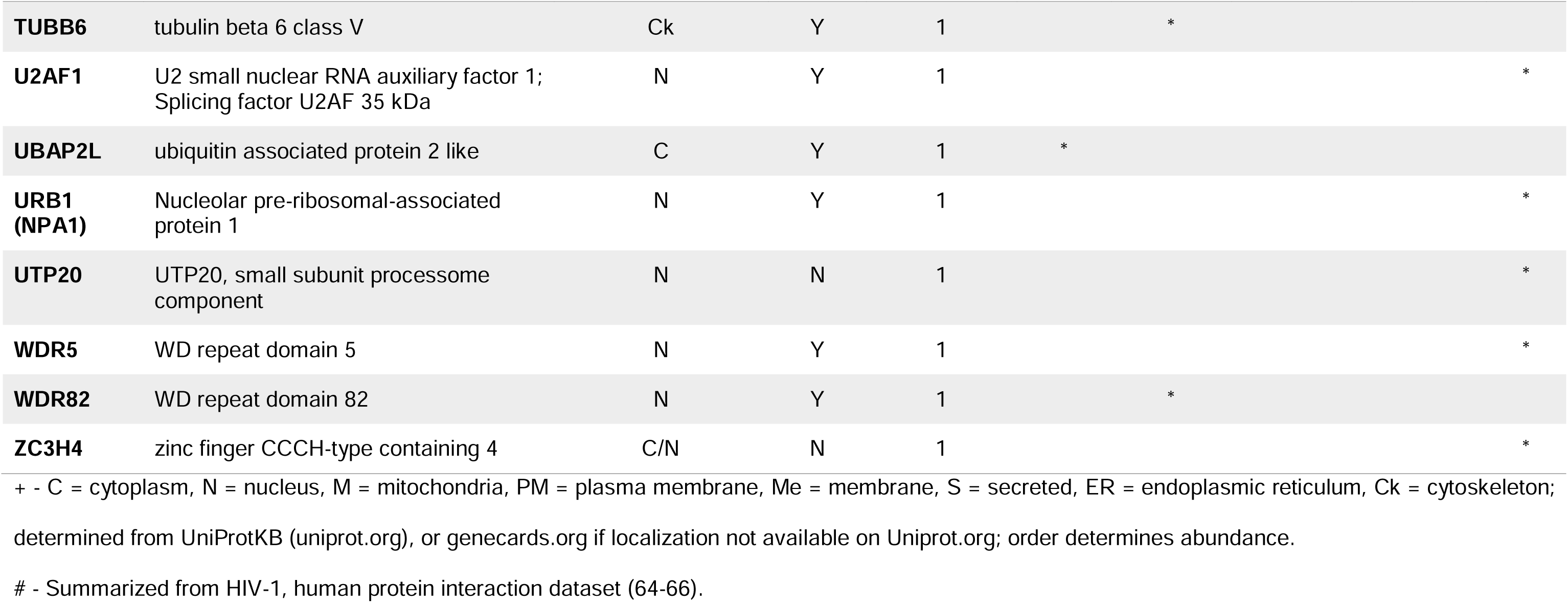
Proteins identified in the present HIV-1 Gag affinity purification and found in at least one of the previously published HIV-1 Gag proteomic publications.

These overlapping proteins were analyzed using Ingenuity Pathway Analysis (IPA) (**Fig 3**), and the interacting proteins found in this report and at least one of the previously published lists were categorized based on their functions. **Figure 3** shows the distribution of the proteins among 13 basic categories identified by Lippé (57). Proteins may belong to more than one functional category, which results in the number of proteins present in each category exceeding the total number of proteins. To address this issue, the number of proteins in each category was divided by the total number of proteins to yield a percentage for each of the categories represented in the protein list analyzed. **Figure 3** displays the relative values of the Ingenuity Pathway Analysis rather than the raw data.

**Figure 3.**
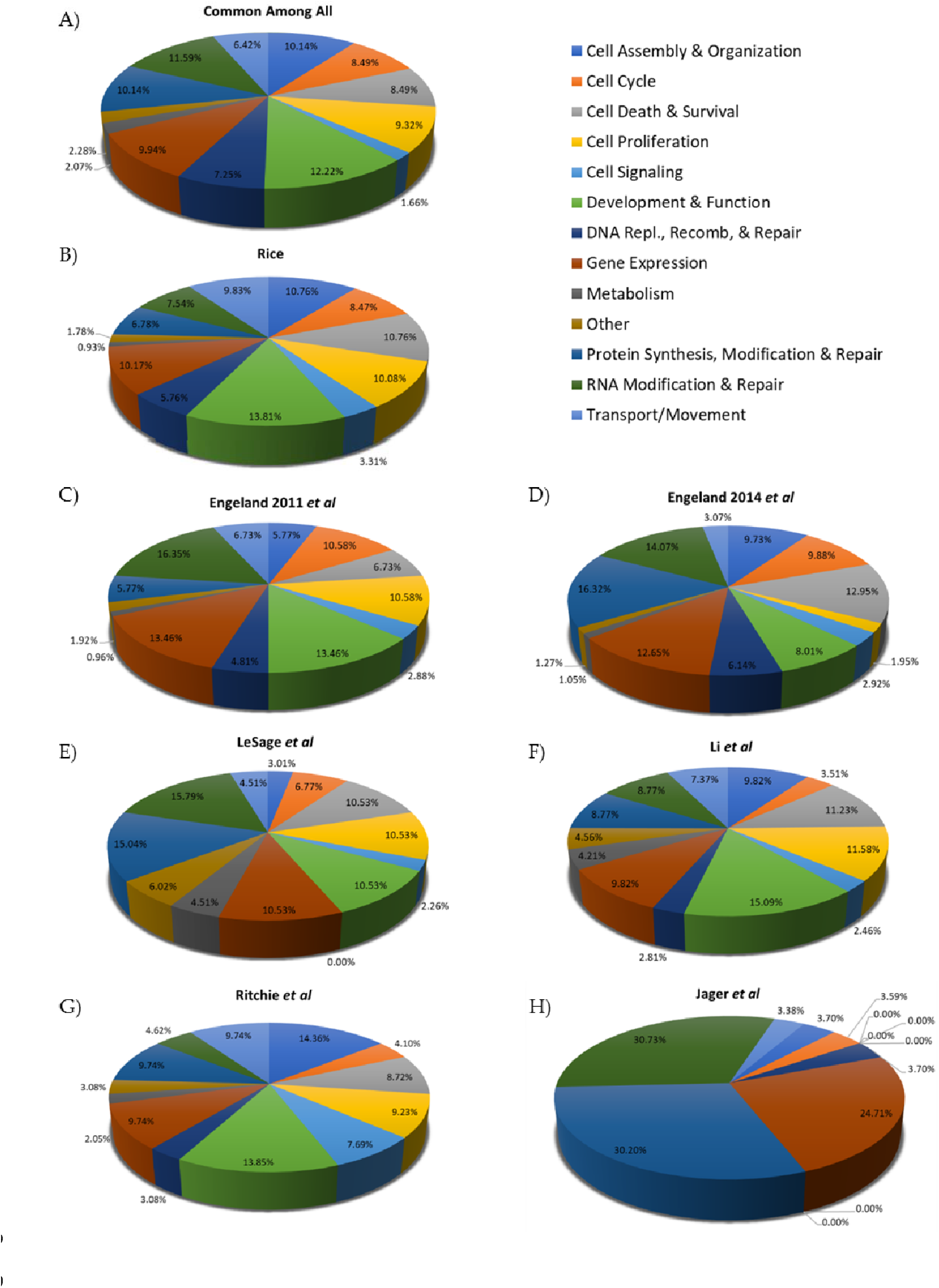
The relative representation of the IPA categories present in each protein list. (**A**) The proteins identified in our HIV-1 Gag pulldown that were also found in at least one of the previously published reports were analyzed for molecular and cellular functions. The color key on the right is the same for each pie graph. Protein functions identified in this publication (Rice) (**B**), Engeland 2011 (31)(**C**), Engeland 2014 (30)(**D**), Le Sage (33)(**E**), Li (34)(**F**), Ritchie (35)(**G**), and Jäger (32)(**H**).

### 3.4. Closer examination of Gag interactions with host proteins involved in transcription and splicing

Numerous Gag-interacting proteins were found to be involved in RNA polymerase II transcription (GO term: 0006366 transcription from RNA polymerase II promoter) and splicing (GO term: 0008380 RNA splicing). We were interested in these two GO terms because our previous work demonstrated that RSV and HIV-1 Gag localize to the perichromatin space and associate with newly-transcribed USvRNA (8, 9), suggesting that they may interface with transcription and splicing processes. In addition, in Rice *et al.*, we demonstrated that RSV Gag.L219A colocalizes with the splicing factors SF2 and SC35, and has similar mobility and dynamics as proteins residing in splicing speckles (67). Maldonado *et al.* subsequently showed that RSV Gag forms discrete nuclear foci and interacts with USvRNA at active transcription sites (8). Taken together with the proteomic data presented here, it is reasonable to propose that RSV Gag interacts with cellular proteins involved in transcription and co-transcriptional processes such as splicing and RNA processing, at or near transcription sites. **Tables 3** and **4** list the proteins involved in transcription and splicing, respectively, for the RSV pulldowns. Similarly, Tuffy *et al.* and Chang *et al.* (9, 10) both demonstrated that HIV-1 Gag localizes to transcriptionally active regions in HeLa cells and T cells reactivated from latency. Given these findings, it is feasible that HIV-1 Gag interacts with host nuclear factors involved in transcription, RNA processing, and chromatin remodeling. **Tables 5** and **6** list the proteins in these categories identified in our HIV-1 pulldowns and indicate whether each protein was identified in any of the other publications. Of note, proteins involved in processes such as splicing are included in the DAVID category of transcription interactome (68, 69) because transcription and splicing have been shown to be linked and occur simultaneously (70–73).

**Table 3.**
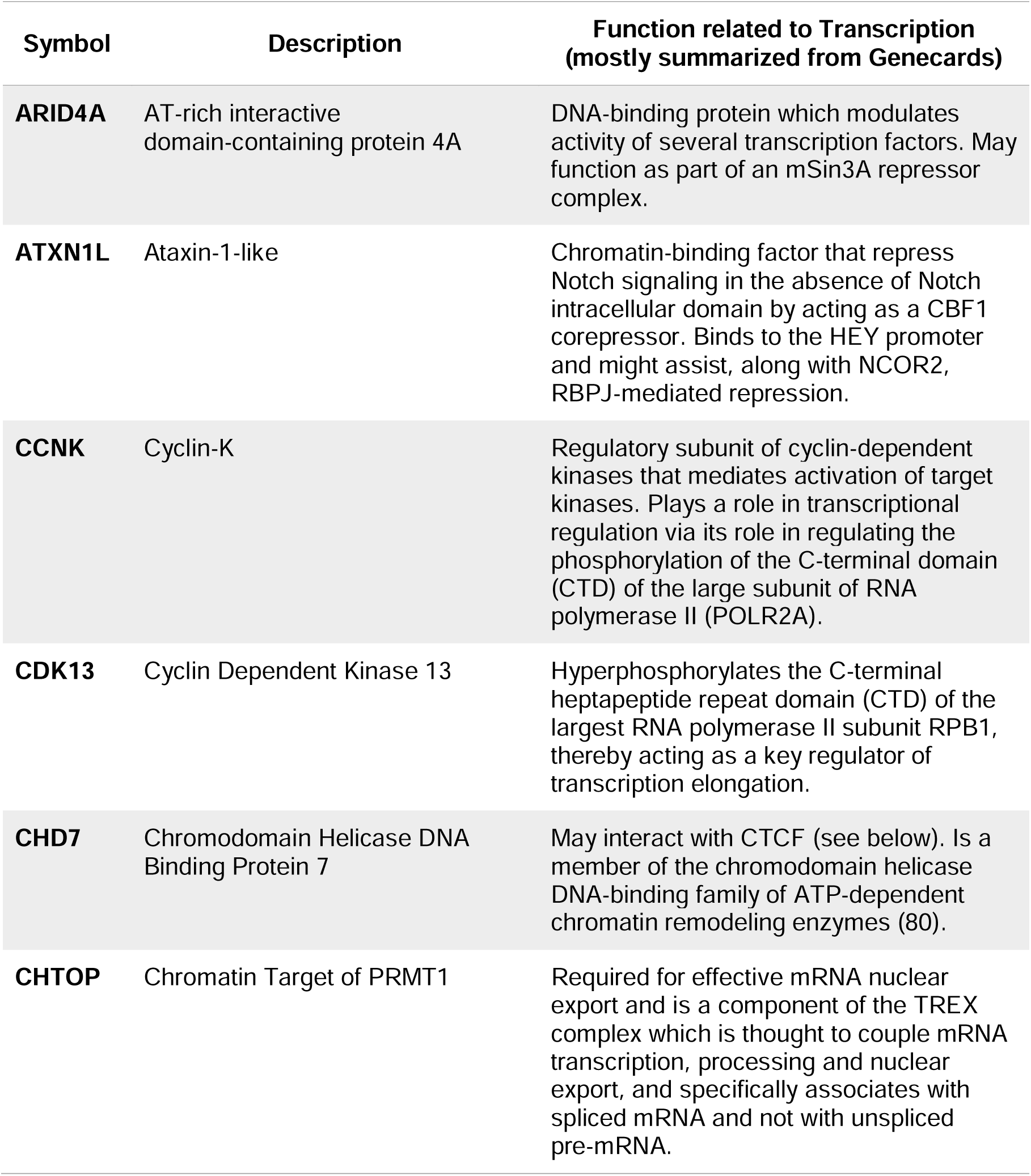
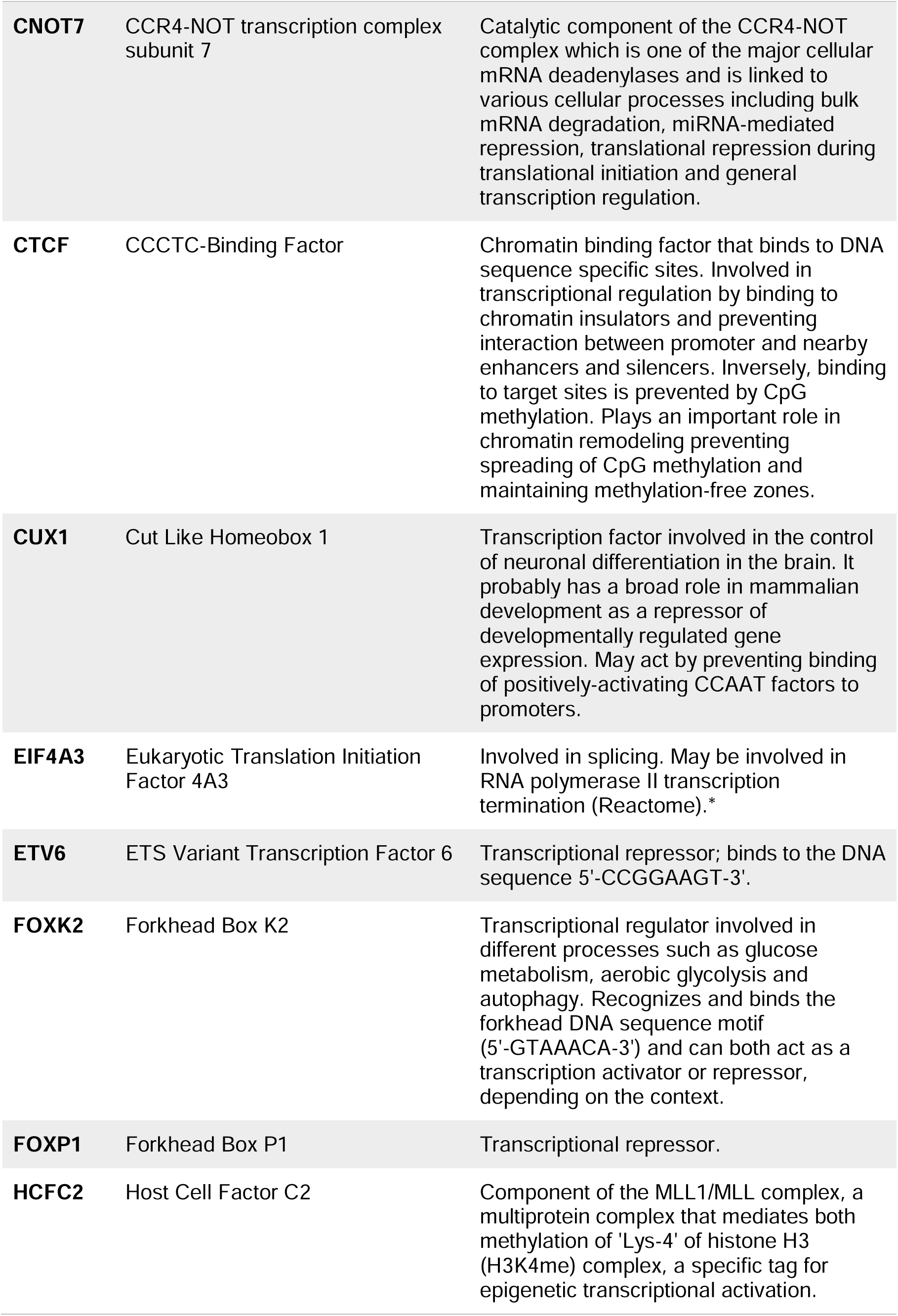
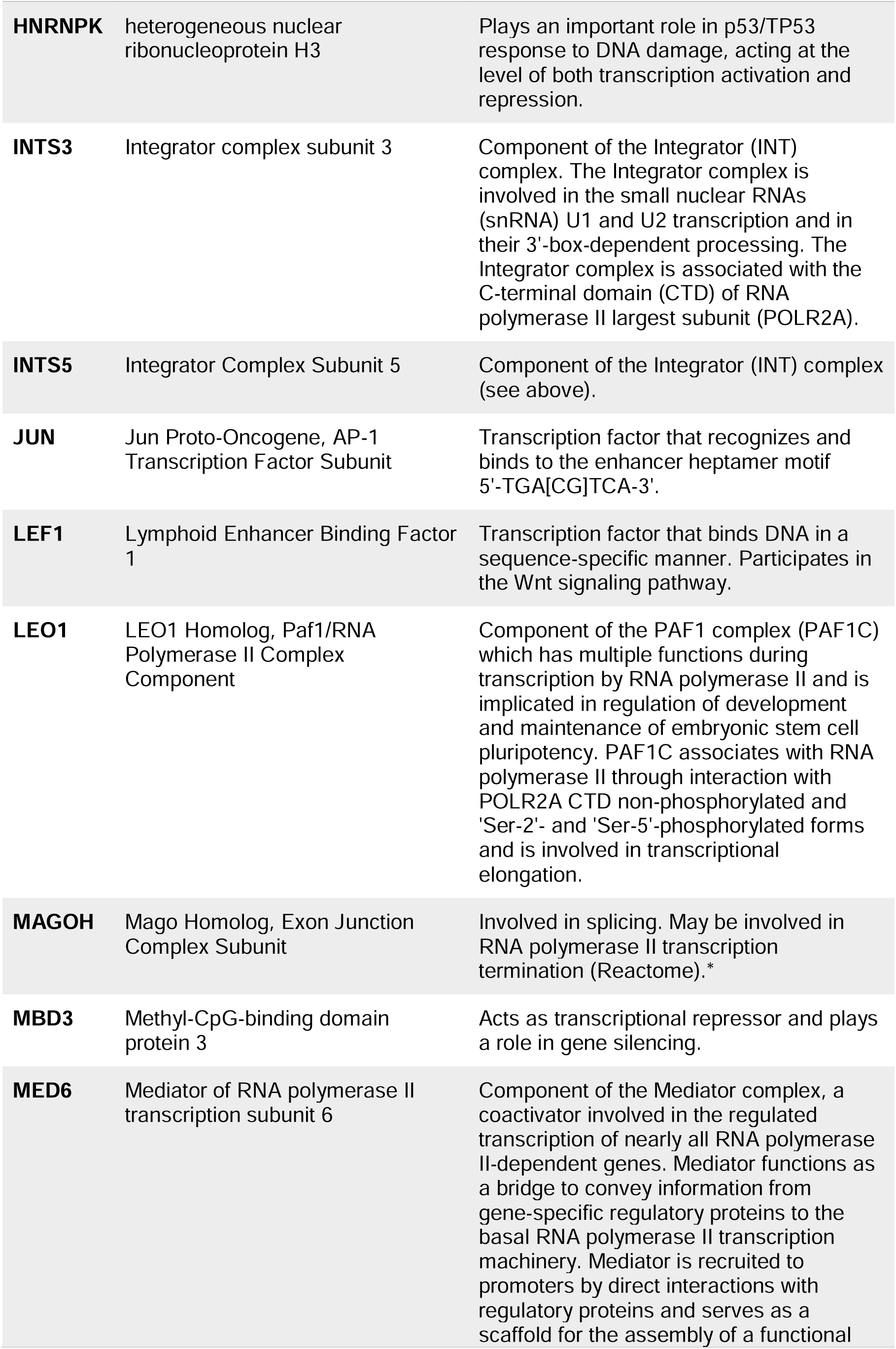
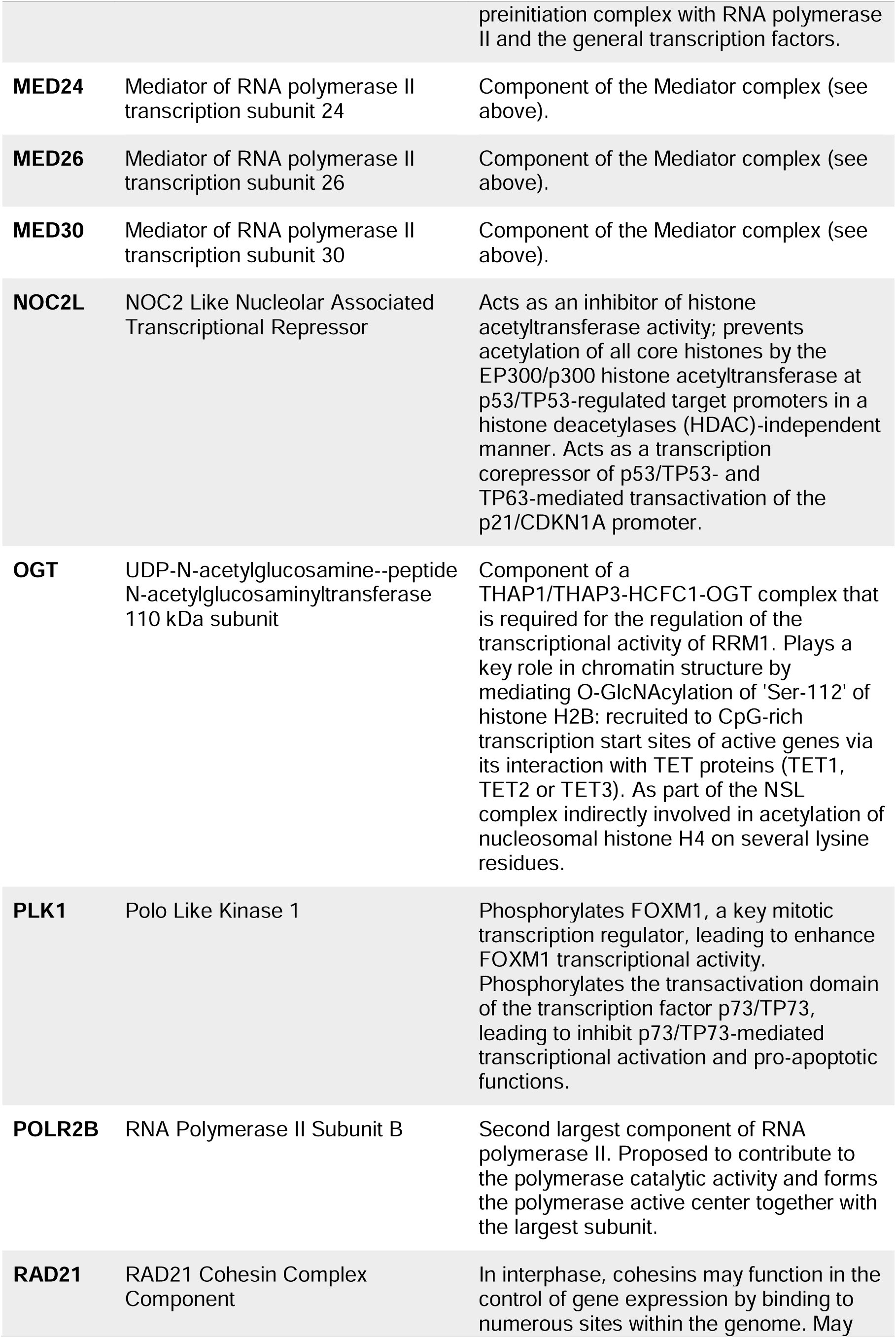
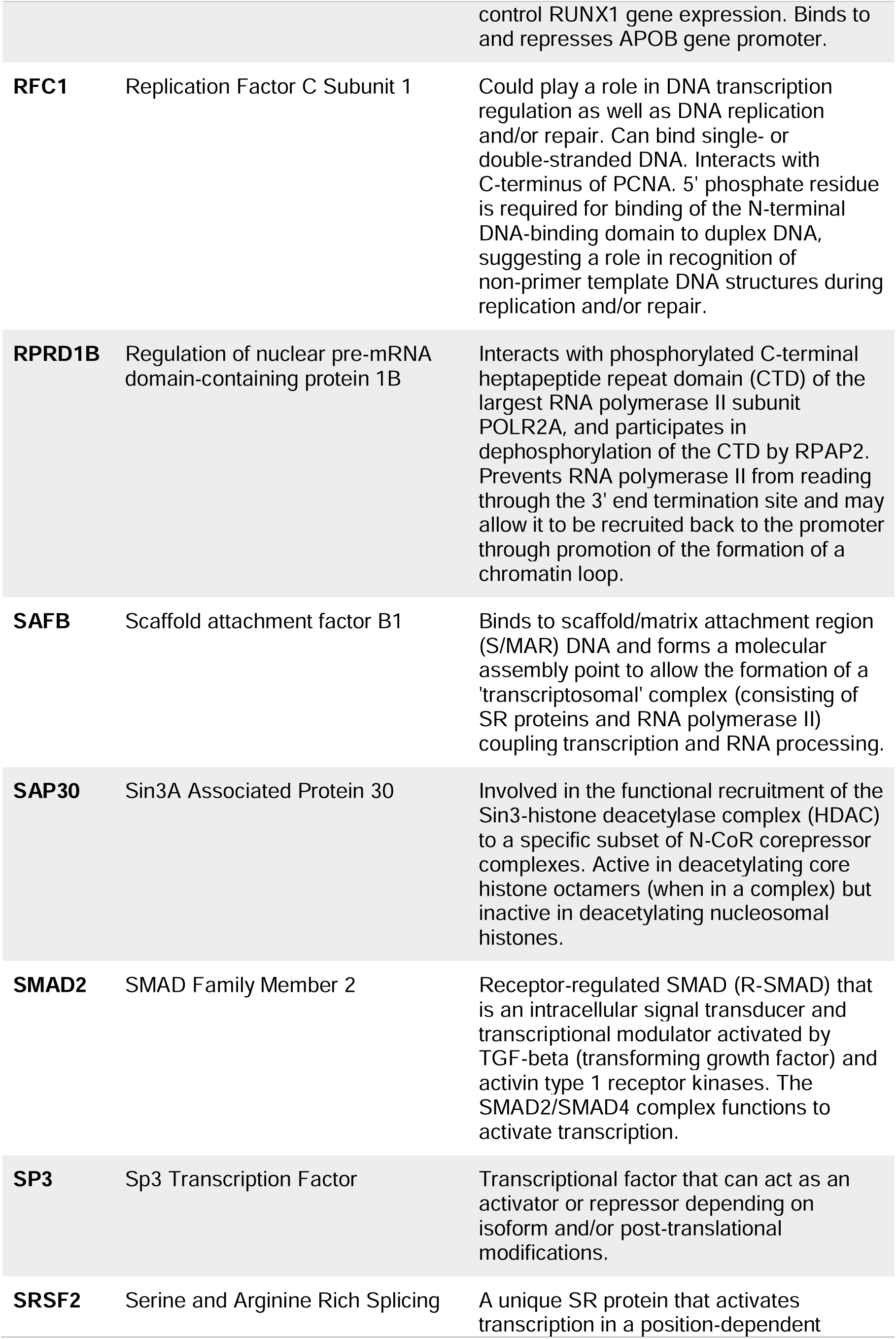
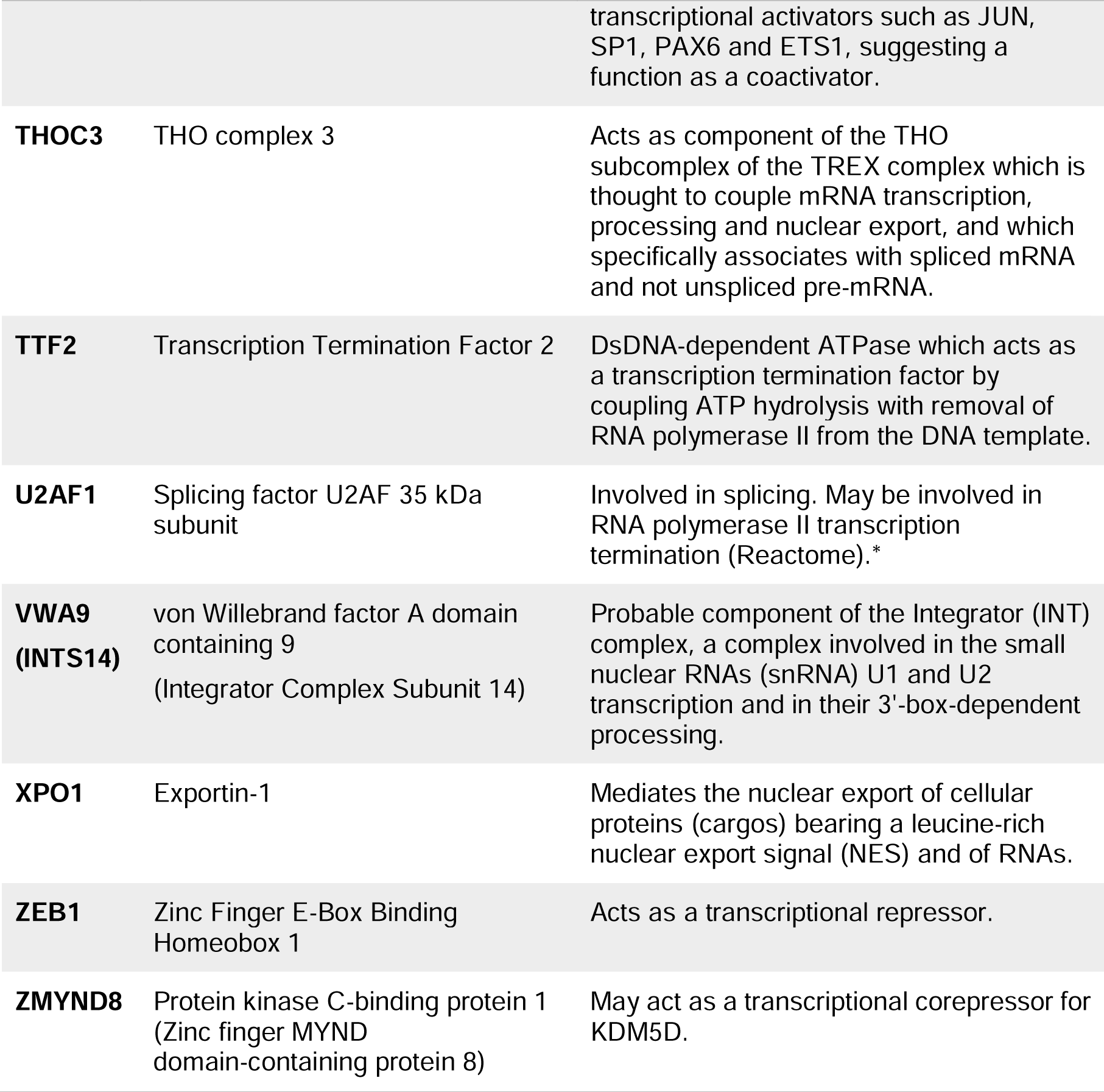
Names and functions of the proteins identified in the RSV proteomics list under GO:0006366∼ transcription from RNA polymerase II promoter.

**Table 4.**
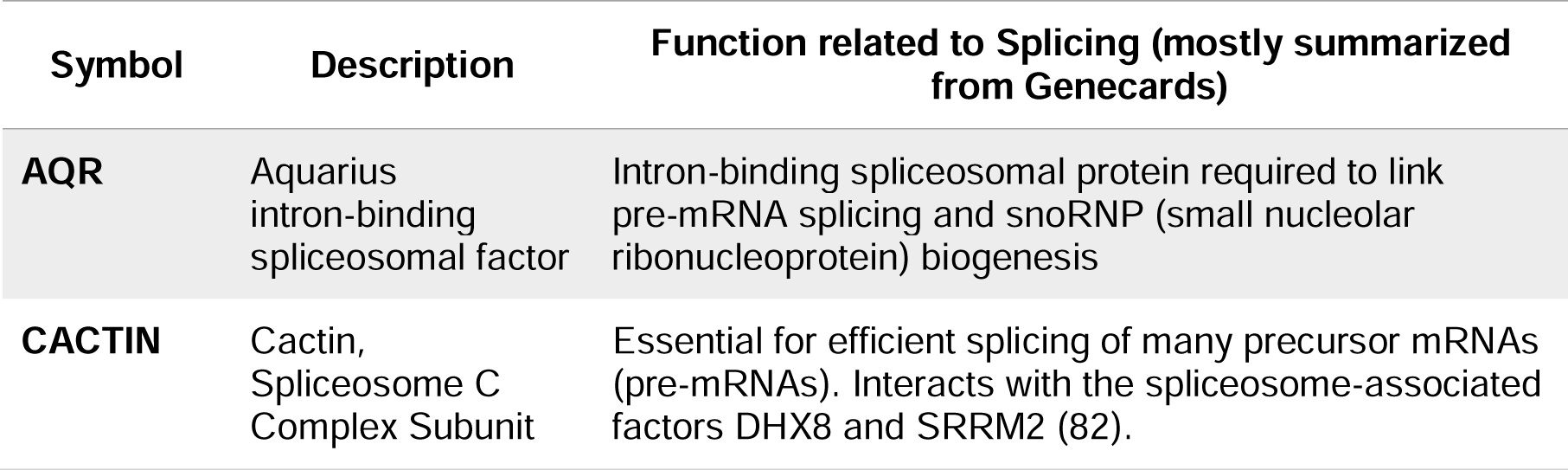
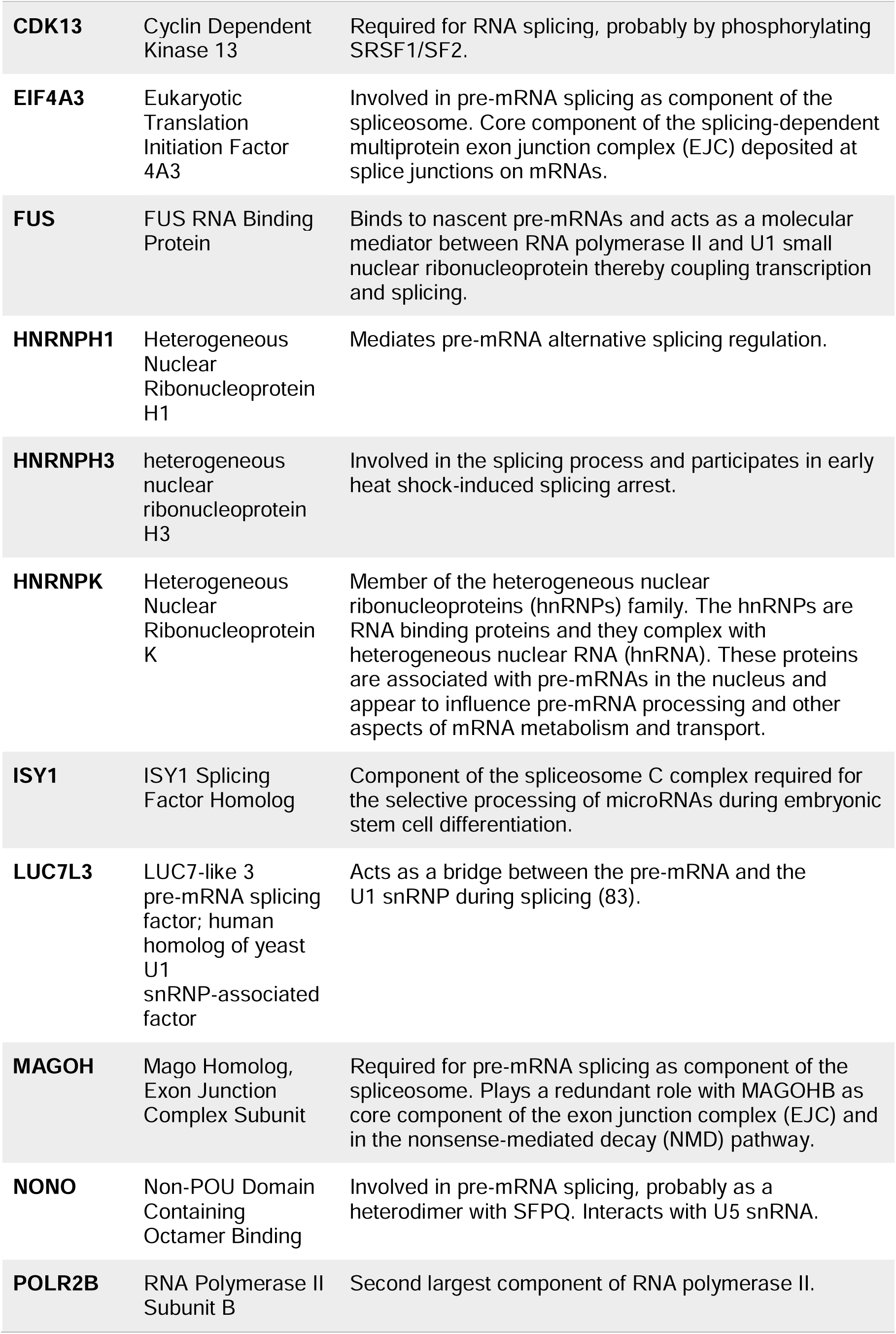
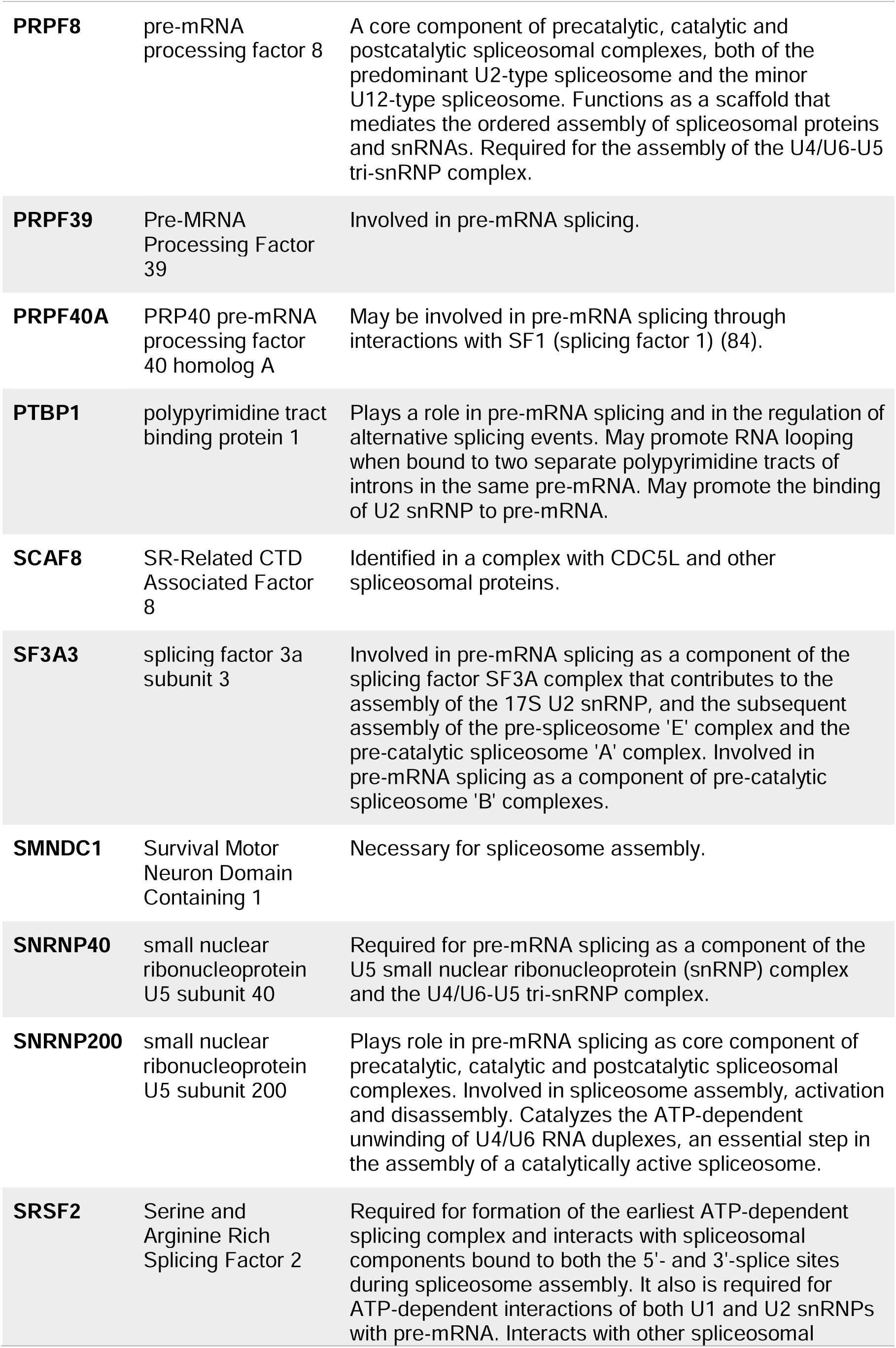
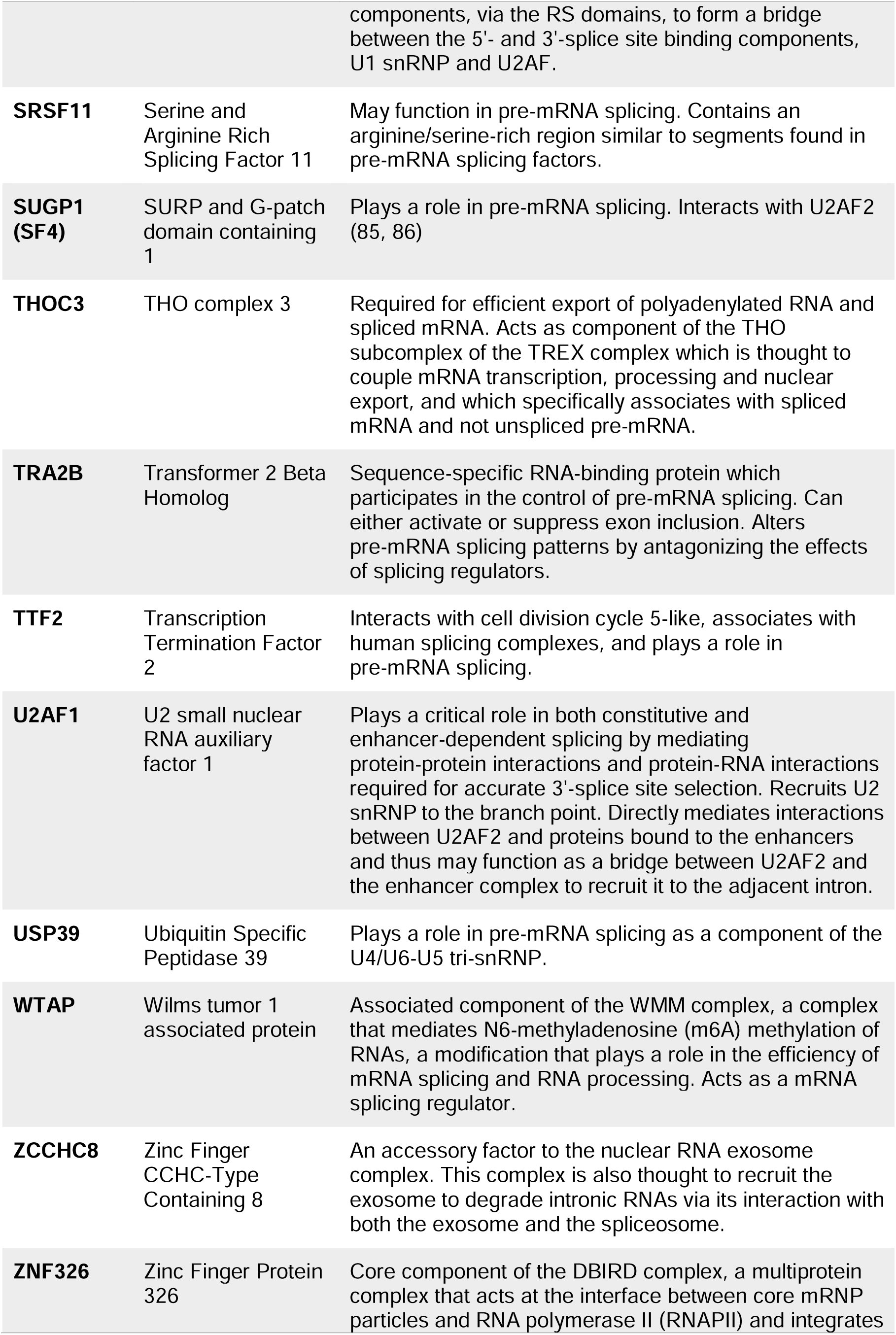
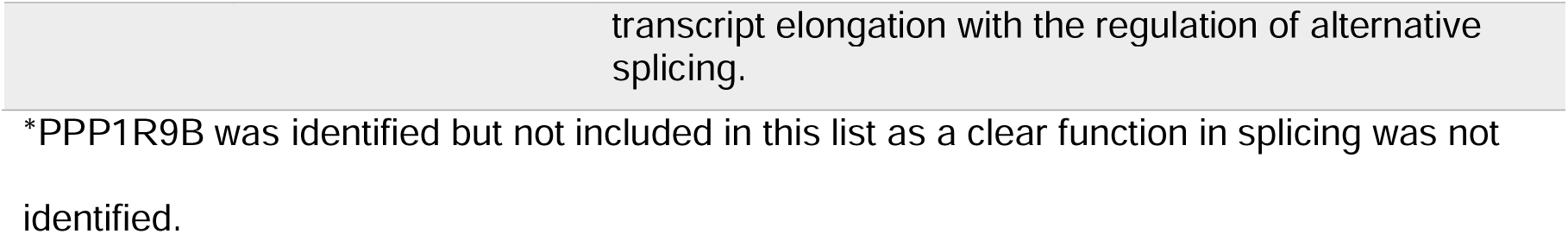
Names and functions of the proteins identified in the RSV proteomics list under GO:0008380∼RNA splicing.

| Symbol | Description | Function related to Splicing (mostly summarized from Genecards) |
| --- | --- | --- |
| **AQR** | Aquarius intron-binding spliceosomal factor | Intron-binding spliceosomal protein required to link pre-mRNA splicing and snoRNP (small nucleolar ribonucleoprotein) biogenesis |
| **CACTIN** | Cactin, Spliceosome C Complex Subunit | Essential for efficient splicing of many precursor mRNAs (pre-mRNAs). Interacts with the spliceosome-associated factors DHX8 and SRRM2 (82). |
| **CDK13** | Cyclin Dependent Kinase 13 | Required for RNA splicing, probably by phosphorylating SRSF1/SF2. |
| **EIF4A3** | Eukaryotic Translation Initiation Factor 4A3 | Involved in pre-mRNA splicing as component of the spliceosome. Core component of the splicing-dependent multiprotein exon junction complex (EJC) deposited at splice junctions on mRNAs. |
| **FUS** | FUS RNA Binding Protein | Binds to nascent pre-mRNAs and acts as a molecular mediator between RNA polymerase II and U1 small nuclear ribonucleoprotein thereby coupling transcription and splicing. |
| **HNRNPH1** | Heterogeneous Nuclear Ribonucleoprotein H1 | Mediates pre-mRNA alternative splicing regulation. |
| **HNRNPH3** | heterogeneous nuclear ribonucleoprotein H3 | Involved in the splicing process and participates in early heat shock-induced splicing arrest. |
| **HNRNPK** | Heterogeneous Nuclear Ribonucleoprotein K | Member of the heterogeneous nuclear ribonucleoproteins (hnRNPs) family. The hnRNPs are RNA binding proteins and they complex with heterogeneous nuclear RNA (hnRNA). These proteins are associated with pre-mRNAs in the nucleus and appear to influence pre-mRNA processing and other aspects of mRNA metabolism and transport. |
| **ISY1** | ISY1 Splicing Factor Homolog | Component of the spliceosome C complex required for the selective processing of microRNAs during embryonic stem cell differentiation. |
| **LUC7L3** | LUC7-like 3 pre-mRNA splicing factor; human homolog of yeast U1 snRNP-associated factor | Acts as a bridge between the pre-mRNA and the U1 snRNP during splicing (83). |
| **MAGOH** | Mago Homolog, Exon Junction Complex Subunit | Required for pre-mRNA splicing as component of the spliceosome. Plays a redundant role with MAGOHB as core component of the exon junction complex (EJC) and in the nonsense-mediated decay (NMD) pathway. |
| **NONO** | Non-POU Domain Containing Octamer Binding | Involved in pre-mRNA splicing, probably as a heterodimer with SFPQ. Interacts with U5 snRNA. |
| **POLR2B** | RNA Polymerase II Subunit B | Second largest component of RNA polymerase II. |
| **PRPF8** | pre-mRNA processing factor 8 | A core component of precatalytic, catalytic and postcatalytic spliceosomal complexes, both of the predominant U2-type spliceosome and the minor U12-type spliceosome. Functions as a scaffold that mediates the ordered assembly of spliceosomal proteins and snRNAs. Required for the assembly of the U4/U6-U5 tri-snRNP complex. |
| **PRPF39** | Pre-mRNA Processing Factor 39 | Involved in pre-mRNA splicing. |
| **PRPF40A** | PRP40 pre-mRNA processing factor 40 homolog A | May be involved in pre-mRNA splicing through interactions with SF1 (splicing factor 1) (84). |
| **PTBP1** | polypyrimidine tract binding protein 1 | Plays a role in pre-mRNA splicing and in the regulation of alternative splicing events. May promote RNA looping when bound to two separate polypyrimidine tracts of introns in the same pre-mRNA. May promote the binding of U2 snRNP to pre-mRNA. |
| **SCAF8** | SR-Related CTD Associated Factor 8 | Identified in a complex with CDC5L and other spliceosomal proteins. |
| **SF3A3** | splicing factor 3a subunit 3 | Involved in pre-mRNA splicing as a component of the splicing factor SF3A complex that contributes to the assembly of the 17S U2 snRNP, and the subsequent assembly of the pre-spliceosome 'E' complex and the pre-catalytic spliceosome 'A' complex. Involved in pre-mRNA splicing as a component of pre-catalytic spliceosome 'B' complexes. |
| **SMNDC1** | Survival Motor Neuron Domain Containing 1 | Necessary for spliceosome assembly. |
| **SNRNP40** | small nuclear ribonucleoprotein U5 subunit 40 | Required for pre-mRNA splicing as a component of the U5 small nuclear ribonucleoprotein (snRNP) complex and the U4/U6-U5 tri-snRNP complex. |
| **SNRNP200** | small nuclear ribonucleoprotein U5 subunit 200 | Plays role in pre-mRNA splicing as core component of precatalytic, catalytic and postcatalytic spliceosomal complexes. Involved in spliceosome assembly, activation and disassembly. Catalyzes the ATP-dependent unwinding of U4/U6 RNA duplexes, an essential step in the assembly of a catalytically active spliceosome. |
| **SRSF2** | Serine and Arginine Rich Splicing Factor 2 | Required for formation of the earliest ATP-dependent splicing complex and interacts with spliceosomal components bound to both the 5'- and 3'-splice sites during spliceosome assembly. It also is required for ATP-dependent interactions of both U1 and U2 snRNPs with pre-mRNA. Interacts with other spliceosomal |
|  |  | components, via the RS domains, to form a bridge between the 5'- and 3'-splice site binding components, U1 snRNP and U2AF. |
| **SRSF11** | Serine and Arginine Rich Splicing Factor 11 | May function in pre-mRNA splicing. Contains an arginine/serine-rich region similar to segments found in pre-mRNA splicing factors. |
| **SUGP1 (SF4)** | SURP and G-patch domain containing 1 | Plays a role in pre-mRNA splicing. Interacts with U2AF2 (85, 86) |
| **THOC3** | THO complex 3 | Required for efficient export of polyadenylated RNA and spliced mRNA. Acts as component of the THO subcomplex of the TREX complex which is thought to couple mRNA transcription, processing and nuclear export, and which specifically associates with spliced mRNA and not unspliced pre-mRNA. |
| **TRA2B** | Transformer 2 Beta Homolog | Sequence-specific RNA-binding protein which participates in the control of pre-mRNA splicing. Can either activate or suppress exon inclusion. Alters pre-mRNA splicing patterns by antagonizing the effects of splicing regulators. |
| **TTF2** | Transcription Termination Factor 2 | Interacts with cell division cycle 5-like, associates with human splicing complexes, and plays a role in pre-mRNA splicing. |
| **U2AF1** | U2 small nuclear RNA auxiliary factor 1 | Plays a critical role in both constitutive and enhancer-dependent splicing by mediating protein-protein interactions and protein-RNA interactions required for accurate 3'-splice site selection. Recruits U2 snRNP to the branch point. Directly mediates interactions between U2AF2 and proteins bound to the enhancers and thus may function as a bridge between U2AF2 and the enhancer complex to recruit it to the adjacent intron. |
| **USP39** | Ubiquitin Specific Peptidase 39 | Plays a role in pre-mRNA splicing as a component of the U4/U6-U5 tri-snRNP. |
| **WTAP** | Wilms tumor 1 associated protein | Associated component of the WMM complex, a complex that mediates N6-methyladenosine (m6A) methylation of RNAs, a modification that plays a role in the efficiency of mRNA splicing and RNA processing. Acts as a mRNA splicing regulator. |
| **ZCCHC8** | Zinc Finger CCHC-Type Containing 8 | An accessory factor to the nuclear RNA exosome complex. This complex is also thought to recruit the exosome to degrade intronic RNAs via its interaction with both the exosome and the spliceosome. |
| **ZNF326** | Zinc Finger Protein 326 | Core component of the DBIRD complex, a multiprotein complex that acts at the interface between core mRNP particles and RNA polymerase II (RNAPII) and integrates |

\*PPP1R9B was identified but not included in this list as a clear function in splicing was not
identified.

### 3.5 Validation of RSV Gag-Med26 Interaction

We were intrigued by the finding that our proteomic analyses of interactors with RSV and HIV-1 Gag proteins identified several components of the Mediator complex (RSV: Med6, Med13L, Med22, Med24, Med26, Med30; HIV-1: Med9, Med13, Med15, Med21, Med23, Med26, Med28)(**Tables 3, 5, S1-S4**), a coactivator involved in the regulated transcription of nearly all RNAPII genes (59, 74–77). The presence of multiple Mediator proteins in our dataset raised the likelihood that Gag may interact with this multiprotein complex. Furthermore, Mediator proteins have been shown to be exploited by other viruses and endogenous retroelements (36–43). Interestingly, Med26 and Med30 are both metazoan-specific Mediator proteins, implying that Gag may display selectivity for metazoan-specific Mediator complexes over those with protozoan orthologs. Med26 and Med30 both also have critical roles in transcription, with Med26 responsible for the recruitment of important elongation factors to sites of transcription, and Med30 providing stabilization of the intact Mediator core (78, 79).

**Table 5.**
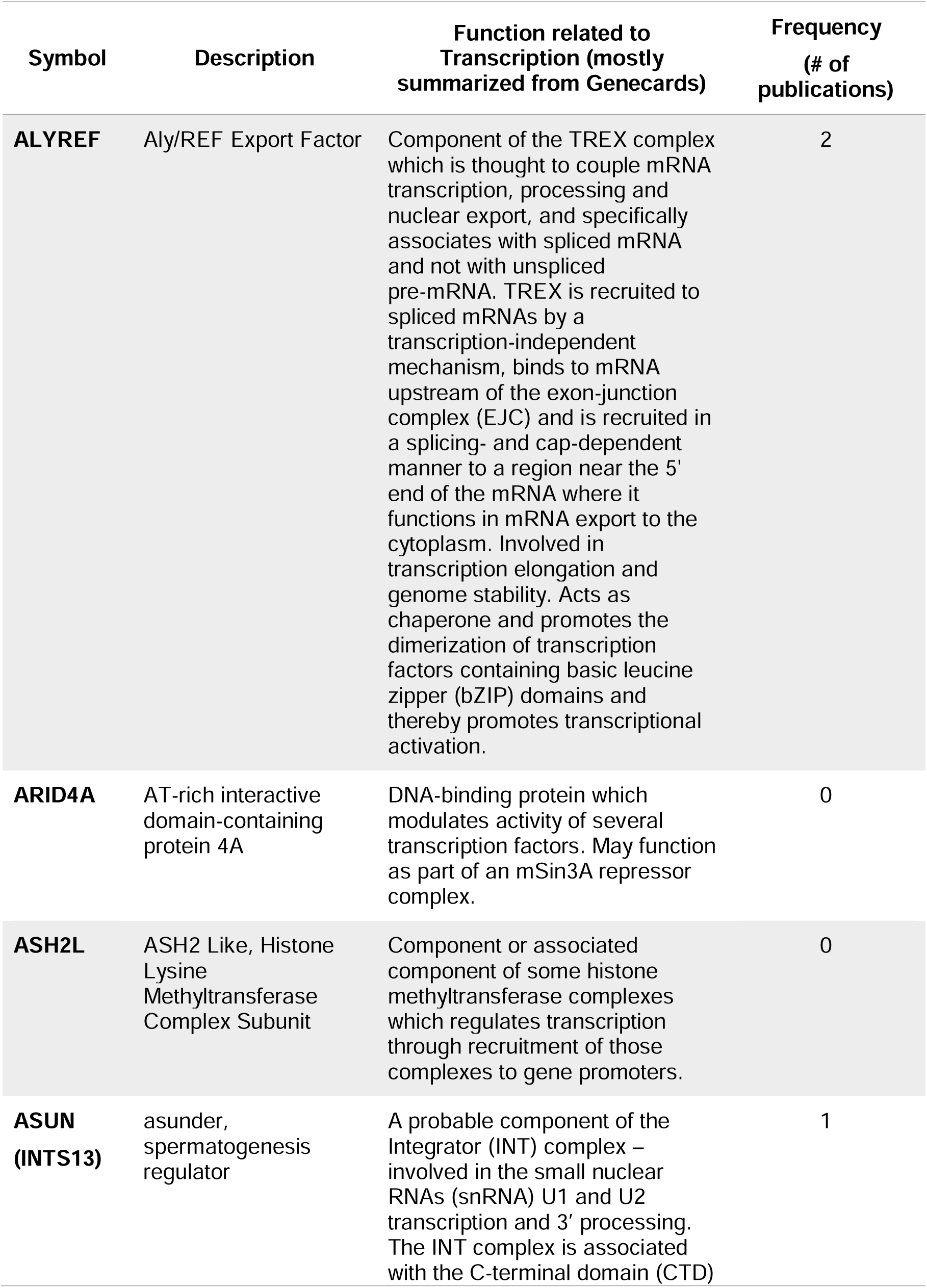
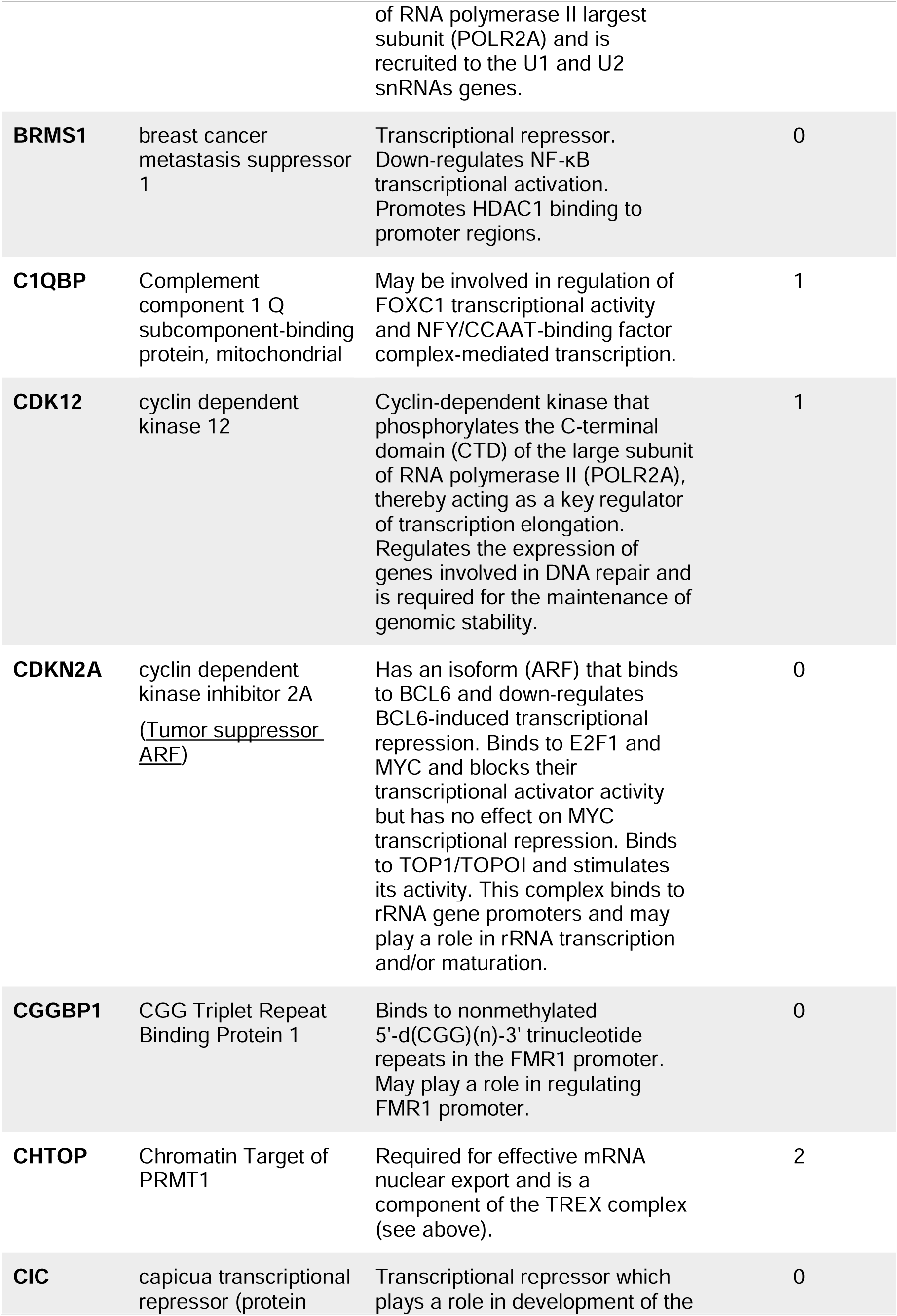
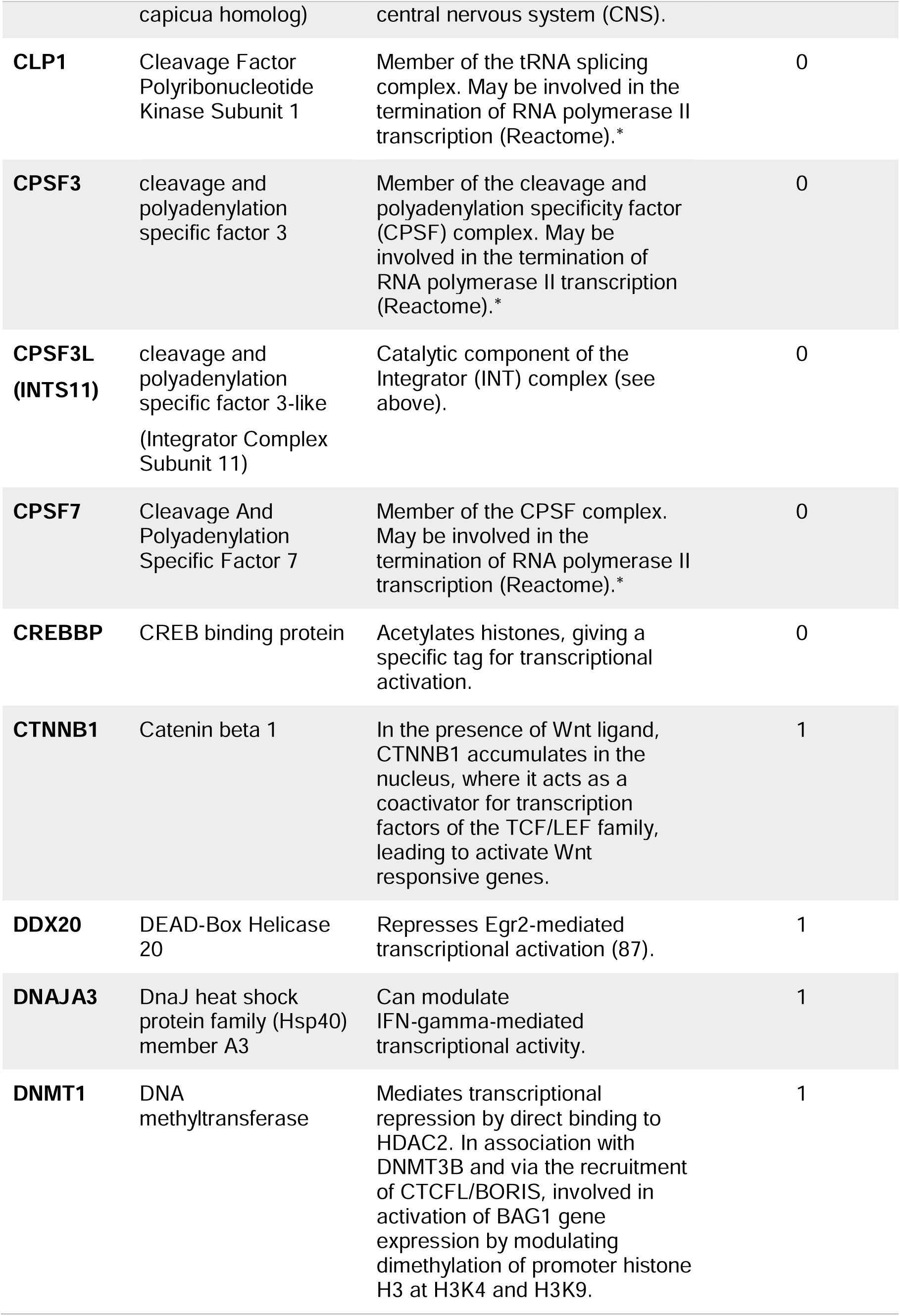
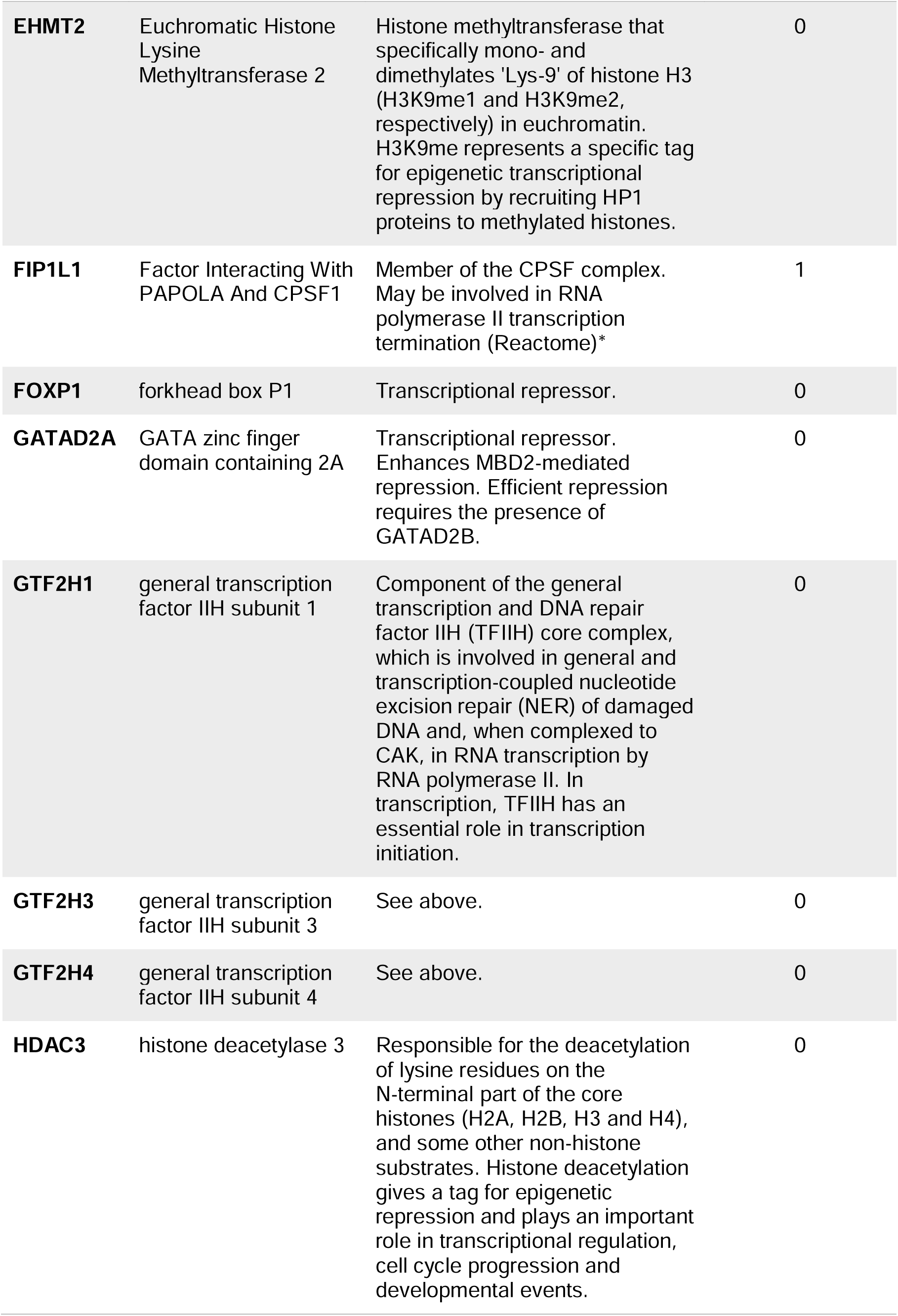
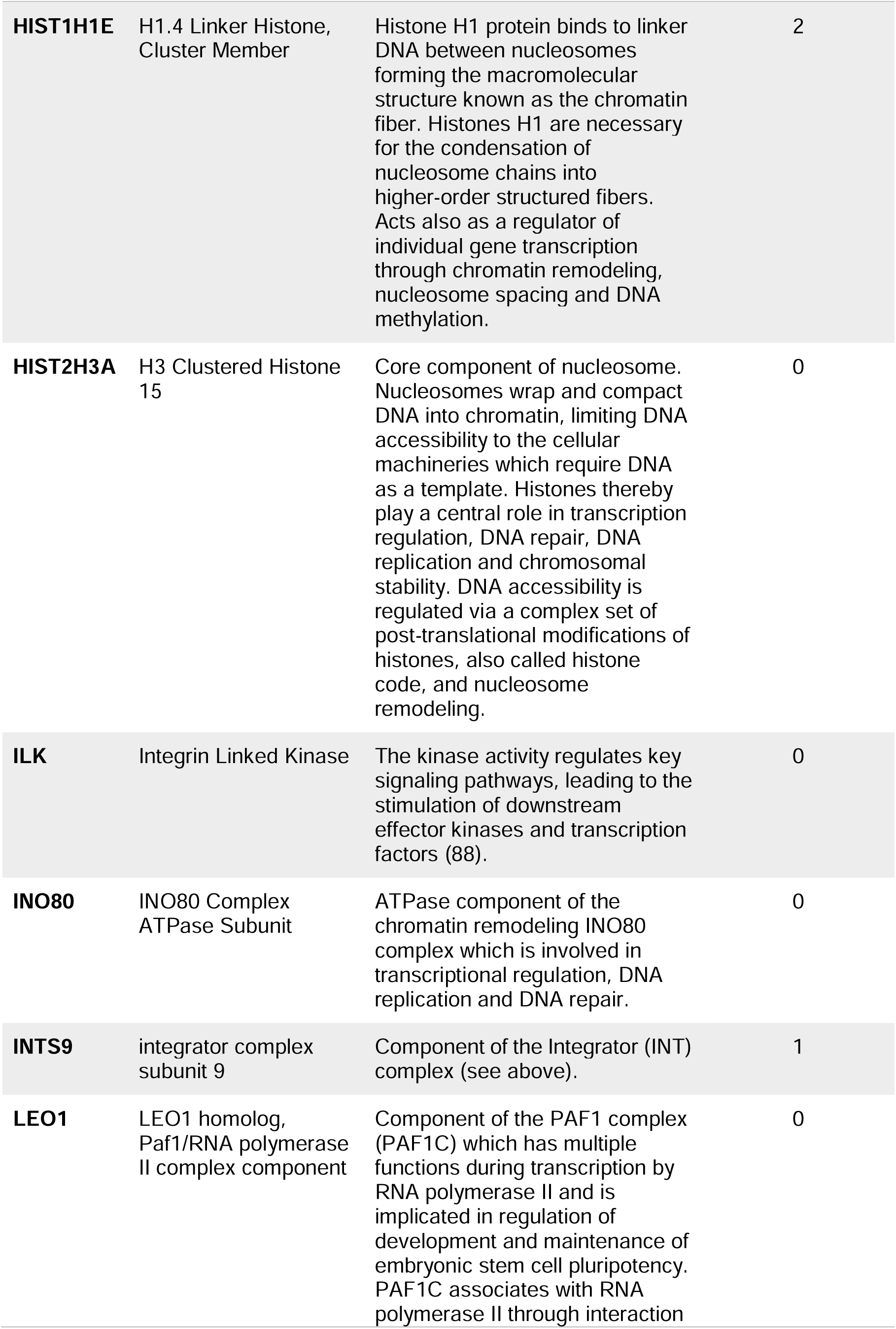
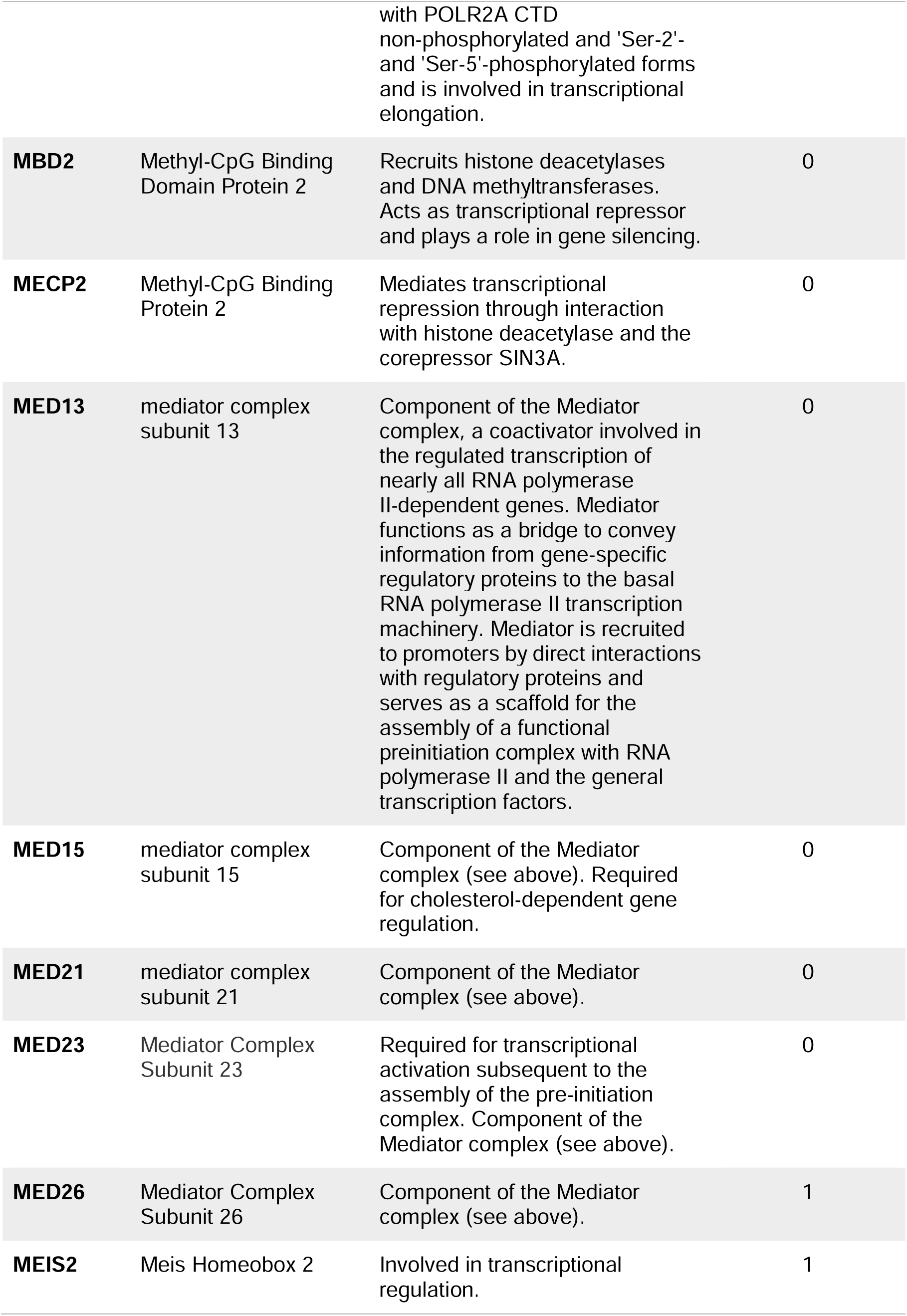
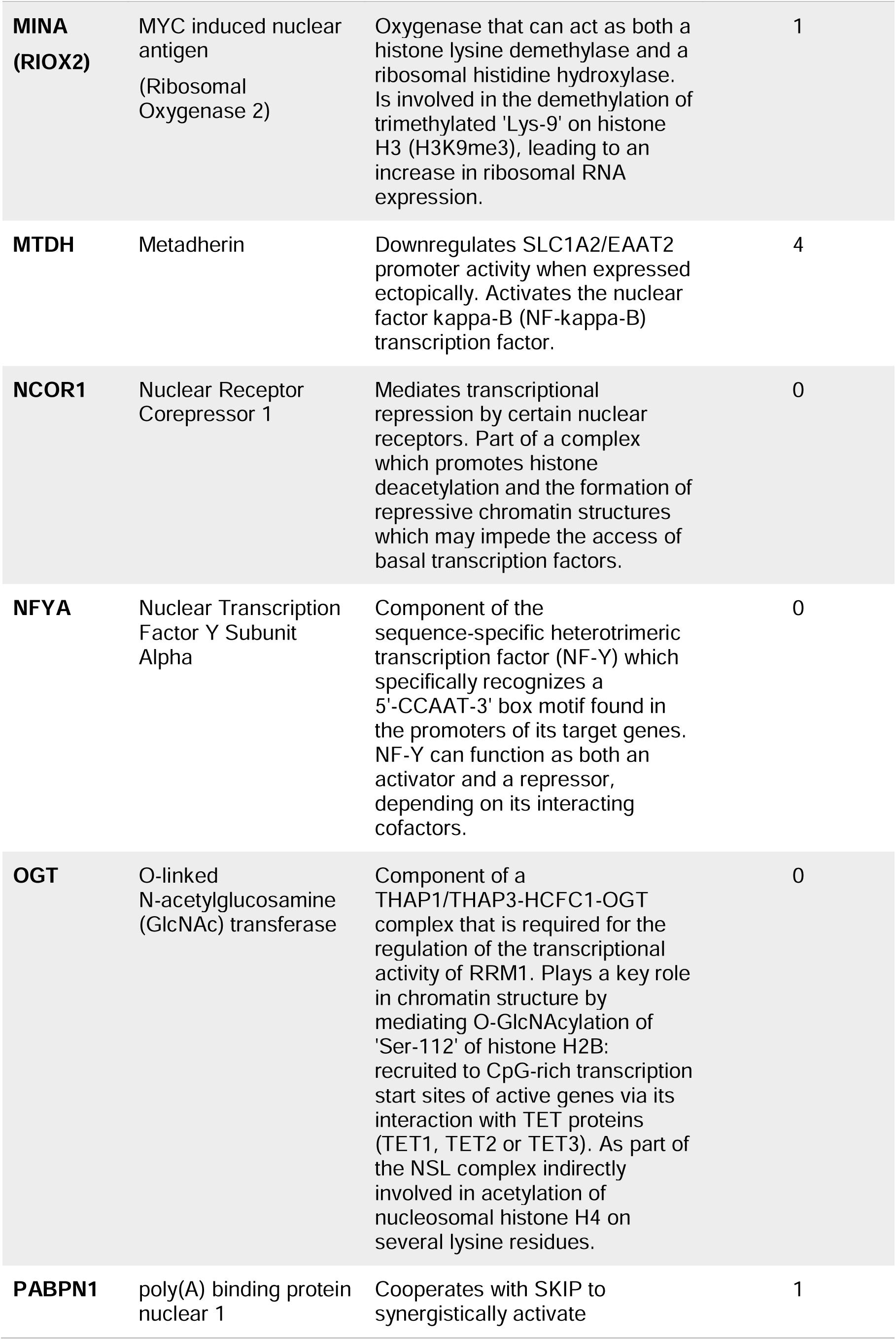
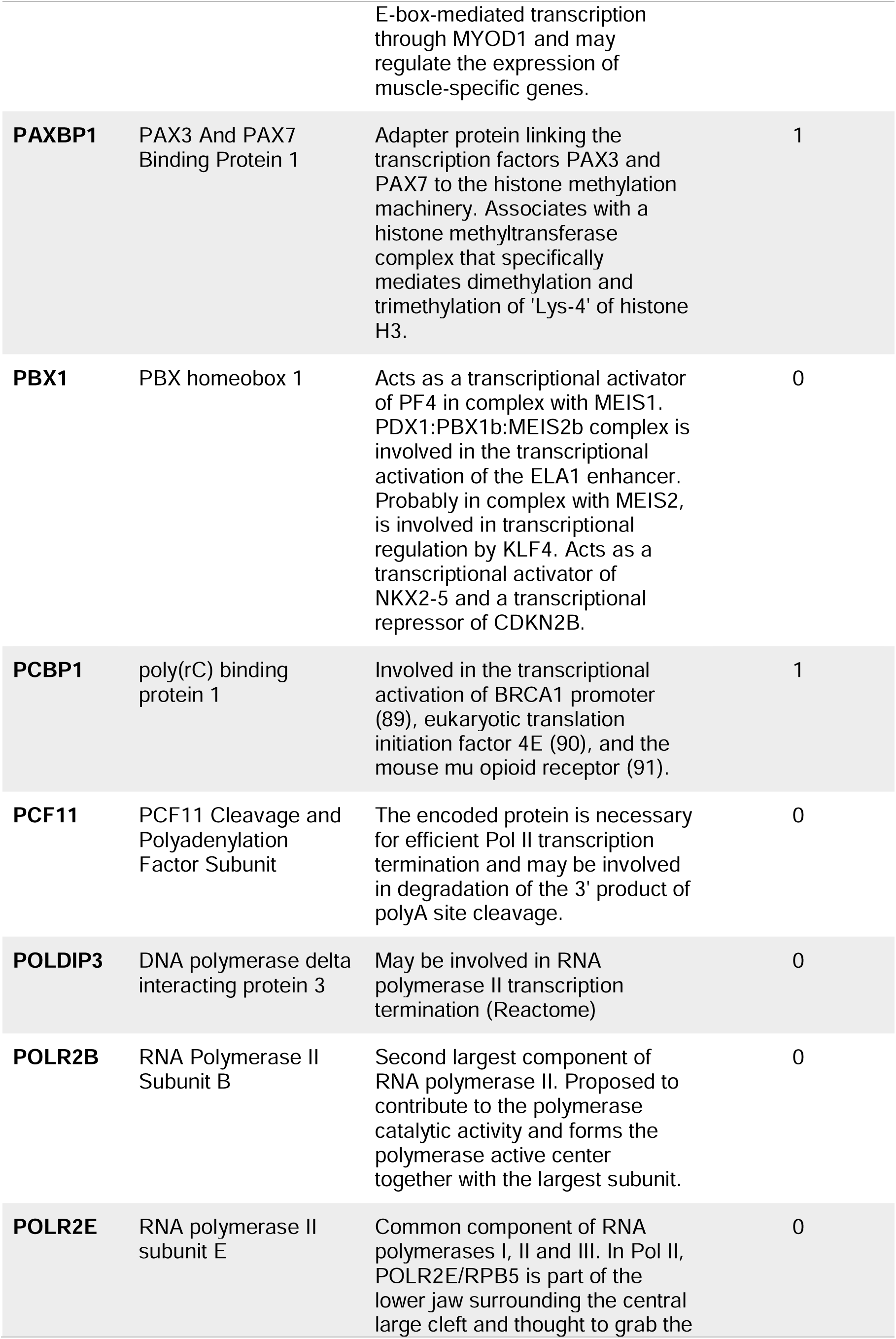
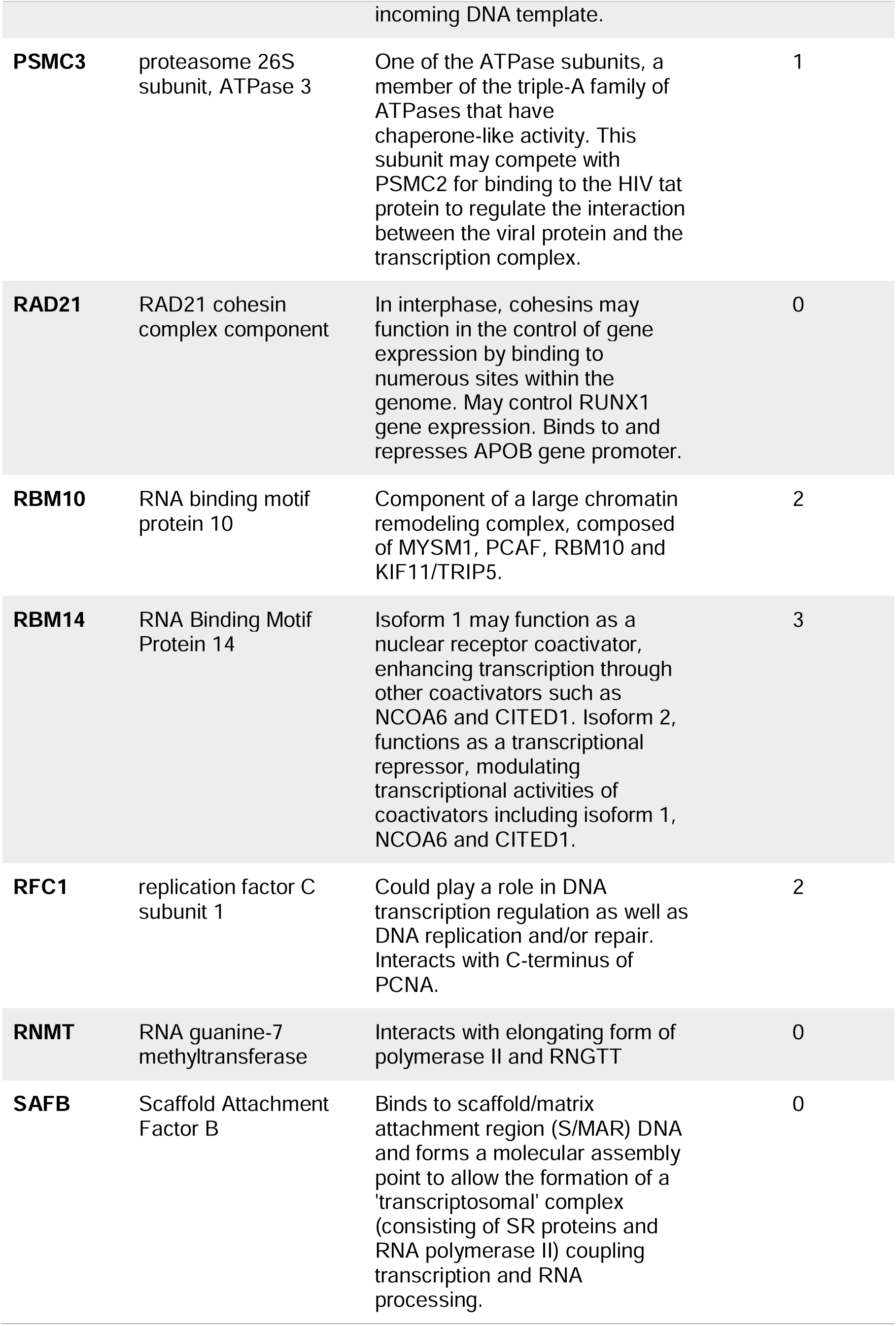
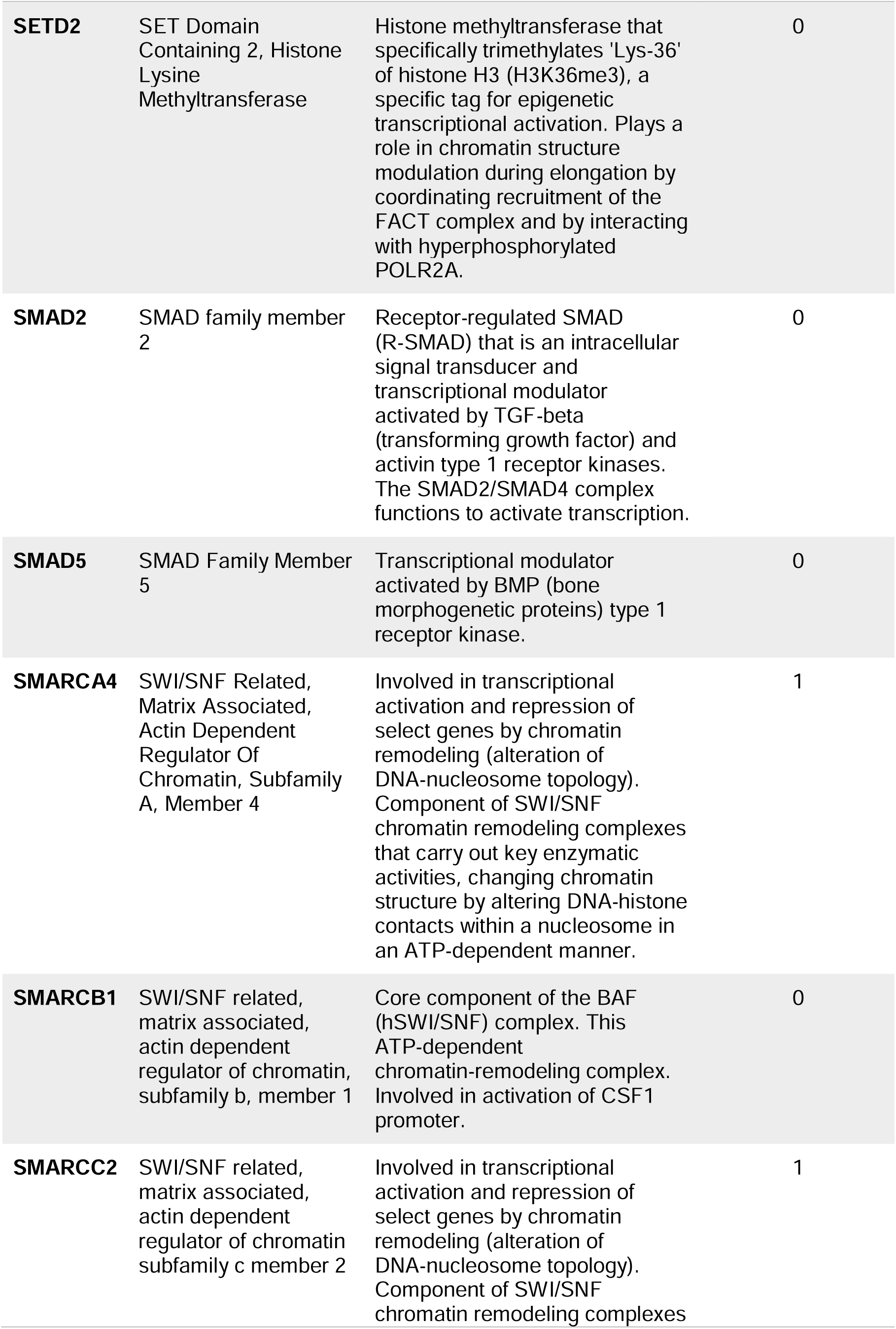
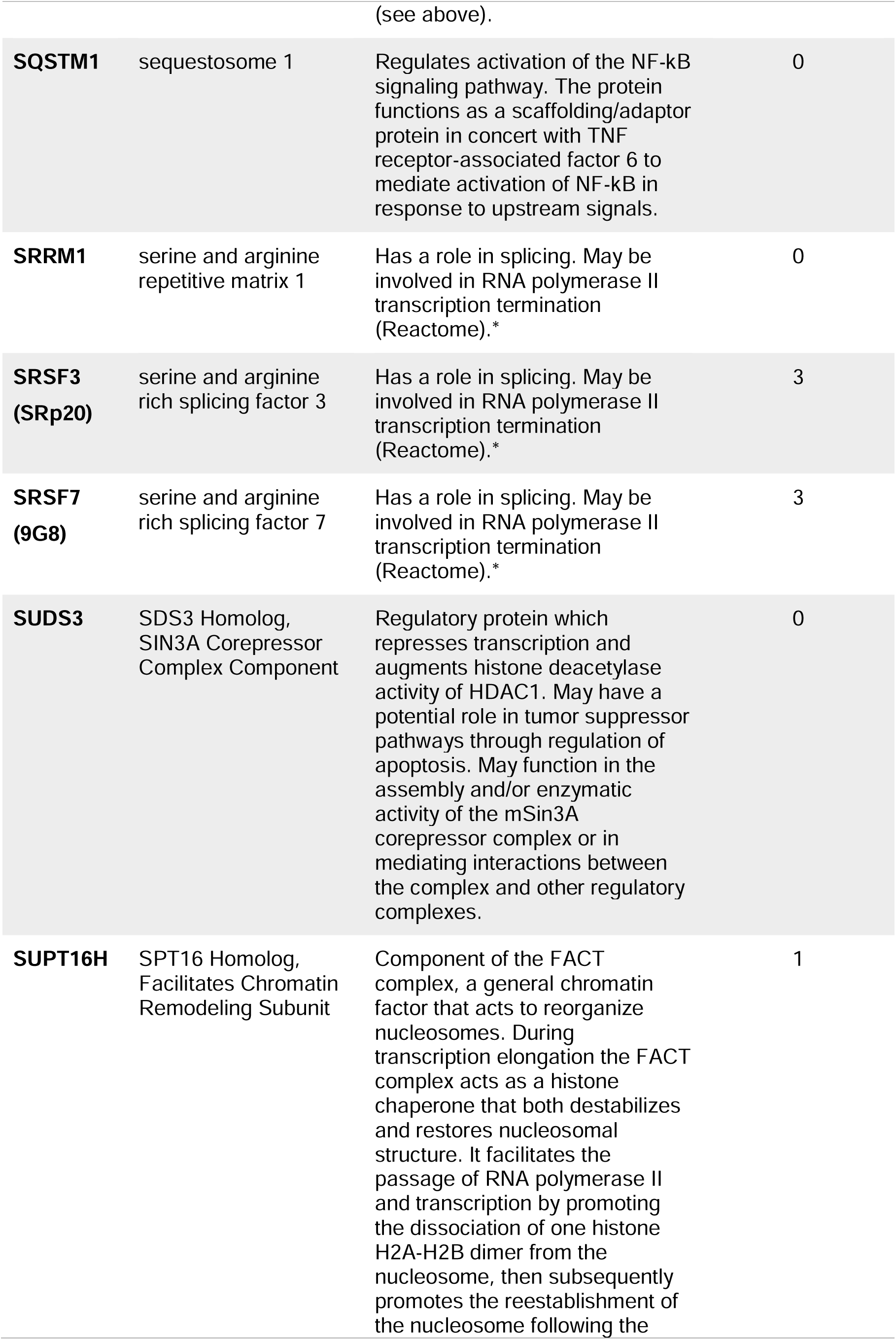
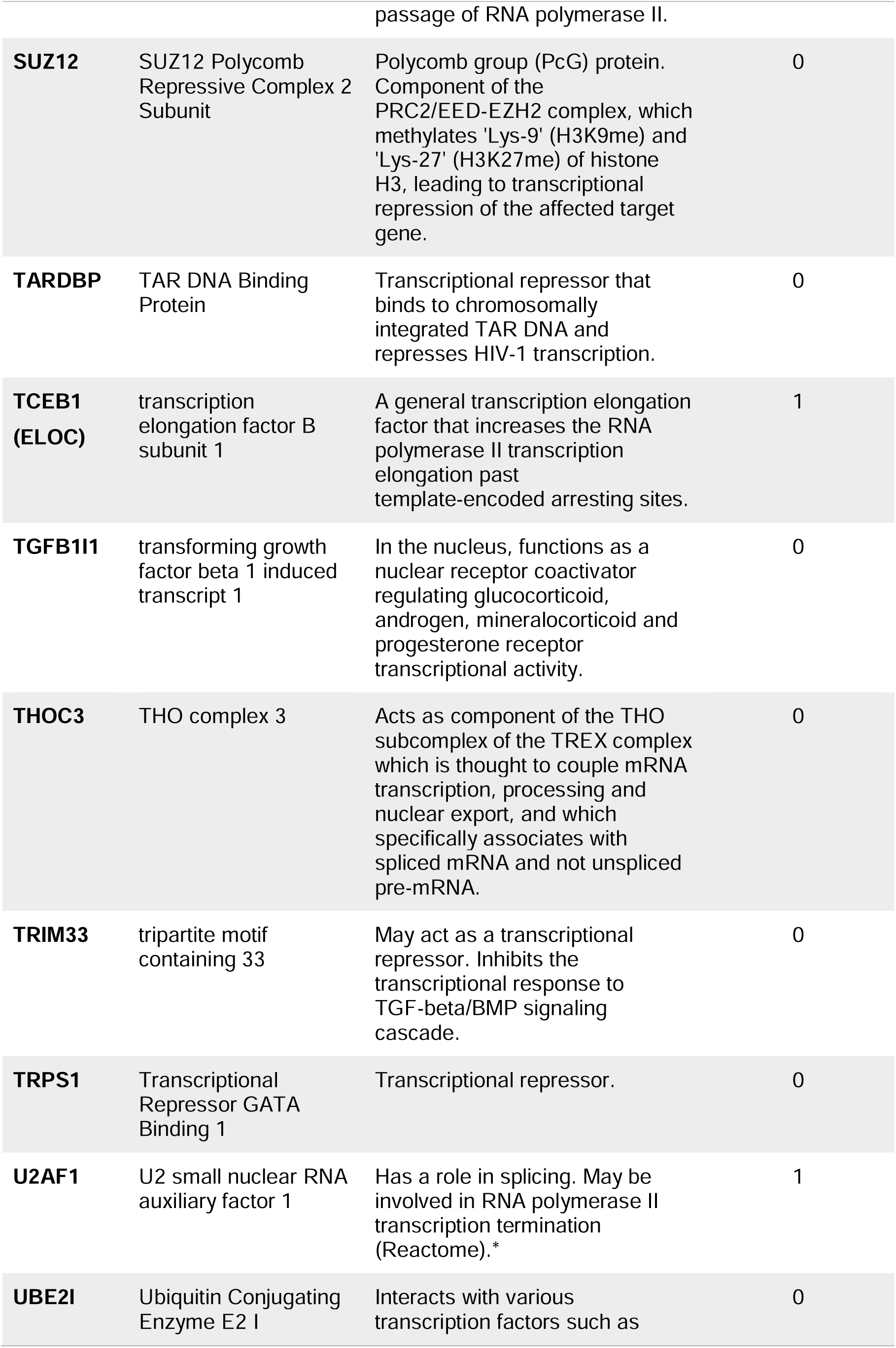
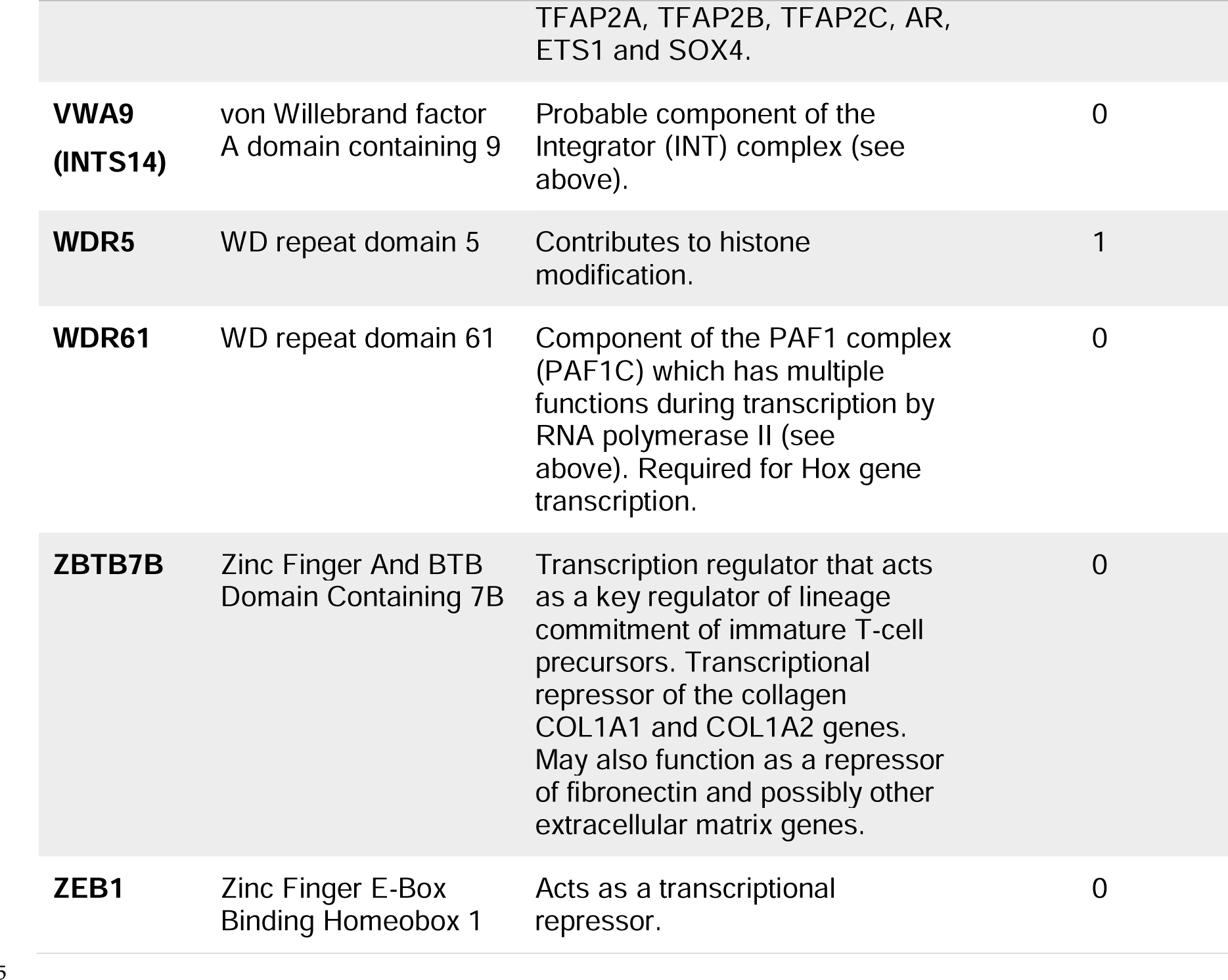
Names and functions of the proteins identified in the HIV-1 proteomics list under GO:0006366∼ transcription from RNA polymerase II promoter.

| Symbol | Description | Function related to Transcription (mostly summarized from Genecards) | Frequency (# of publications) |
| --- | --- | --- | --- |
| **ALYREF** | Aly/REF Export Factor | Component of the TREX complex which is thought to couple mRNA transcription, processing and nuclear export, and specifically associates with spliced mRNA and not with unspliced pre-mRNA. TREX is recruited to spliced mRNAs by a transcription-independent mechanism, binds to mRNA upstream of the exon-junction complex (EJC) and is recruited in a splicing- and cap-dependent manner to a region near the 5' end of the mRNA where it functions in mRNA export to the cytoplasm. Involved in transcription elongation and genome stability. Acts as chaperone and promotes the dimerization of transcription factors containing basic leucine zipper (bZIP) domains and thereby promotes transcriptional activation. | 2 |
| **ARID4A** | AT-rich interactive domain-containing protein 4A | DNA-binding protein which modulates activity of several transcription factors. May function as part of an mSin3A repressor complex. | 0 |
| **ASH2L** | ASH2 Like, Histone Lysine Methyltransferase Complex Subunit | Component or associated component of some histone methyltransferase complexes which regulates transcription through recruitment of those complexes to gene promoters. | 0 |
| **ASUN (INTS13)** | asunder, spermatogenesis regulator | A probable component of the Integrator (INT) complex – involved in the small nuclear RNAs (snRNA) U1 and U2 transcription and 3' processing. The INT complex is associated with the C-terminal domain (CTD) | 1 |
|  |  | of RNA polymerase II largest subunit (POLR2A) and is recruited to the U1 and U2 snRNAs genes. |  |
| **BRMS1** | breast cancer metastasis suppressor 1 | Transcriptional repressor. Down-regulates NF-κB transcriptional activation. Promotes HDAC1 binding to promoter regions. | 0 |
| **C1QBP** | Complement component 1 Q subcomponent-binding protein, mitochondrial | May be involved in regulation of FOXC1 transcriptional activity and NFY/CCAAT-binding factor complex-mediated transcription. | 1 |
| **CDK12** | cyclin dependent kinase 12 | Cyclin-dependent kinase that phosphorylates the C-terminal domain (CTD) of the large subunit of RNA polymerase II (POLR2A), thereby acting as a key regulator of transcription elongation. Regulates the expression of genes involved in DNA repair and is required for the maintenance of genomic stability. | 1 |
| **CDKN2A** | cyclin dependent kinase inhibitor 2A<br>(<u>Tumor suppressor ARF</u>) | Has an isoform (ARF) that binds to BCL6 and down-regulates BCL6-induced transcriptional repression. Binds to E2F1 and MYC and blocks their transcriptional activator activity but has no effect on MYC transcriptional repression. Binds to TOP1/TOPOI and stimulates its activity. This complex binds to rRNA gene promoters and may play a role in rRNA transcription and/or maturation. | 0 |
| **CGGBP1** | CGG Triplet Repeat Binding Protein 1 | Binds to nonmethylated 5'-d(CGG)(n)-3' trinucleotide repeats in the FMR1 promoter. May play a role in regulating FMR1 promoter. | 0 |
| **CHTOP** | Chromatin Target of PRMT1 | Required for effective mRNA nuclear export and is a component of the TREX complex (see above). | 2 |
| **CIC** | capicua transcriptional repressor (protein | Transcriptional repressor which plays a role in development of the | 0 |
|  | capicua homolog) | central nervous system (CNS). |  |
| **CLP1** | Cleavage Factor Polyribonucleotide Kinase Subunit 1 | Member of the tRNA splicing complex. May be involved in the termination of RNA polymerase II transcription (Reactome).* | 0 |
| **CPSF3** | cleavage and polyadenylation specific factor 3 | Member of the cleavage and polyadenylation specificity factor (CPSF) complex. May be involved in the termination of RNA polymerase II transcription (Reactome).* | 0 |
| **CPSF3L (INTS11)** | cleavage and polyadenylation specific factor 3-like (Integrator Complex Subunit 11) | Catalytic component of the Integrator (INT) complex (see above). | 0 |
| **CPSF7** | Cleavage And Polyadenylation Specific Factor 7 | Member of the CPSF complex. May be involved in the termination of RNA polymerase II transcription (Reactome).* | 0 |
| **CREBBP** | CREB binding protein | Acetylates histones, giving a specific tag for transcriptional activation. | 0 |
| **CTNNB1** | Catenin beta 1 | In the presence of Wnt ligand, CTNNB1 accumulates in the nucleus, where it acts as a coactivator for transcription factors of the TCF/LEF family, leading to activate Wnt responsive genes. | 1 |
| **DDX20** | DEAD-Box Helicase 20 | Represses Egr2-mediated transcriptional activation (87). | 1 |
| **DNAJA3** | DnaJ heat shock protein family (Hsp40) member A3 | Can modulate IFN-gamma-mediated transcriptional activity. | 1 |
| **DNMT1** | DNA methyltransferase | Mediates transcriptional repression by direct binding to HDAC2. In association with DNMT3B and via the recruitment of CTCFL/BORIS, involved in activation of BAG1 gene expression by modulating dimethylation of promoter histone H3 at H3K4 and H3K9. | 1 |
| **EHMT2** | Euchromatic Histone Lysine Methyltransferase 2 | Histone methyltransferase that specifically mono- and dimethylates 'Lys-9' of histone H3 (H3K9me1 and H3K9me2, respectively) in euchromatin. H3K9me represents a specific tag for epigenetic transcriptional repression by recruiting HP1 proteins to methylated histones. | 0 |
| **FIP1L1** | Factor Interacting With PAPOLA And CPSF1 | Member of the CPSF complex. May be involved in RNA polymerase II transcription termination (Reactome)* | 1 |
| **FOXP1** | forkhead box P1 | Transcriptional repressor. | 0 |
| **GATAD2A** | GATA zinc finger domain containing 2A | Transcriptional repressor. Enhances MBD2-mediated repression. Efficient repression requires the presence of GATAD2B. | 0 |
| **GTF2H1** | general transcription factor IIH subunit 1 | Component of the general transcription and DNA repair factor IIH (TFIIH) core complex, which is involved in general and transcription-coupled nucleotide excision repair (NER) of damaged DNA and, when complexed to CAK, in RNA transcription by RNA polymerase II. In transcription, TFIIH has an essential role in transcription initiation. | 0 |
| **GTF2H3** | general transcription factor IIH subunit 3 | See above. | 0 |
| **GTF2H4** | general transcription factor IIH subunit 4 | See above. | 0 |
| **HDAC3** | histone deacetylase 3 | Responsible for the deacetylation of lysine residues on the N-terminal part of the core histones (H2A, H2B, H3 and H4), and some other non-histone substrates. Histone deacetylation gives a tag for epigenetic repression and plays an important role in transcriptional regulation, cell cycle progression and developmental events. | 0 |
| **HIST1H1E** | H1.4 Linker Histone, Cluster Member | Histone H1 protein binds to linker DNA between nucleosomes forming the macromolecular structure known as the chromatin fiber. Histones H1 are necessary for the condensation of nucleosome chains into higher-order structured fibers. Acts also as a regulator of individual gene transcription through chromatin remodeling, nucleosome spacing and DNA methylation. | 2 |
| **HIST2H3A** | H3 Clustered Histone 15 | Core component of nucleosome. Nucleosomes wrap and compact DNA into chromatin, limiting DNA accessibility to the cellular machineries which require DNA as a template. Histones thereby play a central role in transcription regulation, DNA repair, DNA replication and chromosomal stability. DNA accessibility is regulated via a complex set of post-translational modifications of histones, also called histone code, and nucleosome remodeling. | 0 |
| **ILK** | Integrin Linked Kinase | The kinase activity regulates key signaling pathways, leading to the stimulation of downstream effector kinases and transcription factors (88). | 0 |
| **INO80** | INO80 Complex ATPase Subunit | ATPase component of the chromatin remodeling INO80 complex which is involved in transcriptional regulation, DNA replication and DNA repair. | 0 |
| **INTS9** | integrator complex subunit 9 | Component of the Integrator (INT) complex (see above). | 1 |
| **LEO1** | LEO1 homolog, Paf1/RNA polymerase II complex component | Component of the PAF1 complex (PAF1C) which has multiple functions during transcription by RNA polymerase II and is implicated in regulation of development and maintenance of embryonic stem cell pluripotency. PAF1C associates with RNA polymerase II through interaction with POLR2A CTD non-phosphorylated and 'Ser-2'- and 'Ser-5'-phosphorylated forms and is involved in transcriptional elongation. | 0 |
| **MBD2** | Methyl-CpG Binding Domain Protein 2 | Recruits histone deacetylases and DNA methyltransferases. Acts as transcriptional repressor and plays a role in gene silencing. | 0 |
| **MECP2** | Methyl-CpG Binding Protein 2 | Mediates transcriptional repression through interaction with histone deacetylase and the corepressor SIN3A. | 0 |
| **MED13** | mediator complex subunit 13 | Component of the Mediator complex, a coactivator involved in the regulated transcription of nearly all RNA polymerase II-dependent genes. Mediator functions as a bridge to convey information from gene-specific regulatory proteins to the basal RNA polymerase II transcription machinery. Mediator is recruited to promoters by direct interactions with regulatory proteins and serves as a scaffold for the assembly of a functional preinitiation complex with RNA polymerase II and the general transcription factors. | 0 |
| **MED15** | mediator complex subunit 15 | Component of the Mediator complex (see above). Required for cholesterol-dependent gene regulation. | 0 |
| **MED21** | mediator complex subunit 21 | Component of the Mediator complex (see above). | 0 |
| **MED23** | Mediator Complex Subunit 23 | Required for transcriptional activation subsequent to the assembly of the pre-initiation complex. Component of the Mediator complex (see above). | 0 |
| **MED26** | Mediator Complex Subunit 26 | Component of the Mediator complex (see above). | 1 |
| **MEIS2** | Meis Homeobox 2 | Involved in transcriptional regulation. | 1 |
| **MINA**<br>**(RIOX2)** | MYC induced nuclear antigen<br>(Ribosomal Oxygenase 2) | Oxygenase that can act as both a histone lysine demethylase and a ribosomal histidine hydroxylase. Is involved in the demethylation of trimethylated 'Lys-9' on histone H3 (H3K9me3), leading to an increase in ribosomal RNA expression. | 1 |
| **MTDH** | Metadherin | Downregulates SLC1A2/EAAT2 promoter activity when expressed ectopically. Activates the nuclear factor kappa-B (NF-kappa-B) transcription factor. | 4 |
| **NCOR1** | Nuclear Receptor Corepressor 1 | Mediates transcriptional repression by certain nuclear receptors. Part of a complex which promotes histone deacetylation and the formation of repressive chromatin structures which may impede the access of basal transcription factors. | 0 |
| **NFYA** | Nuclear Transcription Factor Y Subunit Alpha | Component of the sequence-specific heterotrimeric transcription factor (NF-Y) which specifically recognizes a 5'-CCAAT-3' box motif found in the promoters of its target genes. NF-Y can function as both an activator and a repressor, depending on its interacting cofactors. | 0 |
| **OGT** | O-linked N-acetylglucosamine (GlcNAc) transferase | Component of a THAP1/THAP3-HCFC1-OGT complex that is required for the regulation of the transcriptional activity of RRM1. Plays a key role in chromatin structure by mediating O-GlcNAcylation of 'Ser-112' of histone H2B: recruited to CpG-rich transcription start sites of active genes via its interaction with TET proteins (TET1, TET2 or TET3). As part of the NSL complex indirectly involved in acetylation of nucleosomal histone H4 on several lysine residues. | 0 |
| **PABPN1** | poly(A) binding protein nuclear 1 | Cooperates with SKIP to synergistically activate | 1 |
|  |  | E-box-mediated transcription through MYOD1 and may regulate the expression of muscle-specific genes. |  |
| **PAXBP1** | PAX3 And PAX7 Binding Protein 1 | Adapter protein linking the transcription factors PAX3 and PAX7 to the histone methylation machinery. Associates with a histone methyltransferase complex that specifically mediates dimethylation and trimethylation of 'Lys-4' of histone H3. | 1 |
| **PBX1** | PBX homeobox 1 | Acts as a transcriptional activator of PF4 in complex with MEIS1. PDX1:PBX1b:MEIS2b complex is involved in the transcriptional activation of the ELA1 enhancer. Probably in complex with MEIS2, is involved in transcriptional regulation by KLF4. Acts as a transcriptional activator of NKX2-5 and a transcriptional repressor of CDKN2B. | 0 |
| **PCBP1** | poly(rC) binding protein 1 | Involved in the transcriptional activation of BRCA1 promoter (89), eukaryotic translation initiation factor 4E (90), and the mouse mu opioid receptor (91). | 1 |
| **PCF11** | PCF11 Cleavage and Polyadenylation Factor Subunit | The encoded protein is necessary for efficient Pol II transcription termination and may be involved in degradation of the 3' product of polyA site cleavage. | 0 |
| **POLDIP3** | DNA polymerase delta interacting protein 3 | May be involved in RNA polymerase II transcription termination (Reactome) | 0 |
| **POLR2B** | RNA Polymerase II Subunit B | Second largest component of RNA polymerase II. Proposed to contribute to the polymerase catalytic activity and forms the polymerase active center together with the largest subunit. | 0 |
| **POLR2E** | RNA polymerase II subunit E | Common component of RNA polymerases I, II and III. In Pol II, POLR2E/RPB5 is part of the lower jaw surrounding the central large cleft and thought to grab the incoming DNA template. | 0 |
| **PSMC3** | proteasome 26S subunit, ATPase 3 | One of the ATPase subunits, a member of the triple-A family of ATPases that have chaperone-like activity. This subunit may compete with PSMC2 for binding to the HIV tat protein to regulate the interaction between the viral protein and the transcription complex. | 1 |
| **RAD21** | RAD21 cohesin complex component | In interphase, cohesins may function in the control of gene expression by binding to numerous sites within the genome. May control RUNX1 gene expression. Binds to and represses APOB gene promoter. | 0 |
| **RBM10** | RNA binding motif protein 10 | Component of a large chromatin remodeling complex, composed of MYSM1, PCAF, RBM10 and KIF11/TRIP5. | 2 |
| **RBM14** | RNA Binding Motif Protein 14 | Isoform 1 may function as a nuclear receptor coactivator, enhancing transcription through other coactivators such as NCOA6 and CITED1. Isoform 2, functions as a transcriptional repressor, modulating transcriptional activities of coactivators including isoform 1, NCOA6 and CITED1. | 3 |
| **RFC1** | replication factor C subunit 1 | Could play a role in DNA transcription regulation as well as DNA replication and/or repair. Interacts with C-terminus of PCNA. | 2 |
| **RNMT** | RNA guanine-7 methyltransferase | Interacts with elongating form of polymerase II and RNGTT | 0 |
| **SAFB** | Scaffold Attachment Factor B | Binds to scaffold/matrix attachment region (S/MAR) DNA and forms a molecular assembly point to allow the formation of a 'transcriptosomal' complex (consisting of SR proteins and RNA polymerase II) coupling transcription and RNA processing. | 0 |
| **SETD2** | SET Domain<br>Containing 2, Histone<br>Lysine<br>Methyltransferase | Histone methyltransferase that specifically trimethylates 'Lys-36' of histone H3 (H3K36me3), a specific tag for epigenetic transcriptional activation. Plays a role in chromatin structure modulation during elongation by coordinating recruitment of the FACT complex and by interacting with hyperphosphorylated POLR2A. | 0 |
| **SMAD2** | SMAD family member<br>2 | Receptor-regulated SMAD (R-SMAD) that is an intracellular signal transducer and transcriptional modulator activated by TGF-beta (transforming growth factor) and activin type 1 receptor kinases. The SMAD2/SMAD4 complex functions to activate transcription. | 0 |
| **SMAD5** | SMAD Family Member<br>5 | Transcriptional modulator activated by BMP (bone morphogenetic proteins) type 1 receptor kinase. | 0 |
| **SMARCA4** | SWI/SNF Related,<br>Matrix Associated,<br>Actin Dependent<br>Regulator Of<br>Chromatin, Subfamily<br>A, Member 4 | Involved in transcriptional activation and repression of select genes by chromatin remodeling (alteration of DNA-nucleosome topology). Component of SWI/SNF chromatin remodeling complexes that carry out key enzymatic activities, changing chromatin structure by altering DNA-histone contacts within a nucleosome in an ATP-dependent manner. | 1 |
| **SMARCB1** | SWI/SNF related,<br>matrix associated,<br>actin dependent<br>regulator of chromatin,<br>subfamily b, member 1 | Core component of the BAF (hSWI/SNF) complex. This ATP-dependent chromatin-remodeling complex. Involved in activation of CSF1 promoter. | 0 |
| **SMARCC2** | SWI/SNF related,<br>matrix associated,<br>actin dependent<br>regulator of chromatin<br>subfamily c member 2 | Involved in transcriptional activation and repression of select genes by chromatin remodeling (alteration of DNA-nucleosome topology). Component of SWI/SNF chromatin remodeling complexes | 1 |
| **SQSTM1** | sequestosome 1 | Regulates activation of the NF-κB signaling pathway. The protein functions as a scaffolding/adaptor protein in concert with TNF receptor-associated factor 6 to mediate activation of NF-κB in response to upstream signals. | 0 |
| **SRRM1** | serine and arginine repetitive matrix 1 | Has a role in splicing. May be involved in RNA polymerase II transcription termination (Reactome).* | 0 |
| **SRSF3 (SRp20)** | serine and arginine rich splicing factor 3 | Has a role in splicing. May be involved in RNA polymerase II transcription termination (Reactome).* | 3 |
| **SRSF7 (9G8)** | serine and arginine rich splicing factor 7 | Has a role in splicing. May be involved in RNA polymerase II transcription termination (Reactome).* | 3 |
| **SUDS3** | SDS3 Homolog, SIN3A Corepressor Complex Component | Regulatory protein which represses transcription and augments histone deacetylase activity of HDAC1. May have a potential role in tumor suppressor pathways through regulation of apoptosis. May function in the assembly and/or enzymatic activity of the mSin3A corepressor complex or in mediating interactions between the complex and other regulatory complexes. | 0 |
| **SUPT16H** | SPT16 Homolog, Facilitates Chromatin Remodeling Subunit | Component of the FACT complex, a general chromatin factor that acts to reorganize nucleosomes. During transcription elongation the FACT complex acts as a histone chaperone that both destabilizes and restores nucleosomal structure. It facilitates the passage of RNA polymerase II and transcription by promoting the dissociation of one histone H2A-H2B dimer from the nucleosome, then subsequently promotes the reestablishment of the nucleosome following the passage of RNA polymerase II. | 1 |
| **SUZ12** | SUZ12 Polycomb Repressive Complex 2 Subunit | Polycomb group (PcG) protein. Component of the PRC2/EED-EZH2 complex, which methylates 'Lys-9' (H3K9me) and 'Lys-27' (H3K27me) of histone H3, leading to transcriptional repression of the affected target gene. | 0 |
| **TARDBP** | TAR DNA Binding Protein | Transcriptional repressor that binds to chromosomally integrated TAR DNA and represses HIV-1 transcription. | 0 |
| **TCEB1 (ELOC)** | transcription elongation factor B subunit 1 | A general transcription elongation factor that increases the RNA polymerase II transcription elongation past template-encoded arresting sites. | 1 |
| **TGFB1I1** | transforming growth factor beta 1 induced transcript 1 | In the nucleus, functions as a nuclear receptor coactivator regulating glucocorticoid, androgen, mineralocorticoid and progesterone receptor transcriptional activity. | 0 |
| **THOC3** | THO complex 3 | Acts as component of the THO subcomplex of the TREX complex which is thought to couple mRNA transcription, processing and nuclear export, and which specifically associates with spliced mRNA and not unspliced pre-mRNA. | 0 |
| **TRIM33** | tripartite motif containing 33 | May act as a transcriptional repressor. Inhibits the transcriptional response to TGF-beta/BMP signaling cascade. | 0 |
| **TRPS1** | Transcriptional Repressor GATA Binding 1 | Transcriptional repressor. | 0 |
| **U2AF1** | U2 small nuclear RNA auxiliary factor 1 | Has a role in splicing. May be involved in RNA polymerase II transcription termination (Reactome).* | 1 |
| **UBE2I** | Ubiquitin Conjugating Enzyme E2 I | Interacts with various transcription factors such as | 0 |
|  |  | TFAP2A, TFAP2B, TFAP2C, AR, ETS1 and SOX4. |  |
| **VWA9 (INTS14)** | von Willebrand factor A domain containing 9 | Probable component of the Integrator (INT) complex (see above). | 0 |
| **WDR5** | WD repeat domain 5 | Contributes to histone modification. | 1 |
| **WDR61** | WD repeat domain 61 | Component of the PAF1 complex (PAF1C) which has multiple functions during transcription by RNA polymerase II (see above). Required for Hox gene transcription. | 0 |
| **ZBTB7B** | Zinc Finger And BTB Domain Containing 7B | Transcription regulator that acts as a key regulator of lineage commitment of immature T-cell precursors. Transcriptional repressor of the collagen COL1A1 and COL1A2 genes. May also function as a repressor of fibronectin and possibly other extracellular matrix genes. | 0 |
| **ZEB1** | Zinc Finger E-Box Binding Homeobox 1 | Acts as a transcriptional repressor. | 0 |

We previously showed that nuclear localization of Gag contributes to efficient USvRNA selection for packaging (7). One possible model is that interaction of Gag with Mediator proteins could tether Gag at active transcription sites, increasing the chance that it would find and associate with nascent USvRNA. To examine the interaction of RSV Gag and Med26, we utilized confocal microscopy to assess colocalization between transfected RSV Gag and Med26 in QT6 cells and observed that these two proteins did in fact colocalize (**Fig 4A-B**, **Video S1**) (M1, Med26∩RSV Gag: 0.0364 ± 0.004; M2, RSV Gag∩Med26: 0.471 ± 0.067).

**Figure 4.**
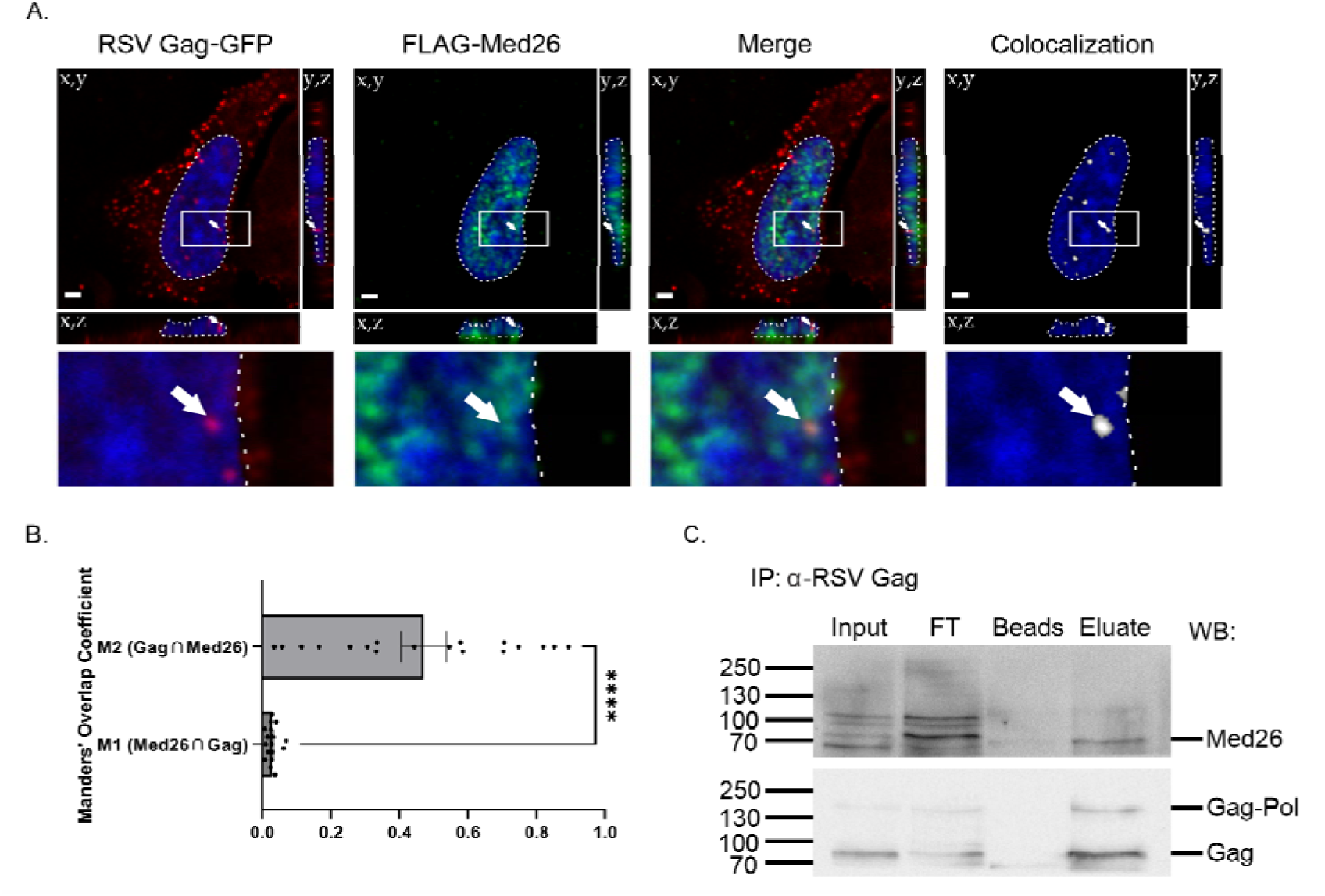
RSV Gag colocalized and co-immunoprecipitated with Med26. (**A**) Transfected RSV Gag-GFP (red) and FLAG-Med26 (green) colocalize (white) within QT6 cells. Image representative of average colocalization, as quantified in panel (**B**). Nuclei (blue) are outlined by a dotted white line, and regions boxed in the main images are enlarged below. White arrows are included to guide the eye. Scale bars = 2 µm. (**B**) Manders’ Overlap Coefficient values for image set represented by (**A**). Individual values are shown in addition mean ± SEM, n ≥ 17; ****, p<0.0001 by unpaired two-tailed t-test. (**C**) 500 µg of RC.V8-infected QT6 nuclear lysates were incubated with an α-RSV Gag antibody (mouse α-RSV CA.A11, gift from Neil Christensen, Penn State College of Medicine), followed by antibody capture on Pierce™ Protein G Magnetic Beads. After extensive washing, proteins were eluted from beads by boiling in 1X SDS-PAGE sample buffer and run on a 10% SDS-PAGE gel, transferred to PVDF, and Western blotted first for Med26 (top) followed by RSV Gag (bottom). The position of molecular weight markers, in kilodaltons, are indicated on the left. FT, flow through; Beads, lysate only; Eluate, lysate plus antibody. Images representative of three independent experiments.

Next, immunoprecipitation of endogenous RSV Gag from RC.V8-infected QT6 cell nuclear lysates was undertaken, and samples were resolved via SDS-PAGE and subjected to western blotting, as described in the Methods section (**Fig 4C**). In line with our proteomic results, we observed that Med26 and RSV Gag co-immunoprecipitated (**Fig 4C**, **Eluate lane**). The molecular mechanisms underlying this interaction and the possible involvement of additional Mediator proteins will be pursued in future studies.

## 4. Discussion

Several different laboratories have observed that the retroviral Gag proteins of HIV-1, RSV, MMTV, MLV, FIV, PFV, and MPMV undergo nuclear localization (9–23, 25, 26, 100–103). As described in detail in this report, six previously published proteomic studies searching for binding partners of HIV-1 Gag identified many nuclear proteins (30–35). As we were specifically interested in the nuclear interactomes of RSV and HIV-1 Gag, we took a different approach than previous groups, using nuclear lysates incubated with recombinant Gag proteins to perform affinity purification of complexes followed by mass spectrometry for our proteomic analysis. Despite these differences in methodology, a set of overlapping factors were identified for HIV-1 Gag that included a large number of proteins involved in nuclear processes such as transcription/gene expression, RNA processing, splicing, and chromatin remodeling. Comparison of the potential binding partners of RSV and HIV-1 Gag indicated that 57 proteins were found to be in common (**Fig 2**). Whether these factors have similar functions in RSV or HIV-1 replication remains to be examined. When the HIV-1 Gag interactomes identified by other laboratories were compared to the interacting proteins identified by us in this report, 190 common proteins were found by at least two independent laboratory groups. Further experimentation will be needed to validate each of these factors to determine whether they play important roles in retrovirus replication or pathogenesis **(Fig 5)**.

**Figure 5.**
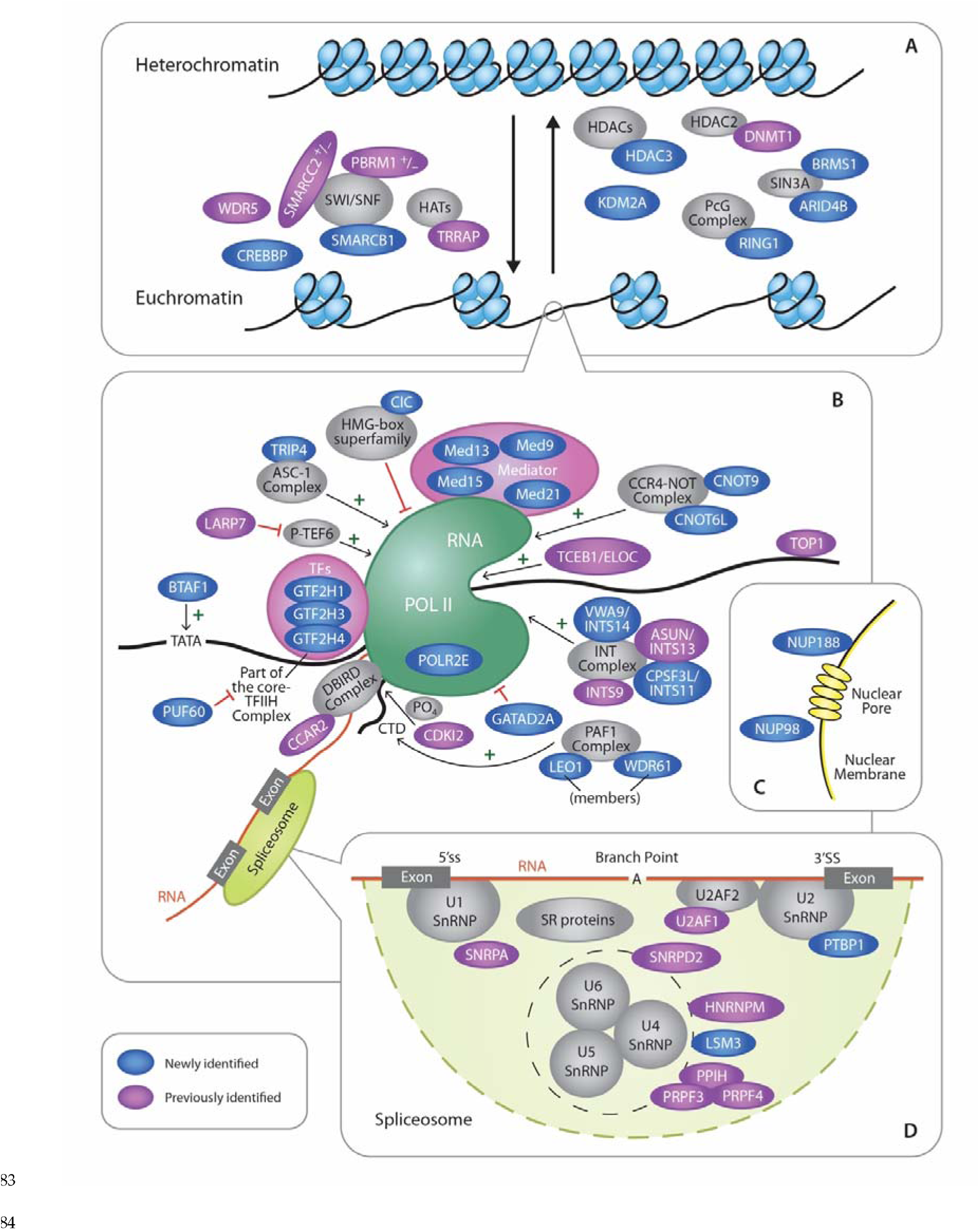
HIV-1 interactome pathway analysis. This diagram illustrates the HIV-1 Gag-interacting nuclear host factors discussed in this study. Factors shaded blue were uniquely identified in the present study (newly identified). Proteins shaded in purple were identified in this publication as well as at least one other published report (previously identified). Gray shading was used to show host protein complexes that are involved with Gag-interacting factors in chromatin remodeling, gene expression, nuclear export, and splic-ing. (**A**) HIV-1 Gag interacting factors that promote an open chromatin structure (euchromatin state) are on the left, whereas proteins involved in condensing chromatin are shown on the right. (**B**) Proteins involved in regulation of gene expression are depicted. Factors that promote transcription initiation are indicated by arrows and green plus signs. Factors that suppress or inhibit transcription are demarcated by red blocking lines. (**C**) The two nucleoporin proteins NUP98 and NUP188 were identified and are involved in trafficking between the nucleus and the cytoplasm through the nuclear pore complex. (**D**) Proteins that localize to the spliceosome and are involved in RNA splicing are shown.

Our past and present cell fractionation experiments demonstrated that, in addition to being present in the nucleoplasm, both the RSV and HIV-1 Gag proteins can be extracted from euchromatin and heterochromatin fractions, complementing our recently published report indicating that HIV-1 Gag localizes with euchromatin marks at the nuclear periphery (9, 10). These results suggest that HIV-1 Gag may be specifically targeted to a chromatin-associated compartment through interactions with host nuclear binding partners. Indeed, many of the proteins identified by us and others are chromatin-associated, raising the importance of further exploring the signals that target chromatin-associated regions, examining Gag interactions with chromatin factors, and elucidating possible roles for Gag in chromatin-related functions. To better understand how these chromatin-associated factors interact, and to identify which histone-associated protein networks appear most closely associated with HIV-1 Gag, we employed the use of the STRING Consortium database (**Fig 6**). Of those identified, histones H1 (8 proteins), H2A (14 proteins), and H2B (11 proteins) displayed the strongest evidence for association. Histones H3 and H4 were also identified, but each only displayed a single association. Additionally, four histone deacetylases (HDACs)—including the transcription-repressing class II HDACs 4 and 6—and seven members of the SWI/SNF chromatin remodeling complex (e.g. SMARCC2, SMARCA5, DEK, BAZ1A, MAZ1B, ARID1B, and ARID2) were identified (104). SWI/SNF complexes are highly enriched at transcriptional enhancer regions, where they modulate accessibility to promote gene activations (105). Interestingly, SWI/SNF complexes have been shown to play a role in HIV-1 infection (106–108), and also interact with histones H2A, H2B, and H4. Given the spatial organization of the nucleosome, it is possible that HIV-1 Gag may be interacting with factors residing near histones H1, H2A, H2B, and H4, to locate viral transcription sites, influence transcription of viral or cellular genes, or facilitate chromatin remodeling to expose the integrated provirus.

**Figure 6.**
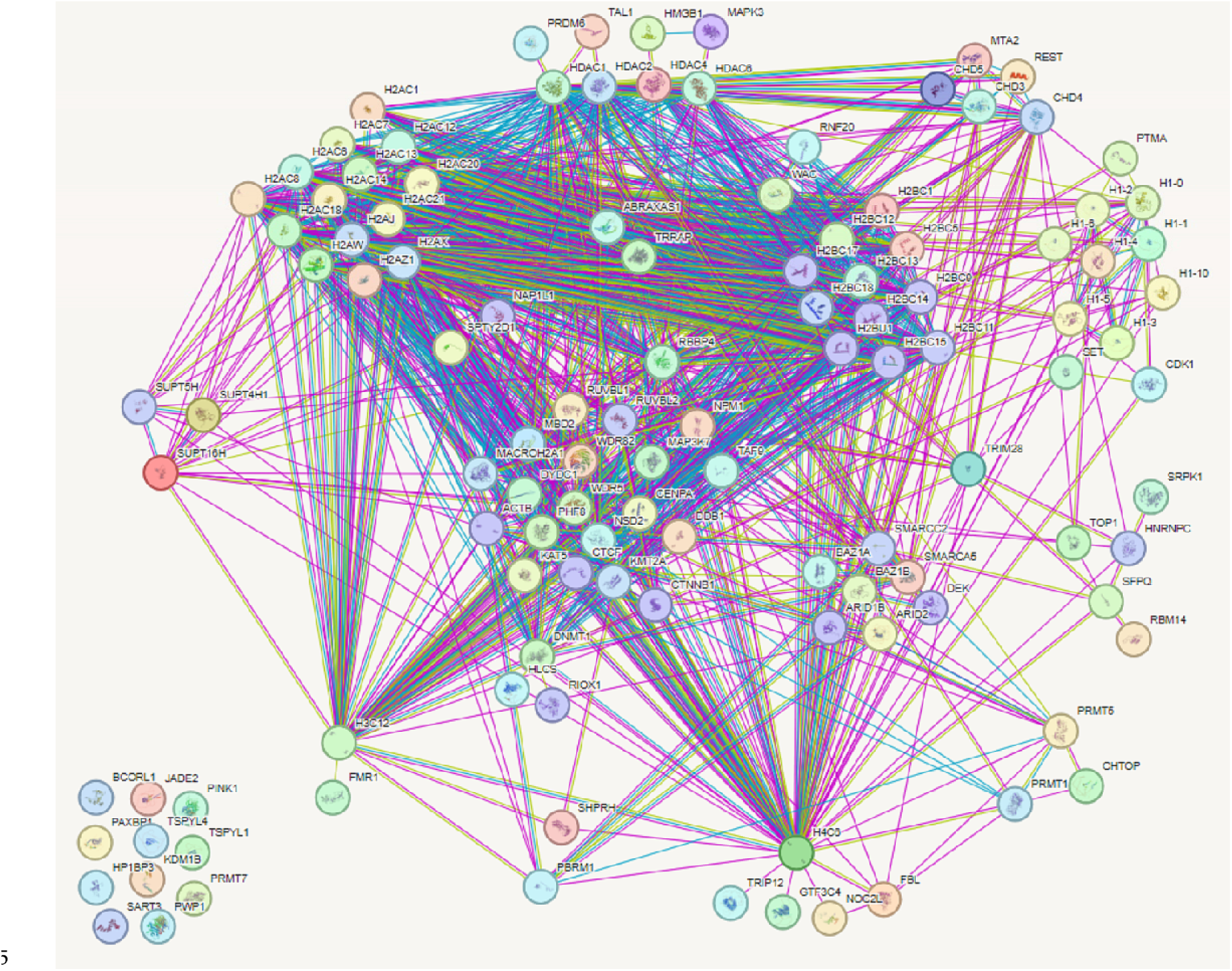
STRING protein network map of HIV-1 Gag interacting host chromatin proteins. Protein lists were generated from mass spectrometry experiments of Gag interacting proteins. Gene lists were then categorized into gene ontology (GO) terms to identify those which were chromatin associated. This refined list was input into STRING Consortium v12.0 (https://string-db.org/) to generate a protein-protein interaction map. A total of 129 proteins were queried and the following physical protein-protein interactions were made. Among the 129 proteins queried, 117 proteins are displayed after a minimum required interaction score of 0.4 was applied. Proteins that do not have any interacting partners are shown at the bottom left corner of the map. Lines are generated based on known interactions (purple lines), predicted gene fusions (red lines), predicted gene neighborhoods (green lines), and predicted gene co-occurrences (blue lines) from the literature.

It will be important to investigate whether these interactions facilitate viral replication steps occurring in the nucleus, such as gene expression, viral RNA processing and splicing, unspliced viral RNA binding by Gag, and/or nuclear export of viral RNA complexes, or whether Gag influences cellular processes for the benefit of the virus, for example by modulating expression of cellular genes involved in the immune response or altering splicing of host genes to change their localization or function. **Figure 5** highlights the various nuclear processes that HIV-1 Gag could be involved in based on the proteins identified in the HIV-1 proteomic studies.

Our published data indicating that RSV and HIV-1 Gag each interact specifically with their cognate USvRNAs to form discrete foci to form viral RNP complexes in the interchromatin space raise the possibility that interaction with host factors facilitates co-transcriptional retroviral genome selection (8, 9). In both viruses, these findings were confirmed in infected cells, where Gag was targeted to the USvRNA transcription site, presumably at the site of proviral integration. Among other transcription-related proteins, in this study we identified multiple members of the Mediator complex, which has been shown to be hijacked by other viruses and retroelements, and we demonstrated that RSV Gag colocalized and co-immunoprecipitated with Med26 (**Fig 4 and Video S1**) (36–43). Quantitative analysis of our imaging experiments indicated that nearly half of the nuclear Gag signal was colocalized with Med26. Visual inspection of the colocalization channel revealed that much of this colocalization was at Gag foci in the perichromatin space (**Fig 4A**). Our previous work demonstrating that RSV and HIV-1 Gag proteins can form biomolecular condensates (BMCs) (109, 110), combined with evidence for the existence of Mediator-containing transcriptional condensates (111, 112), argues that Gag may be utilizing condensate-driven compartmentalization to increase its chances of encountering nascent viral RNA. Further studies will address this intriguing possibility.

Additionally, we identified two components of the WMM complex, CBLL1 (Cbl Pro-to-Oncogene Like 1; present in RSV and HIV-1 datasets) and WTAP (Wilms tumor 1 associated protein; RSV dataset only), which mediates m6-methyladenosine (m6A) methylation of RNAs. This RNA modification affects RNA splicing and processing, and full length HIV-1 RNA containing m6A has displayed a bias towards serving as template RNA for the translation of viral proteins, as opposed to being packaged as gRNA (103). Interaction with components of the WMM complex could simply serve to localize Gag to sites where viral RNA may exist, but it is intriguing to consider the possibility that Gag antagonizes the function of this complex to maintain a pool of genomic RNA.

For RSV, we have observed movement of viral RNPs across the nuclear membrane, suggesting that these complexes may ultimately be packaged into assembling virions at the plasma membrane (8). We also previously found that RSV Gag colocalizes with splicing factors SF2 and SC35 (67) in splicing speckles, which are located in the perichromatin space, raising the possibility that Gag may influence splicing of viral and/or cellular RNAs in a co-transcriptional manner. It would be of interest to test the hypothesis that RSV Gag suppresses splicing at viral intron/exon junctions to promote retention of full-length viral RNA for use as the viral genome.

**Tables 3-6** list the proteins involved in transcription and splicing identified in the RSV and HIV-1 Gag interactomes. We noted that there were Gag-interacting proteins identified at different stages of transcription, including initiation, elongation, and termination. Interestingly, PolR2B, the second largest subunit of RNA polymerase II (RNAPII), was identified in both the RSV and HIV-1 interactome datasets, raising the possibility that Gag interacts with RNAPII itself (113, 114). Each dataset also contained a member of the Elongin complex (TCEB1, or Elongin C, for HIV-1; TCEB3, or Elongin A, for RSV), which forms a complex with PolR2B to promote elongation (115). Numerous members of the Integrator complex were also identified in the datasets (VWA9, RSV and HIV-1; INTS3 and INTS5, RSV only; ASUN, INTS5, INTS9, and CPSF3L, HIV-1 only). Integrator regulates transcription by binding to the C-terminal tail of PolR2A (the largest subunit of RNAPII) during pausing, which results in cleavage of newly synthesized RNA and termination of transcription (116–118). Other factors that modulate transcription identified were LEO1 and SCAF8 (both RSV and HIV-1), CDK13 and ZNF326 (RSV only), and ALYREF, CCAR2, CDK12, SETD2, and SUPT16H (HIV-1 only). One intriguing idea is that Gag could alter transcription elongation rates or regulate pausing of nascent viral RNA synthesis to promote folding of the psi region, recruit essential host RNA binding factors, modulate or suppress termination or splicing of viral RNA, or alter other co-transcriptional processes that could have downstream effects on the fate of viral RNA.

**Table 6.**
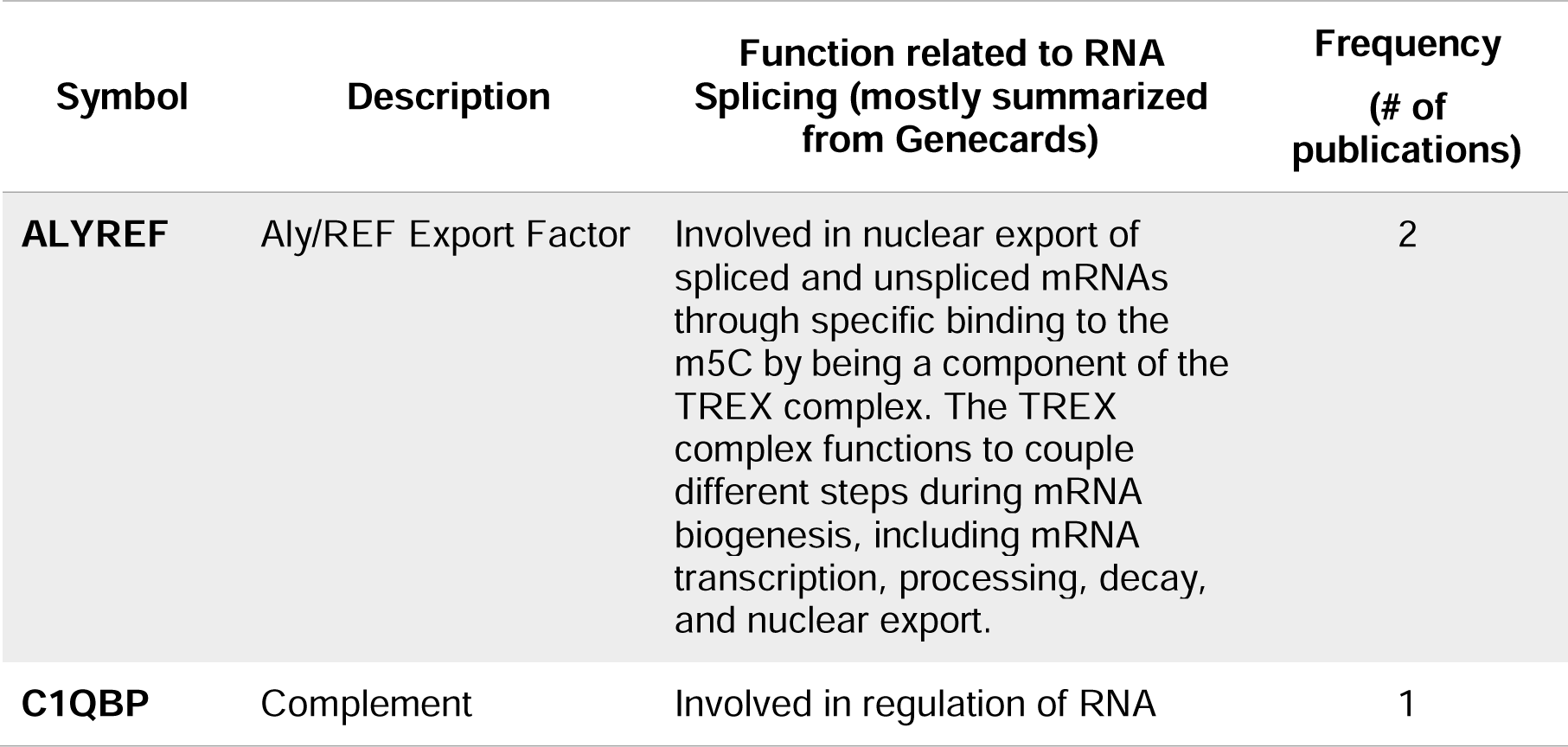
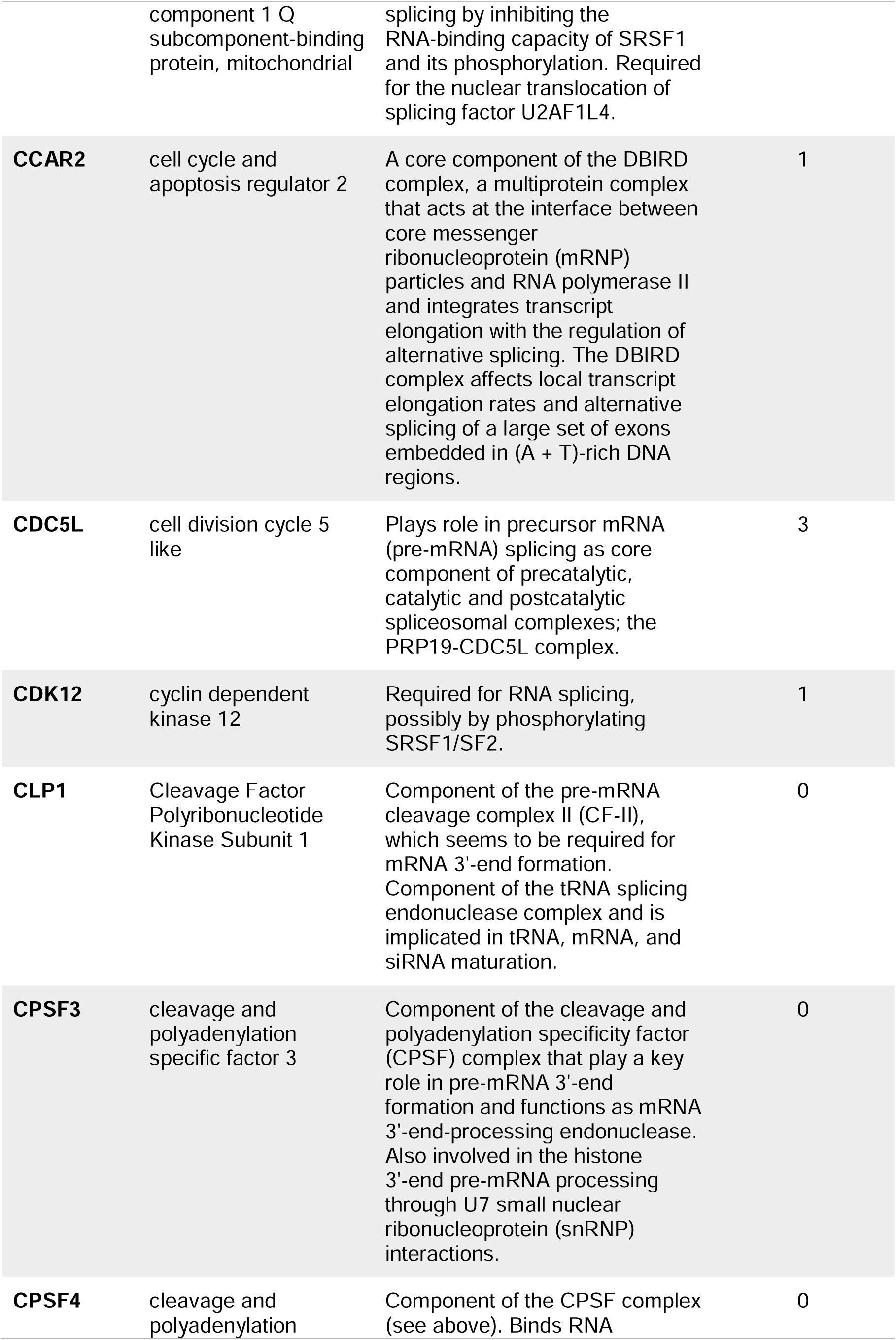
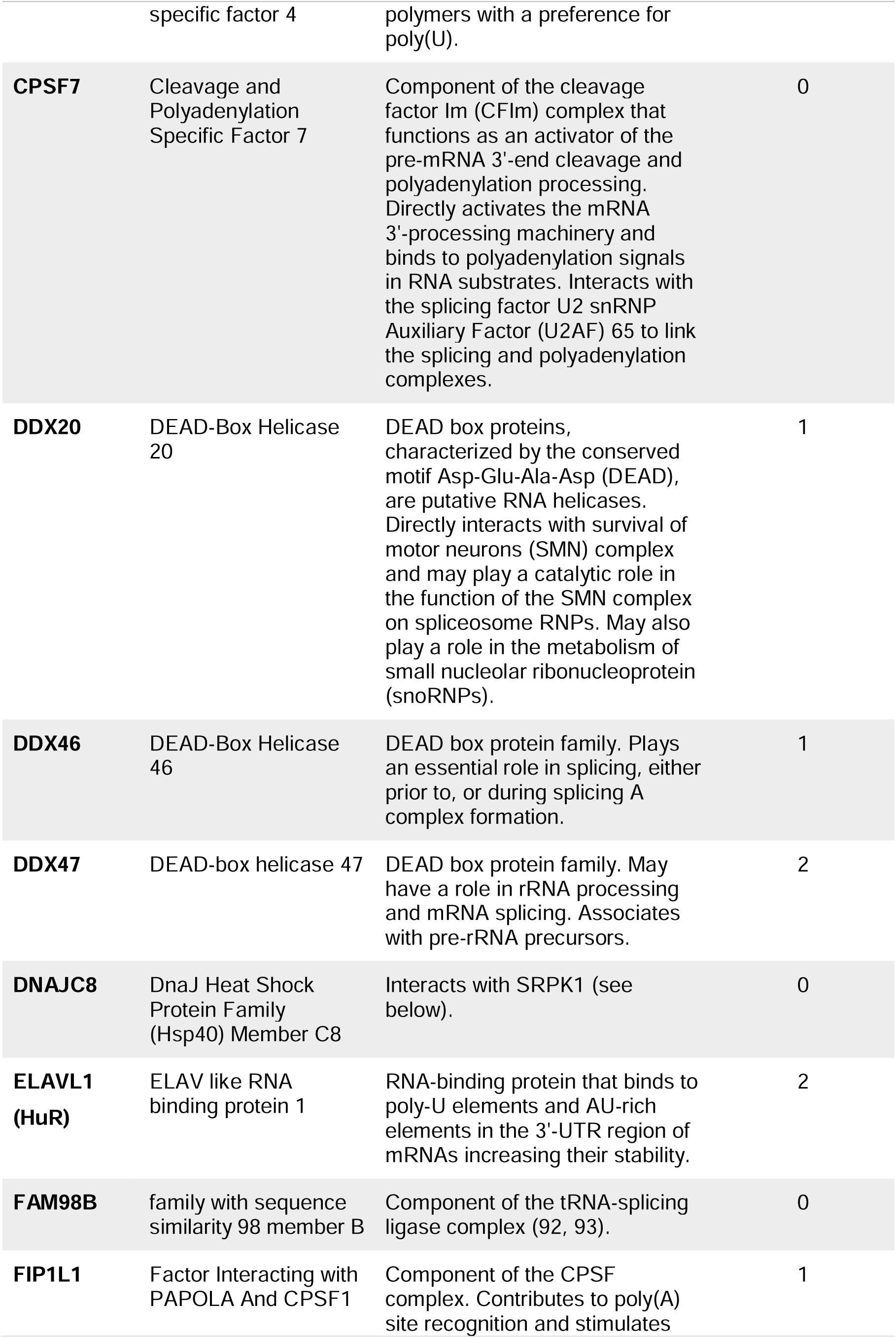
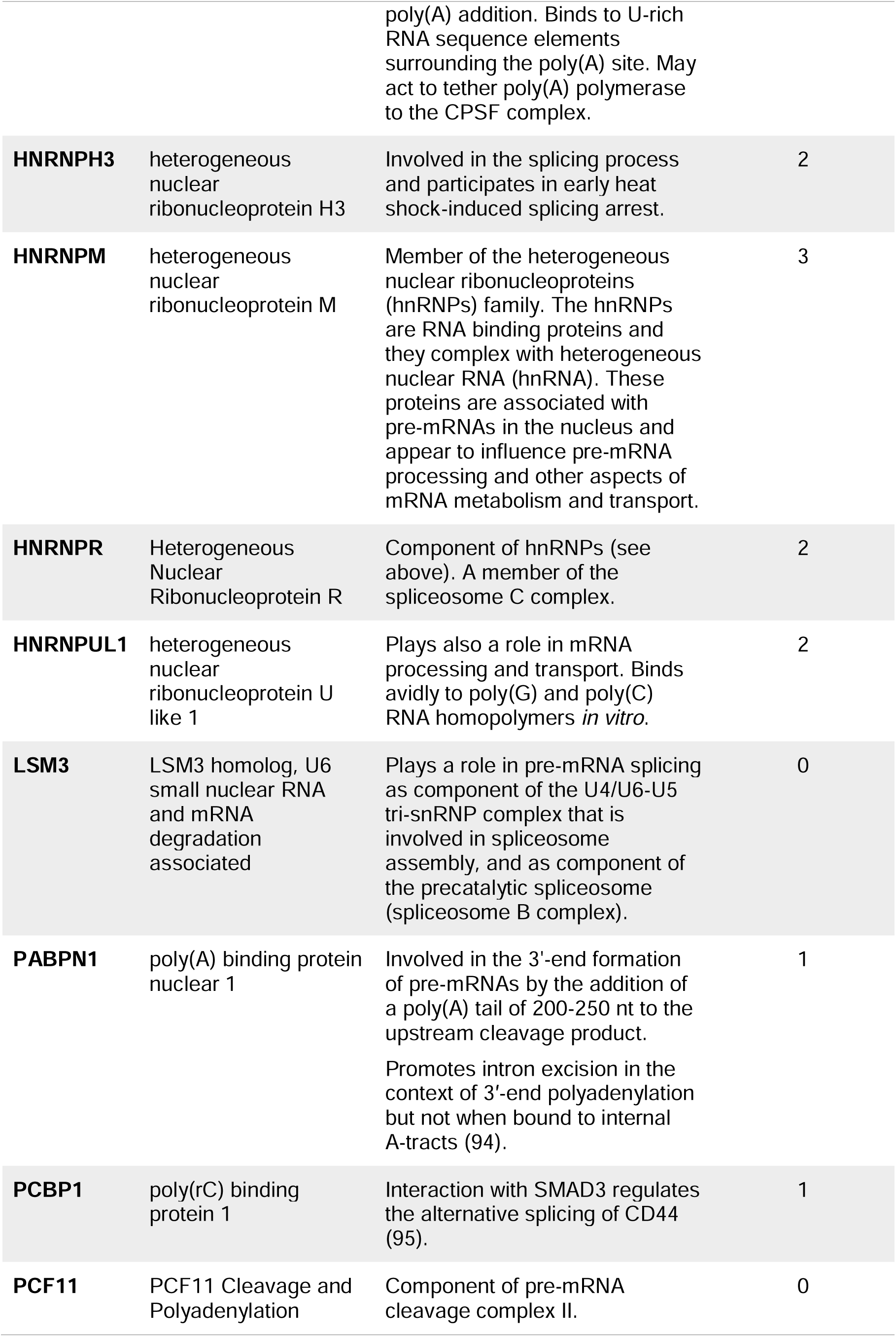
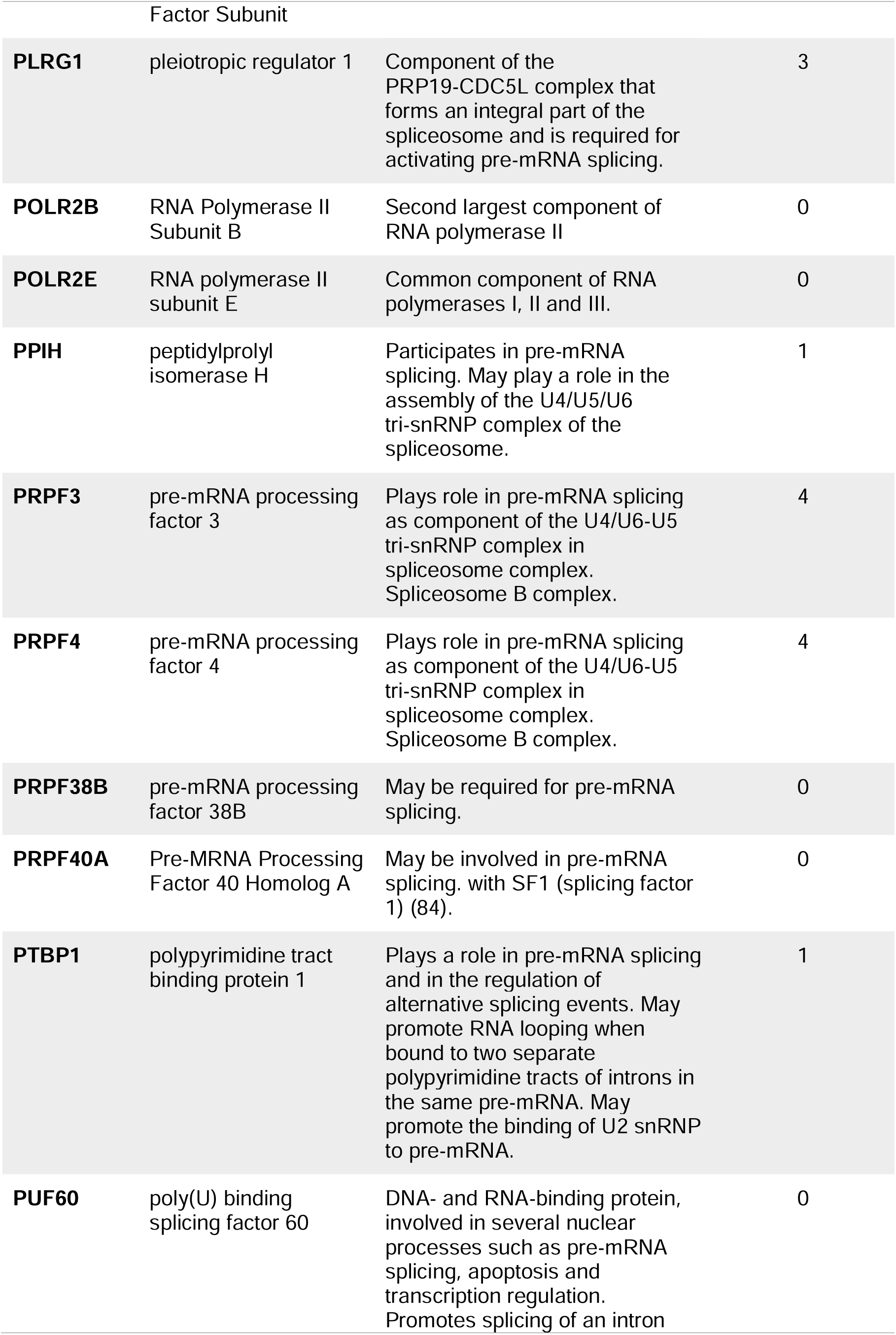
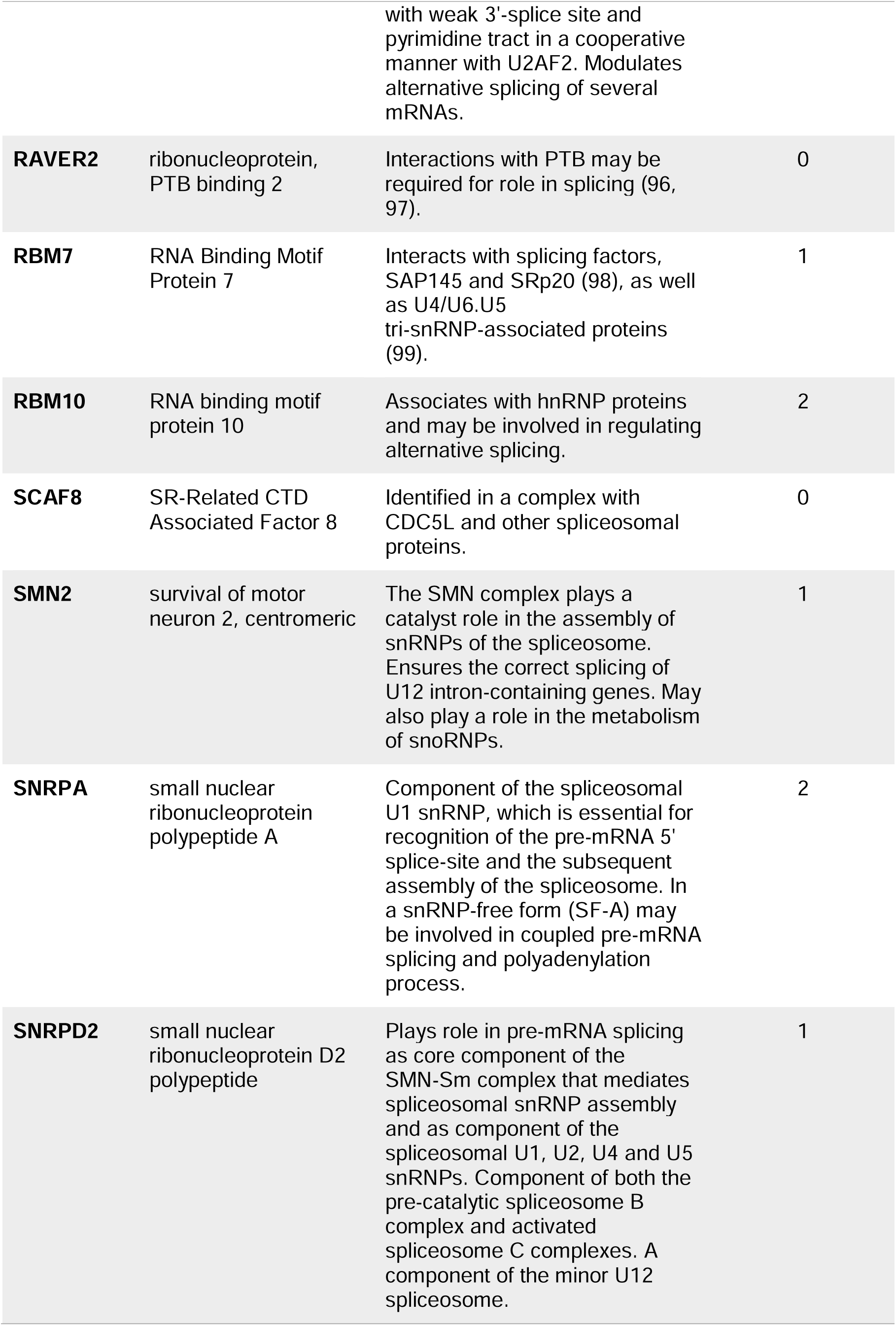
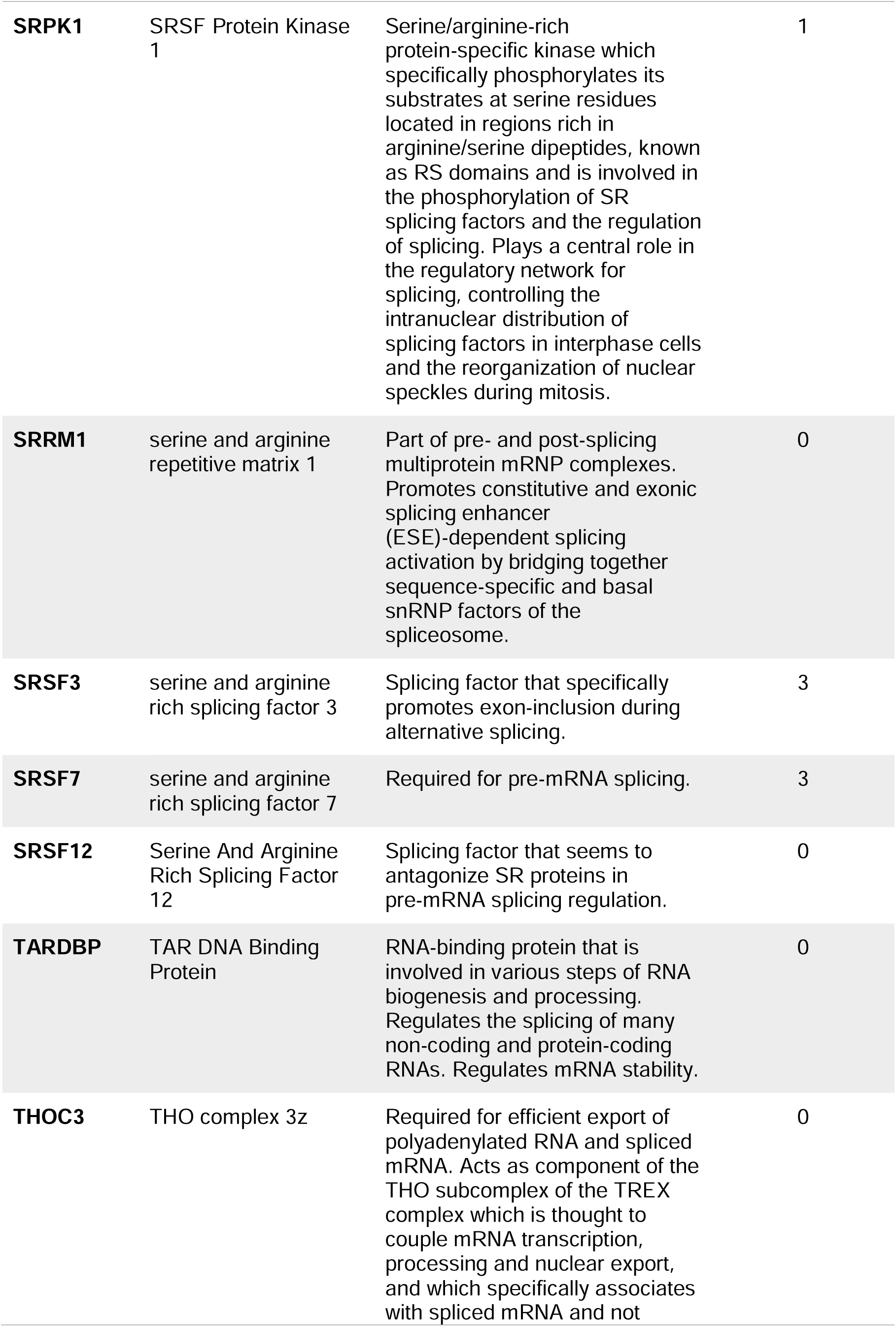
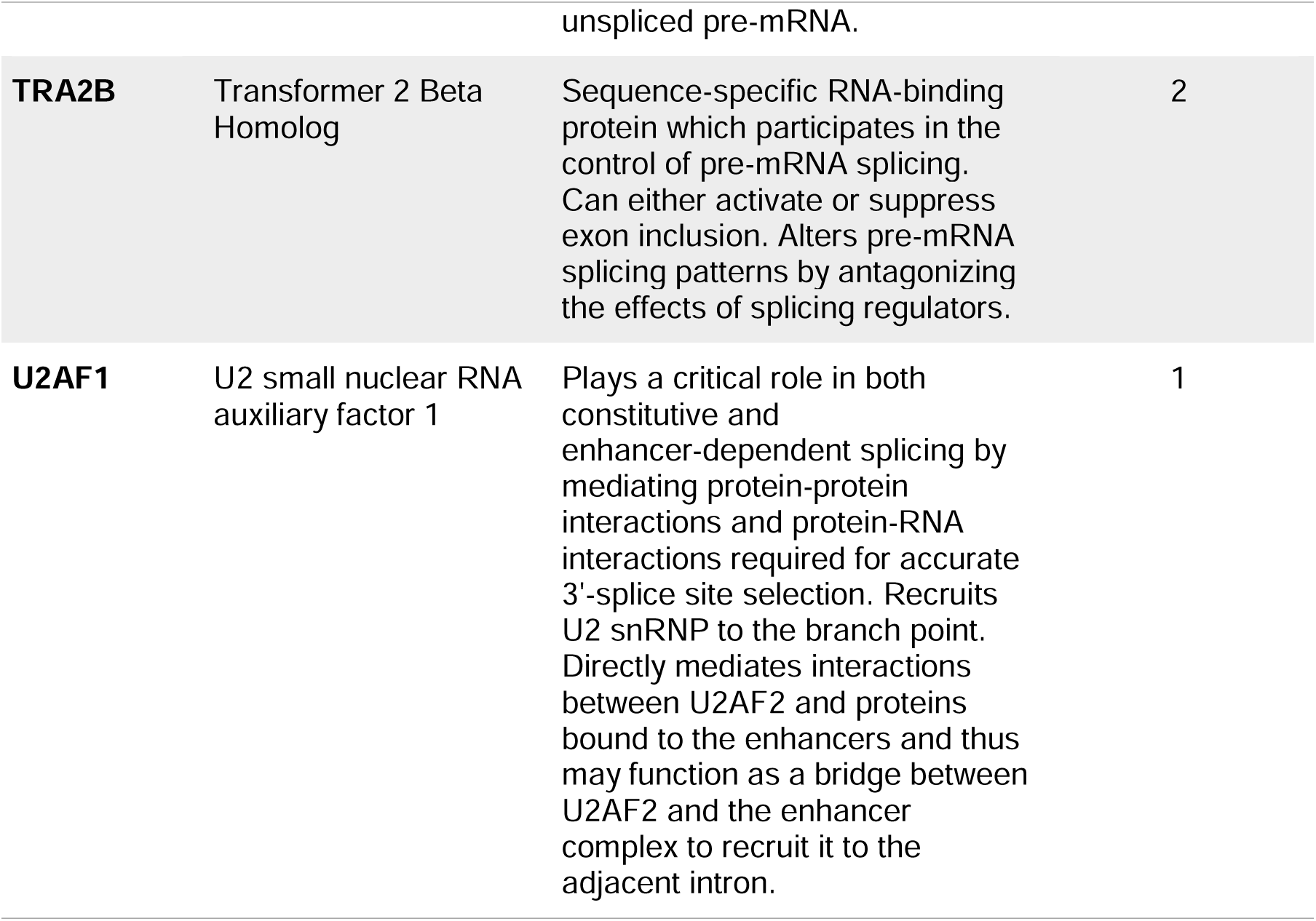
Names and function of the proteins identified in the HIV-1 proteomics list under the GO:0008380∼RNA splicing.

The data presented here, along with our analysis of previously published HIV-1 Gag interactomes, indicate that many potential host protein partners of RSV and HIV-1 Gag reside in the nucleus in association with chromatin. As both RSV and HIV-1 Gag have nuclear populations that form vRNPs in the perichromatin region, the interaction of Gag proteins with chromatin-associated nuclear factors merits further investigation. Novel roles for Gag proteins of RSV, HIV-1, and other retroviruses may be uncovered, shedding light on previously unknown aspects of the replication cycle that could be targeted by antiviral therapies.

## Supplementary Materials

Spreadsheet S1. RSV Gag Dataset 1.xlsx;

Spreadsheet S2. RSV Gag Dataset 2.xlsx;

Spreadsheet S3. HIV Gag Dataset 1.xlsx;

Spreadsheet S4. HIV Gag Dataset 2.xlsx

Table S1. Top 10 DAVID biological processes for proteins isolated from the RSV Gag affinity purifications from DF1 nuclear lysates;

Table S2. Top 10 nuclear enriched DAVID biological processes for proteins isolated from the RSV Gag affinity purifications from DF1 nuclear lysates;

Table S3. Top 10 DAVID biological processes for proteins isolated from the purified HIV Gag affinity purifications from HeLa nuclear lysates;

Table S4. Top 10 nuclear enriched DAVID biological processes for proteins isolated from the purified HIV Gag affinity purifications from HeLa nuclear lysates;

Table S5. Top 10 DAVID biological processes of nuclear proteins identified in Engeland *et al*., 2011 (31);

Table S6. Top 10 DAVID biological processes of nuclear proteins identified in Engeland *et al*., 2014 (30);

Table S7. Top 10 DAVID biological processes of nuclear proteins identified in Jäger *et al*., 2012 (32);

Table S8. Top 10 DAVID biological processes of nuclear proteins identified in Ritchie *et al*., 2015 (35);

Table S9. Top 10 DAVID biological processes of nuclear proteins identified in Le Sage *et al*., 2015 (33);

Table S10. Top 10 DAVID biological processes of nuclear proteins identified in Li *et al*., 2016 (34);

Video S1. RSV Gag and Mediator complex subunit 26 (Med26) colocalize in transfected QT6 cells

## Funding

This project was funded in part by NIH awards P50GM103297 (LJP), R01CA076534 (LJP), R01 GM139392 (LJP), R21/R33 DA053689 (LJP), T32 CA60395 (BLR), and F31 CA196292 (BLR). This project was also supported by the Penn State College of Medicine’s Comprehensive Health Studies Program. This content is solely the responsibility of the authors and does not necessarily represent the official views of the National Institutes of Health.

## Acknowledgments

We would like to acknowledge the expert assistance of Anne Stanley and Bruce Stanley in the PSU College of Medicine Mass Spectrometry and Proteomics Core, for their assistance with the mass spectrometry experiments and analysis. We greatly appreciate and acknowledge John Flanagan in the Department of Biochemistry and Molecular Biology at Penn State College of Medicine for purification of the recombinant RSV and HIV-1 Gag proteins. We acknowledge Bradley Winters at the Penn State College of Medicine for creating the illustration in Figure 5. We thank members of the Parent laboratory, Eunice Chen, Malgorzata Sudol, and Kevin Tuffy, for insightful discussions and suggestions. The Mass Spectrometry and Proteomics Core (RRID:SCR_017831) and Advanced Light Microscopy core (RRID:SCR_022526) services and instruments used in this project were funded, in part, by the Pennsylvania State University College of Medicine via the Office of the Vice Dean of Research and Graduate Students and the Pennsylvania Department of Health using Tobacco Settlement Funds (CURE). The content is solely the responsibility of the authors and does not necessarily represent the official views of the University or College of Medicine. The Pennsylvania Department of Health specifically disclaims responsibility for any analyses, interpretations or conclusions.

## Conflicts of Interest

The authors declare no conflict of interest.

## Notes

### Competing Interest Statement

The authors have declared no competing interest.

### Summary of Updates

References were reformatted; mass spectrometry proteomics data were deposited to the ProteomeXchange Consortium PRIDE partner repository.

## References

1. Butterfield-Gerson KL, Scheifele LZ, Ryan EP, Hopper AK, Parent LJ. Importin-β Family Members Mediate Alpharetrovirus Gag Nuclear Entry via Interactions with Matrix and Nucleocapsid. J Virol. 2006;80(4):1798–806.

2. Gudleski N, Flanagan JM, Ryan EP, Bewly MC, Parent LJ. Directionality of nucleocytoplasmic transport of the retroviral Gag protein depends on sequential binding of karyopherins and viral RNA. Proceedings of the National Academy of Sciences. 2010;107(20):9358–63.

3. Scheifele LZ, Garbitt RA, Rhoads JD, Parent LJ. Nuclear entry and CRM1-dependent nuclear export of the Rous sarcoma virus Gag polyprotein. Proceedings of the National Academy of Sciences. 2002;99(6):3944–9.

4. Scheifele LZ, Ryan EP, Parent LJ. Detailed Mapping of the Nuclear Export Signal in the Rous Sarcoma Virus Gag Protein. J Virol. 2005;79(14):8732–41.

5. Scheifele LZ, Kenney SP, Cairns TM, Craven RC, Parent LJ. Overlapping roles of the Rous sarcoma virus Gag p10 domain in nuclear export and virion core morphology. J Virol. 2007;81(19):10718–28.

6. Rice BL, Stake MS, Parent LJ. TNPO3-Mediated Nuclear Entry of the Rous Sarcoma Virus Gag Protein Is Independent of the Cargo-Binding Domain. J Virol. 2020;94(17).

7. Garbitt-Hirst R, Kenney SP, Parent LJ. Genetic evidence for a connection between Rous sarcoma virus Gag nuclear trafficking and genomic RNA packaging. JVirol. 2009;83(13):6790–7.

8. Maldonado RJK, Rice B, Chen EC, Tuffy KM, Chiari EF, Fahrbach KM, et al. Visualizing Association of the Retroviral Gag Protein with Unspliced Viral RNA in the Nucleus. mBio. 2020;11(2):e00524–20.

9. Tuffy KM, Maldonado RJK, Chang J, Rosenfeld P, Cochrane A, Parent LJ. HIV-1 Gag Forms Ribonucleoprotein Complexes with Unspliced Viral RNA at Transcription Sites. Viruses. 2020;12(11).

10. Chang J, Parent LJ. HIV-1 Gag colocalizes with euchromatin histone marks at the nuclear periphery. bioRxiv. 2023.

11. Lochmann TL, Bann DV, Ryan EP, Beyer AR, Mao A, Cochrane A, et al. NC-Mediated Nucleolar Localization of Retroviral Gag Proteins. Virus research. 2013;171(2):304–18.

12. Gallay P, Swingler S, Song J, Bushman F, Trono D. HIV nuclear import is governed by the phosphotyrosine-mediated binding of matrix to the core domain of integrase. Cell. 1995;83(4):569–76.

13. Zhang J, Crumpacker CS. Human immunodeficiency virus type 1 nucleocapsid protein nuclear localization mediates early viral mRNA expression. J Virol. 2002;76(20):10444–54.

14. Kemler I, Meehan A, Poeschla EM. Live-Cell Coimaging of the Genomic RNAs and Gag Proteins of Two Lentiviruses. J Virol. 2010;84(13):6352–66.

15. Kemler I, Saenz D, Poeschla E. Feline Immunodeficiency Virus Gag Is a Nuclear Shuttling Protein. J Virol. 2012;86(16):8402–11.

16. Bohl CR, Brown SM, Weldon RA. The pp24 phosphoprotein of Mason-Pfizer monkey virus contributes to viral genome packaging. Retrovirology. 2005;2(1):68.

17. Weldon RA, Sarkar P, Brown SM, Weldon SK. Mason–Pfizer monkey virus Gag proteins interact with the human sumo conjugating enzyme, hUbc9. Virology. 2003;314(1):62–73.

18. Baluyot MF, Grosse SA, Lyddon TD, Janaka SK, Johnson MC. CRM1-dependent trafficking of retroviral Gag proteins revisited. J Virol. 2012;86(8):4696–700.

19. Beyer AR, Bann DV, Rice B, Pultz IS, Kane M, Goff SP, et al. Nucleolar Trafficking of the Mouse Mammary Tumor Virus Gag Protein Induced by Interaction with Ribosomal Protein L9. J Virol. 2013;87(2):1069–82.

20. Elis E, Ehrlich M, Prizan-Ravid A, Laham-Karam N, Bacharach E. p12 Tethers the Murine Leukemia Virus Pre-integration Complex to Mitotic Chromosomes. PLOS Pathogens. 2012;8(12):e1003103.

21. Nash MA, Meyer MK, Decker GL, Arlinghaus RB. A subset of Pr65Gag is nucleus associated in murine leukemia virus-infected cells. J Virol. 1993;67(3):1350–6.

22. Risco C, Menendez-Arias L, Copeland TD, Pinto da Silva P, Oroszlan S. Intracellular transport of the murine leukemia virus during acute infection of NIH 3T3 cells: nuclear import of nucleocapsid protein and integrase. Journal of Cell Science. 1995;108(9):3039.

23. Schneider WM, Brzezinski JD, Aiyer S, Malani N, Gyuricza M, Bushman FD, et al. Viral DNA tethering domains complement replication-defective mutations in the p12 protein of MuLV Gag. Proceedings of the National Academy of Sciences. 2013;110(23):9487–92.

24. Müllers E, Stirnnagel K, Kaulfuss S, Lindemann D. Prototype foamy virus Gag nuclear localization: a novel pathway among retroviruses. J Virol. 2011;85(18):9276–85.

25. Renault N, Tobaly-Tapiero J, Paris J, Giron M-L, Coiffic A, Roingeard P, et al. A nuclear export signal within the structural Gag protein is required for prototype foamy virus replication. Retrovirology. 2011;8(1):6.

26. Schliephake AW, Rethwilm A. Nuclear localization of foamy virus Gag precursor protein. J Virol. 1994;68(8):4946–54.

27. Tobaly-Tapiero J, Bittoun P, Lehmann-Che J, Delelis O, Giron ML, de Thé H, et al. Chromatin tethering of incoming foamy virus by the structural Gag protein. Traffic. 2008;9(10):1717–27.

28. Yu KL, Lee SH, Lee ES, You JC. HIV-1 nucleocapsid protein localizes efficiently to the nucleus and nucleolus. Virology. 2016;492:204–12.

29. Brzezinski JD, Modi A, Liu M, Roth MJ. Repression of the Chromatin-Tethering Domain of Murine Leukemia Virus p12. J Virol. 2016;90(24):11197–207.

30. Engeland CE, Brown NP, Börner K, Schümann M, Krause E, Kaderali L, et al. Proteome analysis of the HIV-1 Gag interactome. Virology. 2014;460–461:194-206.

31. Engeland CE, Oberwinkler H, Schümann M, Krause E, Müller GA, Kräusslich H-G. The Cellular Protein Lyric Interacts with HIV-1 Gag. J Virol. 2011;85(24):13322–32.

32. Jäger S, Cimermancic P, Gulbahce N, Johnson JR, McGovern KE, Clarke SC, et al. Global landscape of HIV–human protein complexes. Nature. 2011;481:365.

33. Le Sage V, Cinti A, Valiente-Echeverría F, Mouland AJ. Proteomic analysis of HIV-1 Gag interacting partners using proximity-dependent biotinylation. Virology Journal. 2015;12(1):138.

34. Li Y, Frederick KM, Haverland NA, Ciborowski P, Belshan M. Investigation of the HIV-1 matrix interactome during virus replication. PROTEOMICS – Clinical Applications. 2016;10(2):156–63.

35. Ritchie C, Cylinder I, Platt EJ, Barklis E. Analysis of HIV-1 Gag Protein Interactions via Biotin Ligase Tagging. J Virol. 2015;89(7):3988–4001.

36. Kornberg RD. Mediator and the mechanism of transcriptional activation. Trends Biochem Sci. 2005;30(5):235–9.

37. Boyer TG, Martin ME, Lees E, Ricciardi RP, Berk AJ. Mammalian Srb/Mediator complex is targeted by adenovirus E1A protein. Nature. 1999;399(6733):276-9.

38. Yang M, Hay J, Ruyechan WT. Varicella-zoster virus IE62 protein utilizes the human mediator complex in promoter activation. J Virol. 2008;82(24):12154–63.

39. Lester JT, DeLuca NA. Herpes simplex virus 1 ICP4 forms complexes with TFIID and mediator in virus-infected cells. J Virol. 2011;85(12):5733–44.

40. Vijayalingam S, Chinnadurai G. Adenovirus L-E1A activates transcription through mediator complex-dependent recruitment of the super elongation complex. J Virol. 2013;87(6):3425–34.

41. Pei J, Beri NR, Zou AJ, Hubel P, Dorando HK, Bergant V, et al. Nuclear-localized human respiratory syncytial virus NS1 protein modulates host gene transcription. Cell Rep. 2021;37(2):109803.

42. Rovnak J, Quackenbush SL. Exploitation of the Mediator complex by viruses. PLoS Pathog. 2022;18(4):e1010422.

43. Asimi V, Sampath Kumar A, Niskanen H, Riemenschneider C, Hetzel S, Naderi J, et al. Hijacking of transcriptional condensates by endogenous retroviruses. Nat Genet. 2022.

44. Himly M, Foster DN, Bottoli I, Iacovoni JS, Vogt PK. The DF-1 Chicken Fibroblast Cell Line: Transformation Induced by Diverse Oncogenes and Cell Death Resulting from Infection by Avian Leukosis Viruses. Virology. 1998;248(2):295–304.

45. Craven RC, Leure-duPree AE, Weldon RA, Wills JW. Genetic analysis of the major homology region of the Rous sarcoma virus Gag protein. J Virol. 1995;69(7):4213–27.

46. Kenney SP, Lochmann TL, Schmid CL, Parent LJ. Intermolecular Interactions between Retroviral Gag Proteins in the Nucleus. J Virol. 2008;82(2):683–91.

47. Bewley MC, Reinhart L, Stake MS, Nadaraia-Hoke S, Parent LJ, Flanagan JM. A non-cleavable hexahistidine affinity tag at the carboxyl-terminus of the HIV-1 Pr55Gag polyprotein alters nucleic acid binding properties. Protein Expression and Purification. 2017;130(Supplement C):137–45.

48. Rye-McCurdy TD, Nadaraia-Hoke S, Gudleski-O’Regan N, Flanagan JM, Parent LJ, Musier-Forsyth K. Mechanistic Differences between Nucleic Acid Chaperone Activities of the Gag Proteins of Rous Sarcoma Virus and Human Immunodeficiency Virus Type 1 Are Attributed to the MA Domain. J Virol. 2014;88(14):7852–61.

49. Fujiwara T, Oda K, Yokota S, Takatsuki A, Ikehara Y. Brefeldin A causes disassembly of the Golgi complex and accumulation of secretory proteins in the endoplasmic reticulum. Journal of Biological Chemistry. 1988;263(34):18545–52.

50. Chase GP, Rameix-Welti M-A, Zvirbliene A, Zvirblis G, Götz V, Wolff T, et al. Influenza Virus Ribonucleoprotein Complexes Gain Preferential Access to Cellular Export Machinery through Chromatin Targeting. PLOS Pathogens. 2011;7(9):e1002187.

51. Weldon RA, Erdie CR, Oliver MG, Wills JW. Incorporation of chimeric Gag protein into retroviral particles. J Virol. 1990;64(9):4169–79.

52. Shilov IV, Seymour SL, Patel AA, Loboda A, Tang WH, Keating SP, et al. The Paragon Algorithm, a Next Generation Search Engine That Uses Sequence Temperature Values and Feature Probabilities to Identify Peptides from Tandem Mass Spectra. Molecular & Cellular Proteomics. 2007;6(9):1638–55.

53. Tang WH, Shilov IV, Seymour SL. Nonlinear Fitting Method for Determining Local False Discovery Rates from Decoy Database Searches. Journal of Proteome Research. 2008;7(9):3661–7.

54. Perez-Riverol Y, Bai J, Bandla C, García-Seisdedos D, Hewapathirana S, Kamatchinathan S, et al. The PRIDE database resources in 2022: a hub for mass spectrometry-based proteomics evidences. Nucleic Acids Res. 2022;50(D1):D543–D52.

55. Huang DW, Sherman BT, Lempicki RA. Systematic and integrative analysis of large gene lists using DAVID bioinformatics resources. Nature Protocols. 2008;4:44.

56. Huang DW, Sherman BT, Lempicki RA. Bioinformatics enrichment tools: paths toward the comprehensive functional analysis of large gene lists. Nucleic Acids Research. 2009;37(1):1–13.

57. Lippé R. Deciphering Novel Host–Herpesvirus Interactions by Virion Proteomics. Frontiers in Microbiology. 2012;3(181).

58. Nojima T, Gomes T, Grosso ARF, Kimura H, Dye MJ, Dhir S, et al. Mammalian NET-Seq Reveals Genome-wide Nascent Transcription Coupled to RNA Processing. Cell. 2015;161(3):526–40.

59. Sato S, Tomomori-Sato C, Parmely TJ, Florens L, Zybailov B, Swanson SK, et al. A set of consensus mammalian mediator subunits identified by multidimensional protein identification technology. Mol Cell. 2004;14(5):685–91.

60. Kim DI, KC B, Zhu W, Motamedchaboki K, Doye V, Roux KJ. Probing nuclear pore complex architecture with proximity-dependent biotinylation. Proceedings of the National Academy of Sciences. 2014:201406459.

61. Mechold U, Gilbert C, Ogryzko V. Codon optimization of the BirA enzyme gene leads to higher expression and an improved efficiency of biotinylation of target proteins in mammalian cells. J Biotechnol. 2005;116(3):245–9.

62. Roux KJ, Kim DI, Raida M, Burke B. A promiscuous biotin ligase fusion protein identifies proximal and interacting proteins in mammalian cells. Journal of Cell Biology. 2012;196(6):801–10.

63. Gao W, Li M, Zhang J. Tandem immunoprecipitation approach to identify HIV-1 Gag associated host factors. J Virol Methods. 2014;203:116–9.

64. Fu W, Sanders-Beer BE, Katz KS, Maglott DR, Pruitt KD, Ptak RG. Human immunodeficiency virus type 1, human protein interaction database at NCBI. Nucleic Acids Res. 2009;37(Database issue):15.

65. Ptak RG, Fu W, Sanders-Beer BE, Dickerson JE, Pinney JW, Robertson DL, et al. Cataloguing the HIV type 1 human protein interaction network. AIDS Res Hum Retroviruses. 2008;24(12):1497–502.

66. Pinney JW, Dickerson JE, Fu W, Sanders-Beer BE, Ptak RG, Robertson DL. HIV-host interactions: a map of viral perturbation of the host system. Aids. 2009;23(5):549–54.

67. Rice B, Kaddis R, Stake M, Lochmann T, Parent L. Interplay between the alpharetroviral Gag protein and SR Proteins SF2 and SC35 in the nucleus. Frontiers in Microbiology. 2015;6(925).

68. Jassal B, Matthews L, Viteri G, Gong C, Lorente P, Fabregat A, et al. The reactome pathway knowledgebase. Nucleic Acids Res. 2020;48(D1):D498–D503.

69. Wu G, Haw R. Functional Interaction Network Construction and Analysis for Disease Discovery. Methods Mol Biol. 2017:6783–4_11.

70. Kornblihtt AR, de la Mata M, Fededa JP, Muñoz MJ, Nogués G. Multiple links between transcription and splicing. RNA. 2004;10(10):1489–98.

71. Brody Y, Shav-Tal Y. Transcription and splicing. Transcription. 2011;2(5):216–20.

72. Herzel L, Ottoz DSM, Alpert T, Neugebauer KM. Splicing and transcription touch base: co-transcriptional spliceosome assembly and function. Nature Reviews Molecular Cell Biology. 2017;18(10):637–50.

73. Tellier M, Maudlin I, Murphy S. Transcription and splicing: A two-way street. WIREs RNA. 2020;11(5):e1593.

74. Soutourina J. Transcription regulation by the Mediator complex. Nat Rev Mol Cell Biol. 2018;19(4):262–74.

75. Conaway RC, Conaway JW. Function and regulation of the Mediator complex. Curr Opin Genet Dev. 2011;21(2):225–30.

76. Conaway RC, Conaway JW. The Mediator complex and transcription elongation. Biochim Biophys Acta. 2013;1829(1):69–75.

77. Ansari SA, Morse RH. Mechanisms of Mediator complex action in transcriptional activation. Cell Mol Life Sci. 2013;70(15):2743–56.

78. Takahashi H, Parmely TJ, Sato S, Tomomori-Sato C, Banks CA, Kong SE, et al. Human mediator subunit MED26 functions as a docking site for transcription elongation factors. Cell. 2011;146(1):92–104.

79. Tan C, Zhu S, Chen Z, Liu C, Li YE, Zhu M, et al. Mediator complex proximal Tail subunit MED30 is critical for Mediator core stability and cardiomyocyte transcriptional network. PLoS Genet. 2021;17(9):e1009785.

80. Basson MA, van Ravenswaaij-Arts C. Functional Insights into Chromatin Remodelling from Studies on CHARGE Syndrome. Trends in Genetics. 2015;31(10):600–11.

81. Mo S, Ji X, Fu X-D. Unique role of SRSF2 in transcription activation and diverse functions of the SR and hnRNP proteins in gene expression regulation. Transcription. 2013;4(5):251–9.

82. Zanini IMY, Soneson C, Lorenzi LE, Azzalin CM. Human cactin interacts with DHX8 and SRRM2 to assure efficient pre-mRNA splicing and sister chromatid cohesion. Journal of Cell Science. 2017;130(4):767.

83. Fortes P, Bilbao-Cortés D, Fornerod M, Rigaut G, Raymond W, Séraphin B, et al. Luc7p, a novel yeast U1 snRNP protein with a role in 5′ splice site recognition. Genes & Development. 1999;13(18):2425–38.

84. Crisci A, Raleff F, Bagdiul I, Raabe M, Urlaub H, Rain J-C, et al. Mammalian splicing factor SF1 interacts with SURP domains of U2 snRNP-associated proteins. Nucleic Acids Research. 2015;43(21):10456–73.

85. Hegele A, Kamburov A, Grossmann A, Sourlis C, Wowro S, Weimann M, et al. Dynamic protein-protein interaction wiring of the human spliceosome. Mol Cell. 2012;45(4):567–80.

86. Wang J, Huo K, Ma L, Tang L, Li D, Huang X, et al. Toward an understanding of the protein interaction network of the human liver. Mol Syst Biol. 2011;7(536):67.

87. Gillian AL, Svaren J. The Ddx20/DP103 Dead Box Protein Represses Transcriptional Activation by Egr2/Krox-20. Journal of Biological Chemistry. 2004;279(10):9056–63.

88. Wu C, Dedhar S. Integrin-linked kinase (ILK) and its interactors : a new paradigm for the coupling of extracellular matrix to actin cytoskeleton and signaling complexes. Journal of Cell Biology. 2001;155(4):505–10.

89. Thakur S, Nakamura T, Calin G, Russo A, Tamburrino JF, Shimizu M, et al. Regulation of BRCA1 Transcription by Specific Single-Stranded DNA Binding Factors. Molecular and Cellular Biology. 2003;23(11):3774.

90. Lynch M, Chen L, Ravitz MJ, Mehtani S, Korenblat K, Pazin MJ, et al. hnRNP K Binds a Core Polypyrimidine Element in the Eukaryotic Translation Initiation Factor 4E (eIF4E) Promoter, and Its Regulation of eIF4E Contributes to Neoplastic Transformation. Molecular and Cellular Biology. 2005;25(15):6436.

91. Choi HS, Song KY, Hwang CK, Kim CS, Law P-Y, Wei L-N, et al. A proteomic approach for identification of single-strand DNA-binding proteins involved in transcriptional regulation of mouse mu-opioid receptor gene. Molecular & Cellular Proteomics. 2008.

92. Popow J, Englert M, Weitzer S, Schleiffer A, Mierzwa B, Mechtler K, et al. HSPC117 Is the Essential Subunit of a Human tRNA Splicing Ligase Complex. Science. 2011;331(6018):760-4.

93. Popow J, Jurkin J, Schleiffer A, Martinez J. Analysis of orthologous groups reveals archease and DDX1 as tRNA splicing factors. Nature. 2014;511(7507):104-7.

94. Muniz L, Davidson L, West S. Poly(A) Polymerase and the Nuclear Poly(A) Binding Protein, PABPN1, Coordinate the Splicing and Degradation of a Subset of Human Pre-mRNAs. Molecular and Cellular Biology. 2015;35(13):2218.

95. Tripathi V, Sixt KM, Gao S, Xu X, Huang J, Weigert R, et al. Direct Regulation of Alternative Splicing by SMAD3 through PCBP1 Is Essential to the Tumor-Promoting Role of TGF-beta. Mol Cell. 2016;64(3):549–64.

96. Kleinhenz B, Fabienke M, Swiniarski S, Wittenmayer N, Kirsch J, Jockusch BM, et al. Raver2, a new member of the hnRNP family. FEBS Lett. 2005;579(20):4254–8.

97. Henneberg B, Swiniarski S, Sabine B, Illenberger S. A conserved peptide motif in Raver2 mediates its interaction with the polypyrimidine tract-binding protein. Exp Cell Res. 2010;316(6):966–79.

98. Guo TB, Boros LG, Chan KC, Hikim APS, Hudson AP, Swerdloff RS, et al. Spermatogenetic Expression of RNA-Binding Motif Protein 7, a Protein That Interacts With Splicing Factors. Journal of Andrology. 2003;24(2):204–14.

99. Nag A, Steitz JA. Tri-snRNP-associated proteins interact with subunits of the TRAMP and nuclear exosome complexes, linking RNA decay and pre-mRNA splicing. RNA Biology. 2012;9(3):334–42.

100. Lesbats P, Serrao E, Maskell DP, Pye VE, O’Reilly N, Lindemann D, et al. Structural basis for spumavirus GAG tethering to chromatin. Proceedings of the National Academy of Sciences. 2017;114(21):5509.

101. Müllers E, Stirnnagel K, Kaulfuss S, Lindemann D. Prototype Foamy Virus Gag Nuclear Localization: a Novel Pathway among Retroviruses. J Virol. 2011;85(18):9276–85.

102. Tobaly-Tapiero J, Bittoun P, Lehmann-Che J, Delelis O, Giron ML, Thé HD, et al. Chromatin Tethering of Incoming Foamy Virus by the Structural Gag Protein. Traffic. 2008;9(10):1717–27.

103. Pereira-Montecinos C, Toro-Ascuy D, Ananías-Sáez C, Gaete-Argel A, Rojas-Fuentes C, Riquelme-Barrios S, et al. Epitranscriptomic regulation of HIV-1 full-length RNA packaging. Nucleic Acids Res. 2022;50(4):2302–18.

104. Thiagalingam S, Cheng KH, Lee HJ, Mineva N, Thiagalingam A, Ponte JF. Histone deacetylases: unique players in shaping the epigenetic histone code. Ann N Y Acad Sci. 2003;983:84–100.

105. Mittal P, Roberts CWM. The SWI/SNF complex in cancer - biology, biomarkers and therapy. Nat Rev Clin Oncol. 2020;17(7):435–48.

106. La Porte A, Cano J, Wu X, Mitra D, Kalpana GV. An Essential Role of INI1/hSNF5 Chromatin Remodeling Protein in HIV-1 Posttranscriptional Events and Gag/Gag-Pol Stability. J Virol. 2016;90(21):9889–904.

107. Lesbats P, Botbol Y, Chevereau G, Vaillant C, Calmels C, Arneodo A, et al. Functional coupling between HIV-1 integrase and the SWI/SNF chromatin remodeling complex for efficient in vitro integration into stable nucleosomes. PLoS Pathog. 2011;7(2):e1001280.

108. Mahmoudi T, Parra M, Vries RG, Kauder SE, Verrijzer CP, Ott M, et al. The SWI/SNF chromatin-remodeling complex is a cofactor for Tat transactivation of the HIV promoter. J Biol Chem. 2006;281(29):19960–8.

109. Kaddis Maldonado R, Lambert GS, Rice BL, Sudol M, Flanagan JM, Parent LJ. The Rous sarcoma virus Gag Polyprotein Forms Biomolecular Condensates Driven by Intrinsically-disordered Regions. J Mol Biol. 2023;435(16):168182.

110. Monette A, Niu M, Maldonado RK, Chang J, Lambert GS, Flanagan JM, et al. Influence of HIV-1 Genomic RNA on the Formation of Gag Biomolecular Condensates. J Mol Biol. 2023;435(16):168190.

111. Sabari BR, Dall’Agnese A, Boija A, Klein IA, Coffey EL, Shrinivas K, et al. Coactivator condensation at super-enhancers links phase separation and gene control. Science. 2018;361(6400).

112. Sabari BR, Dall’Agnese A, Young RA. Biomolecular Condensates in the Nucleus. Trends Biochem Sci. 2020;45(11):961–77.

113. Cramer P, Bushnell DA, Kornberg RD. Structural basis of transcription: RNA polymerase II at 2.8 angstrom resolution. Science. 2001;292(5523):1863–76.

114. Acker J, Wintzerith M, Vigneron M, Kédinger C. Primary structure of the second largest subunit of human RNA polymerase II (or B). J Mol Biol. 1992;226(4):1295–9.

115. Chen Y, Kokic G, Dienemann C, Dybkov O, Urlaub H, Cramer P. Structure of the transcribing RNA polymerase II-Elongin complex. Nat Struct Mol Biol. 2023;30(12):1925–35.

116. Fianu I, Chen Y, Dienemann C, Dybkov O, Linden A, Urlaub H, et al. Structural basis of Integrator-mediated transcription regulation. Science. 2021;374(6569):883-7.

117. Welsh SA, Gardini A. Genomic regulation of transcription and RNA processing by the multitasking Integrator complex. Nat Rev Mol Cell Biol. 2023;24(3):204–20.

118. Wagner EJ, Tong L, Adelman K. Integrator is a global promoter-proximal termination complex. Mol Cell. 2023;83(3):416–27.

