## Supplemental Captions for "Comparative analysis of retroviral Gag-host cell interactions: focus on the nuclear interactome"

Spreadsheet S1. RSV Gag Dataset 1.xlsx;

Spreadsheet S2. RSV Gag Dataset 2.xlsx;

Spreadsheet S3. HIV Gag Dataset 1.xlsx;

Spreadsheet S4. HIV Gag Dataset 2.xlsx

Table S1. Top 10 DAVID biological processes for proteins isolated from the RSV Gag affinity purifications from DF1 nuclear lysates;

Table S2. Top 10 nuclear enriched DAVID biological processes for proteins isolated from the RSV Gag affinity purifications from DF1 nuclear lysates;

Table S3. Top 10 DAVID biological processes for proteins isolated from the purified HIV Gag affinity purifications from HeLa nuclear lysates;

Table S4. Top 10 nuclear enriched DAVID biological processes for proteins isolated from the purified HIV Gag affinity purifications from HeLa nuclear lysates;

Table S5. Top 10 DAVID biological processes of nuclear proteins identified in Engeland *et al*., 2011;

Table S6. Top 10 DAVID biological processes of nuclear proteins identified in Engeland *et al*., 2014;

Table S7. Top 10 DAVID biological processes of nuclear proteins identified in Jäger *et al*., 2012;

Table S8. Top 10 DAVID biological processes of nuclear proteins identified in Ritchie *et al*., 2015;

Table S9. Top 10 DAVID biological processes of nuclear proteins identified in Le Sage *et al*., 2015

Table S10. Top 10 DAVID biological processes of nuclear proteins identified in Li *et al*., 2016;

Video S1. RSV Gag-GFP and Mediator complex subunit 26 (FLAG-Med26, detected by FLAG antibody) colocalize in transfected QT6 cells. Imaris was used to generate a 3D image, which was rotated for the first 13 seconds to show Gag (red), Med26 (green), and DAPI (blue) signal. For the next segment (14-24 seconds), the nucleus was masked using DAPI signal, and colocalization (white) was displayed. For the final segment (25-41 seconds), a 3D surface rendering was created using the masked nucleus, and the spot function was used to identify areas of colocalization in white. The masked nucleus was rotated and scanned through using the orthogonal clipping plane tool.
