## Supplementary material for "Comparative analysis of retroviral Gag-host cell interactions: focus on the nuclear interactome": Table S1

| **Table S1.**  Top 10 DAVID biological processes for proteins isolated from the RSV Gag affinity purifications from DF1 nuclear lysates. | | |
| --- | --- | --- |
| **Gene Ontology Term (protein count)** | **Fold Enrichment**  **(p-value)** | **Gene names for proteins isolated** |
| GO:0006139  Nucleobase-containing compound metabolic process  (185) | 1.78  (7.66E-22) | XPO1, ASCC1, HIRA, MED24, MED22, INTS3, CNOT7, WTAP, CACTIN, SMNDC1, NONO, MAP3K7, RAD21, INTS5, MIER1, MED26, NT5C2, SMARCD1, U2AF1, ORC5, NUP37, OGT, RPRD1B, BCL7A, LUC7L3, TWIST2, MCM3AP, KRR1, MCRS1, CHTOP, GTPBP4, RRP7A, STRN3, MAGOH, PTBP1, MTA1, ACTN1, SLTM, CHRAC1, RFC5, AQR, AAAS, KDM2A, ASCC2, RFC1, DHX29, JUN, ZNF384, ATXN1L, YARS2, SCAF8, FUS, COX7B, SRSF11, AHCTF1, ZMYND8, RPS25, ZFC3H1, TCF20, HNRNPK, RPS29, ZNF326, ISY1, LEO1, IDH1, THAP12, RPL10A, TCF3, THAP11, HIP1, PRPF40A, TAF2, TBL3, TSR1, TAF8, TAF7, SMG1, SMAD2, MED13L, FOXP1, SAFB2, SRSF2, HNRNPH3, EIF4E, PLK1, PTPN1, HNRNPH1, SMC1A, ATP5D, ARID4A, FOXK2, PRKAG2, CTCF, ZEB1, PKM, SAP30, NDUFS4, RRP1B, VWA9, AEN, NDUFS8, ZNF407, RANBP2, LRWD1, ATP5H, NDUFS2, NDUFS1, CDK13, ZCCHC8, NOL6, CCNK, SSBP1, DDB1, ZFX, G3BP1, PRKCI, TLE3, EXOSC3, PRPF39, LEF1, NUP85, MBD3, EXOSC1, MCM3, RAD50, TTF2, NOC2L, MED6, EIF4A3, RECQL, TAF13, FANCD2, EIF4A2, SNRNP200, UCHL5, CNOT11, SNRNP40, JMJD1C, CUX1, THOC3, TOP3B, C1D, NDUFB3, ING2, ZBTB10, TRA2B, NDUFB9, UTP6, NOB1, HK2, HCFC2, POLR2B, CHD9, SUMO1, CHD7, PRPF8, SAFB, LANCL2, USP39, CAMK2D, MYCBP, ETV6, TRIP12, CHD3, NSA2, NDUFA4, NDUFA5, PDCD11, PDS5B, NDUFA9, NF1, PPP1R10, RPL27, PHF10, SAP30BP, SF3A3, PPP1R9B, ATXN2, MED30, SUGP1, RPL23, SP3, TCEB3, CALM3, IGFBP4 |
| GO:0034641  Cellular nitrogen compound metabolic process  (198) | 1.70  (1.19E-21) | ASCC1, HIRA, MED24, INTS3, MED22, WTAP, MAP3K7, NONO, INTS5, MED26, NT5C2, U2AF1, NUP37, OGT, BCL7A, TWIST2, LUC7L3, KRR1, MCRS1, CHTOP, GTPBP4, RRP7A, STRN3, PTBP1, RFC5, CHRAC1, AAAS, AQR, ASCC2, DHX29, RFC1, JUN, EIF2S2, ZNF384, ATXN1L, SCAF8, FUS, SRSF11, AHCTF1, IGF2BP3, RPS25, TCF20, RPS29, ISY1, LEO1, TMED10, IDH1, TCF3, HIP1, PRPF40A, SMAD2, SRSF2, ATP5D, ZEB1, PKM, VWA9, RRP1B, AEN, SLC25A3, ATP5H, CCNK, TLE3, EXOSC3, PRPF39, EXOSC1, MBD3, RAD50, TTF2, NOC2L, MED6, EIF4A3, TAF13, FANCD2, EIF4A2, CNOT11, SNRNP40, EIF5A2, THOC3, TOP3B, ZBTB10, NOB1, HK2, POLR2B, SUMO1, LANCL2, USP39, ETV6, TRIP12, NSA2, PDS5B, PPP1R10, PHF10, RPL27, SF3A3, SUGP1, RPL23, TCEB3, CALM3, XPO1, EIF5B, CNOT7, CACTIN, MTHFD1L, SMNDC1, RAD21, MIER1, SMARCD1, ORC5, RPRD1B, MCM3AP, MAGOH, MTA1, ACTN1, EEF2, SLTM, KDM2A, YARS2, COX7B, ZMYND8, RPL10L, ZFC3H1, HNRNPK, ZNF326, EIF3K, THAP12, RPL10A, THAP11, TAF2, TBL3, TSR1, TAF8, TAF7, SMG1, MED13L, FOXP1, SAFB2, HNRNPH3, EIF4E, PLK1, CIRBP, PTPN1, HNRNPH1, SMC1A, ARID4A, FOXK2, PRKAG2, CTCF, SAP30, NDUFS4, NDUFS8, ZNF407, RANBP2, LRWD1, NDUFS2, NDUFS1, CDK13, ZCCHC8, NOL6, SSBP1, DDB1, ZFX, G3BP1, PRKCI, LEF1, NUP85, MCM3, RECQL, EIF4G2, SNRNP200, UCHL5, JMJD1C, CUX1, C1D, NDUFB3, ING2, TRA2B, NDUFB9, UTP6, HCFC2, CHD9, CHD7, SAFB, PRPF8, CAMK2D, MYCBP, CHD3, NDUFA4, NDUFA5, PDCD11, NDUFA9, NF1, SAP30BP, PPP1R9B, ATXN2, MED30, SP3, IGFBP4, MGST1 |
| GO:0046483  Heterocycle metabolic process  (186) | 1.75  (3.65E-21) | XPO1, ASCC1, HIRA, MED24, MED22, INTS3, CNOT7, WTAP, CACTIN, MTHFD1L, SMNDC1, NONO, MAP3K7, RAD21, INTS5, MIER1, MED26, NT5C2, SMARCD1, U2AF1, ORC5, NUP37, OGT, RPRD1B, BCL7A, LUC7L3, TWIST2, MCM3AP, KRR1, MCRS1, CHTOP, GTPBP4, RRP7A, STRN3, MAGOH, PTBP1, MTA1, ACTN1, SLTM, CHRAC1, RFC5, AQR, AAAS, KDM2A, ASCC2, RFC1, DHX29, JUN, ZNF384, ATXN1L, YARS2, SCAF8, FUS, COX7B, SRSF11, AHCTF1, ZMYND8, RPS25, ZFC3H1, TCF20, HNRNPK, RPS29, ZNF326, ISY1, LEO1, IDH1, THAP12, RPL10A, TCF3, THAP11, HIP1, PRPF40A, TAF2, TBL3, TSR1, TAF8, TAF7, SMG1, SMAD2, MED13L, FOXP1, SAFB2, SRSF2, HNRNPH3, EIF4E, PLK1, PTPN1, HNRNPH1, SMC1A, ATP5D, ARID4A, FOXK2, PRKAG2, CTCF, ZEB1, PKM, SAP30, NDUFS4, RRP1B, VWA9, AEN, NDUFS8, ZNF407, RANBP2, LRWD1, ATP5H, NDUFS2, NDUFS1, CDK13, ZCCHC8, NOL6, CCNK, SSBP1, DDB1, ZFX, G3BP1, PRKCI, TLE3, EXOSC3, PRPF39, LEF1, NUP85, MBD3, EXOSC1, MCM3, RAD50, TTF2, NOC2L, MED6, EIF4A3, RECQL, TAF13, FANCD2, EIF4A2, SNRNP200, UCHL5, CNOT11, SNRNP40, JMJD1C, CUX1, THOC3, TOP3B, C1D, NDUFB3, ING2, ZBTB10, TRA2B, NDUFB9, UTP6, NOB1, HK2, HCFC2, POLR2B, CHD9, SUMO1, CHD7, PRPF8, SAFB, LANCL2, USP39, CAMK2D, MYCBP, ETV6, TRIP12, CHD3, NSA2, NDUFA4, NDUFA5, PDCD11, PDS5B, NDUFA9, NF1, PPP1R10, RPL27, PHF10, SAP30BP, SF3A3, PPP1R9B, ATXN2, MED30, SUGP1, RPL23, SP3, TCEB3, CALM3, IGFBP4 |
| GO:1901360  Organic cyclic compound metabolic process  (190) | 1.72  (5.48E-21) | XPO1, ASCC1, HIRA, MED24, MED22, INTS3, CNOT7, WTAP, CACTIN, MTHFD1L, SMNDC1, NONO, MAP3K7, RAD21, INTS5, MIER1, MED26, NT5C2, SMARCD1, U2AF1, ORC5, NUP37, OGT, RPRD1B, BCL7A, LUC7L3, TWIST2, MCM3AP, KRR1, MCRS1, CHTOP, GTPBP4, RRP7A, STRN3, MAGOH, PTBP1, MTA1, ACTN1, SLTM, CHRAC1, RFC5, AQR, AAAS, KDM2A, ASCC2, RFC1, DHX29, JUN, ZNF384, ATXN1L, YARS2, SCAF8, FUS, HDLBP, COX7B, SRSF11, AHCTF1, ZMYND8, RPS25, ZFC3H1, TCF20, HNRNPK, RPS29, ZNF326, ISY1, LEO1, IDH1, THAP12, RPL10A, TCF3, THAP11, HIP1, PRPF40A, TAF2, TBL3, MSMO1, TSR1, TAF8, TAF7, FDPS, SMG1, SMAD2, MED13L, FOXP1, SAFB2, SRSF2, HNRNPH3, EIF4E, PLK1, PTPN1, HNRNPH1, SMC1A, ATP5D, ARID4A, FOXK2, PRKAG2, CTCF, ZEB1, PKM, SAP30, NDUFS4, RRP1B, VWA9, AEN, NDUFS8, ZNF407, RANBP2, LRWD1, ATP5H, NDUFS2, NDUFS1, CDK13, ZCCHC8, NOL6, CCNK, SSBP1, DDB1, ZFX, G3BP1, PRKCI, TLE3, EXOSC3, PRPF39, LEF1, NUP85, MBD3, EXOSC1, MCM3, RAD50, TTF2, NOC2L, MED6, EIF4A3, RECQL, TAF13, FANCD2, EIF4A2, SNRNP200, UCHL5, CNOT11, SNRNP40, JMJD1C, CUX1, THOC3, TOP3B, C1D, NDUFB3, ING2, ZBTB10, TRA2B, NDUFB9, UTP6, NOB1, HK2, HCFC2, POLR2B, CHD9, SUMO1, CHD7, PRPF8, SAFB, LANCL2, USP39, CAMK2D, MYCBP, ETV6, TRIP12, CHD3, NSA2, NDUFA4, NDUFA5, PDCD11, PDS5B, NDUFA9, NF1, PPP1R10, RPL27, PHF10, ATP1A1, SAP30BP, SF3A3, PPP1R9B, ATXN2, MED30, SUGP1, RPL23, SP3, TCEB3, CALM3, IGFBP4 |
| GO:0006725  Cellular aromatic compound metabolic process  (186) | 1.74  (9.57E-210 | XPO1, ASCC1, HIRA, MED24, MED22, INTS3, CNOT7, WTAP, CACTIN, MTHFD1L, SMNDC1, NONO, MAP3K7, RAD21, INTS5, MIER1, MED26, NT5C2, SMARCD1, U2AF1, ORC5, NUP37, OGT, RPRD1B, BCL7A, LUC7L3, TWIST2, MCM3AP, KRR1, MCRS1, CHTOP, GTPBP4, RRP7A, STRN3, MAGOH, PTBP1, MTA1, ACTN1, SLTM, CHRAC1, RFC5, AQR, AAAS, KDM2A, ASCC2, RFC1, DHX29, JUN, ZNF384, ATXN1L, YARS2, SCAF8, FUS, COX7B, SRSF11, AHCTF1, ZMYND8, RPS25, ZFC3H1, TCF20, HNRNPK, RPS29, ZNF326, ISY1, LEO1, IDH1, THAP12, RPL10A, TCF3, THAP11, HIP1, PRPF40A, TAF2, TBL3, TSR1, TAF8, TAF7, SMG1, SMAD2, MED13L, FOXP1, SAFB2, SRSF2, HNRNPH3, EIF4E, PLK1, PTPN1, HNRNPH1, SMC1A, ATP5D, ARID4A, FOXK2, PRKAG2, CTCF, ZEB1, PKM, SAP30, NDUFS4, RRP1B, VWA9, AEN, NDUFS8, ZNF407, RANBP2, LRWD1, ATP5H, NDUFS2, NDUFS1, CDK13, ZCCHC8, NOL6, CCNK, SSBP1, DDB1, ZFX, G3BP1, PRKCI, TLE3, EXOSC3, PRPF39, LEF1, NUP85, MBD3, EXOSC1, MCM3, RAD50, TTF2, NOC2L, MED6, EIF4A3, RECQL, TAF13, FANCD2, EIF4A2, SNRNP200, UCHL5, CNOT11, SNRNP40, JMJD1C, CUX1, THOC3, TOP3B, C1D, NDUFB3, ING2, ZBTB10, TRA2B, NDUFB9, UTP6, NOB1, HK2, HCFC2, POLR2B, CHD9, SUMO1, CHD7, PRPF8, SAFB, LANCL2, USP39, CAMK2D, MYCBP, ETV6, TRIP12, CHD3, NSA2, NDUFA4, NDUFA5, PDCD11, PDS5B, NDUFA9, NF1, PPP1R10, RPL27, PHF10, SAP30BP, SF3A3, PPP1R9B, ATXN2, MED30, SUGP1, RPL23, SP3, TCEB3, CALM3, IGFBP4 |
| GO:0006807  Nitrogen compound metabolic process  (201) | 1.61  (2.95E-19) | ASCC1, HIRA, MED24, INTS3, MED22, WTAP, NONO, MAP3K7, INTS5, MED26, NT5C2, U2AF1, NUP37, OGT, BCL7A, TWIST2, LUC7L3, KRR1, MCRS1, CHTOP, GTPBP4, RRP7A, STRN3, CRTAP, PTBP1, RFC5, CHRAC1, AQR, AAAS, ASCC2, DHX29, RFC1, JUN, EIF2S2, ZNF384, ATXN1L, SCAF8, FUS, SRSF11, AHCTF1, IGF2BP3, RPS25, TCF20, RPS29, ISY1, LEO1, TMED10, IDH1, TCF3, HIP1, PRPF40A, SMAD2, SRSF2, ATP5D, ZEB1, PKM, VWA9, RRP1B, P4HA1, AEN, SLC25A3, ATP5H, CCNK, TLE3, EXOSC3, PRPF39, EXOSC1, MBD3, RAD50, TTF2, NOC2L, MED6, EIF4A3, TAF13, FANCD2, EIF4A2, CNOT11, SNRNP40, EIF5A2, THOC3, TOP3B, BCAT1, ZBTB10, NOB1, HK2, POLR2B, SUMO1, LANCL2, USP39, ETV6, TRIP12, NSA2, PDS5B, PPP1R10, PHF10, RPL27, SF3A3, SUGP1, RPL23, TCEB3, CALM3, XPO1, EIF5B, CNOT7, CACTIN, MTHFD1L, SMNDC1, RAD21, MIER1, SMARCD1, ORC5, RPRD1B, MCM3AP, MAGOH, MTA1, ACTN1, EEF2, SLTM, KDM2A, YARS2, COX7B, ZMYND8, RPL10L, ZFC3H1, HNRNPK, ZNF326, EIF3K, THAP12, RPL10A, THAP11, TAF2, TBL3, TSR1, TAF8, TAF7, SMG1, MED13L, FOXP1, SAFB2, HNRNPH3, EIF4E, PLK1, CIRBP, PTPN1, HNRNPH1, SMC1A, ARID4A, FOXK2, PRKAG2, CTCF, SAP30, NDUFS4, NDUFS8, ZNF407, RANBP2, LRWD1, NDUFS2, NDUFS1, CDK13, ZCCHC8, NOL6, SSBP1, DDB1, ZFX, G3BP1, PRKCI, LEF1, NUP85, MCM3, RECQL, EIF4G2, SNRNP200, UCHL5, JMJD1C, CUX1, C1D, NDUFB3, ING2, TRA2B, NDUFB9, UTP6, HCFC2, CHD9, CHD7, SAFB, PRPF8, CAMK2D, MYCBP, CHD3, NDUFA4, NDUFA5, PDCD11, NDUFA9, NF1, SAP30BP, PPP1R9B, ATXN2, MED30, SP3, IGFBP4, MGST1 |
| GO:0006396  RNA processing  (61) | 3.74  (5.01E-19) | INTS3, WTAP, CACTIN, SMNDC1, NONO, RRP1B, INTS5, U2AF1, LUC7L3, CDK13, ZCCHC8, NOL6, KRR1, GTPBP4, CHTOP, RRP7A, MAGOH, PTBP1, PRPF39, EXOSC3, EXOSC1, TTF2, EIF4A3, AQR, DHX29, SNRNP200, SNRNP40, SCAF8, THOC3, C1D, FUS, TRA2B, UTP6, SRSF11, NOB1, POLR2B, ZFC3H1, RPS25, CHD7, HNRNPK, RPS29, ZNF326, PRPF8, USP39, ISY1, LEO1, RPL10A, NSA2, PRPF40A, TBL3, PDCD11, TSR1, RPL27, SMAD2, SF3A3, SRSF2, PPP1R9B, HNRNPH3, SUGP1, RPL23, HNRNPH1 |
| GO:0090304  Nucleic acid metabolic process  (165) | 1.78  (2.22E-18) | XPO1, ASCC1, HIRA, MED24, MED22, INTS3, CNOT7, WTAP, CACTIN, SMNDC1, NONO, MAP3K7, RAD21, INTS5, MIER1, MED26, SMARCD1, U2AF1, ORC5, NUP37, OGT, RPRD1B, BCL7A, LUC7L3, TWIST2, MCM3AP, KRR1, MCRS1, CHTOP, GTPBP4, RRP7A, STRN3, MAGOH, PTBP1, MTA1, ACTN1, SLTM, CHRAC1, RFC5, AQR, AAAS, KDM2A, ASCC2, RFC1, DHX29, JUN, ZNF384, ATXN1L, YARS2, SCAF8, FUS, SRSF11, AHCTF1, ZMYND8, RPS25, ZFC3H1, TCF20, HNRNPK, RPS29, ZNF326, ISY1, LEO1, THAP12, RPL10A, TCF3, THAP11, HIP1, PRPF40A, TAF2, TBL3, TSR1, TAF8, TAF7, SMG1, SMAD2, MED13L, FOXP1, SAFB2, SRSF2, HNRNPH3, EIF4E, PLK1, HNRNPH1, SMC1A, ARID4A, FOXK2, CTCF, ZEB1, SAP30, RRP1B, VWA9, AEN, ZNF407, RANBP2, LRWD1, CDK13, ZCCHC8, NOL6, CCNK, SSBP1, DDB1, ZFX, G3BP1, PRKCI, TLE3, EXOSC3, PRPF39, LEF1, NUP85, MBD3, EXOSC1, MCM3, RAD50, TTF2, NOC2L, MED6, EIF4A3, RECQL, TAF13, FANCD2, EIF4A2, SNRNP200, UCHL5, CNOT11, SNRNP40, JMJD1C, CUX1, THOC3, C1D, TOP3B, ING2, ZBTB10, TRA2B, UTP6, NOB1, HCFC2, POLR2B, CHD9, SUMO1, CHD7, PRPF8, SAFB, LANCL2, USP39, CAMK2D, MYCBP, ETV6, TRIP12, CHD3, NSA2, PDCD11, PDS5B, PPP1R10, RPL27, PHF10, SAP30BP, SF3A3, PPP1R9B, ATXN2, MED30, SUGP1, RPL23, SP3, TCEB3, IGFBP4 |
| GO:0016071  mRNA metabolic process  (51) | 4.23  (4.83E-18) | CNOT7, WTAP, CACTIN, SMNDC1, NONO, U2AF1, LUC7L3, CDK13, ZCCHC8, CHTOP, MAGOH, PTBP1, EXOSC3, PRPF39, EXOSC1, TTF2, SLTM, EIF4A3, AQR, EIF4A2, SNRNP200, CNOT11, SNRNP40, SCAF8, THOC3, FUS, TRA2B, SRSF11, POLR2B, RPS25, HNRNPK, RPS29, ZNF326, SAFB, PRPF8, USP39, ISY1, LEO1, RPL10A, PRPF40A, PDCD11, SMG1, RPL27, SF3A3, SAFB2, SRSF2, HNRNPH3, SUGP1, EIF4E, RPL23, HNRNPH1 |
| GO:0010467  Gene expression  (163) | 1.71  (5.18E-16) | XPO1, EIF5B, ASCC1, HIRA, MED24, MED22, INTS3, CNOT7, WTAP, CACTIN, SMNDC1, NONO, MAP3K7, RAD21, INTS5, MIER1, MED26, SMARCD1, U2AF1, NUP37, OGT, RPRD1B, BCL7A, LUC7L3, TWIST2, KRR1, MCRS1, CHTOP, GTPBP4, RRP7A, STRN3, MAGOH, PTBP1, MTA1, EEF2, PPP1CC, MYH9, PPP1CB, SLTM, AQR, AAAS, KDM2A, ASCC2, RFC1, DHX29, SERBP1, JUN, EIF2S2, ZNF384, ATXN1L, YARS2, SCAF8, GLG1, FUS, SRSF11, AHCTF1, IGF2BP3, ZMYND8, RPS25, RPL10L, ZFC3H1, TCF20, HNRNPK, RPS29, ZNF326, ISY1, EIF3K, LEO1, THAP12, RPL10A, THAP11, TCF3, HIP1, PRPF40A, TAF2, TBL3, TSR1, TAF8, TAF7, SMG1, SMAD2, MED13L, FOXP1, SAFB2, SRSF2, HNRNPH3, EIF4E, PLK1, CIRBP, HNRNPH1, ARID4A, FOXK2, CTCF, ZEB1, SAP30, RRP1B, VWA9, SLC25A3, ZNF407, RANBP2, CDK13, ZCCHC8, NOL6, CCNK, ZFX, PRKCI, TLE3, EXOSC3, PRPF39, LEF1, NUP85, MBD3, EXOSC1, TTF2, NOC2L, MED6, EIF4G2, EIF4A3, TAF13, FANCD2, EIF4A2, SNRNP200, UCHL5, CNOT11, SNRNP40, JMJD1C, CUX1, EIF5A2, THOC3, C1D, ING2, ZBTB10, TRA2B, UTP6, NOB1, HCFC2, POLR2B, CHD9, SUMO1, CHD7, PRPF8, SAFB, LANCL2, USP39, CAMK2D, CNN2, MYCBP, ETV6, CHD3, NSA2, PDCD11, PPP1R10, RPL27, PHF10, SAP30BP, SF3A3, PPP1R9B, ATXN2, MED30, SUGP1, RPL23, SP3, TCEB3 |
