## Supplementary material for "Comparative analysis of retroviral Gag-host cell interactions: focus on the nuclear interactome": Table S2

| **Table S2.**  Top 10 nuclear enriched DAVID biological processes for proteins isolated from the RSV Gag affinity purifications from DF1 nuclear lysates. | | |
| --- | --- | --- |
| **Gene Ontology Term (protein count)** | **Fold Enrichment (p-value)** | **Gene names for proteins isolated** |
| GO:0090304  Nucleic acid metabolic process  (158) | 2.36  (6.38E-37) | XPO1, ASCC1, HIRA, MED24, MED22, INTS3, CNOT7, WTAP, CACTIN, SMNDC1, NONO, MAP3K7, RAD21, INTS5, MIER1, MED26, SMARCD1, U2AF1, ORC5, NUP37, OGT, RPRD1B, LUC7L3, TWIST2, MCM3AP, KRR1, MCRS1, CHTOP, GTPBP4, RRP7A, STRN3, MAGOH, PTBP1, MTA1, SLTM, CHRAC1, RFC5, AQR, AAAS, KDM2A, ASCC2, RFC1, DHX29, JUN, ZNF384, ATXN1L, SCAF8, FUS, SRSF11, AHCTF1, ZMYND8, RPS25, TCF20, HNRNPK, RPS29, ZNF326, ISY1, LEO1, THAP12, RPL10A, THAP11, TCF3, HIP1, PRPF40A, TAF2, TBL3, TSR1, TAF8, TAF7, SMG1, SMAD2, MED13L, FOXP1, SAFB2, SRSF2, HNRNPH3, PLK1, HNRNPH1, SMC1A, ARID4A, FOXK2, CTCF, ZEB1, SAP30, RRP1B, VWA9, AEN, ZNF407, RANBP2, LRWD1, CDK13, ZCCHC8, NOL6, CCNK, SSBP1, DDB1, ZFX, G3BP1, PRKCI, TLE3, EXOSC3, PRPF39, LEF1, NUP85, MBD3, EXOSC1, MCM3, RAD50, TTF2, NOC2L, MED6, EIF4A3, RECQL, TAF13, FANCD2, SNRNP200, UCHL5, CNOT11, SNRNP40, JMJD1C, CUX1, THOC3, C1D, TOP3B, ING2, ZBTB10, TRA2B, UTP6, NOB1, HCFC2, POLR2B, CHD9, SUMO1, CHD7, PRPF8, SAFB, LANCL2, USP39, CAMK2D, MYCBP, ETV6, TRIP12, CHD3, NSA2, PDCD11, PDS5B, PPP1R10, RPL27, PHF10, SAP30BP, SF3A3, PPP1R9B, ATXN2, MED30, SUGP1, RPL23, SP3, TCEB3 |
| GO:0034641  Cellular nitrogen compound metabolic process  (175) | 2.07  (3.10E-36) | XPO1, EIF5B, ASCC1, HIRA, MED24, MED22, INTS3, CNOT7, WTAP, CACTIN, SMNDC1, NONO, MAP3K7, RAD21, INTS5, MIER1, MED26, SMARCD1, U2AF1, ORC5, NUP37, OGT, RPRD1B, LUC7L3, TWIST2, MCM3AP, KRR1, MCRS1, CHTOP, GTPBP4, RRP7A, STRN3, MAGOH, PTBP1, MTA1, EEF2, SLTM, CHRAC1, RFC5, AQR, AAAS, KDM2A, ASCC2, RFC1, DHX29, JUN, EIF2S2, ZNF384, ATXN1L, SCAF8, FUS, SRSF11, AHCTF1, IGF2BP3, ZMYND8, RPS25, RPL10L, TCF20, HNRNPK, RPS29, ZNF326, ISY1, EIF3K, LEO1, THAP12, RPL10A, TCF3, THAP11, HIP1, PRPF40A, TAF2, TBL3, TSR1, TAF8, TAF7, SMG1, SMAD2, MED13L, FOXP1, SAFB2, SRSF2, HNRNPH3, PLK1, CIRBP, HNRNPH1, SMC1A, ARID4A, FOXK2, PRKAG2, CTCF, ZEB1, PKM, SAP30, RRP1B, VWA9, AEN, SLC25A3, ZNF407, RANBP2, LRWD1, ATP5H, NDUFS2, CDK13, ZCCHC8, NOL6, CCNK, SSBP1, DDB1, ZFX, G3BP1, PRKCI, TLE3, EXOSC3, PRPF39, LEF1, NUP85, MBD3, EXOSC1, MCM3, RAD50, TTF2, NOC2L, MED6, EIF4A3, RECQL, TAF13, FANCD2, SNRNP200, UCHL5, CNOT11, SNRNP40, JMJD1C, CUX1, EIF5A2, THOC3, C1D, TOP3B, ING2, ZBTB10, TRA2B, UTP6, NOB1, HCFC2, POLR2B, CHD9, SUMO1, CHD7, PRPF8, SAFB, LANCL2, USP39, CAMK2D, MYCBP, ETV6, TRIP12, CHD3, NSA2, PDCD11, PDS5B, NDUFA9, NF1, PPP1R10, RPL27, PHF10, SAP30BP, SF3A3, PPP1R9B, ATXN2, MED30, SUGP1, RPL23, SP3, TCEB3, CALM3, MGST1 |
| GO:0006139  Nucleobase-containing compound metabolic process  (165) | 2.19  (1.60E-35) | XPO1, ASCC1, HIRA, MED24, MED22, INTS3, CNOT7, WTAP, CACTIN, SMNDC1, NONO, MAP3K7, RAD21, INTS5, MIER1, MED26, SMARCD1, U2AF1, ORC5, NUP37, OGT, RPRD1B, LUC7L3, TWIST2, MCM3AP, KRR1, MCRS1, CHTOP, GTPBP4, RRP7A, STRN3, MAGOH, PTBP1, MTA1, SLTM, CHRAC1, RFC5, AQR, AAAS, KDM2A, ASCC2, RFC1, DHX29, JUN, ZNF384, ATXN1L, SCAF8, FUS, SRSF11, AHCTF1, ZMYND8, RPS25, TCF20, HNRNPK, RPS29, ZNF326, ISY1, LEO1, THAP12, RPL10A, TCF3, THAP11, HIP1, PRPF40A, TAF2, TBL3, TSR1, TAF8, TAF7, SMG1, SMAD2, MED13L, FOXP1, SAFB2, SRSF2, HNRNPH3, PLK1, HNRNPH1, SMC1A, ARID4A, FOXK2, PRKAG2, CTCF, ZEB1, PKM, SAP30, RRP1B, VWA9, AEN, ZNF407, RANBP2, LRWD1, ATP5H, NDUFS2, CDK13, ZCCHC8, NOL6, CCNK, SSBP1, DDB1, ZFX, G3BP1, PRKCI, TLE3, EXOSC3, PRPF39, LEF1, NUP85, MBD3, EXOSC1, MCM3, RAD50, TTF2, NOC2L, MED6, EIF4A3, RECQL, TAF13, FANCD2, SNRNP200, UCHL5, CNOT11, SNRNP40, JMJD1C, CUX1, THOC3, C1D, TOP3B, ING2, ZBTB10, TRA2B, UTP6, NOB1, HCFC2, POLR2B, CHD9, SUMO1, CHD7, PRPF8, SAFB, LANCL2, USP39, CAMK2D, MYCBP, ETV6, TRIP12, CHD3, NSA2, PDCD11, PDS5B, NDUFA9, NF1, PPP1R10, RPL27, PHF10, SAP30BP, SF3A3, PPP1R9B, ATXN2, MED30, SUGP1, RPL23, SP3, CALM3, TCEB3 |
| GO:0046483  Heterocycle metabolic process  (165) | 2.15  (3.03E-34) | XPO1, ASCC1, HIRA, MED24, MED22, INTS3, CNOT7, WTAP, CACTIN, SMNDC1, NONO, MAP3K7, RAD21, INTS5, MIER1, MED26, SMARCD1, U2AF1, ORC5, NUP37, OGT, RPRD1B, LUC7L3, TWIST2, MCM3AP, KRR1, MCRS1, CHTOP, GTPBP4, RRP7A, STRN3, MAGOH, PTBP1, MTA1, SLTM, CHRAC1, RFC5, AQR, AAAS, KDM2A, ASCC2, RFC1, DHX29, JUN, ZNF384, ATXN1L, SCAF8, FUS, SRSF11, AHCTF1, ZMYND8, RPS25, TCF20, HNRNPK, RPS29, ZNF326, ISY1, LEO1, THAP12, RPL10A, TCF3, THAP11, HIP1, PRPF40A, TAF2, TBL3, TSR1, TAF8, TAF7, SMG1, SMAD2, MED13L, FOXP1, SAFB2, SRSF2, HNRNPH3, PLK1, HNRNPH1, SMC1A, ARID4A, FOXK2, PRKAG2, CTCF, ZEB1, PKM, SAP30, RRP1B, VWA9, AEN, ZNF407, RANBP2, LRWD1, ATP5H, NDUFS2, CDK13, ZCCHC8, NOL6, CCNK, SSBP1, DDB1, ZFX, G3BP1, PRKCI, TLE3, EXOSC3, PRPF39, LEF1, NUP85, MBD3, EXOSC1, MCM3, RAD50, TTF2, NOC2L, MED6, EIF4A3, RECQL, TAF13, FANCD2, SNRNP200, UCHL5, CNOT11, SNRNP40, JMJD1C, CUX1, THOC3, C1D, TOP3B, ING2, ZBTB10, TRA2B, UTP6, NOB1, HCFC2, POLR2B, CHD9, SUMO1, CHD7, PRPF8, SAFB, LANCL2, USP39, CAMK2D, MYCBP, ETV6, TRIP12, CHD3, NSA2, PDCD11, PDS5B, NDUFA9, NF1, PPP1R10, RPL27, PHF10, SAP30BP, SF3A3, PPP1R9B, ATXN2, MED30, SUGP1, RPL23, SP3, CALM3, TCEB3 |
| GO:0006725  Cellular aromatic compound metabolic process  (165) | 2.13  (8.76E-34) | XPO1, ASCC1, HIRA, MED24, MED22, INTS3, CNOT7, WTAP, CACTIN, SMNDC1, NONO, MAP3K7, RAD21, INTS5, MIER1, MED26, SMARCD1, U2AF1, ORC5, NUP37, OGT, RPRD1B, LUC7L3, TWIST2, MCM3AP, KRR1, MCRS1, CHTOP, GTPBP4, RRP7A, STRN3, MAGOH, PTBP1, MTA1, SLTM, CHRAC1, RFC5, AQR, AAAS, KDM2A, ASCC2, RFC1, DHX29, JUN, ZNF384, ATXN1L, SCAF8, FUS, SRSF11, AHCTF1, ZMYND8, RPS25, TCF20, HNRNPK, RPS29, ZNF326, ISY1, LEO1, THAP12, RPL10A, TCF3, THAP11, HIP1, PRPF40A, TAF2, TBL3, TSR1, TAF8, TAF7, SMG1, SMAD2, MED13L, FOXP1, SAFB2, SRSF2, HNRNPH3, PLK1, HNRNPH1, SMC1A, ARID4A, FOXK2, PRKAG2, CTCF, ZEB1, PKM, SAP30, RRP1B, VWA9, AEN, ZNF407, RANBP2, LRWD1, ATP5H, NDUFS2, CDK13, ZCCHC8, NOL6, CCNK, SSBP1, DDB1, ZFX, G3BP1, PRKCI, TLE3, EXOSC3, PRPF39, LEF1, NUP85, MBD3, EXOSC1, MCM3, RAD50, TTF2, NOC2L, MED6, EIF4A3, RECQL, TAF13, FANCD2, SNRNP200, UCHL5, CNOT11, SNRNP40, JMJD1C, CUX1, THOC3, C1D, TOP3B, ING2, ZBTB10, TRA2B, UTP6, NOB1, HCFC2, POLR2B, CHD9, SUMO1, CHD7, PRPF8, SAFB, LANCL2, USP39, CAMK2D, MYCBP, ETV6, TRIP12, CHD3, NSA2, PDCD11, PDS5B, NDUFA9, NF1, PPP1R10, RPL27, PHF10, SAP30BP, SF3A3, PPP1R9B, ATXN2, MED30, SUGP1, RPL23, SP3, CALM3, TCEB3 |
| GO:1901360  Organic cyclic compound metabolic process  (167) | 2.09  (1.77E-33) | XPO1, ASCC1, HIRA, MED24, MED22, INTS3, CNOT7, WTAP, CACTIN, SMNDC1, NONO, MAP3K7, RAD21, INTS5, MIER1, MED26, SMARCD1, U2AF1, ORC5, NUP37, OGT, RPRD1B, LUC7L3, TWIST2, MCM3AP, KRR1, MCRS1, CHTOP, GTPBP4, RRP7A, STRN3, MAGOH, PTBP1, MTA1, SLTM, CHRAC1, RFC5, AQR, AAAS, KDM2A, ASCC2, RFC1, DHX29, JUN, ZNF384, ATXN1L, SCAF8, FUS, HDLBP, SRSF11, AHCTF1, ZMYND8, RPS25, TCF20, HNRNPK, RPS29, ZNF326, ISY1, LEO1, THAP12, RPL10A, TCF3, THAP11, HIP1, PRPF40A, TAF2, TBL3, TSR1, TAF8, TAF7, FDPS, SMG1, SMAD2, MED13L, FOXP1, SAFB2, SRSF2, HNRNPH3, PLK1, HNRNPH1, SMC1A, ARID4A, FOXK2, PRKAG2, CTCF, ZEB1, PKM, SAP30, RRP1B, VWA9, AEN, ZNF407, RANBP2, LRWD1, ATP5H, NDUFS2, CDK13, ZCCHC8, NOL6, CCNK, SSBP1, DDB1, ZFX, G3BP1, PRKCI, TLE3, EXOSC3, PRPF39, LEF1, NUP85, MBD3, EXOSC1, MCM3, RAD50, TTF2, NOC2L, MED6, EIF4A3, RECQL, TAF13, FANCD2, SNRNP200, UCHL5, CNOT11, SNRNP40, JMJD1C, CUX1, THOC3, C1D, TOP3B, ING2, ZBTB10, TRA2B, UTP6, NOB1, HCFC2, POLR2B, CHD9, SUMO1, CHD7, PRPF8, SAFB, LANCL2, USP39, CAMK2D, MYCBP, ETV6, TRIP12, CHD3, NSA2, PDCD11, PDS5B, NDUFA9, NF1, PPP1R10, RPL27, PHF10, SAP30BP, SF3A3, PPP1R9B, ATXN2, MED30, SUGP1, RPL23, SP3, CALM3, TCEB3 |
| GO:0010467  Gene expression  (155) | 2.25  (4.66E-33) | XPO1, EIF5B, ASCC1, HIRA, MED24, MED22, INTS3, CNOT7, WTAP, CACTIN, SMNDC1, NONO, MAP3K7, RAD21, INTS5, MIER1, MED26, SMARCD1, U2AF1, NUP37, OGT, RPRD1B, LUC7L3, TWIST2, KRR1, MCRS1, CHTOP, GTPBP4, RRP7A, STRN3, MAGOH, PTBP1, MTA1, EEF2, PPP1CC, MYH9, PPP1CB, SLTM, AQR, AAAS, KDM2A, ASCC2, RFC1, DHX29, SERBP1, JUN, EIF2S2, ZNF384, ATXN1L, SCAF8, FUS, SRSF11, AHCTF1, IGF2BP3, ZMYND8, RPS25, RPL10L, TCF20, HNRNPK, RPS29, ZNF326, ISY1, EIF3K, LEO1, THAP12, RPL10A, THAP11, TCF3, HIP1, PRPF40A, TAF2, TBL3, TSR1, TAF8, TAF7, SMG1, SMAD2, MED13L, FOXP1, SAFB2, SRSF2, HNRNPH3, PLK1, CIRBP, HNRNPH1, ARID4A, FOXK2, CTCF, ZEB1, SAP30, RRP1B, VWA9, SLC25A3, ZNF407, RANBP2, CDK13, ZCCHC8, NOL6, CCNK, ZFX, PRKCI, TLE3, EXOSC3, PRPF39, LEF1, NUP85, MBD3, EXOSC1, TTF2, NOC2L, MED6, EIF4A3, TAF13, FANCD2, SNRNP200, UCHL5, CNOT11, SNRNP40, JMJD1C, CUX1, EIF5A2, THOC3, C1D, ING2, ZBTB10, TRA2B, UTP6, NOB1, HCFC2, POLR2B, CHD9, SUMO1, CHD7, SAFB, LANCL2, PRPF8, USP39, CAMK2D, MYCBP, ETV6, CHD3, NSA2, PDCD11, PPP1R10, RPL27, PHF10, SAP30BP, SF3A3, PPP1R9B, ATXN2, MED30, SUGP1, RPL23, SP3, TCEB3 |
| GO:0006807  Nitrogen compound metabolic process  (175) | 1.94  (5.14E-32) | XPO1, EIF5B, ASCC1, HIRA, MED24, MED22, INTS3, CNOT7, WTAP, CACTIN, SMNDC1, NONO, MAP3K7, RAD21, INTS5, MIER1, MED26, SMARCD1, U2AF1, ORC5, NUP37, OGT, RPRD1B, LUC7L3, TWIST2, MCM3AP, KRR1, MCRS1, CHTOP, GTPBP4, RRP7A, STRN3, MAGOH, PTBP1, MTA1, EEF2, SLTM, CHRAC1, RFC5, AQR, AAAS, KDM2A, ASCC2, RFC1, DHX29, JUN, EIF2S2, ZNF384, ATXN1L, SCAF8, FUS, SRSF11, AHCTF1, IGF2BP3, ZMYND8, RPS25, RPL10L, TCF20, HNRNPK, RPS29, ZNF326, ISY1, EIF3K, LEO1, THAP12, RPL10A, TCF3, THAP11, HIP1, PRPF40A, TAF2, TBL3, TSR1, TAF8, TAF7, SMG1, SMAD2, MED13L, FOXP1, SAFB2, SRSF2, HNRNPH3, PLK1, CIRBP, HNRNPH1, SMC1A, ARID4A, FOXK2, PRKAG2, CTCF, ZEB1, PKM, SAP30, RRP1B, VWA9, AEN, SLC25A3, ZNF407, RANBP2, LRWD1, ATP5H, NDUFS2, CDK13, ZCCHC8, NOL6, CCNK, SSBP1, DDB1, ZFX, G3BP1, PRKCI, TLE3, EXOSC3, PRPF39, LEF1, NUP85, MBD3, EXOSC1, MCM3, RAD50, TTF2, NOC2L, MED6, EIF4A3, RECQL, TAF13, FANCD2, SNRNP200, UCHL5, CNOT11, SNRNP40, JMJD1C, CUX1, EIF5A2, THOC3, C1D, TOP3B, ING2, ZBTB10, TRA2B, UTP6, NOB1, HCFC2, POLR2B, CHD9, SUMO1, CHD7, PRPF8, SAFB, LANCL2, USP39, CAMK2D, MYCBP, ETV6, TRIP12, CHD3, NSA2, PDCD11, PDS5B, NDUFA9, NF1, PPP1R10, RPL27, PHF10, SAP30BP, SF3A3, PPP1R9B, ATXN2, MED30, SUGP1, RPL23, SP3, TCEB3, CALM3, MGST1 |
| GO:0016070  RNA metabolic process  (142) | 2.35  (1.02E-30) | XPO1, ASCC1, HIRA, MED24, MED22, INTS3, CNOT7, WTAP, CACTIN, SMNDC1, NONO, MAP3K7, RAD21, INTS5, MIER1, MED26, SMARCD1, U2AF1, NUP37, OGT, RPRD1B, LUC7L3, TWIST2, KRR1, MCRS1, CHTOP, GTPBP4, RRP7A, STRN3, MAGOH, PTBP1, MTA1, SLTM, AQR, AAAS, KDM2A, ASCC2, RFC1, DHX29, JUN, ZNF384, ATXN1L, SCAF8, FUS, SRSF11, AHCTF1, ZMYND8, RPS25, TCF20, HNRNPK, RPS29, ZNF326, ISY1, LEO1, THAP12, RPL10A, THAP11, TCF3, HIP1, PRPF40A, TAF2, TBL3, TSR1, TAF8, TAF7, SMG1, SMAD2, MED13L, FOXP1, SAFB2, SRSF2, HNRNPH3, PLK1, HNRNPH1, ARID4A, FOXK2, CTCF, ZEB1, SAP30, RRP1B, VWA9, ZNF407, RANBP2, CDK13, ZCCHC8, NOL6, CCNK, ZFX, TLE3, PRKCI, EXOSC3, PRPF39, LEF1, NUP85, MBD3, EXOSC1, TTF2, NOC2L, MED6, EIF4A3, TAF13, FANCD2, SNRNP200, UCHL5, CNOT11, SNRNP40, JMJD1C, CUX1, THOC3, C1D, ING2, ZBTB10, TRA2B, UTP6, NOB1, HCFC2, POLR2B, CHD9, SUMO1, CHD7, SAFB, LANCL2, PRPF8, USP39, CAMK2D, MYCBP, ETV6, CHD3, NSA2, PDCD11, PPP1R10, RPL27, PHF10, SAP30BP, SF3A3, PPP1R9B, ATXN2, MED30, SUGP1, RPL23, SP3, TCEB3 |
| GO:0006396  RNA processing  (60) | 5.08  (6.93E-26) | INTS3, WTAP, CACTIN, SMNDC1, NONO, RRP1B, INTS5, U2AF1, LUC7L3, CDK13, ZCCHC8, NOL6, KRR1, GTPBP4, CHTOP, RRP7A, MAGOH, PTBP1, PRPF39, EXOSC3, EXOSC1, TTF2, EIF4A3, AQR, DHX29, SNRNP200, SNRNP40, SCAF8, THOC3, C1D, FUS, TRA2B, UTP6, SRSF11, NOB1, POLR2B, RPS25, CHD7, HNRNPK, RPS29, ZNF326, PRPF8, USP39, ISY1, LEO1, RPL10A, NSA2, PRPF40A, TBL3, PDCD11, TSR1, RPL27, SMAD2, SF3A3, SRSF2, PPP1R9B, HNRNPH3, SUGP1, RPL23, HNRNPH1 |
