## Supplementary material for "Comparative analysis of retroviral Gag-host cell interactions: focus on the nuclear interactome": Table S3

| **Table S3.**  Top 10 DAVID biological processes for proteins isolated from the purified HIV Gag affinity purifications from HeLa nuclear lysates. | | |
| --- | --- | --- |
| **Gene Ontology Term (protein count)** | **Fold Enrichment (p-value)** | **Gene names for proteins isolated** |
| GO:0006396  RNA processing  (105) | 4.62  (3.10E-41) | RNMT, AAR2, RPL15, SNRPD2, RBM7, CMTR1, RPS27L, INTS9, DDX27, IMP3, CDKN2A, RAVER2, CLP1, TARDBP, DNAJC8, DHX34, U2AF1, FAU, SRRM1, LSM3, DDX20, RBM10, DHX30, CCAR2, PABPN1, CHTOP, BYSL, PTBP1, RRP9, HNRNPR, RSL1D1, PCF11, NOP2, SNRPA, SCAF8, CPSF3L, FIP1L1, SRSF12, ZFC3H1, HNRNPM, DDX47, RPS28, DDX46, CNOT6L, LEO1, NAT10, RPL5, PRPF40A, RPS24, CCDC86, NOC4L, SMAD2, SRSF3, PPIH, HNRNPH3, SRSF7, WDR3, RPL37A, DHX40, UTP20, NCOR1, PUF60, PLRG1, DKC1, PCBP1, CDK12, MDN1, SYMPK, EXOSC4, EXOSC2, GTF2H4, GTF2H3, PRPF3, CDC5L, SMN2, PRPF4, SRPK1, GTF2H1, SENP3, MRPS9, C1QBP, LARP7, CPSF7, CPSF4, CPSF3, THOC3, NHP2, PRPF38B, POLR2E, FAM98B, TRA2B, POLR2B, DIMT1, RBM26, PDCD11, ALYREF, GRSF1, ELAVL1, HNRNPDL, FBL, HNRNPUL1, POLDIP3, NGDN, PES1, TEX10 |
| GO:0016071  mRNA metabolic process  (84) | 5.00  (4.71E-35) | CNOT9, RNMT, AAR2, ZC3HAV1, RPL15, SNRPD2, RBM7, CMTR1, DKC1, PLRG1, RAVER2, PCBP1, CLP1, TARDBP, DNAJC8, DHX34, U2AF1, CDK12, SRRM1, FAU, LSM3, DDX20, RBM10, CCAR2, PABPN1, SYMPK, CHTOP, EXOSC4, PTBP1, EXOSC2, GTF2H4, GTF2H3, PRPF3, CDC5L, SMN2, PRPF4, HNRNPR, SRPK1, GTF2H1, PCF11, C1QBP, CPSF7, SNRPA, CDK11B, CPSF4, CPSF3, SCAF8, THOC3, XRN1, PRPF38B, FIP1L1, POLR2E, SRSF12, TRA2B, SKIV2L, POLR2B, HNRNPM, MOV10, DDX47, RPS28, DDX46, CNOT6L, SAFB, LEO1, RPL5, RBM26, RPS24, PRPF40A, SMG9, PDCD11, UPF1, ALYREF, GRSF1, ELAVL1, ETF1, SAFB2, SRSF3, HNRNPH3, PPIH, SRSF7, HNRNPUL1, POLDIP3, RPL37A, PUF60 |
| GO:0010467  Gene expression  (253) | 1.91  (1.66E-34) | AAR2, RPL15, MED23, CMTR1, MED21, CTNNB1, INTS9, MED28, RAVER2, MED26, ILK, DHX34, U2AF1, FAU, LSM3, OGT, DHX30, CCAR2, MRPL53, CHTOP, BYSL, PTBP1, MECP2, HMG20B, MED13, MAPK1, PCF11, PPP1CA, NOP2, BAZ1A, MED15, RFC1, PTRF, BAZ1B, EIF2S1, SERBP1, TGFB1I1, SCAF8, SUDS3, CPSF3L, FIP1L1, SRSF12, IGF2BP1, IGF2BP3, MOV10, RPS28, LEO1, PBRM1, NAT10, MEPCE, RPS24, PRPF40A, CCDC86, NOC4L, SMAD5, CREBBP, SMAD2, AFG3L2, FXR1, SRSF3, SRSF7, WDR61, PTCD3, TRPS1, SUPT16H, DHX40, UTP20, NCOR1, ABCF1, SPIN1, NARS, QARS, ZEB1, VWA9, ASH2L, SLC25A3, SYMPK, HIST1H1E, SLC25A4, EXOSC4, EXOSC2, GTF2H4, GTF2H3, MBD2, SMN2, SRPK1, GTF2H1, TRAP1, SENP3, MED9, CDK11B, NHP2, THOC3, SRP9, PRPF38B, POLR2E, MTDH, ASUN, FAM98B, CIC, POLR2B, MRPL10, SQSTM1, NUP210, GATAD2A, MLLT3, RBM26, UPF1, ZNF24, ELAVL1, ILF3, ETF1, HNRNPDL, BRMS1, ILF2, HNRNPUL1, POLDIP3, PBX1, TCEB1, PES1, RBM14, TEX10, DNM2, CNOT9, MRPS34, RNMT, INO80, SNRPD2, RBM7, RPS27L, MRPS31, CSNK2A2, DDX27, TOP1, IMP3, RAD21, CDKN2A, SMARCD2, SMARCD3, CGGBP1, TARDBP, CLP1, DNAJC8, PUM1, SRRM1, ASPH, DDX20, RBM10, PABPN1, RRP9, HNRNPR, RSL1D1, SUZ12, TRIM33, KDM2A, HSPB1, SNRPA, ERC1, XRN1, SMARCA4, EEF1B2, MRPS14, SETD1A, CLU, TRRAP, ZFC3H1, HNRNPM, DDX47, DDX46, MEIS2, SMARCB1, CNOT6L, EIF3F, UBAP2L, RPL5, DNAJA3, HELLS, HIST2H3A, MRPS27, TRIP4, MRPS23, MYO1C, MRPS22, WDR5, EHMT2, FOXP1, SAFB2, SLC25A11, HNRNPH3, PPIH, HDAC3, SLC25A13, PSMC3, SMARCC2, DNMT1, WDR3, CARS2, RPL37A, TMPO, PUF60, DAP3, HP1BP3, ARID4B, NUP188, DKC1, PLRG1, PCBP1, PSMD2, CDK12, PSMD5, MDN1, AHNAK, ELMSAN1, RING1, PRPF3, UBE2I, CDC5L, PRPF4, PURA, CD3EAP, EIF4G2, MRPS9, C1QBP, NCOA5, LARP7, CPSF7, CPSF4, CPSF3, CTSG, DPF2, BTAF1, NUP98, ING2, TRA2B, RDX, NFYA, PAXBP1, IARS, MINA, DIMT1, SAFB, GTF3C2, GAPDH, ZBTB7B, PDCD11, ALYREF, GRSF1, CENPF, TAB1, FBL, ATXN2, NGDN, SETD2 |
| GO:0034641  Cellular nitrogen compound metabolic process  (283) | 1.74  (1.79E-33) | AAR2, RPL15, MED23, CMTR1, MED21, CTNNB1, INTS9, MED28, RAVER2, MED26, ILK, DHX34, U2AF1, FAU, LSM3, OGT, DHX30, CCAR2, MRPL53, CHTOP, BYSL, PTBP1, MECP2, HMG20B, MED13, RFC5, PCF11, MAPK1, PPP1CA, NOP2, NNT, BAZ1A, HUWE1, MED15, RFC1, PTRF, BAZ1B, EIF2S1, ADSL, TGFB1I1, SCAF8, SUDS3, CPSF3L, FIP1L1, SRSF12, IGF2BP1, IGF2BP3, MOV10, RPS28, LEO1, PBRM1, NAT10, MEPCE, RPS24, PRPF40A, CCDC86, NOC4L, MKI67, INIP, SMAD5, CREBBP, SMAD2, FXR1, CCT7, SRSF3, CCT5, CCT4, SRSF7, WDR61, PTCD3, TRPS1, CCT8, SUPT16H, DHX40, UTP20, NCOR1, ABCF1, SPIN1, NARS, QARS, ZEB1, ANKRD17, VWA9, ASH2L, FANCI, SLC25A3, SYMPK, HIST1H1E, NDUFB10, SLC25A4, EXOSC4, EXOSC2, GTF2H4, GTF2H3, CCT6A, MBD2, SMN2, SRPK1, GTF2H1, TRAP1, SENP3, MED9, CDK11B, THOC3, NHP2, SRP9, PRPF38B, ASUN, POLR2E, MTDH, FAM98B, SKIV2L, CIC, POLR2B, MRPL10, SQSTM1, NUP210, GATAD2A, TRIP12, MLLT3, RBM26, UPF1, ZNF24, ELAVL1, ILF3, ETF1, HNRNPDL, BRMS1, ILF2, TFRC, HNRNPUL1, POLDIP3, PBX1, TCEB1, RBM14, PES1, TEX10, CALM2, DNM2, CNOT9, MRPS34, RNMT, ZC3HAV1, INO80, SNRPD2, RBM7, RPS27L, MRPS31, CSNK2A2, DDX27, TOP1, IMP3, RAD21, CDKN2A, SMARCD2, SMARCD3, CGGBP1, TARDBP, CLP1, DNAJC8, DDX24, PUM1, ORC5, SRRM1, ASPH, DDX20, RBM10, ORC3, PABPN1, ACTN4, ACTN1, RRP9, HNRNPR, RSL1D1, SUZ12, TRIM33, KDM2A, HSPB1, RAD18, SNRPA, DDX31, ERC1, XRN1, SMARCA4, EEF1B2, MRPS14, SETD1A, CLU, TRRAP, ZFC3H1, HNRNPM, DDX47, DDX46, MEIS2, SMARCB1, CNOT6L, EIF3F, RPL5, DNAJA3, HELLS, HIST2H3A, MRPS27, SMG9, TRIP4, MRPS23, MRPS22, WDR5, EHMT2, FOXP1, SAFB2, SLC25A11, HNRNPH3, PPIH, HDAC3, SLC25A13, PSMC3, SMARCC2, DNMT1, WDR3, CARS2, RPL37A, TMPO, SMC1A, PUF60, DAP3, HP1BP3, ARID4B, NUP188, UQCRQ, PLRG1, DKC1, PCBP1, PSMD2, CDK12, PSMD5, NDUFS3, MDN1, AHNAK, ELMSAN1, RING1, PRPF3, UBE2I, CDC5L, PRPF4, MCM4, PURA, CD3EAP, EIF4G2, MRPS9, C1QBP, RIF1, NCOA5, LARP7, CPSF7, CPSF4, CPSF3, CTSG, DPF2, BTAF1, NUP98, ING2, TRA2B, NFYA, PAXBP1, IARS, MINA, DIMT1, SAFB, GTF3C2, GAPDH, ZBTB7B, TCP1, PDCD11, NDUFA9, ALYREF, GRSF1, CENPF, TAB1, FBL, ATXN2, CUL4A, NGDN, SETD2 |
| GO:0090304  Nucleic acid metabolic process  (245) | 1.90  (1.70E-32) | AAR2, RPL15, MED23, CMTR1, MED21, CTNNB1, INTS9, MED28, RAVER2, MED26, ILK, DHX34, U2AF1, FAU, LSM3, OGT, DHX30, CCAR2, CHTOP, BYSL, PTBP1, MECP2, HMG20B, MED13, RFC5, MAPK1, PCF11, NOP2, BAZ1A, MED15, RFC1, PTRF, BAZ1B, HUWE1, TGFB1I1, SCAF8, SUDS3, CPSF3L, FIP1L1, SRSF12, MOV10, RPS28, LEO1, PBRM1, NAT10, MEPCE, RPS24, PRPF40A, CCDC86, NOC4L, MKI67, INIP, SMAD5, CREBBP, SMAD2, SRSF3, CCT7, CCT5, CCT4, SRSF7, WDR61, TRPS1, CCT8, SUPT16H, DHX40, UTP20, NCOR1, SPIN1, NARS, QARS, ZEB1, ANKRD17, VWA9, ASH2L, FANCI, SYMPK, HIST1H1E, EXOSC4, EXOSC2, GTF2H4, GTF2H3, CCT6A, MBD2, SMN2, SRPK1, GTF2H1, SENP3, MED9, CDK11B, NHP2, THOC3, PRPF38B, POLR2E, MTDH, ASUN, FAM98B, SKIV2L, CIC, POLR2B, SQSTM1, NUP210, GATAD2A, TRIP12, MLLT3, RBM26, UPF1, ZNF24, ELAVL1, ILF3, ETF1, HNRNPDL, BRMS1, ILF2, TFRC, HNRNPUL1, POLDIP3, PBX1, TCEB1, PES1, RBM14, TEX10, DNM2, CNOT9, RNMT, ZC3HAV1, INO80, SNRPD2, RBM7, RPS27L, CSNK2A2, DDX27, TOP1, IMP3, RAD21, CDKN2A, SMARCD2, SMARCD3, CGGBP1, TARDBP, CLP1, DDX24, DNAJC8, ORC5, SRRM1, ASPH, DDX20, RBM10, ORC3, PABPN1, ACTN4, ACTN1, RRP9, HNRNPR, RSL1D1, SUZ12, TRIM33, KDM2A, RAD18, SNRPA, DDX31, ERC1, XRN1, SMARCA4, SETD1A, CLU, TRRAP, ZFC3H1, HNRNPM, DDX47, DDX46, MEIS2, CNOT6L, SMARCB1, RPL5, DNAJA3, HELLS, HIST2H3A, SMG9, TRIP4, WDR5, EHMT2, FOXP1, SAFB2, HNRNPH3, PPIH, HDAC3, PSMC3, SMARCC2, DNMT1, WDR3, CARS2, RPL37A, TMPO, SMC1A, PUF60, HP1BP3, ARID4B, NUP188, DKC1, PLRG1, PCBP1, CDK12, MDN1, AHNAK, ELMSAN1, RING1, PRPF3, UBE2I, CDC5L, PRPF4, MCM4, PURA, CD3EAP, MRPS9, C1QBP, RIF1, NCOA5, LARP7, CPSF7, CPSF4, CPSF3, DPF2, BTAF1, NUP98, ING2, TRA2B, NFYA, PAXBP1, IARS, MINA, DIMT1, SAFB, GTF3C2, ZBTB7B, TCP1, PDCD11, ALYREF, GRSF1, CENPF, TAB1, FBL, ATXN2, CUL4A, NGDN, SETD2 |
| GO:0006807  Nitrogen compound metabolic process  (289) | 1.67  (8.23E-31) | AAR2, RPL15, MED23, CMTR1, MED21, CTNNB1, INTS9, CD44, MED28, RAVER2, MED26, ILK, DHX34, U2AF1, FAU, LSM3, OGT, DHX30, CCAR2, MRPL53, CHTOP, BYSL, PTBP1, MECP2, HMG20B, MED13, RFC5, PCF11, MAPK1, PPP1CA, NOP2, NNT, BAZ1A, HUWE1, MED15, RFC1, PTRF, BAZ1B, EIF2S1, ADSL, TGFB1I1, SCAF8, SUDS3, CPSF3L, FIP1L1, SRSF12, IGF2BP1, IGF2BP3, MOV10, RPS28, LEO1, PBRM1, NAT10, MEPCE, RPS24, PRPF40A, CCDC86, NOC4L, MKI67, INIP, SMAD5, CREBBP, SMAD2, FXR1, CCT7, SRSF3, CCT5, CCT4, SRSF7, WDR61, PTCD3, TRPS1, CCT8, SUPT16H, DHX40, UTP20, NCOR1, ABCF1, SPIN1, NARS, QARS, ZEB1, ANKRD17, VWA9, ASH2L, FANCI, SLC25A3, SYMPK, HIST1H1E, NDUFB10, SLC25A4, EXOSC4, EXOSC2, GTF2H4, GTF2H3, CCT6A, MBD2, SMN2, SRPK1, GTF2H1, TRAP1, SENP3, MED9, CDK11B, THOC3, NHP2, SRP9, PRPF38B, ASUN, POLR2E, MTDH, FAM98B, SKIV2L, CIC, POLR2B, MRPL10, SQSTM1, NUP210, GATAD2A, TRIP12, MLLT3, RBM26, UPF1, ZNF24, ELAVL1, ILF3, ETF1, HNRNPDL, BRMS1, ILF2, TFRC, HNRNPUL1, POLDIP3, PBX1, TCEB1, RBM14, PES1, TEX10, CALM2, DNM2, CNOT9, MRPS34, RNMT, ZC3HAV1, INO80, SNRPD2, RBM7, RPS27L, MRPS31, CSNK2A2, DDX27, TOP1, IMP3, RAD21, CDKN2A, SMARCD2, SMARCD3, CGGBP1, TARDBP, CLP1, DNAJC8, DDX24, PUM1, ORC5, SRRM1, ASPH, DDX20, RBM10, ORC3, PABPN1, ACTN4, ACTN1, RRP9, HNRNPR, RSL1D1, SUZ12, PYCR1, PYCR2, TRIM33, KDM2A, HSPB1, RAD18, SNRPA, DDX31, ERC1, XRN1, SMARCA4, EEF1B2, MRPS14, SETD1A, CLU, TRRAP, ZFC3H1, HNRNPM, CEPT1, DDX47, DDX46, MEIS2, SMARCB1, CNOT6L, EIF3F, RPL5, DNAJA3, HELLS, HIST2H3A, MRPS27, SMG9, TRIP4, MRPS23, MRPS22, WDR5, EHMT2, FOXP1, SAFB2, SLC25A11, PPIH, HNRNPH3, HDAC3, SLC25A13, PSMC3, SMARCC2, DNMT1, WDR3, CARS2, RPL37A, TMPO, SMC1A, PUF60, DAP3, HP1BP3, ARID4B, NUP188, CLTC, UQCRQ, PLRG1, DKC1, PCBP1, PSMD2, CDK12, PSMD5, NDUFS3, PTDSS1, MDN1, AHNAK, ELMSAN1, RING1, PRPF3, UBE2I, CDC5L, PRPF4, MCM4, PURA, CD3EAP, EIF4G2, MRPS9, C1QBP, RIF1, NCOA5, LARP7, CPSF7, CPSF4, CPSF3, CTSG, DPF2, BTAF1, NUP98, ING2, TRA2B, NFYA, PAXBP1, IARS, MINA, DIMT1, SAFB, GTF3C2, GAPDH, ZBTB7B, TCP1, PDCD11, NDUFA9, ALYREF, GRSF1, CENPF, TAB1, FBL, ATXN2, CUL4A, NGDN, SETD2 |
| GO:0016070  RNA metabolic process  (223) | 1.92  (7.99E-29) | AAR2, RPL15, MED23, CMTR1, MED21, CTNNB1, INTS9, MED28, RAVER2, MED26, ILK, DHX34, U2AF1, FAU, LSM3, OGT, DHX30, CCAR2, CHTOP, BYSL, PTBP1, MECP2, HMG20B, MED13, MAPK1, PCF11, NOP2, BAZ1A, MED15, RFC1, PTRF, BAZ1B, TGFB1I1, SCAF8, SUDS3, CPSF3L, FIP1L1, SRSF12, MOV10, RPS28, LEO1, PBRM1, NAT10, MEPCE, RPS24, PRPF40A, CCDC86, NOC4L, SMAD5, CREBBP, SMAD2, SRSF3, SRSF7, WDR61, TRPS1, SUPT16H, DHX40, UTP20, NCOR1, SPIN1, NARS, QARS, ZEB1, VWA9, ASH2L, SYMPK, HIST1H1E, EXOSC4, EXOSC2, GTF2H4, GTF2H3, MBD2, SMN2, SRPK1, GTF2H1, SENP3, MED9, CDK11B, NHP2, THOC3, PRPF38B, POLR2E, MTDH, ASUN, FAM98B, SKIV2L, CIC, POLR2B, SQSTM1, NUP210, GATAD2A, MLLT3, RBM26, UPF1, ZNF24, ELAVL1, ILF3, ETF1, HNRNPDL, BRMS1, ILF2, HNRNPUL1, POLDIP3, PBX1, TCEB1, PES1, RBM14, TEX10, DNM2, CNOT9, RNMT, ZC3HAV1, SNRPD2, INO80, RBM7, RPS27L, CSNK2A2, DDX27, IMP3, CDKN2A, RAD21, SMARCD2, SMARCD3, CGGBP1, TARDBP, CLP1, DDX24, DNAJC8, SRRM1, DDX20, ASPH, RBM10, PABPN1, ACTN4, ACTN1, RRP9, HNRNPR, SUZ12, RSL1D1, TRIM33, KDM2A, SNRPA, DDX31, ERC1, XRN1, SMARCA4, SETD1A, CLU, TRRAP, ZFC3H1, HNRNPM, DDX47, DDX46, MEIS2, CNOT6L, SMARCB1, RPL5, DNAJA3, HELLS, HIST2H3A, SMG9, TRIP4, WDR5, EHMT2, FOXP1, SAFB2, HNRNPH3, PPIH, HDAC3, PSMC3, SMARCC2, DNMT1, WDR3, CARS2, RPL37A, TMPO, PUF60, HP1BP3, ARID4B, NUP188, DKC1, PLRG1, PCBP1, CDK12, MDN1, AHNAK, ELMSAN1, RING1, PRPF3, UBE2I, CDC5L, PRPF4, PURA, CD3EAP, MRPS9, C1QBP, NCOA5, LARP7, CPSF7, CPSF4, CPSF3, DPF2, BTAF1, NUP98, ING2, TRA2B, PAXBP1, NFYA, IARS, DIMT1, MINA, SAFB, GTF3C2, ZBTB7B, PDCD11, ALYREF, GRSF1, CENPF, TAB1, FBL, ATXN2, NGDN, SETD2 |
| GO:0006139  Nucleobase-containing compound metabolic process  (254) | 1.76  (2.80E-28) | AAR2, RPL15, MED23, CMTR1, MED21, CTNNB1, INTS9, MED28, RAVER2, MED26, ILK, DHX34, U2AF1, FAU, LSM3, OGT, DHX30, CCAR2, CHTOP, BYSL, PTBP1, MECP2, HMG20B, MED13, RFC5, MAPK1, PCF11, NOP2, NNT, BAZ1A, MED15, RFC1, PTRF, BAZ1B, HUWE1, ADSL, TGFB1I1, SCAF8, SUDS3, CPSF3L, FIP1L1, SRSF12, MOV10, RPS28, LEO1, PBRM1, NAT10, MEPCE, RPS24, PRPF40A, CCDC86, NOC4L, MKI67, INIP, SMAD5, CREBBP, SMAD2, CCT7, SRSF3, CCT5, CCT4, SRSF7, WDR61, TRPS1, CCT8, SUPT16H, DHX40, UTP20, NCOR1, SPIN1, NARS, QARS, ZEB1, ANKRD17, VWA9, ASH2L, FANCI, SYMPK, HIST1H1E, NDUFB10, EXOSC4, EXOSC2, GTF2H4, GTF2H3, CCT6A, MBD2, SMN2, SRPK1, GTF2H1, SENP3, MED9, CDK11B, NHP2, THOC3, PRPF38B, ASUN, POLR2E, MTDH, FAM98B, SKIV2L, CIC, POLR2B, SQSTM1, NUP210, GATAD2A, TRIP12, MLLT3, RBM26, UPF1, ZNF24, ELAVL1, ILF3, ETF1, HNRNPDL, BRMS1, ILF2, TFRC, HNRNPUL1, POLDIP3, PBX1, TCEB1, PES1, RBM14, TEX10, CALM2, DNM2, CNOT9, RNMT, ZC3HAV1, INO80, SNRPD2, RBM7, RPS27L, CSNK2A2, DDX27, TOP1, IMP3, RAD21, CDKN2A, SMARCD2, SMARCD3, CGGBP1, TARDBP, CLP1, DDX24, DNAJC8, ORC5, SRRM1, ASPH, DDX20, RBM10, ORC3, PABPN1, ACTN4, ACTN1, RRP9, HNRNPR, RSL1D1, SUZ12, TRIM33, KDM2A, RAD18, SNRPA, DDX31, ERC1, XRN1, SMARCA4, SETD1A, CLU, TRRAP, ZFC3H1, HNRNPM, DDX47, DDX46, MEIS2, CNOT6L, SMARCB1, RPL5, DNAJA3, HELLS, HIST2H3A, SMG9, TRIP4, WDR5, EHMT2, FOXP1, SAFB2, HNRNPH3, PPIH, HDAC3, SLC25A13, PSMC3, SMARCC2, DNMT1, WDR3, CARS2, RPL37A, TMPO, SMC1A, PUF60, HP1BP3, ARID4B, NUP188, UQCRQ, PLRG1, DKC1, PCBP1, CDK12, NDUFS3, MDN1, AHNAK, ELMSAN1, RING1, PRPF3, UBE2I, CDC5L, PRPF4, MCM4, PURA, CD3EAP, MRPS9, C1QBP, RIF1, NCOA5, LARP7, CPSF7, CPSF4, CPSF3, DPF2, BTAF1, NUP98, ING2, TRA2B, NFYA, PAXBP1, IARS, MINA, DIMT1, SAFB, GTF3C2, GAPDH, ZBTB7B, TCP1, PDCD11, NDUFA9, ALYREF, GRSF1, CENPF, TAB1, FBL, ATXN2, CUL4A, NGDN, SETD2 |
| GO:0006397  mRNA processing  (63) | 5.36  (1.18E-27) | RNMT, AAR2, SNRPD2, RBM7, CMTR1, PLRG1, RAVER2, PCBP1, CLP1, TARDBP, DNAJC8, U2AF1, CDK12, SRRM1, LSM3, DDX20, RBM10, CCAR2, PABPN1, SYMPK, CHTOP, PTBP1, GTF2H4, GTF2H3, PRPF3, CDC5L, HNRNPR, PRPF4, SMN2, SRPK1, GTF2H1, PCF11, C1QBP, CPSF7, SNRPA, CPSF4, CPSF3, SCAF8, THOC3, PRPF38B, FIP1L1, POLR2E, SRSF12, TRA2B, POLR2B, HNRNPM, DDX47, DDX46, CNOT6L, LEO1, PRPF40A, RBM26, PDCD11, ALYREF, GRSF1, ELAVL1, SRSF3, HNRNPH3, PPIH, SRSF7, HNRNPUL1, POLDIP3, PUF60 |
| GO:0046483  Heterocycle metabolic process  (254) | 1.72  (1.04E-26) | AAR2, RPL15, MED23, CMTR1, MED21, CTNNB1, INTS9, MED28, RAVER2, MED26, ILK, DHX34, U2AF1, FAU, LSM3, OGT, DHX30, CCAR2, CHTOP, BYSL, PTBP1, MECP2, HMG20B, MED13, RFC5, MAPK1, PCF11, NOP2, NNT, BAZ1A, MED15, RFC1, PTRF, BAZ1B, HUWE1, ADSL, TGFB1I1, SCAF8, SUDS3, CPSF3L, FIP1L1, SRSF12, MOV10, RPS28, LEO1, PBRM1, NAT10, MEPCE, RPS24, PRPF40A, CCDC86, NOC4L, MKI67, INIP, SMAD5, CREBBP, SMAD2, CCT7, SRSF3, CCT5, CCT4, SRSF7, WDR61, TRPS1, CCT8, SUPT16H, DHX40, UTP20, NCOR1, SPIN1, NARS, QARS, ZEB1, ANKRD17, VWA9, ASH2L, FANCI, SYMPK, HIST1H1E, NDUFB10, EXOSC4, EXOSC2, GTF2H4, GTF2H3, CCT6A, MBD2, SMN2, SRPK1, GTF2H1, SENP3, MED9, CDK11B, NHP2, THOC3, PRPF38B, ASUN, POLR2E, MTDH, FAM98B, SKIV2L, CIC, POLR2B, SQSTM1, NUP210, GATAD2A, TRIP12, MLLT3, RBM26, UPF1, ZNF24, ELAVL1, ILF3, ETF1, HNRNPDL, BRMS1, ILF2, TFRC, HNRNPUL1, POLDIP3, PBX1, TCEB1, PES1, RBM14, TEX10, CALM2, DNM2, CNOT9, RNMT, ZC3HAV1, INO80, SNRPD2, RBM7, RPS27L, CSNK2A2, DDX27, TOP1, IMP3, RAD21, CDKN2A, SMARCD2, SMARCD3, CGGBP1, TARDBP, CLP1, DDX24, DNAJC8, ORC5, SRRM1, ASPH, DDX20, RBM10, ORC3, PABPN1, ACTN4, ACTN1, RRP9, HNRNPR, RSL1D1, SUZ12, TRIM33, KDM2A, RAD18, SNRPA, DDX31, ERC1, XRN1, SMARCA4, SETD1A, CLU, TRRAP, ZFC3H1, HNRNPM, DDX47, DDX46, MEIS2, CNOT6L, SMARCB1, RPL5, DNAJA3, HELLS, HIST2H3A, SMG9, TRIP4, WDR5, EHMT2, FOXP1, SAFB2, HNRNPH3, PPIH, HDAC3, SLC25A13, PSMC3, SMARCC2, DNMT1, WDR3, CARS2, RPL37A, TMPO, SMC1A, PUF60, HP1BP3, ARID4B, NUP188, UQCRQ, PLRG1, DKC1, PCBP1, CDK12, NDUFS3, MDN1, AHNAK, ELMSAN1, RING1, PRPF3, UBE2I, CDC5L, PRPF4, MCM4, PURA, CD3EAP, MRPS9, C1QBP, RIF1, NCOA5, LARP7, CPSF7, CPSF4, CPSF3, DPF2, BTAF1, NUP98, ING2, TRA2B, NFYA, PAXBP1, IARS, MINA, DIMT1, SAFB, GTF3C2, GAPDH, ZBTB7B, TCP1, PDCD11, NDUFA9, ALYREF, GRSF1, CENPF, TAB1, FBL, ATXN2, CUL4A, NGDN, SETD2 |
