## Supplementary material for "Comparative analysis of retroviral Gag-host cell interactions: focus on the nuclear interactome": Table S4

| **Table S4.**  Top 10 nuclear enriched DAVID biological processes for proteins isolated from the purified HIV Gag affinity purifications from HeLa nuclear lysates. | | |
| --- | --- | --- |
| **Gene Ontology Term (protein count)** | **Fold Enrichment (p-value)** | **Gene names for proteins isolated** |
| GO:0090304  Nucleic acid metabolic process  (229) | 2.46  (9.02E-59) | RPL15, MED23, CMTR1, MED21, CTNNB1, INTS9, MED28, RAVER2, MED26, ILK, DHX34, U2AF1, FAU, LSM3, OGT, DHX30, CCAR2, CHTOP, BYSL, PTBP1, MECP2, HMG20B, MED13, RFC5, MAPK1, PCF11, NOP2, BAZ1A, MED15, RFC1, PTRF, BAZ1B, HUWE1, TGFB1I1, SCAF8, SUDS3, CPSF3L, FIP1L1, SRSF12, RPS28, LEO1, PBRM1, NAT10, RPS24, PRPF40A, CCDC86, NOC4L, MKI67, INIP, SMAD5, CREBBP, SMAD2, SRSF3, CCT5, CCT4, SRSF7, WDR61, TRPS1, CCT8, SUPT16H, DHX40, UTP20, NCOR1, SPIN1, ZEB1, ANKRD17, VWA9, ASH2L, FANCI, SYMPK, HIST1H1E, EXOSC4, EXOSC2, GTF2H4, GTF2H3, MBD2, SMN2, SRPK1, GTF2H1, SENP3, MED9, CDK11B, NHP2, THOC3, PRPF38B, POLR2E, MTDH, ASUN, FAM98B, SKIV2L, CIC, POLR2B, SQSTM1, NUP210, GATAD2A, TRIP12, MLLT3, UPF1, ZNF24, ELAVL1, ILF3, ETF1, HNRNPDL, BRMS1, ILF2, HNRNPUL1, POLDIP3, PBX1, TCEB1, PES1, RBM14, TEX10, DNM2, CNOT9, RNMT, ZC3HAV1, SNRPD2, INO80, RBM7, RPS27L, CSNK2A2, DDX27, TOP1, IMP3, RAD21, CDKN2A, SMARCD2, SMARCD3, CGGBP1, TARDBP, CLP1, DDX24, DNAJC8, ORC5, SRRM1, DDX20, RBM10, ORC3, PABPN1, ACTN4, RRP9, HNRNPR, RSL1D1, SUZ12, TRIM33, KDM2A, RAD18, SNRPA, DDX31, XRN1, SMARCA4, SETD1A, CLU, TRRAP, HNRNPM, DDX47, DDX46, MEIS2, CNOT6L, SMARCB1, RPL5, DNAJA3, HELLS, HIST2H3A, TRIP4, WDR5, EHMT2, FOXP1, SAFB2, HNRNPH3, PPIH, HDAC3, PSMC3, SMARCC2, DNMT1, WDR3, RPL37A, TMPO, SMC1A, PUF60, HP1BP3, ARID4B, NUP188, DKC1, PLRG1, PCBP1, CDK12, MDN1, AHNAK, ELMSAN1, RING1, PRPF3, UBE2I, CDC5L, PRPF4, MCM4, PURA, CD3EAP, MRPS9, C1QBP, RIF1, NCOA5, LARP7, CPSF7, CPSF4, CPSF3, DPF2, BTAF1, NUP98, ING2, TRA2B, NFYA, PAXBP1, IARS, MINA, DIMT1, SAFB, GTF3C2, ZBTB7B, TCP1, PDCD11, ALYREF, CENPF, TAB1, FBL, ATXN2, CUL4A, NGDN, SETD2 |
| GO:0034641  Cellular nitrogen compound metabolic process  (253) | 2.16  (1.64E-58) | RPL15, MED23, CMTR1, MED21, CTNNB1, INTS9, MED28, RAVER2, MED26, ILK, DHX34, U2AF1, FAU, LSM3, OGT, DHX30, CCAR2, CHTOP, BYSL, PTBP1, MECP2, HMG20B, MED13, RFC5, MAPK1, PCF11, PPP1CA, NOP2, BAZ1A, MED15, RFC1, PTRF, BAZ1B, HUWE1, EIF2S1, TGFB1I1, SCAF8, SUDS3, CPSF3L, FIP1L1, SRSF12, IGF2BP1, IGF2BP3, RPS28, LEO1, PBRM1, NAT10, RPS24, PRPF40A, CCDC86, NOC4L, MKI67, INIP, SMAD5, CREBBP, SMAD2, FXR1, SRSF3, CCT5, CCT4, SRSF7, WDR61, TRPS1, CCT8, SUPT16H, DHX40, UTP20, NCOR1, ABCF1, SPIN1, ZEB1, ANKRD17, VWA9, ASH2L, FANCI, SLC25A3, SYMPK, HIST1H1E, SLC25A4, EXOSC4, EXOSC2, GTF2H4, GTF2H3, MBD2, SMN2, SRPK1, GTF2H1, TRAP1, SENP3, MED9, CDK11B, NHP2, THOC3, PRPF38B, ASUN, POLR2E, MTDH, FAM98B, SKIV2L, CIC, POLR2B, MRPL10, SQSTM1, NUP210, GATAD2A, TRIP12, MLLT3, UPF1, ZNF24, ELAVL1, ILF3, ETF1, HNRNPDL, BRMS1, ILF2, HNRNPUL1, POLDIP3, PBX1, TCEB1, PES1, RBM14, TEX10, CALM2, DNM2, CNOT9, RNMT, ZC3HAV1, INO80, SNRPD2, RBM7, RPS27L, MRPS31, CSNK2A2, DDX27, TOP1, IMP3, RAD21, CDKN2A, SMARCD2, SMARCD3, CGGBP1, TARDBP, CLP1, DNAJC8, DDX24, ORC5, SRRM1, DDX20, RBM10, ORC3, PABPN1, ACTN4, RRP9, HNRNPR, RSL1D1, SUZ12, TRIM33, KDM2A, HSPB1, RAD18, SNRPA, DDX31, XRN1, SMARCA4, EEF1B2, MRPS14, SETD1A, CLU, TRRAP, HNRNPM, DDX47, DDX46, MEIS2, SMARCB1, CNOT6L, RPL5, DNAJA3, HELLS, HIST2H3A, TRIP4, MRPS23, WDR5, EHMT2, FOXP1, SAFB2, SLC25A11, HNRNPH3, PPIH, HDAC3, PSMC3, SMARCC2, DNMT1, WDR3, RPL37A, TMPO, SMC1A, PUF60, DAP3, HP1BP3, ARID4B, NUP188, PLRG1, DKC1, PCBP1, PSMD2, CDK12, PSMD5, NDUFS3, MDN1, AHNAK, ELMSAN1, RING1, PRPF3, UBE2I, CDC5L, PRPF4, MCM4, PURA, CD3EAP, MRPS9, C1QBP, RIF1, NCOA5, LARP7, CPSF7, CPSF4, CPSF3, CTSG, DPF2, BTAF1, NUP98, ING2, TRA2B, NFYA, PAXBP1, IARS, MINA, DIMT1, SAFB, GTF3C2, GAPDH, ZBTB7B, TCP1, PDCD11, NDUFA9, ALYREF, CENPF, TAB1, FBL, ATXN2, CUL4A, NGDN, SETD2 |
| GO:0010467  Gene expression  (230) | 2.40  (3.66E-57) | RPL15, MED23, CMTR1, MED21, CTNNB1, INTS9, MED28, RAVER2, MED26, ILK, DHX34, U2AF1, FAU, LSM3, OGT, DHX30, CCAR2, CHTOP, BYSL, PTBP1, MECP2, HMG20B, MED13, MAPK1, PCF11, PPP1CA, NOP2, BAZ1A, MED15, RFC1, PTRF, BAZ1B, EIF2S1, SERBP1, TGFB1I1, SCAF8, SUDS3, CPSF3L, FIP1L1, SRSF12, IGF2BP1, IGF2BP3, RPS28, LEO1, PBRM1, NAT10, RPS24, PRPF40A, CCDC86, NOC4L, SMAD5, CREBBP, SMAD2, FXR1, SRSF3, SRSF7, WDR61, TRPS1, SUPT16H, DHX40, UTP20, NCOR1, ABCF1, SPIN1, ZEB1, VWA9, ASH2L, SLC25A3, SYMPK, HIST1H1E, SLC25A4, EXOSC4, EXOSC2, GTF2H4, GTF2H3, MBD2, SMN2, SRPK1, GTF2H1, SENP3, TRAP1, MED9, CDK11B, NHP2, THOC3, PRPF38B, POLR2E, MTDH, ASUN, FAM98B, CIC, POLR2B, MRPL10, SQSTM1, NUP210, GATAD2A, MLLT3, UPF1, ZNF24, ELAVL1, ILF3, ETF1, HNRNPDL, BRMS1, ILF2, HNRNPUL1, POLDIP3, PBX1, TCEB1, PES1, RBM14, TEX10, DNM2, CNOT9, RNMT, SNRPD2, INO80, RBM7, RPS27L, MRPS31, CSNK2A2, DDX27, TOP1, IMP3, RAD21, CDKN2A, SMARCD2, SMARCD3, CGGBP1, TARDBP, CLP1, DNAJC8, SRRM1, DDX20, RBM10, PABPN1, RRP9, HNRNPR, RSL1D1, SUZ12, TRIM33, KDM2A, HSPB1, SNRPA, XRN1, SMARCA4, EEF1B2, MRPS14, SETD1A, CLU, TRRAP, HNRNPM, DDX47, DDX46, MEIS2, CNOT6L, SMARCB1, RPL5, UBAP2L, DNAJA3, HELLS, HIST2H3A, TRIP4, MRPS23, MYO1C, WDR5, EHMT2, FOXP1, SAFB2, SLC25A11, HNRNPH3, PPIH, HDAC3, PSMC3, SMARCC2, DNMT1, WDR3, RPL37A, TMPO, PUF60, DAP3, HP1BP3, ARID4B, NUP188, DKC1, PLRG1, PCBP1, PSMD2, CDK12, PSMD5, MDN1, AHNAK, ELMSAN1, RING1, PRPF3, UBE2I, CDC5L, PRPF4, PURA, CD3EAP, MRPS9, C1QBP, NCOA5, LARP7, CPSF7, CPSF4, CPSF3, CTSG, DPF2, BTAF1, NUP98, ING2, TRA2B, NFYA, PAXBP1, IARS, MINA, DIMT1, SAFB, GTF3C2, GAPDH, ZBTB7B, PDCD11, ALYREF, CENPF, TAB1, FBL, ATXN2, NGDN, SETD2 |
| GO:0006807  Nitrogen compound metabolic process  (254) | 2.02  (5.43E-53) | RPL15, MED23, CMTR1, MED21, CTNNB1, INTS9, MED28, RAVER2, MED26, ILK, DHX34, U2AF1, FAU, LSM3, OGT, DHX30, CCAR2, CHTOP, BYSL, PTBP1, MECP2, HMG20B, MED13, RFC5, MAPK1, PCF11, PPP1CA, NOP2, BAZ1A, MED15, RFC1, PTRF, BAZ1B, HUWE1, EIF2S1, TGFB1I1, SCAF8, SUDS3, CPSF3L, FIP1L1, SRSF12, IGF2BP1, IGF2BP3, RPS28, LEO1, PBRM1, NAT10, RPS24, PRPF40A, CCDC86, NOC4L, MKI67, INIP, SMAD5, CREBBP, SMAD2, FXR1, SRSF3, CCT5, CCT4, SRSF7, WDR61, TRPS1, CCT8, SUPT16H, DHX40, UTP20, NCOR1, ABCF1, SPIN1, ZEB1, ANKRD17, VWA9, ASH2L, FANCI, SLC25A3, SYMPK, HIST1H1E, SLC25A4, EXOSC4, EXOSC2, GTF2H4, GTF2H3, MBD2, SMN2, SRPK1, GTF2H1, TRAP1, SENP3, MED9, CDK11B, NHP2, THOC3, PRPF38B, ASUN, POLR2E, MTDH, FAM98B, SKIV2L, CIC, POLR2B, MRPL10, SQSTM1, NUP210, GATAD2A, TRIP12, MLLT3, UPF1, ZNF24, ELAVL1, ILF3, ETF1, HNRNPDL, BRMS1, ILF2, HNRNPUL1, POLDIP3, PBX1, TCEB1, PES1, RBM14, TEX10, CALM2, DNM2, CNOT9, RNMT, ZC3HAV1, INO80, SNRPD2, RBM7, RPS27L, MRPS31, CSNK2A2, DDX27, TOP1, IMP3, RAD21, CDKN2A, SMARCD2, SMARCD3, CGGBP1, TARDBP, CLP1, DNAJC8, DDX24, ORC5, SRRM1, DDX20, RBM10, ORC3, PABPN1, ACTN4, RRP9, HNRNPR, RSL1D1, SUZ12, TRIM33, KDM2A, HSPB1, RAD18, SNRPA, DDX31, XRN1, SMARCA4, EEF1B2, MRPS14, SETD1A, CLU, TRRAP, HNRNPM, CEPT1, DDX47, DDX46, MEIS2, SMARCB1, CNOT6L, RPL5, DNAJA3, HELLS, HIST2H3A, TRIP4, MRPS23, WDR5, EHMT2, FOXP1, SAFB2, SLC25A11, HNRNPH3, PPIH, HDAC3, PSMC3, SMARCC2, DNMT1, WDR3, RPL37A, TMPO, SMC1A, PUF60, DAP3, HP1BP3, ARID4B, NUP188, PLRG1, DKC1, PCBP1, PSMD2, CDK12, PSMD5, NDUFS3, MDN1, AHNAK, ELMSAN1, RING1, PRPF3, UBE2I, CDC5L, PRPF4, MCM4, PURA, CD3EAP, MRPS9, C1QBP, RIF1, NCOA5, LARP7, CPSF7, CPSF4, CPSF3, CTSG, DPF2, BTAF1, NUP98, ING2, TRA2B, NFYA, PAXBP1, IARS, MINA, DIMT1, SAFB, GTF3C2, GAPDH, ZBTB7B, TCP1, PDCD11, NDUFA9, ALYREF, CENPF, TAB1, FBL, ATXN2, CUL4A, NGDN, SETD2 |
| GO:0006396  RNA processing  (101) | 6.15  (4.43E-52) | RNMT, RPL15, SNRPD2, CMTR1, RPS27L, RBM7, DDX27, INTS9, IMP3, CDKN2A, RAVER2, CLP1, TARDBP, DNAJC8, DHX34, U2AF1, FAU, SRRM1, LSM3, DDX20, RBM10, DHX30, CCAR2, PABPN1, CHTOP, BYSL, PTBP1, RRP9, HNRNPR, RSL1D1, PCF11, NOP2, SNRPA, SCAF8, CPSF3L, FIP1L1, SRSF12, HNRNPM, DDX47, DDX46, RPS28, CNOT6L, LEO1, NAT10, RPL5, PRPF40A, RPS24, CCDC86, NOC4L, SMAD2, SRSF3, PPIH, HNRNPH3, SRSF7, WDR3, RPL37A, DHX40, UTP20, NCOR1, PUF60, PLRG1, DKC1, PCBP1, CDK12, MDN1, SYMPK, EXOSC4, EXOSC2, GTF2H4, GTF2H3, PRPF3, CDC5L, SMN2, PRPF4, SRPK1, GTF2H1, SENP3, C1QBP, MRPS9, LARP7, CPSF7, CPSF4, CPSF3, THOC3, NHP2, PRPF38B, POLR2E, FAM98B, TRA2B, POLR2B, DIMT1, PDCD11, ALYREF, ELAVL1, HNRNPDL, FBL, HNRNPUL1, POLDIP3, NGDN, PES1, TEX10 |
| GO:0006139  Nucleobase-containing compound metabolic process  (233) | 2.23  (5.25E-52) | RPL15, MED23, CMTR1, MED21, CTNNB1, INTS9, MED28, RAVER2, MED26, ILK, DHX34, U2AF1, FAU, LSM3, OGT, DHX30, CCAR2, CHTOP, BYSL, PTBP1, MECP2, HMG20B, MED13, RFC5, MAPK1, PCF11, NOP2, BAZ1A, MED15, RFC1, PTRF, BAZ1B, HUWE1, TGFB1I1, SCAF8, SUDS3, CPSF3L, FIP1L1, SRSF12, RPS28, LEO1, PBRM1, NAT10, RPS24, PRPF40A, CCDC86, NOC4L, MKI67, INIP, SMAD5, CREBBP, SMAD2, SRSF3, CCT5, CCT4, SRSF7, WDR61, TRPS1, CCT8, SUPT16H, DHX40, UTP20, NCOR1, SPIN1, ZEB1, ANKRD17, VWA9, ASH2L, FANCI, SYMPK, HIST1H1E, EXOSC4, EXOSC2, GTF2H4, GTF2H3, MBD2, SMN2, SRPK1, GTF2H1, SENP3, MED9, CDK11B, NHP2, THOC3, PRPF38B, POLR2E, MTDH, ASUN, FAM98B, SKIV2L, CIC, POLR2B, SQSTM1, NUP210, GATAD2A, TRIP12, MLLT3, UPF1, ZNF24, ELAVL1, ILF3, ETF1, HNRNPDL, BRMS1, ILF2, HNRNPUL1, POLDIP3, PBX1, TCEB1, PES1, RBM14, TEX10, CALM2, DNM2, CNOT9, RNMT, ZC3HAV1, INO80, SNRPD2, RBM7, RPS27L, CSNK2A2, DDX27, TOP1, IMP3, RAD21, CDKN2A, SMARCD2, SMARCD3, CGGBP1, TARDBP, CLP1, DDX24, DNAJC8, ORC5, SRRM1, DDX20, RBM10, ORC3, PABPN1, ACTN4, RRP9, HNRNPR, RSL1D1, SUZ12, TRIM33, KDM2A, RAD18, SNRPA, DDX31, XRN1, SMARCA4, SETD1A, CLU, TRRAP, HNRNPM, DDX47, DDX46, MEIS2, CNOT6L, SMARCB1, RPL5, DNAJA3, HELLS, HIST2H3A, TRIP4, WDR5, EHMT2, FOXP1, SAFB2, HNRNPH3, PPIH, HDAC3, PSMC3, SMARCC2, DNMT1, WDR3, RPL37A, TMPO, SMC1A, PUF60, HP1BP3, ARID4B, NUP188, DKC1, PLRG1, PCBP1, CDK12, NDUFS3, MDN1, AHNAK, ELMSAN1, RING1, PRPF3, UBE2I, CDC5L, PRPF4, MCM4, PURA, CD3EAP, MRPS9, C1QBP, RIF1, NCOA5, LARP7, CPSF7, CPSF4, CPSF3, DPF2, BTAF1, NUP98, ING2, TRA2B, NFYA, PAXBP1, IARS, MINA, DIMT1, SAFB, GTF3C2, GAPDH, ZBTB7B, TCP1, PDCD11, NDUFA9, ALYREF, CENPF, TAB1, FBL, ATXN2, CUL4A, NGDN, SETD2 |
| GO:0016070  RNA metabolic process  (210) | 2.50  (9.32E-52) | RPL15, MED23, CMTR1, MED21, CTNNB1, INTS9, MED28, RAVER2, MED26, ILK, DHX34, U2AF1, FAU, LSM3, OGT, DHX30, CCAR2, CHTOP, BYSL, PTBP1, MECP2, HMG20B, MED13, MAPK1, PCF11, NOP2, BAZ1A, MED15, RFC1, PTRF, BAZ1B, TGFB1I1, SCAF8, SUDS3, CPSF3L, FIP1L1, SRSF12, RPS28, LEO1, PBRM1, NAT10, PRPF40A, RPS24, CCDC86, NOC4L, SMAD5, CREBBP, SMAD2, SRSF3, SRSF7, WDR61, TRPS1, SUPT16H, DHX40, UTP20, NCOR1, SPIN1, ZEB1, VWA9, ASH2L, SYMPK, HIST1H1E, EXOSC4, EXOSC2, GTF2H4, GTF2H3, MBD2, SMN2, SRPK1, GTF2H1, SENP3, MED9, CDK11B, NHP2, THOC3, PRPF38B, POLR2E, MTDH, ASUN, FAM98B, SKIV2L, CIC, POLR2B, SQSTM1, NUP210, GATAD2A, MLLT3, UPF1, ZNF24, ELAVL1, ILF3, ETF1, HNRNPDL, BRMS1, ILF2, HNRNPUL1, POLDIP3, PBX1, TCEB1, PES1, RBM14, TEX10, DNM2, CNOT9, RNMT, ZC3HAV1, SNRPD2, INO80, RBM7, RPS27L, CSNK2A2, DDX27, IMP3, CDKN2A, RAD21, SMARCD2, SMARCD3, CGGBP1, TARDBP, CLP1, DDX24, DNAJC8, SRRM1, DDX20, RBM10, PABPN1, ACTN4, RRP9, HNRNPR, SUZ12, RSL1D1, TRIM33, KDM2A, SNRPA, DDX31, XRN1, SMARCA4, SETD1A, CLU, TRRAP, HNRNPM, DDX47, DDX46, MEIS2, CNOT6L, SMARCB1, RPL5, DNAJA3, HELLS, HIST2H3A, TRIP4, WDR5, EHMT2, FOXP1, SAFB2, HNRNPH3, PPIH, HDAC3, PSMC3, SMARCC2, DNMT1, WDR3, RPL37A, TMPO, PUF60, HP1BP3, ARID4B, NUP188, DKC1, PLRG1, PCBP1, CDK12, MDN1, AHNAK, ELMSAN1, RING1, UBE2I, PRPF3, CDC5L, PRPF4, PURA, CD3EAP, MRPS9, C1QBP, NCOA5, LARP7, CPSF7, CPSF4, CPSF3, DPF2, BTAF1, NUP98, ING2, TRA2B, PAXBP1, NFYA, IARS, DIMT1, MINA, SAFB, GTF3C2, ZBTB7B, PDCD11, ALYREF, CENPF, TAB1, FBL, ATXN2, NGDN, SETD2 |
| GO:0046483  Heterocycle metabolic process  (233) | 2.18  (3.68E-50) | RPL15, MED23, CMTR1, MED21, CTNNB1, INTS9, MED28, RAVER2, MED26, ILK, DHX34, U2AF1, FAU, LSM3, OGT, DHX30, CCAR2, CHTOP, BYSL, PTBP1, MECP2, HMG20B, MED13, RFC5, MAPK1, PCF11, NOP2, BAZ1A, MED15, RFC1, PTRF, BAZ1B, HUWE1, TGFB1I1, SCAF8, SUDS3, CPSF3L, FIP1L1, SRSF12, RPS28, LEO1, PBRM1, NAT10, RPS24, PRPF40A, CCDC86, NOC4L, MKI67, INIP, SMAD5, CREBBP, SMAD2, SRSF3, CCT5, CCT4, SRSF7, WDR61, TRPS1, CCT8, SUPT16H, DHX40, UTP20, NCOR1, SPIN1, ZEB1, ANKRD17, VWA9, ASH2L, FANCI, SYMPK, HIST1H1E, EXOSC4, EXOSC2, GTF2H4, GTF2H3, MBD2, SMN2, SRPK1, GTF2H1, SENP3, MED9, CDK11B, NHP2, THOC3, PRPF38B, POLR2E, MTDH, ASUN, FAM98B, SKIV2L, CIC, POLR2B, SQSTM1, NUP210, GATAD2A, TRIP12, MLLT3, UPF1, ZNF24, ELAVL1, ILF3, ETF1, HNRNPDL, BRMS1, ILF2, HNRNPUL1, POLDIP3, PBX1, TCEB1, PES1, RBM14, TEX10, CALM2, DNM2, CNOT9, RNMT, ZC3HAV1, INO80, SNRPD2, RBM7, RPS27L, CSNK2A2, DDX27, TOP1, IMP3, RAD21, CDKN2A, SMARCD2, SMARCD3, CGGBP1, TARDBP, CLP1, DDX24, DNAJC8, ORC5, SRRM1, DDX20, RBM10, ORC3, PABPN1, ACTN4, RRP9, HNRNPR, RSL1D1, SUZ12, TRIM33, KDM2A, RAD18, SNRPA, DDX31, XRN1, SMARCA4, SETD1A, CLU, TRRAP, HNRNPM, DDX47, DDX46, MEIS2, CNOT6L, SMARCB1, RPL5, DNAJA3, HELLS, HIST2H3A, TRIP4, WDR5, EHMT2, FOXP1, SAFB2, HNRNPH3, PPIH, HDAC3, PSMC3, SMARCC2, DNMT1, WDR3, RPL37A, TMPO, SMC1A, PUF60, HP1BP3, ARID4B, NUP188, DKC1, PLRG1, PCBP1, CDK12, NDUFS3, MDN1, AHNAK, ELMSAN1, RING1, PRPF3, UBE2I, CDC5L, PRPF4, MCM4, PURA, CD3EAP, MRPS9, C1QBP, RIF1, NCOA5, LARP7, CPSF7, CPSF4, CPSF3, DPF2, BTAF1, NUP98, ING2, TRA2B, NFYA, PAXBP1, IARS, MINA, DIMT1, SAFB, GTF3C2, GAPDH, ZBTB7B, TCP1, PDCD11, NDUFA9, ALYREF, CENPF, TAB1, FBL, ATXN2, CUL4A, NGDN, SETD2 |
| GO:0006725  Cellular aromatic compound metabolic process  (233) | 2.16  (1.71E-49) | RPL15, MED23, CMTR1, MED21, CTNNB1, INTS9, MED28, RAVER2, MED26, ILK, DHX34, U2AF1, FAU, LSM3, OGT, DHX30, CCAR2, CHTOP, BYSL, PTBP1, MECP2, HMG20B, MED13, RFC5, MAPK1, PCF11, NOP2, BAZ1A, MED15, RFC1, PTRF, BAZ1B, HUWE1, TGFB1I1, SCAF8, SUDS3, CPSF3L, FIP1L1, SRSF12, RPS28, LEO1, PBRM1, NAT10, RPS24, PRPF40A, CCDC86, NOC4L, MKI67, INIP, SMAD5, CREBBP, SMAD2, SRSF3, CCT5, CCT4, SRSF7, WDR61, TRPS1, CCT8, SUPT16H, DHX40, UTP20, NCOR1, SPIN1, ZEB1, ANKRD17, VWA9, ASH2L, FANCI, SYMPK, HIST1H1E, EXOSC4, EXOSC2, GTF2H4, GTF2H3, MBD2, SMN2, SRPK1, GTF2H1, SENP3, MED9, CDK11B, NHP2, THOC3, PRPF38B, POLR2E, MTDH, ASUN, FAM98B, SKIV2L, CIC, POLR2B, SQSTM1, NUP210, GATAD2A, TRIP12, MLLT3, UPF1, ZNF24, ELAVL1, ILF3, ETF1, HNRNPDL, BRMS1, ILF2, HNRNPUL1, POLDIP3, PBX1, TCEB1, PES1, RBM14, TEX10, CALM2, DNM2, CNOT9, RNMT, ZC3HAV1, INO80, SNRPD2, RBM7, RPS27L, CSNK2A2, DDX27, TOP1, IMP3, RAD21, CDKN2A, SMARCD2, SMARCD3, CGGBP1, TARDBP, CLP1, DDX24, DNAJC8, ORC5, SRRM1, DDX20, RBM10, ORC3, PABPN1, ACTN4, RRP9, HNRNPR, RSL1D1, SUZ12, TRIM33, KDM2A, RAD18, SNRPA, DDX31, XRN1, SMARCA4, SETD1A, CLU, TRRAP, HNRNPM, DDX47, DDX46, MEIS2, CNOT6L, SMARCB1, RPL5, DNAJA3, HELLS, HIST2H3A, TRIP4, WDR5, EHMT2, FOXP1, SAFB2, HNRNPH3, PPIH, HDAC3, PSMC3, SMARCC2, DNMT1, WDR3, RPL37A, TMPO, SMC1A, PUF60, HP1BP3, ARID4B, NUP188, DKC1, PLRG1, PCBP1, CDK12, NDUFS3, MDN1, AHNAK, ELMSAN1, RING1, PRPF3, UBE2I, CDC5L, PRPF4, MCM4, PURA, CD3EAP, MRPS9, C1QBP, RIF1, NCOA5, LARP7, CPSF7, CPSF4, CPSF3, DPF2, BTAF1, NUP98, ING2, TRA2B, NFYA, PAXBP1, IARS, MINA, DIMT1, SAFB, GTF3C2, GAPDH, ZBTB7B, TCP1, PDCD11, NDUFA9, ALYREF, CENPF, TAB1, FBL, ATXN2, CUL4A, NGDN, SETD2 |
| GO:1901360  Organic cyclic compound metabolic process  (234) | 2.11  (1.14E-47) | RPL15, MED23, CMTR1, MED21, CTNNB1, INTS9, MED28, RAVER2, MED26, ILK, DHX34, U2AF1, FAU, LSM3, OGT, DHX30, CCAR2, CHTOP, BYSL, PTBP1, MECP2, HMG20B, MED13, RFC5, MAPK1, PCF11, NOP2, BAZ1A, MED15, RFC1, PTRF, BAZ1B, HUWE1, TGFB1I1, SCAF8, SUDS3, CPSF3L, FIP1L1, SRSF12, RPS28, LEO1, PBRM1, NAT10, RPS24, PRPF40A, CCDC86, NOC4L, MKI67, INIP, SMAD5, CREBBP, SMAD2, SRSF3, CCT5, CCT4, SRSF7, WDR61, TRPS1, CCT8, SUPT16H, DHX40, UTP20, NCOR1, SPIN1, ZEB1, ANKRD17, VWA9, ASH2L, FANCI, SYMPK, HIST1H1E, EXOSC4, EXOSC2, GTF2H4, GTF2H3, MBD2, SMN2, SRPK1, GTF2H1, SENP3, MED9, CDK11B, NHP2, THOC3, PRPF38B, POLR2E, MTDH, ASUN, FAM98B, SKIV2L, CIC, POLR2B, SQSTM1, NUP210, GATAD2A, TRIP12, MLLT3, UPF1, ZNF24, ELAVL1, ACLY, ILF3, ETF1, HNRNPDL, BRMS1, ILF2, HNRNPUL1, POLDIP3, PBX1, TCEB1, PES1, RBM14, TEX10, CALM2, DNM2, CNOT9, RNMT, ZC3HAV1, INO80, SNRPD2, RBM7, RPS27L, CSNK2A2, DDX27, TOP1, IMP3, RAD21, CDKN2A, SMARCD2, SMARCD3, CGGBP1, TARDBP, CLP1, DDX24, DNAJC8, ORC5, SRRM1, DDX20, RBM10, ORC3, PABPN1, ACTN4, RRP9, HNRNPR, RSL1D1, SUZ12, TRIM33, KDM2A, RAD18, SNRPA, DDX31, XRN1, SMARCA4, SETD1A, CLU, TRRAP, HNRNPM, DDX47, DDX46, MEIS2, CNOT6L, SMARCB1, RPL5, DNAJA3, HELLS, HIST2H3A, TRIP4, WDR5, EHMT2, FOXP1, SAFB2, HNRNPH3, PPIH, HDAC3, PSMC3, SMARCC2, DNMT1, WDR3, RPL37A, TMPO, SMC1A, PUF60, HP1BP3, ARID4B, NUP188, DKC1, PLRG1, PCBP1, CDK12, NDUFS3, MDN1, AHNAK, ELMSAN1, RING1, PRPF3, UBE2I, CDC5L, PRPF4, MCM4, PURA, CD3EAP, MRPS9, C1QBP, RIF1, NCOA5, LARP7, CPSF7, CPSF4, CPSF3, DPF2, BTAF1, NUP98, ING2, TRA2B, NFYA, PAXBP1, IARS, MINA, DIMT1, SAFB, GTF3C2, GAPDH, ZBTB7B, TCP1, PDCD11, NDUFA9, ALYREF, CENPF, TAB1, FBL, ATXN2, CUL4A, NGDN, SETD2 |
