## Supplementary material for "Comparative analysis of retroviral Gag-host cell interactions: focus on the nuclear interactome": Table S5

| **Table S5.**  Top 10 DAVID biological processes of nuclear proteins identified in Engeland *et al* 2011. | | |
| --- | --- | --- |
| **Gene Ontology Term (protein count)** | **Fold Enrichment (p-value)** | **Gene names for proteins isolated** |
| GO:0006396  RNA processing  (14) | 10.8  (2.20E-11) | GTPBP4, EXOSC5, PRPF3, CDC5L, PRPF4, SRPK1, SRSF3, EXOSC10, SRSF2, TRMT1L, SRSF7, SRSF9, POP1, NAT10 |
| GO:0010467  Gene expression  (22) | 2.9  (3.75E-09) | ABCF1, GTPBP4, MTDH, EXOSC5, PRPF3, CDC5L, PRPF4, SRPK1, LARP1, SRSF3, EXOSC10, SRSF2, TOP1, TROVE2, TRMT1L, SRSF7, SRSF9, POP1, LARS, NAT10, RBM14, MYBBP1A |
| GO:0016071  mRNA metabolic process  (11) | 11.5  (7.03E-09) | SRSF3, EXOSC10, SRSF2, ZC3HAV1, SRSF7, SRSF9, EXOSC5, PRPF3, CDC5L, PRPF4, SRPK1 |
| GO:0034641  Cellular nitrogen compound metabolic process  (23) | 2.5  (1.23E-08) | ABCF1, GTPBP4, MTDH, ZC3HAV1, EXOSC5, PRPF3, CDC5L, PRPF4, SRPK1, LARP1, SRSF3, EXOSC10, SRSF2, TOP1, TROVE2, TRMT1L, SRSF7, SRSF9, POP1, LARS, NAT10, RBM14, MYBBP1A |
| GO:0090304  Nucleic acid metabolic process  (21) | 2.8  (3.43E-08) | GTPBP4, MTDH, ZC3HAV1, EXOSC5, PRPF3, CDC5L, PRPF4, SRPK1, SRSF3, EXOSC10, SRSF2, TOP1, TROVE2, TRMT1L, SRSF7, SRSF9, POP1, LARS, NAT10, RBM14, MYBBP1A |
| GO:0006807  Nitrogen compound metabolic process  (23) | 2.3  (5.05E-08) | ABCF1, GTPBP4, MTDH, ZC3HAV1, EXOSC5, PRPF3, CDC5L, PRPF4, SRPK1, LARP1, SRSF3, EXOSC10, SRSF2, TOP1, TROVE2, TRMT1L, SRSF7, SRSF9, POP1, LARS, NAT10, RBM14, MYBBP1A |
| GO:0016070  RNA metabolic process  (20) | 3.0  (6.45E-08) | GTPBP4, MTDH, ZC3HAV1, EXOSC5, PRPF3, CDC5L, PRPF4, SRPK1, SRSF3, EXOSC10, SRSF2, TROVE2, TRMT1L, SRSF7, SRSF9, POP1, LARS, NAT10, RBM14, MYBBP1A |
| GO:0006139  Nucleobase-containing compound metabolic process  (21) | 2.5  (3.02E-07) | GTPBP4, MTDH, ZC3HAV1, EXOSC5, PRPF3, CDC5L, PRPF4, SRPK1, SRSF3, EXOSC10, SRSF2, TOP1, TROVE2, TRMT1L, SRSF7, SRSF9, POP1, LARS, NAT10, RBM14, MYBBP1A |
| GO:0046483  Heterocycle metabolic process  (21) | 2.5  (4.48E-07) | GTPBP4, MTDH, ZC3HAV1, EXOSC5, PRPF3, CDC5L, PRPF4, SRPK1, SRSF3, EXOSC10, SRSF2, TOP1, TROVE2, TRMT1L, SRSF7, SRSF9, POP1, LARS, NAT10, RBM14, MYBBP1A |
| GO:0006725  Cellular aromatic compound metabolic process  (21) | 2.5  (5.17E-07) | GTPBP4, MTDH, ZC3HAV1, EXOSC5, PRPF3, CDC5L, PRPF4, SRPK1, SRSF3, EXOSC10, SRSF2, TOP1, TROVE2, TRMT1L, SRSF7, SRSF9, POP1, LARS, NAT10, RBM14, MYBBP1A |
