## Supplementary material for "Comparative analysis of retroviral Gag-host cell interactions: focus on the nuclear interactome": Table S7

| **Table S7.**  Top 10 DAVID biological processes of nuclear proteins identified in Jäger *et al* 2012. | | |
| --- | --- | --- |
| **Gene Ontology Term (protein count)** | **Fold Enrichment (p-value)** | **Gene names for proteins isolated** |
| GO:0006396  RNA processing  (119) | 12.2  (5.22E-106) | RPL18, RPL17, RPL19, RPL13, RBM4, RPL15, SYNCRIP, NONO, DDX27, DDX17, DHX37, SRRM2, RPLP0, DDX21, RPL11, RPL12, RBM10, DHX30, RPS27A, SNRPA1, CHTOP, RRP1, HNRNPA2B1, HNRNPR, HNRNPU, MRTO4, RPS18, RPS19, NOP2, RPS16, RPS17, RPS14, RPS12, SNRPB, RPS13, SNRPA, RPS10, RPS11, PABPC4, LIN28B, RPS25, HNRNPA3, HNRNPL, RPS26, HNRNPM, DDX47, RPS28, HNRNPK, RPL7, RPL6, RPL9, HNRNPF, RPL8, HNRNPD, RPL3, RPL5, RPS20, RPL10A, PABPC1, RPL7A, RPL4, RPS23, RPS24, RPS9, RPL23A, RPS6, DDX5, RPS5, HNRNPA1, RPF2, RPS8, HNRNPA0, RPS7, SRSF3, SRSF2, HNRNPH3, SRSF7, SRSF6, WDR3, RRS1, RPL37A, HNRNPH1, ADAR, DICER1, RPS2, YBX1, RPS3, WDR36, PLRG1, RRP1B, RPS3A, SRPK2, EXOSC5, PRPF3, CDC5L, RPS4X, PRPF4, SRPK1, KHSRP, RPL35, RPS15A, RPL36, BMS1, SF3B3, DIMT1, RPL30, NSUN2, PDCD11, ALYREF, PNO1, ELAVL1, RPL27, PWP2, FBL, RPL23, RPL13A, HNRNPUL1, RPL21, SFPQ |
| GO:0016071  mRNA metabolic process  (102) | 14.2  (2.28E-94) | RPL18, RPL17, RPL19, ZC3HAV1, RPL13, RBM4, RPL15, SYNCRIP, NONO, RPLP0, SRRM2, RPL11, RPL12, RBM10, RPS27A, SNRPA1, CHTOP, HNRNPA2B1, HNRNPR, HNRNPU, MRTO4, RPS18, RPS19, RPS16, RPS17, RPS14, RPS12, SNRPB, RPS13, SNRPA, RPS10, RPS11, XRN1, HNRNPA3, HNRNPL, RPS25, HNRNPM, RPS26, DDX47, HNRNPK, RPS28, RPL7, RPL6, RPL9, HNRNPF, RPL8, HNRNPD, RPL3, RPL5, RPL7A, RPL10A, PABPC1, RPL4, RPS20, RPS23, RPS24, RPS9, RPL23A, DDX5, RPS6, RPS5, HNRNPA1, RPS8, HNRNPA0, RPS7, SRSF3, SRSF2, HNRNPH3, SRSF7, SRSF6, RPL37A, HNRNPH1, ADAR, RPS2, YBX1, RPS3, PLRG1, RPS3A, SRPK2, EXOSC5, PRPF3, CDC5L, RPS4X, PRPF4, SRPK1, EIF4G1, KHSRP, RPL35, RPS15A, RPL36, SF3B3, RPL30, PDCD11, UPF1, ALYREF, ELAVL1, RPL27, RPL23, HNRNPUL1, RPL13A, RPL21, SFPQ |
| GO:0042254  Ribosome biogenesis  (78) | 21.7  (1.83E-84) | RPL18, RPL17, RPL19, RPL13, SURF6, RPL15, RPS2, RPS3, DDX27, WDR36, RRP1B, DHX37, RPS3A, RPLP0, DDX21, RPL11, RPL12, GNL2, DHX30, RPS27A, GNL3, RRP1, EXOSC5, RPS4X, MRTO4, RPS18, RPS19, NOP2, RPS16, RPS17, RPS14, RPS12, RPS13, RPS10, RPS11, RPL35, RPS15A, RPL36, BMS1, DIMT1, RPS25, RPS26, RPL30, DDX47, RPS28, RPL7, RPL6, RPL9, NPM1, RPL8, RPL3, RPL5, RPS20, RPL4, RSL24D1, RPL10A, RPL7A, RPS23, RPS24, PDCD11, PNO1, RPL27, RPS9, RPL23A, RPS6, RPS5, RPF2, PWP2, RPS8, FBL, RPS7, RPL23, RPL13A, NOP16, RPL21, WDR3, RRS1, RPL37A |
| GO:0022613  Ribonucleoprotein complex biogenesis  (85) | 16.5  (3.73E-82) | RPL18, RPL17, RPL19, RPL13, SURF6, DICER1, RPL15, RPS2, RPS3, DDX27, WDR36, RRP1B, DHX37, RPS3A, RPLP0, DDX21, RPL11, RPL12, GNL2, DHX30, RPS27A, GNL3, SRPK2, RRP1, EXOSC5, PRPF3, RPS4X, MRTO4, RPS18, RPS19, NOP2, RPS16, RPS17, RPS14, SNRPB, RPS12, RPS13, RPS10, RPS11, RPL35, RPS15A, RPL36, BMS1, DIMT1, RPS25, RPS26, RPL30, DDX47, RPS28, ATXN2L, RPL7, RPL6, RPL9, NPM1, RPL8, RPL3, RPL5, RPS20, RPL4, RSL24D1, RPL10A, RPL7A, RPS23, RPS24, PDCD11, PNO1, RPL27, RPS9, RPL23A, RPS6, RPS5, RPF2, PWP2, FBL, RPS8, RPS7, RPL23, RPL13A, SRSF6, NOP16, RPL21, WDR3, RRS1, RPL37A, ADAR |
| GO:0016072  rRNA metabolic process  (72) | 24.3  (3.23E-81) | RPL18, RPL17, RPL19, RPL13, RPL15, RPS2, RPS3, DDX27, WDR36, RRP1B, DHX37, RPS3A, RPLP0, DDX21, RPL11, RPL12, RPS27A, RRP1, EXOSC5, RPS4X, MRTO4, RPS18, RPS19, NOP2, RPS16, RPS17, RPS14, RPS12, RPS13, RPS10, RPS11, XRN1, RPL35, RPS15A, RPL36, BMS1, DIMT1, RPS25, RPS26, RPL30, DDX47, RPS28, RPL7, RPL6, RPL9, RPL8, RPL3, RPL5, RPS20, RPL7A, RPL10A, RPL4, RPS23, RPS24, PDCD11, PNO1, RPS9, RPL27, RPL23A, RPS6, RPS5, RPF2, PWP2, RPS8, FBL, RPS7, RPL23, RPL13A, RPL21, WDR3, RRS1, RPL37A |
| GO:0006364  rRNA processing  (71) | 24.6  (2.20E-80) | RPL18, RPL17, RPL19, RPL13, RPL15, RPS2, RPS3, DDX27, WDR36, RRP1B, DHX37, RPS3A, RPLP0, DDX21, RPL11, RPL12, RPS27A, RRP1, EXOSC5, RPS4X, MRTO4, RPS18, RPS19, NOP2, RPS16, RPS17, RPS14, RPS12, RPS13, RPS10, RPS11, RPL35, RPS15A, RPL36, BMS1, DIMT1, RPS25, RPS26, RPL30, DDX47, RPS28, RPL7, RPL6, RPL9, RPL8, RPL3, RPL5, RPS20, RPL7A, RPL10A, RPL4, RPS23, RPS24, PDCD11, PNO1, RPS9, RPL27, RPL23A, RPS6, RPS5, RPF2, PWP2, RPS8, FBL, RPS7, RPL23, RPL13A, RPL21, WDR3, RRS1, RPL37A |
| GO:0006614  SRP-dependent cotranslational protein targeting to membrane  (53) | 51.5  (2.61E-79) | RPL18, RPL17, SRP14, RPL19, RPL13, RPL15, RPS2, RPS3, RPS3A, RPLP0, RPL11, RPL12, RPS27A, RPS4X, RPS18, RPS19, RPS16, RPS17, RPS14, RPS12, RPS13, RPS10, RPS11, RPL35, RPS15A, RPL36, RPS25, RPS26, RPL30, RPS28, RPL7, RPL6, RPL9, RPL8, RPL3, RPL5, RPL7A, RPL10A, RPL4, RPS20, RPS23, RPS24, RPS9, RPL27, RPL23A, RPS6, RPS5, RPS8, RPS7, RPL23, RPL13A, RPL21, RPL37A |
| GO:0006613  Cotranslational protein targeting to membrane  (53) | 48.0  (4.24E-77) | RPL18, RPL17, SRP14, RPL19, RPL13, RPL15, RPS2, RPS3, RPS3A, RPLP0, RPL11, RPL12, RPS27A, RPS4X, RPS18, RPS19, RPS16, RPS17, RPS14, RPS12, RPS13, RPS10, RPS11, RPL35, RPS15A, RPL36, RPS25, RPS26, RPL30, RPS28, RPL7, RPL6, RPL9, RPL8, RPL3, RPL5, RPL7A, RPL10A, RPL4, RPS20, RPS23, RPS24, RPS9, RPL27, RPL23A, RPS6, RPS5, RPS8, RPS7, RPL23, RPL13A, RPL21, RPL37A |
| GO:0045047  Protein targeting to ER  (53) | 47.5  (8.41E-77) | RPL18, RPL17, SRP14, RPL19, RPL13, RPL15, RPS2, RPS3, RPS3A, RPLP0, RPL11, RPL12, RPS27A, RPS4X, RPS18, RPS19, RPS16, RPS17, RPS14, RPS12, RPS13, RPS10, RPS11, RPL35, RPS15A, RPL36, RPS25, RPS26, RPL30, RPS28, RPL7, RPL6, RPL9, RPL8, RPL3, RPL5, RPL7A, RPL10A, RPL4, RPS20, RPS23, RPS24, RPS9, RPL27, RPL23A, RPS6, RPS5, RPS8, RPS7, RPL23, RPL13A, RPL21, RPL37A |
| GO:0000184  Nuclear-transcribed mRNA catabolic process, nonsense-mediated decay  (55) | 41.7  (8.26E-76) | RPL18, RPL17, RPL19, RPL13, RPL15, RPS2, RPS3, RPS3A, RPLP0, RPL11, RPL12, RPS27A, RPS4X, EIF4G1, RPS18, RPS19, RPS16, RPS17, RPS14, RPS12, RPS13, RPS10, RPS11, RPL35, RPS15A, RPL36, RPS25, RPS26, RPL30, RPS28, RPL7, RPL6, RPL9, RPL8, RPL3, RPL5, PABPC1, RPL7A, RPL10A, RPL4, RPS20, RPS23, RPS24, UPF1, RPS9, RPL27, RPL23A, RPS6, RPS5, RPS8, RPS7, RPL23, RPL13A, RPL21, RPL37A |
