## Supplementary material for "Comparative analysis of retroviral Gag-host cell interactions: focus on the nuclear interactome": Table S8

| **Table S8.**  Top 10 DAVID biological processes of nuclear proteins identified in Ritchie *et al* 2015. | | |
| --- | --- | --- |
| **Gene Ontology Term (protein count)** | **Fold Enrichment (p-value)** | **Gene names for proteins isolated** |
| GO:0098609  Cell-cell adhesion  (10) | 8.5  (2.63E-07) | EIF4G1, ATXN2L, DDX3X, ZC3HAV1, SERBP1, CCT8, PRKDC, TMPO, FLNA, LARP1 |
| GO:0007155  Cell adhesion  (10) | 5.8  (6.43E-06) | EIF4G1, ATXN2L, DDX3X, ZC3HAV1, SERBP1, CCT8, PRKDC, TMPO, FLNA, LARP1 |
| GO:0022610  Biological adhesion  (10) | 5.8  (6.62E-06) | EIF4G1, ATXN2L, DDX3X, ZC3HAV1, SERBP1, CCT8, PRKDC, TMPO, FLNA, LARP1 |
| GO:0090304  Nucleic acid metabolic process  (13) | 2.5  (3.38E-04) | EIF4G1, HNRNPM, EEF1A1, MTDH, DDX3X, MKI67, ZC3HAV1, HIST1H1C, CCT8, PRKDC, TMPO, XRN1, FLNA |
| GO:0034641  Cellular nitrogen compound metabolic process  (14) | 2.1  (6.50E-04) | EEF1A1, MTDH, MKI67, HIST1H1C, ZC3HAV1, PRKDC, FLNA, LARP1, EIF4G1, HNRNPM, DDX3X, CCT8, TMPO, XRN1 |
| GO:0010608  Posttranscriptional regulation of gene expression  (5) | 10.5  (8.57E-04) | EIF4G1, DDX3X, SERBP1, XRN1, LARP1 |
| GO:0006139  Nucleobase-containing compound metabolic process  (13) | 2.2  (1.12E-03) | EIF4G1, HNRNPM, EEF1A1, MTDH, DDX3X, MKI67, ZC3HAV1, HIST1H1C, CCT8, PRKDC, TMPO, XRN1, FLNA |
| GO:0060255  Regulation of macromolecule metabolic process  (13) | 2.2  (1.16E-03) | EIF4G1, EEF1A1, MTDH, DDX3X, ZC3HAV1, HIST1H1C, SERBP1, CCT8, PRKDC, TMPO, XRN1, FLNA, LARP1 |
| GO:0006807  Nitrogen compound metabolic process  (14) | 2.0  (1.36E-03) | EEF1A1, MTDH, MKI67, HIST1H1C, ZC3HAV1, PRKDC, FLNA, LARP1, EIF4G1, HNRNPM, DDX3X, CCT8, TMPO, XRN1 |
| GO:0046483  Heterocycle metabolic process  (13) | 2.2  (1.39E-03) | EIF4G1, HNRNPM, EEF1A1, MTDH, DDX3X, MKI67, ZC3HAV1, HIST1H1C, CCT8, PRKDC, TMPO, XRN1, FLNA |
