## Supplementary material for "Comparative analysis of retroviral Gag-host cell interactions: focus on the nuclear interactome": Table S9

| **Table S9.**  Top 10 DAVID biological processes of nuclear proteins identified in Le Sage *et al* 2015. | | |
| --- | --- | --- |
| **Gene Ontology Term (protein count)** | **Fold Enrichment (p-value)** | **Gene names for proteins isolated** |
| GO:0006614  SRP-dependent cotranslational protein targeting to membrane  (10) | 92.1  (2.06E-16) | RPS16, RPS3A, RPL34, RPL35, RPL37A, RPL11, RPS2, RPS6, RPS5, RPS3 |
| GO:0006613  Cotranslational protein targeting to membrane  (10) | 85.8  (3.97E-16) | RPS16, RPS3A, RPL34, RPL35, RPL37A, RPL11, RPS2, RPS6, RPS5, RPS3 |
| GO:0006364  rRNA processing  (12) | 39.4  (4.15E-16) | RPS16, RPS3A, FRG1, RPL34, RPL35, RPL37A, RPL11, NAT10, RPS2, RPS6, RPS5, RPS3 |
| GO:0045047  Protein targeting to ER  (10) | 85.0  (4.35E-16) | RPS16, RPS3A, RPL34, RPL35, RPL37A, RPL11, RPS2, RPS6, RPS5, RPS3 |
| GO:0016072  rRNA metabolic process  (12) | 38.4  (5.52E-16) | RPS16, RPS3A, FRG1, RPL34, RPL35, RPL37A, RPL11, NAT10, RPS2, RPS6, RPS5, RPS3 |
| GO:0072599  Establishment of protein localization to endoplasmic reticulum  (10) | 81.8  (6.18E-16) | RPS16, RPS3A, RPL34, RPL35, RPL37A, RPL11, RPS2, RPS6, RPS5, RPS3 |
| GO:0006413  Translational initiation  (11) | 51.4  (1.03E-15) | RPS16, RPS3A, RPL34, EIF5B, RPL35, RPL37A, RPL11, RPS2, RPS6, RPS5, RPS3 |
| GO:0000184  Nuclear-transcribed mRNA catabolic process, nonsense-mediated decay  (10) | 71.9  (2.06E-15) | RPS16, RPS3A, RPL34, RPL35, RPL37A, RPL11, RPS2, RPS6, RPS5, RPS3 |
| GO:0070972  Protein localization to endoplasmic reticulum  (10) | 69.0  (2.98E-15) | RPS16, RPS3A, RPL34, RPL35, RPL37A, RPL11, RPS2, RPS6, RPS5, RPS3 |
| GO:0006396  RNA processing  (15) | 14.6  (4.32E-15) | RPL35, DDX5, RPS6, RPS2, RPS5, RPS3, DDX17, RPS16, RPS3A, FRG1, RPL34, SRRM2, NAT10, RPL11, RPL37A |
