## Supplementary material for "Comparative analysis of retroviral Gag-host cell interactions: focus on the nuclear interactome": Table S10

| **Table S10.**  Top 10 DAVID biological processes of nuclear proteins identified in Li *et al* 2016. | | |
| --- | --- | --- |
| **Gene Ontology Term (protein count)** | **Fold Enrichment (p-value)** | **Gene names for proteins isolated** |
| GO:0006614  SRP-dependent cotranslational protein targeting to membrane  (15) | 41.6  (3.40E-19) | RPL17, RPS9, RPS15A, RPS2, RPS7, RPS25, RPS18, RPS16, RPL23, RPS17, RPS3A, RPS14, RPS12, RPS13, RPS10 |
| GO:0006613  Cotranslational protein targeting to membrane  (15) | 38.8  (9.58E-19) | RPL17, RPS9, RPS15A, RPS2, RPS7, RPS25, RPS18, RPS16, RPL23, RPS17, RPS3A, RPS14, RPS12, RPS13, RPS10 |
| GO:0045047  Protein targeting to ER  (15) | 38.4  (1.10E-18) | RPL17, RPS9, RPS15A, RPS2, RPS7, RPS25, RPS18, RPS16, RPL23, RPS17, RPS3A, RPS14, RPS12, RPS13, RPS10 |
| GO:0072599  Establishment of protein localization to endoplasmic reticulum  (15) | 37.0  (1.92E-18) | RPL17, RPS9, RPS15A, RPS2, RPS7, RPS25, RPS18, RPS16, RPL23, RPS17, RPS3A, RPS14, RPS12, RPS13, RPS10 |
| GO:0000184  Nuclear-transcribed mRNA catabolic process, nonsense-mediated decay  (15) | 32.5  (1.27E-17) | RPL17, RPS9, RPS15A, RPS2, RPS7, RPS25, RPS18, RPS16, RPL23, RPS17, RPS3A, RPS14, RPS12, RPS13, RPS10 |
| GO:0070972  Protein localization to endoplasmic reticulum  (15) | 31.2  (2.25E-17) | RPL17, RPS9, RPS15A, RPS2, RPS7, RPS25, RPS18, RPS16, RPL23, RPS17, RPS3A, RPS14, RPS12, RPS13, RPS10 |
| GO:0090304  Nucleic acid metabolic process  (52) | 2.7  (5.60E-17) | XRCC5, RPL17, FOSL2, XRCC6, LEMD3, DEK, RBM7, NFKB2, RPS2, YBX1, SART1, ZNF207, INTS9, EBNA1BP2, DAB2, PLRG1, RPS3A, BRD7, HIST3H2A, LUC7L3, CTBP1, YY1, MTA2, HNRNPA2B1, PRKCB, MED6, NVL, RPS18, SENP1, RPS16, RPS17, RPS14, RPS12, RPS13, RPS10, MED1, RPS15A, RPS25, TCF20, EZR, DNAJA3, NUP58, MAFF, NACC1, S100A11, RPS9, ZNF668, RPS7, RPL23, UBTF, CCT8, RBM15 |
| GO:0019083  Viral transcription  (16) | 23.6  (9.95E-17) | RPL17, RPS9, RPS15A, RPS2, RPS7, RPS25, RPS18, RPS16, RPL23, RPS17, RPS3A, RPS14, RPS12, RPS13, RPS10, NUP58 |
| GO:0006413  Translational initiation  (16) | 22.6  (1.91E-16) | RPL17, RPS9, RPS15A, RPS2, RPS7, RPS25, RPS18, RPS16, RPL23, RPS17, RPS3A, RPS14, EIF2S2, RPS12, RPS13, RPS10 |
| GO:0019080  Viral gene expression  (16) | 22.2  (2.42E-16) | RPL17, RPS9, RPS15A, RPS2, RPS7, RPS25, RPS18, RPS16, RPL23, RPS17, RPS3A, RPS14, RPS12, RPS13, RPS10, NUP58 |
